# Early Life Adversity Produces Enduring Molecular and Functional Disruption of Developing Vagal Circuits

**DOI:** 10.64898/2026.08.31.748408

**Authors:** Sarah Panvini, Sirajan Kamara, Matthew Hong, Pat Levitt

## Abstract

Early life adversity (ELA) is a leading preventable contributor to morbidity and mortality, increasing risk for mental and physical illness later in life. However, mechanisms linking ELA to comorbid outcomes within both the brain and body remain poorly understood. We tested whether ELA disrupts functional and molecular development of vagal circuitry, a key pathway for brain-body communication, using the limited bedding and nesting (LBN) mouse model of unpredictable maternal care. We measured vagally mediated autonomic stress responses longitudinally using three noninvasive measures of heart rate variability (HRV). This is the first time the development of the vagally-mediated autonomic stress response has been measured in mice. We also performed single-nucleus RNA sequencing of the vagal medulla immediately after LBN and in adulthood, followed by spatial mapping of high-confidence differentially expressed genes. ELA altered trajectories of all HRV measures, with pronounced sex differences. ELA females showed precocious maturation followed by adult declines, whereas males initially exhibited blunted responses but recovered to control levels. Transcriptomic changes were also sex dependent, with female neurons exhibiting signatures of mitochondrial dysfunction, while males showed adaptive mitochondrial responses. Spatial mapping revealed rostro-caudal organization, localizing adaptive male responses to the rostral and intermediate vagal medulla and maladaptive female responses to the intermediate vagal medulla and loose nucleus ambiguus. These findings demonstrate enduring, sex-specific alterations in vagal circuit development centered on mitochondrial pathways.

## Introduction

Early life experiences are essential for shaping brain circuitry that underlies adult behaviors^1–5^. Windows of plasticity create opportunities for experiences to shape developmental trajectories and embed themselves within biology. This occurs across species with conserved biological mechanisms, particularly through changes in gene expression that control cell differentiation, wiring, and ultimately circuit-level function^5, 6^. These sensitive developmental periods are of particular interest for understanding lifespan disorder etiology and heterogeneity, with the goal of designing developmentally appropriate interventions to prevent maladaptive outcomes. Adverse childhood experiences (ACEs) are major factors that cause early life adversity (ELA)^1, 7, 8^. ELAs are risk factors for physical and mental health disturbances, increasing risk, in the absence of protective buffers, for a toxic stress response^8–10^. ELA is considered the leading preventable cause of morbidity and mortality in the United States, particularly related to dysregulated stress systems impacting brain, cardiac, immune, metabolic, and gastrointestinal (GI) function^8–10^.

Given the systemic impact of ELA over time, understanding the functional connections between brain and body is essential to determine mechanisms of action. One understudied candidate is the vagus nerve, a central component of the autonomic nervous system. Brainstem vagal neurons provide approximately 80% of efferent cranial parasympathetic input to visceral organs^11, 12^. The vagus nerve also provides sensory feedback from those organs to the brainstem, which then projects to higher-order forebrain areas to produce a variety of behaviors^13–22^. Components of vagal circuitry converge in the medulla, which contains the sensory nucleus of the vagus nerve, the nucleus of the solitary tract (NTS)^19, 20, 22–26^, and the motor-generating nuclei, the dorsal motor nucleus of vagus (DMV) and nucleus ambiguus (nAmb)^15, 25, 27–29^. A branch of the vagus nerve directly innervates the sinoatrial node of the heart to control beat-to-beat timing of heart rate, facilitating the noninvasive measurement of vagally mediated influence on heart rate via an electrocardiogram (ECG)^30–32^. Several heart rate variability (HRV) metrics extracted from ECG recordings provide complementary information on physiological state^33–36^. Respiratory HRV (respHRV, historically referred to as RSA or HF-HRV), the high frequency component of HRV, is a physiological phenomenon that coordinates respiratory and cardiovascular functions and integrates central and peripheral signals to which the vagus nerve is an important contributor^33–36^. RespHRV is associated with infant neurobehavioral maturation, cognitive and social-emotional growth, and stress adaptation^34, 37–50^. High respHRV reflects the adaptive capacity of the vagus nerve, in concert with other central and peripheral mechanisms, to tune parasympathetic activity to maintain healthy emotional homeostasis^34, 35, 37, 39, 40, 42, 43, 51, 52^. In contrast, low respHRV in the absence of a stressor is correlated to both mental and physical health disorders, including depression, post-traumatic stress disorder, suicidal tendencies and cardiac and GI dysfunction^53–58^, the same disorders exhibiting significantly increased risk in adults with a history of high ACES^8, 59^. Together, these findings suggest that ELA may alter the developing vagal system.

Studies of ELA in animals have focused on the forebrain to determine molecular adaptations, circuit-based dysfunction, and altered behavior. Here, we focused on a detailed characterization of the development of vagally-driven parasympathetic activity. To avoid measures of vagal activity using invasive and/or restraint techniques, which are not compatible developmentally or in the context of ELA^60–62^, we applied non-invasive methods in freely moving rodents, longitudinally across development. We then applied a well-established model of unpredictable maternal care that induces ELA in mouse pups, resulting in enduring cognitive and reward disturbances into adulthood^63–74^. To determine the underlying molecular responses to ELA in vagal circuits, we performed single-nucleus RNA sequencing (snRNAseq) and multiplex in situ hybridization to map transcriptomic changes in the sensory and motor central nuclei of the vagus nerve. These experiments provide new evidence that vagal circuits are sensitive to ELA and are likely important contributors to enduring brain-body dysfunction in response to stress.

## Methods

### 1. Mice and Animal Husbandry

Mouse procedures were approved by the Institutional Animal Care and Use Committee at Children’s Hospital Los Angeles and followed NIH guidelines. Mice were socially housed in a temperature-and humidity-controlled vivarium in Super Mouse 750 ventilated cages (#75050PC-GAW, Lab Products) with LifeSpan Rodent Enrichment modules (#75009PC, Lab Products). The vivarium light cycle was 13/11 hours and food (PicoLab Rodent Diet 20, #5053) and water were provided *ad libitum*. Cages were changed weekly and provided with 100 g bedding (P.J. Murphy Forest Products Sani-Chips, #91100, Newco Distributors) and 2.7 g of nestlet squares (#6002, Newco Distributors). C57BL/6J mice (RRID: IMSR_JAX:000664) were purchased from Jackson Laboratories between 6-8 weeks of age. Mice were acclimated for one week before initiation of harem breeding of three females with one male. Cages were monitored daily, and pregnant dams were separated from the harem. To reduce cannibalism and promote maternal care, following weaning of the initial litters, dams were bred with the same sires to produce second litters that were used for all experiments. Seventy-nine second litters were generated for these studies between February 2023 and July 2025. Fifty-three litters met the behavioral criteria described below for inclusion.

### 2. Limited Bedding and Nesting (LBN) Model of ELA

We utilized a modified version of the LBN model of ELA developed by the Baram Lab^65, 69, 73, 74^. Briefly, second litters were culled to 3 males and 3 females on postnatal day (P) 2 to reduce variability caused by litter size and composition. Dams and litters were randomly assigned to a “care as usual” (CAU) or “early life adversity” (ELA) cage on P2. CAU cages had 100g of bedding and 2.7 g of nestlet material (Supplemental Fig. 1A). ELA cages had 50 g of bedding, 1.35 g of nesting material (Supplemental Fig. 1B). To limit direct access to bedding a wire grid was placed at the bottom of the cage. The elevated hut was also removed. Pups were weighed on P2 (Supplemental Fig. 1C) and then culled and placed in appropriate cages. LBN reliably produces increased nest exits and unpredictable maternal behavior. We used behavioral monitoring and inclusion criteria for our study to ensure the ELA paradigm induced the expected behavioral changes. Litters were observed for 30 minutes between 9 am and 11 am and 30 minutes between 3 pm and 6 pm between P3-P5. Nest exits within a single day were summed, and criteria for inclusion were set to 5 or fewer nest exits for CAU and 8 or more nest exits for ELA (Supplemental Fig. 1D). Approximately two-thirds of the litters met inclusion criteria. On P9 pups were weighed (Supplemental Fig. 1E), and all litters were transferred to CAU cages and weaned into CAU cages with same sex littermates at P21.

### 3. ECGenie Clinic System

#### Establishment of ECGenie Paradigm

The ECGenie Clinic System (Mouse Specifics, Inc) includes a recording tower with an insulated electrical grid that can detect an ECG through paws if the mouse has at least one front and the opposing back paw in contact with the grid. This system has been used reliably and effectively for mice of all ages^75–78^. In piloting the system, when mice were placed atop the ECGenie tower (Supplemental Fig. 2A), they exhibited freezing behavior. Although this eliminated potential movement artifacts and made recordings easy to obtain, data capture occurred only in an anxious state, limiting the ability to measure vagal and behavioral transitions. Thus, we modified the paradigm by placing the sensor pad system in a cage similar to the home cage, but without bedding (Supplemental Fig. 2A). To keep the mouse on the sensor pad and block out light, a cardboard box was placed over the mouse on the sensor pad. This proved effective for generating ECG recordings during an initial stress period followed by acclimation.

ECG recordings at P9 and P21 were performed with the cage on a heating pad. The mice exhibited minimal movement at these ages. To avoid the stress of long maternal separation for P9 mice, the total recording time was 15 minutes: initial 5 minutes, 5 minutes of acclimation, and final 5 minutes. The same ECG recording temporal parameters were used at P21, and mice were then weaned into a new cage with their same sex littermates. In pilot experiments, P35 and P50 mice exhibited optimal stillness between 10-20 minutes; thus the following parameters were used: initial 5 minutes (stressor), 10-minute acclimation, and final 5 minutes. The longer the P120 mice were in the arena, the less they moved (Supplemental Fig. 2B). Therefore, recording parameters were as follows: initial 5 minutes (stress), 20-minute acclimation, and final 5 minutes. Independent confirmation of full acclimation is not possible using this system. It is important to emphasize, however, that mouse heart rate increased initially then decreased, whereas HRV decreased and then increased, respectively, from the first five minutes to the end of the recording period (Supplemental Fig. 2D-E), consistent with an initial period of stress followed by acclimation.

#### Analysis

Signal processing analysis was performed using the EzCG e-MOUSE Software (Mouse Specifics). R-peaks were automatically detected by the Matlab-based software and visually inspected for accuracy. Inclusion parameters were as follows: 1) high quality recordings that are a minimum of one second long, which, as rodent heart rate is very fast (∼600-900 bpm), is sufficient for mouse HRV analysis^60^; 2) recordings of equal to or less than 5% of R peak editing, representing recordings that were 95% free of noise. Time domain analyses included HRV and root mean squared of the successive differences between adjacent R-R intervals (rMSSD) and spectral frequency analyses (respHRV) were extracted from ECG traces. Previous studies used a high frequency range of 1.5-5hz across ages and mouse strains to calculate respHRV^75–78^. Respiration frequency in mice varies both with age and across strains, necessitating new frequency bands that are both age and strain-appropriate^79, 80^. We selected the following frequency bands to extract respHRV at the following ages based on studies that directly measured respiration and HRV: P9 2-5 hz, P21 1.5-5 hz, P35 1.5-5 hz, P50-P55 1.2-3 hz, and P120-P150 .71-5.5 hz^79–83^. HRV adaptability was defined as the within-animal difference in HRV, rMSSD, or respRSA during the last five minutes and the same measures during the first five minutes, reflecting the within-subjects activation in response to increasing time in the arena.

### 4. SnRNAseq

#### Preparation of single nuclear samples

We performed snRNAseq to detect transcriptomic differences between rearing conditions in males and females at the end of the ELA period (P9) and in adults (P120). Mice were anesthetized with saturated isoflurane vapors (Covetrus, cat # 11695067772) using a bell jar, decapitated, and the brain removed into a 5mL conical tube with -80°C isopentane on dry ice. After flash freezing, the brain was stored at -80°C overnight before being sectioned using a brain matrix at -20°C. Two 0.5mm thick sections spanning the rostro-caudal axis of the vagal brainstem (Bregma -6.75mm to -7.67mm^84^) were collected and tissue punches 0.1 mm in diameter taken of the DMV, NTS, and nAmb and combined for a minimum of 3.5 mg of tissue. Tissue from three sex-matched littermates were pooled to generate a single sample, consisting of a minimum of 10.5 mg of tissue. Four samples per age, sex and rearing condition were generated. Samples were stored at -80°C until processing for snRNAseq. For each sequencing batch, two samples per sex and rearing condition at each age were processed in parallel. Nuclei were isolated using the single nucleus isolation kit for neuronal tissues (BN-020, Invent Biotechnologies) according to manufacturer’s instructions, except where noted. Briefly, each sample was homogenized twice with the pestle and tubes provided, followed by several rounds of filtering and ultracentrifugation after which the nuclei were resuspended in 50ul of phosphate-buffered saline (PBS) supplemented with 5% Bovine Serum Albumin (BSA). This kit provides an option for multiple purification steps using a “Buffer B” reagent. In our hands, these optional washes resulted in the loss many nuclei without improvement in the purity of and the amount of debris in the sample. Therefore, the steps involving Buffer B were eliminated. Only samples meeting the following quality control criteria were sequenced: 1) more than 90% of nuclei were separated from the whole cell, and 2) nuclei exhibited intact membranes with limited blebbing (Invent Biotechnologies kit), visualized by Trypan Blue. Samples submitted for sequencing contained between 700-1200 nuclei per microliter, with an average concentration of 900 nuclei/ul, to output approximately 10,000 nuclei for sequencing.

#### Library preparation, sequencing, and data preprocessing

Samples were submitted to the TSRI Spatial Biology and Genomics Core at CHLA between 1-2pm, within an hour of nuclei isolation. Nuclei were sequenced using the Chromium Next GEM Automated Single Cell 3′ Library and Gel Bead Kit v3.1 (10X Genomics, Cat # 1000268) according to manufacturer’s protocol. Library preparation was performed using the Illumina Hiseq X platform (Novogene). The generated FASTQ files were processed through the complete Cell Ranger (RRID:SCR_0177344) pipeline with default settings. Resulting H5AD and gene expression matrix CSV files were used for downstream analysis in R.

#### Data Analysis

#### Quality Control

All analyses were performed using R version 4.4.3. Each sample was run through the SoupX (v1.6.2) pipeline for R integrated with Seurat (v5.5.1) to remove ambient RNA, which commonly contaminates single nuclear preparations. Briefly, both raw and filtered feature matrices for each sample were integrated and underwent PCA. The number of dimensions was decided upon visual inspection of the PCA plot. Nuclei then underwent clustering, and the amount of ambient RNA contamination was estimated per cell compared to background molecules. Two P9 samples were removed from analysis due to a high estimated contamination fraction per cell. Gene expression counts were then corrected by the SoupX pipeline by removing the estimated contamination fraction for each cell.

After correction by SoupX, each sample was run through further quality control using the Seurat (v5.5.1) pipeline for R. Seurat objects (assay version 3) were created for individual samples and the number of counts and features were visualized via scatter plot. Low-quality cells (less than 200 features) and likely doublets (more than 3500 features) were removed. We assessed the percentage of mitochondrial encoded genes expressed as a further measure of sample quality. All samples had very low average mitochondrial gene expression, and all nuclei included in our analysis contained less than 5% mitochondrial-encoded gene expression. Sex chromosome and ribosomal genes were removed. Samples were then saved as H5 Seurat Objects. Following SoupX and our Seurat quality control pipeline, all samples were combined into one Seurat object at each age. The final sample numbers were as follows: P9 CAU Female N=4, CAU Male N=3, ELA Female N=3, ELA Male N=4; P120 CAU Female N=4, CAU Male N=4, ELA Female N=4, ELA Female N=4.

#### Clustering and cell type identification

Dimensionality reduction was performed for both timepoints using principal component analysis, the top 50 of which were carried forward for transformation and normalization of individual samples. Following SCTransform, samples were integrated using Harmony (1.2.3) to control for batch effects. FindNeighbors and FindClusters were run on normalized, integrated datasets at each age with a cluster resolution of .65. Clusters were visualized using UMAP projections. Cluster markers were identified using FindAllMarkers and a Wilcoxon rank sum test. Cell types were identified in a semi-automated manner using the SingleR pipeline (2.14.1) and suggested annotations were confirmed via manual annotation.

#### Differential gene expression (DGE) and downstream analyses

At each age, DGE analysis was performed on a cluster-by-cluster basis, focusing on the neuronal clusters, comparing female CAU to female ELA or male CAU to male ELA, using the EdgeR pipeline with a glmQLFit with post-hoc likelihood ratio tests. Cell-type markers readily identified neuronal and non-neuronal cells but were less successful in identifying individual neuronal subtypes within the dorsal vagal complex (DVC) as many neuronal subtypes clustered together (Supplemental Fig. 3A-B). The broader excitatory and inhibitory neuronal subtypes, however, were identified with confidence (Supplemental Fig. 3C-D) and these categories were used for downstream analysis. Genes were considered significantly differentially expressed if they had an FDR of <.05 and a logFC of +/- .5. For pathway and gene ontology analyses, we used a combination of methods including Ingenuity Pathway Analysis (IPA) and DAVID Gene Ontology. DEGs identified by EdgeR for each cluster were uploaded into both tools. For IPA, pathways were considered significantly enriched if they had a -log(p-adjusted value) of greater than 1.2 and a nonzero z-score for directionality of the pathway. Significant enrichment for DAVID Gene Ontology analysis was pathways that had an FDR <.05.

### 5. *In situ* hybridization

#### Tissue Preparation

At P120, mice that underwent the LBN paradigm were anesthetized with isofluorene, followed by ketamine injection and transcardiac perfusion with 4% paraformaldehyde (PFA). The brain was removed and placed in 4% PFA at 4°C for 24 hours and then switched to 25% sucrose for cryoprotection for 48 hours. Following cryoprotection, brains were embedded in Optimal Cutting Temperature (OCT) compound, flash frozen in liquid nitrogen, and stored at - 80°C for a minimum of 24 hours. Brainstems were then cryosectioned coronally (20µm) at - 20°C, with sections collected on Super Frost Plus slides and left at room temperature until *in situ* hybridization. Sections of the rostral (Bregma -6.75mm to -7.19mm), intermediate (Bregma - 7.43mm to -7.55mm), and caudal (Bregma -7.83mm to -7.91mm) DVC, and loose (Bregma - 7.47mm to -7.67mm) and compact (Bregma -6.59mm to -7.43mm) nAmb, according to Paxinos and Franklin^84^. were used for analysis.

#### Dual RNAscope^TM^ ISH and IHC

We performed spatial analysis of select DEGs identified in snRNAseq using ACD’s RNAscope^TM^ technology for both RNA and protein according to manufacturer’s protocol, as described previously^85, 86^. Briefly, sections went through a graded dehydration by 50%, 75%, and 100% ethanol washes. To unmask RNA targets and improve probe penetration, the tissue underwent target retrieval at 80°C. Treatment with protease 3 for 30 minutes was performed to digest proteins in the tissue. Tissue sections then were hybridized with 12 RNA probes (T1-Tomm7, T2-Phox2b, T3-Chat, T4-Gcg, T5-Gad1, T6-Gabra6, T7-Tac2, T8-S100a8, T9-Hba-a201, T10-Hb-bbs01, T11-Dbh and T12-S100a9), in a HybEZ oven at 40°C for 1 hour. Results for probes Tomm7, Phox2b, Chat, Gad1, Gabra6, S100a8, Hb-ba201, Hb-bs01, and S100a9 are presented in this paper. Data for Gcg, Tac2, and Dbh are not reported. To amplify signals, HiPlex amplification reagents (Amp1-3) were performed for 30 mins each at 40°C. The fluorescence of the RNA probes was captured in four rounds using ACD’s fluoro hybridization solution (T1-T3, T4-T6, T7-T9, T10-T12). After each imaging session, the fluorophores were stripped using the ACD cleaving solution.

After completion of Hi-Plex imaging with 12 RNA targets, we used immunofluorescence staining of the neuronal marker HuC/D to identify vagal neurons of interest. Sections were washed, then blocked with 10% normal donkey serum in 1% BSA and 1X PBS. The samples were then incubated overnight in primary mouse anti-HuC/D antibody (catalog #A-21271, Thermo Fisher Scientific; RID:AB_221448). Sections were then washed with PBS containing 0.5% Tween five times, followed by a one-hour incubation in secondary antibody (Donkey Anti-Mouse Alexa 594, Jackson ImmunoResearch catalog # 715–586-151, RRID:AB_2340858,1:500 in Stock Solution) at room temperature. Nuclei were counterstained with DAPI and sections coverslipped with ProLong Gold.

Sections were imaged on a Stellaris 5 White Light Laser Confocal Microscope (Cellular Imaging Core at Saban Research Institute, CHLA) with a 40x water corrected objective lens at 1024 x 1024 resolution. For consistency, only the left DVC and nucleus ambiguus were imaged. Z-stacks were collected through the 20 µm section and maximum projected for analysis. The operator was blinded to experimental conditions.

#### Single-cell automated multi-plex pipeline for RNA quantification and spatial mapping

SCAMPR is a semi-automated workflow for spatial quantification of gene expression following ACD’s ISH and IHC tissue staining^85, 86^. Twelve RNA probe images were max intensified in FIJI^87, 88^ and saved for further image processing. The registration and alignment of the 12 probes along with DAPI and HuC/D channels was performed using ACD RNAscope HiPlex Image Registration software. Regions of interest (ROIs) of cells within NTS, DMV, and nAmb were generated using a combination of FIJI to outline large regions of interest (sensory and motor nuclei within the medulla) and Cellpose3^89, 90^ to segment individual cells within identified nuclei. The SCAMPR pipeline for FIJI extracts individual area fractions for each cell present within an ROI. After area fractions per cell were generated, we statistically compared summed gene expression values from our regions of interest using R^85, 86^. We first normalized area fraction values by both cell number and cell size within the ROIs, then controlled for batch effects using the ComBat Data function. We then compared the expression of each transcript in CAU vs. ELA conditions using a two-tailed t-test. Males and females were normalized and analyzed separately.

### 6. Statistical Analysis

Statistical analyses were performed using a combination of Graphpad Prism 11.0 and R v4.4.3. Sample sizes for individual experiments were calculated using power analysis assuming an error rate of 0.05 and 80% power, including sex, rearing condition, and age as variables. Means and standard deviations used for power analysis were taken from a combination of pilot experiments and similar published data. All mice were randomly assigned to experimental groups, and the experimenter was blinded to the rearing conditions for data acquisition and analysis. Mice and data points were excluded due to quality control issues (i.e., motion artifacts, cell quality, alignment issues) on a case-by-case basis, as described above. Distributions were tested for normality and variances were tested for equality between CAU and ELA. If data was not normally distributed or equality of variances was not true, the appropriate nonparametric tests were used. When applicable, we used a combination of one-way, two-way, and mixed model ANOVAs controlling for repeated measures and missing data for longitudinal experiments. Bonferroni post-hoc comparisons with p-adjusted values were used to determine individual differences. We also used a general linear model with likelihood ratio tests with false discovery rates for DEG analyses. Non-parametric tests included Wilcoxon rank-sum tests. Significance was determined using a two-tailed alpha, p-adjusted value, or FDR of 0.05. Bar graphs are displayed using the mean, with error bars representing standard deviation.

## Results

### A non-invasive paradigm can measure novelty-induced HRV responses in developing mice

It is well documented that mice experience a stress response in novel environments followed by acclimation^91^. We hypothesized that we could measure this bi-phasic physiological response developmentally. After identifying age-appropriate acclimation periods (Supplemental Fig. 2B, see methods for details), recordings were analyzed during the first 5 minutes of the stress period and the final 5 minutes of the acclimation period (Supplemental Fig. 2C). Mice displayed developmentally appropriate decreases in heart rate following the age-optimized acclimation period (Supplemental Fig. 2D). This was accompanied by increased respHRV (Supplemental Fig. 2E), together indicating an initial stress response that dissipated following acclimation. This was further confirmed upon inspection of raw ECG recordings and the time between successive R-R intervals (Supplemental Fig. 2F&G).

### Sex dependent development of environmental novelty-induced HRV, rMSSD, and respHRV

We performed a developmental study of HRV responses in CAU mice at 5 time points: P9, P21, P35, P50-P55, P120-P150. The longitudinal design did not impact the initial novelty-induced stress response, as there was consistent depression of all HRV metrics during the first five minutes of recording (Fig. 1). Two-way mixed model ANOVAs revealed main effects of age and acclimation for all HRV metrics, with rMSSD and respHRV showing statistically significant interaction effects (Fig. 1). There was an unexpected sex difference in the maturation of HRV and rMSSD responses. We considered a response “mature” when the measures in the final five minutes were significantly greater than the measures in the first five minutes. In females, this functional correlate of maturation occurred prepubertally (P21) for HRV and rMSSD (Fig. 1A-B), while respHRV did not reach maturity until adulthood (P120-150) (Fig. 1C). Males differed, reaching maturity post-pubertally (P50-55) for all metrics (Fig. 1D-F). Establishing the typical developmental trajectories of these measures provided key data to determine potential sex-specifc impacts of ELA on outcomes.

**Figure 1.**
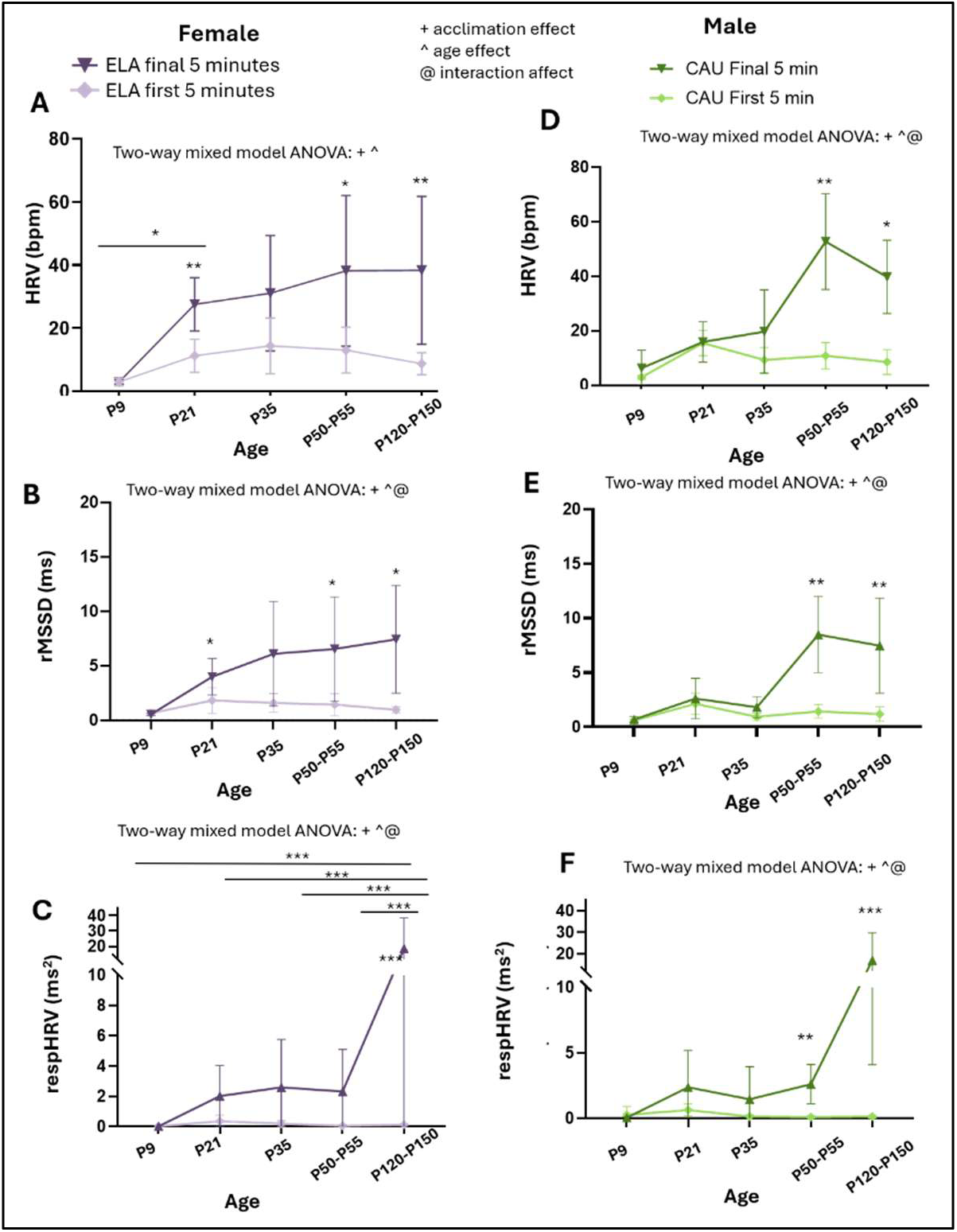
Novelty-Induced HRV Metrics in Typically Developing Mice. Two-way mixed model ANOVAs with Bonferroni post-hoc tests were used for all testing. A) Female HRV showed main effects of age (F(2.277, 25.61)= 6.375, p=0.0043), acclimation F(1,13)= 27.10, p= 0.0002), and an interaction effect (F(4,45)= 2.527, p=0.057). There were significant differences between stressed vs. acclimated HRV at P21 (t(8.122)= 4.106, padj= 0.0033), P50-55 (t(7.105)= 2.670, padj= 0.0316), and P120-150 (t(7.361)= 3.528, padj= 0.0089). There was increased acclimated HRV from P9 to P21 (t(4)= 5.925, padj= 0.210). B) Female rMSSD showed the same pattern (Age, F(2.130, 22.90)= 3,722, p= 0.0375, Acclimation F(1,12)= 27.13, p= 0.0002, Age x Acclimation F(4,43)= 22.868), p=0.0342). There were differences in stressed vs. acclimated rMSSD at P21 (t(9.032)= 2.575, padj= 0.0299), P50-55 (t(6.532)= 2.766, padj= 0.0298), and P120-150 (t(6.036)= 2.766, padj= 0.0133). C) Female respHRV showed main effects of age (F(4, 54) = 4.233), p= 0.0047), acclimation (F(1, 54) = 8.408, p=0.0054) and an interaction effect (F(4,54)= 4.248, p=0.0046). There were differences in stressed vs. acclimated respHRV at P120-150 only (t(54)= 5.070, padj <0.0001). There was increased acclimated respHRV from P9-P120 (t(54)= 4.901, padj< 0.0001), P21-P120 (t(54)= 4.372, padj= 0.0006), P35-P120 (t(54)= 4.221, padj= 0.0009), and P50-P120 (t(54)= 4.468, padj= 0.0004). D) Male HRV displayed main effects of age (F(2.526, 35.36) = 14.27, p<0.0001), acclimation (F(1, 56) = 53.20, p<0.0001), and an interaction effect (F(2.526, 35.36)= 10.87, p<0.0001). There were differences in stressed vs. acclimated HRV at P50-55 (t(4.614)= 5.142, padj= 0.0046) and P120-150 (t(8.582)= 6.227, padj= 0.0002). E) Male rMSSD displayed main effects of age (F(2.146, 20.92)= 10.75, p= 0.0005), acclimation (F(1,14)= 29.79, p<0.0001), and an interaction effect (F(2.146, 20.92)= 9.092 p= 0.0012). There were differences in stressed vs. acclimated rMSSD at P50-55 (t(4.247)= 4.436, padj= 0.0099) and P120-150 (t(7.321)= 4.026, padj= 0.0046). F) Male respHRV displayed main effects of age (F(1.126, 12.11) = 8.920, p= 0.0097), acclimation (F(1, 14) = 11.21, p= 0.0048), and an interaction effect (F(4, 43)= 9.279, p<0.0001). There were differences between stressed vs. acclimated respHRV at P50-55 (t(3.012)= 3.342, padj= 0.0441) and P120-150 (t(7.001)= 3.697, padj= 0.0077).

### ELA induces precocious maturation, followed by blunted HRV responses in female mice

Female ELA mice displayed altered developmental trajectories of HRV response. Specifically, there was a main effect of acclimation, but no main effect of age nor an interaction effect, for HRV in a two-way mixed model ANOVA (Fig. 2A). rMSSD displayed a main effect of acclimation and an interaction effect between age and acclimation, but no main effect of age (Fig. 2B). Strikingly, female ELA mice had mature HRV and rMSSD responses starting at P9, the end of the LBN period. This reflects a significant shift compared to CAU females, which did not show HRV and rMSSD maturation until P21. RespHRV in ELA females displayed significant main effects of age, acclimation, and an interaction effect (Fig. 2C), driven by a significant transient increase in acclimated respHRV at P50-55 (Fig. 2C). Other than P50-55, respHRV in ELA females failed to display mature, typical developmental patterns of acclimation.

**Figure 2.**
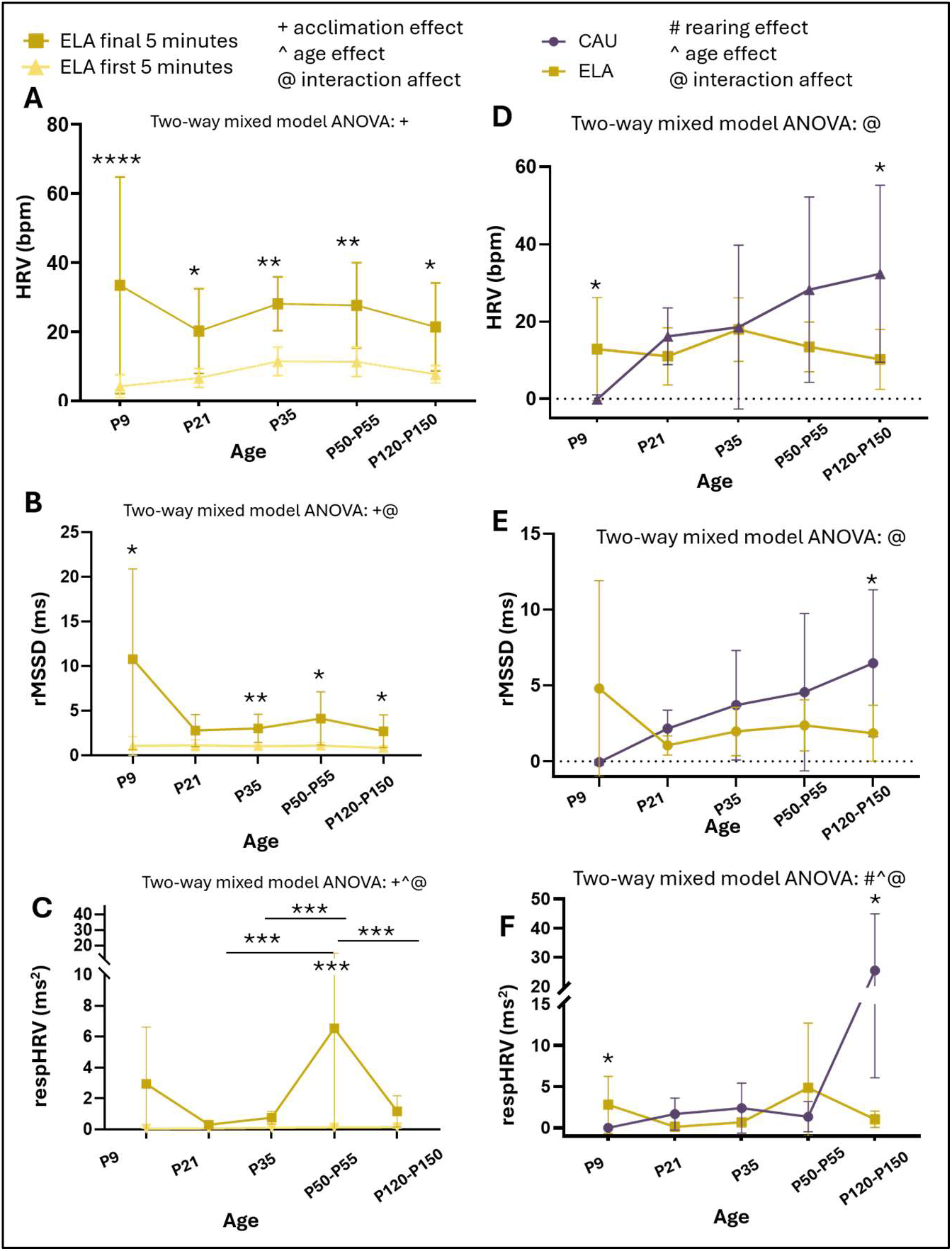
Novelty-Induced HRV Metrics in Female ELA Mice. Two-way mixed model ANOVAs with Bonferroni post-hoc tests were used for all statistical testing. A) HRV displayed a main effect of acclimation (F(1,74)= 40.41, p<0.0001). There were differences between stressed vs. acclimated HRV at P9 (t(74)= 4.516, padj< 0.0001), P21 (t(74)= 2.043, padj= 0.0447), P35 (t(74)= 2.672, padj= 0.0093), P50-55 (t(74)= 2.701, padj= 0.0086), and P120-150 (t(74)= 2.262, padj= 0.0267). B) rMSSD showed a main effect of acclimation (F(1,69)= 23.68, p <0.0001) and an interaction effect of age and acclimation (F(4,69)= 3.608, p=0.0099). There were differences in stressed vs. acclimated rMSSD at P9 (t(6.149)= 2.520, p= 0.0443), P35 (t(7.364)= 3.562, padj= 0.0085), P50-55 (t(8.214)= 3.061, padj= 0.0151), and P120-155 (t(7.180)= 2.874, padj= 0.0232). C) RespHRV displayed main effects of age (F(4,54)= 3.251, p= 0.0184), acclimation (F(1,16)= 8.431, p= 0.0104), and an interaction effect (F(4,54)= 3.209, p= 0.0195). Differences between stressed vs. acclimated respHRV occurred only at P50-55 (t(70.00)= 4.430, p< 0.0001). D) HRV adaptability showed an interaction effect between age and rearing (F(2.474, 29.69)= 3.623, p= 0.0309) and differences between CAU and ELA at P9 (t(7.145)= 2.747, padj = 0.020) and P120-150 (t(7.201)= 2.436, padj= 0.0441). E) rMSSD adaptability showed an interaction effect between age and rearing (F(2.843, 42.65)= 3.321, p= 0.0307) and differences between CAU and ELA at P120-150 only (t(7.511)= 2.374, padj= 0.047). F) respHRV adaptability showed main effects of age (F(1.535, 16.50)= 9.735, p= 0.0029) rearing (F(1, 14)= 7.132, p= 0.0183), and an interaction effect (F(1.535, 16.50)= 11.85, p= 0.0012). There were differences between CAU and ELA at P9 (t(8.001)= 2.434, padj= 0.0409) and P120-150 (t(4.013)= 2.813, p= 0.0480).

Next, we directly compared the effects of age and rearing (ELA vs. CAU) on HRV metrics (Supplemental Fig 3A-C). Two-way mixed model ANOVAs showed main effects of age, rearing, and an interaction effect for HRV (Supplemental Fig. 3A) and respHRV (Supplemental Fig. 3C). rMSSD displayed a main effect of rearing and an interaction effect of age and rearing (Supplemental Fig. 3B). Finally, we compared HRV, rMSSD, and respHRV adaptability between CAU and ELA mice (see methods). HRV and rMSSD adaptability displayed no main effects of age or rearing, but did display an interaction effect in a two-way mixed model ANOVA (Fig. 2D-E). HRV adaptability showed significant post-hoc differences between ELA and CAU at P9 and P120-150 (Fig. 2D). rMSSD adaptability displayed significant differences between ELA and CAU at P120-150 only (Fig. 2E). RespHRV adaptability showed main effects of age, rearing, and an interaction effect (Fig. 2F). Like HRV adaptability, respHRV adaptability showed significant post-hoc differences between ELA and CAU at P9 and P120. These results suggest that, in females, respHRV in particular is altered by ELA, evident just after completion of the LBN paradigm, and continuing through adulthood in female mice.

### ELA induces precocious maturation, followed by normal HRV responses in male mice

Male ELA mice also exhibited changes in the trajectories of HRV responses. Two-way mixed model ANOVAs showed a main effect of acclimation and an interaction effect of age and acclimation in HRV (Fig. 3A). rMSSD displayed main effects of age and acclimation with no interaction effects (Fig. 3B). RespHRV had main effects of age, acclimation, and an interaction effect (Fig. 3C). Post-hoc analyses revealed significant differences between recordings taken in the first five minutes versus the final five minutes at P35 and P120-150 for HRV, rMSSD, and respHRV. These results suggest that although ELA shifts developmental trajectories towards early maturation in males, the systems that drive rMSSD and respHRV recover to CAU levels in adulthood.

**Figure 3.**
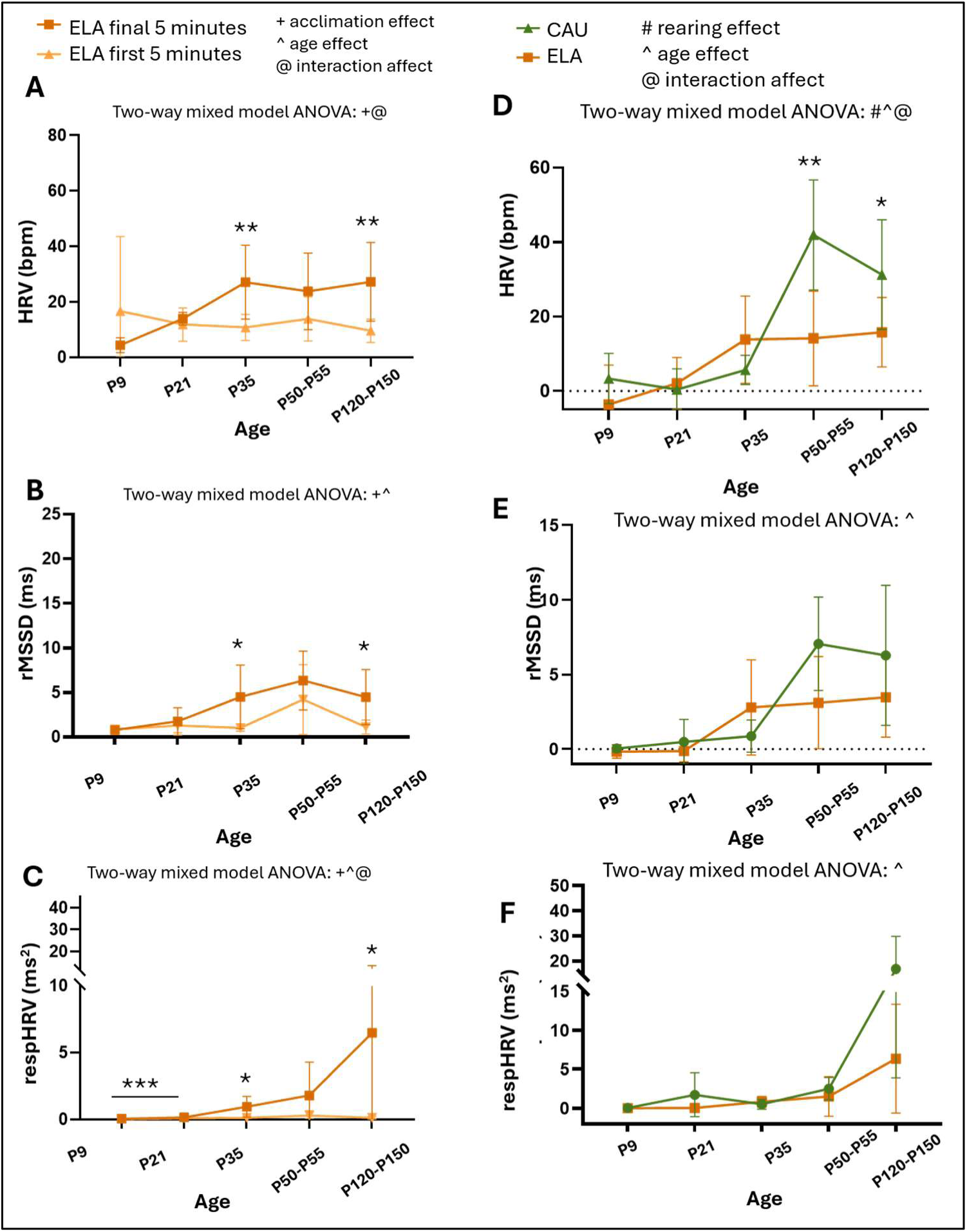
Novelty-Induced HRV Metrics in Male ELA Mice. Two-way mixed model ANOVAs with Bonferroni post-hoc tests were used for all statistical testing. A) HRV showed a main effect of acclimation (F(1,16)= 5.890, p=0.0274) and an interaction effect of age and acclimation (F(2.377, 35.06)= 4.717, p= 0.0114) in HRV. Differences between stressed vs. acclimated HRV occurred at P35 t(10.14)= 3.452, padj= 0.0061) and P120-150 t(9.533)= 3.564, padj= 0.0056). B) rMSSD displayed main effects of age (F(2.342, 42.73)= 8.563, p= 0.0004) and acclimation (F(1, 73)= 13.80, p=0.0004). Differences between stressed vs. acclimated rMSSD occurred at P35 t(8.186)= 2.912, padj= 0.0191 and P120-150 t(9.132)= 3.139, padj= 0.0117 C) respHRV had main effects of age (F (1.145, 16.03) = 5.474; p= 0.0288), acclimation (F (1, 16) = 7.527; p= 0.0144), and an interaction effect (F (1.145, 16.03) = 5.328; p= 0.0308). Differences between stressed vs. acclimated respHRV occurred at P35 t(8.597)= 3.122, padj= 0.0130 and P120-150 t(8.003)= 2.749, padj= 0.0259). D) HRV adaptability revealed main effects of age (F (2.752, 30.96) = 21.63, p< 0.0001), rearing (F (1, 16) = 9.596, p= 0.0069), and an interaction effect (F (2.752, 30.96) = 6.196, p= 0.0025). Differences between ELA and CAU occurred at P50 (t(7.883)= 3.403, padj= 0.0095) and P120P150 (t(11.84)= 2.507, padj= 0.0278). E) rMSSD adaptability had a main effect age (F (2.346, 35.77) = 13.04, p< 0.0001). There were no significant differences between ELA and CAU. F) RespHRV adaptability showed a main effect of age (F(1.150, 13.52)= 15.28, p= 0.012) and no significant differences between ELA and CAU.

Two-way mixed model ANOVAs comparing the effects of age and rearing revealed rearing, age, and interaction effects for HRV, rMSSD, and respHRV in males (Supplemental Fig. 3D-F), with no significant post-hoc comparisons. Similar tests for adaptability revealed main effects of age, rearing, and an interaction effect for HRV only (Fig. 3D). Post-hoc tests revealed significant differences between CAU and ELA HRV adaptability at P50-55 and P120-150. Interestingly, there was a main effect for age, but no main effect of rearing, interaction effects nor significant posthoc effects for rMSSD or respHRV adaptability (Fig. 3E-F). Thus, although ELA impacts the maturation of HRV, rMSSD, and respHRV in both female and male mice, males recover in adulthood and only HRV adaptability is impacted by rearing.

### snRNAseq at P9 reveals acute sex-dependent mitochondrial responses to ELA in vagal neurons

Given the functional changes in HRV responsiveness in the mild stress paradigm, we wondered whether the vagal complex exhibited sex-specific, altered molecular profiles related to the LBN model, as we have reported for the hippocampus^73, 74^. We performed snRNAseq of the vagal brainstem at the end of the LBN paradigm (P9), when functional HRV changes in female mice were observed. We sequenced ∼650,000 cells that included, based on expression of marker genes, those located in the vagal complex including the NTS, DMV, and nAmb (Supplemental Fig. 4A-B). We detected all expected cell types, including oligodendrocytes, oligodendrocyte precursor cells, endothelial cells, astrocytes, microglia, and neurons (Fig. 4A-B). Each cell type exhibited DEGs, with male mice displaying more than double the number of differentially expressed transcripts compared to females (Fig. 4C). We detected many differentially expressed nuclear-encoded mitochondrial genes. Remarkably, ELA induced strikingly different patterns of expression in these genes in females (downregulated) and males (upregulated or unchanged) (Fig. 4D-E). We next focused on DEGs identified in neurons (Fig. 4F), which in this region are directly involved in parasympathetic functions, for IPA and DAVID gene ontology analyses. We further generated excitatory and inhibitory neuron clusters based on the expression of traditional marker genes (Supplemental Figure 4D). For IPA and DAVID analyses, significant pathway enrichment only occurred in excitatory neurons (Supplemental Tables 1-4). IPA revealed downregulation of pathways involved in respiratory electron transport and oxidative phosphorylation (OXPHOS), and upregulation of pathways related to mitochondrial dysfunction in females (Supplemental Table 5). There was concomitant downregulation of the S100 family signaling pathway, with specific DEGs known to aid in cellular antioxidant response (*S100a8* & *S100a9*)^92^ (Supplemental Table 5). In contrast, male ELA mice exhibited upregulation of pathways involved in the NRF2-mediated oxidative stress response, including an extensive list of downstream antioxidants^93^ (Supplemental Table 5). Excitatory neurons also displayed sex-specific patterns of differential expression of GABAergic receptors (Supplemental Table 6), which are important for maintaining excitatory/inhibitory balance^94^. Males displayed downregulation of GABAergic receptor signaling pathways (Fig. 6), although individual receptor subunits were both increased and decreased (Supplemental Table 1). Females also displayed changes in individual receptor subunit expression, but these changes did not reach statistical significance at the pathway level. DAVID revealed significant enrichment of amidation pathways in females only, which can result from oxidative amidation (Supplemental Table 6)^95, 96^. Together, these molecular patterns provide evidence that ELA impacts neuronal mitochondria acutely in a sex-dependent manner.

**Figure 4.**
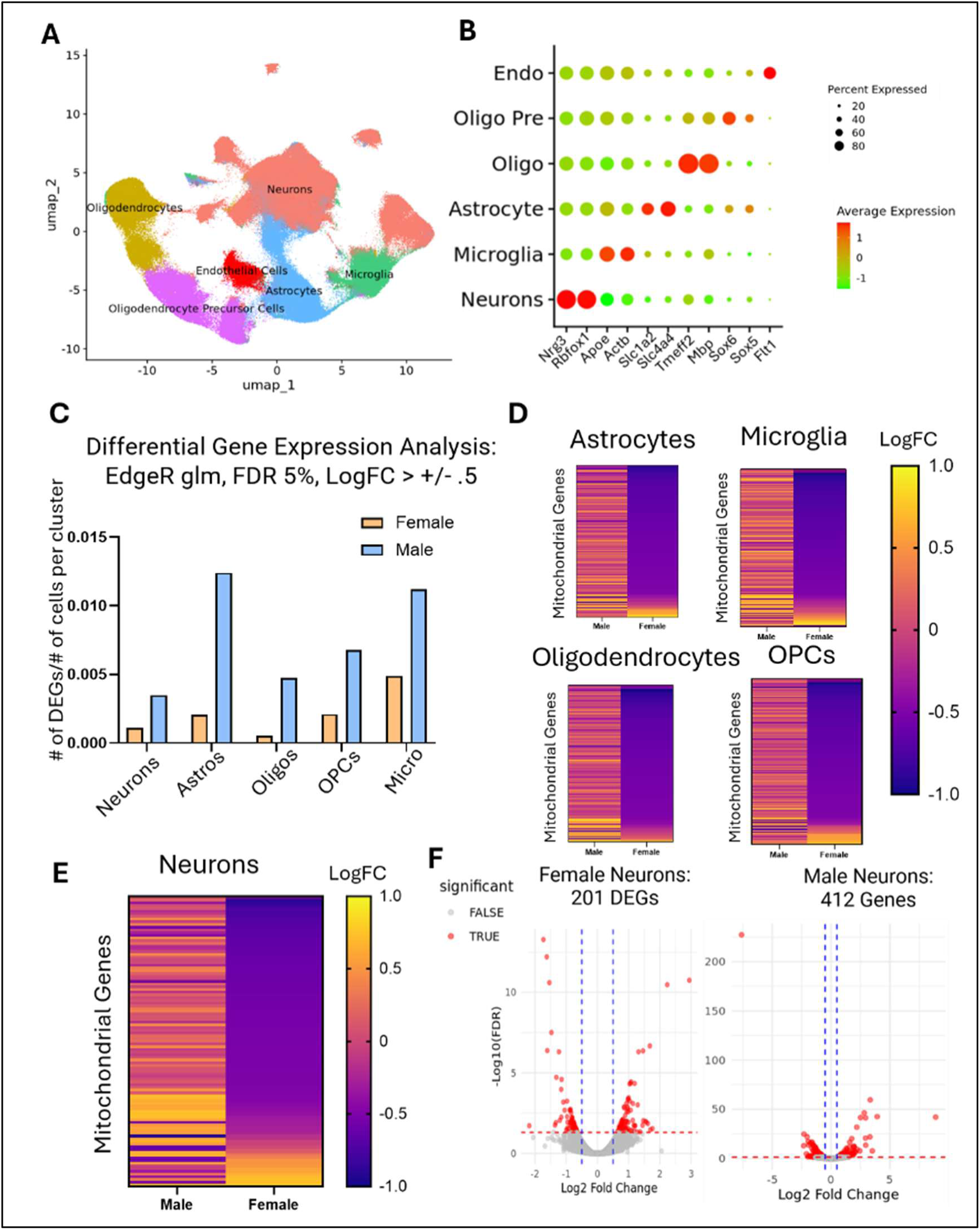
snRNAseq at PG shows robust ELA-induced differential gene expression with marked sex differences. A) UMAP of cell-type clusters present in snRNAseq samples from the medulla. B) Marker genes and expression profiles used to identify major cell types. C) Proportion of significant DEGS (FDR <0.05, logFC > +/- 0.05) normalized to the number of transcripts expressed in each cluster in males and females. D) Heat maps displaying differentially expressed nuclear-encoded mitochondrial transcripts across cell-types, with an emphasis on neurons (E). F) Volcano plots of DEGS in neurons from male and female neuronal subclusters.

**Figure 5.**
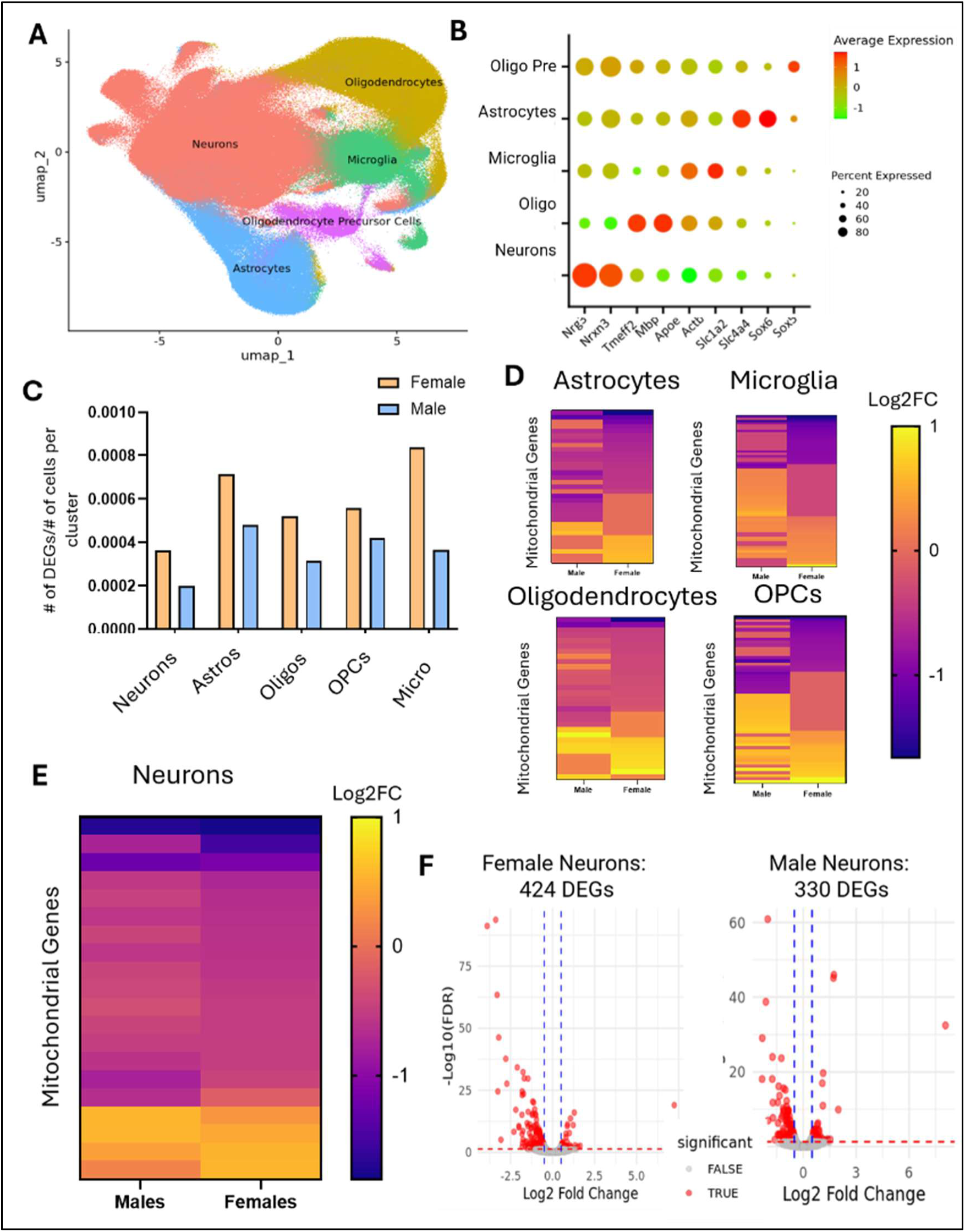
snRNAseq at P120 shows robust ELA-induced differential gene expression with marked sex differences. A) UMAP of cell-type clusters present in snRNAseq samples from the medulla. B) Marker genes and expression profiles used to identify major cell types. C) Proportion of significant DEGS (FDR <0.05, logFC > +/- 0.05) normalized to the number of transcripts expressed in each cluster in males and females. D) Heat maps displaying differentially expressed nuclear-encoded mitochondrial transcripts across cell-types, with an emphasis on neurons (E). F) Volcano plots of DEGS in neurons from male and female neuronal subclusters.

**Figure 6.**
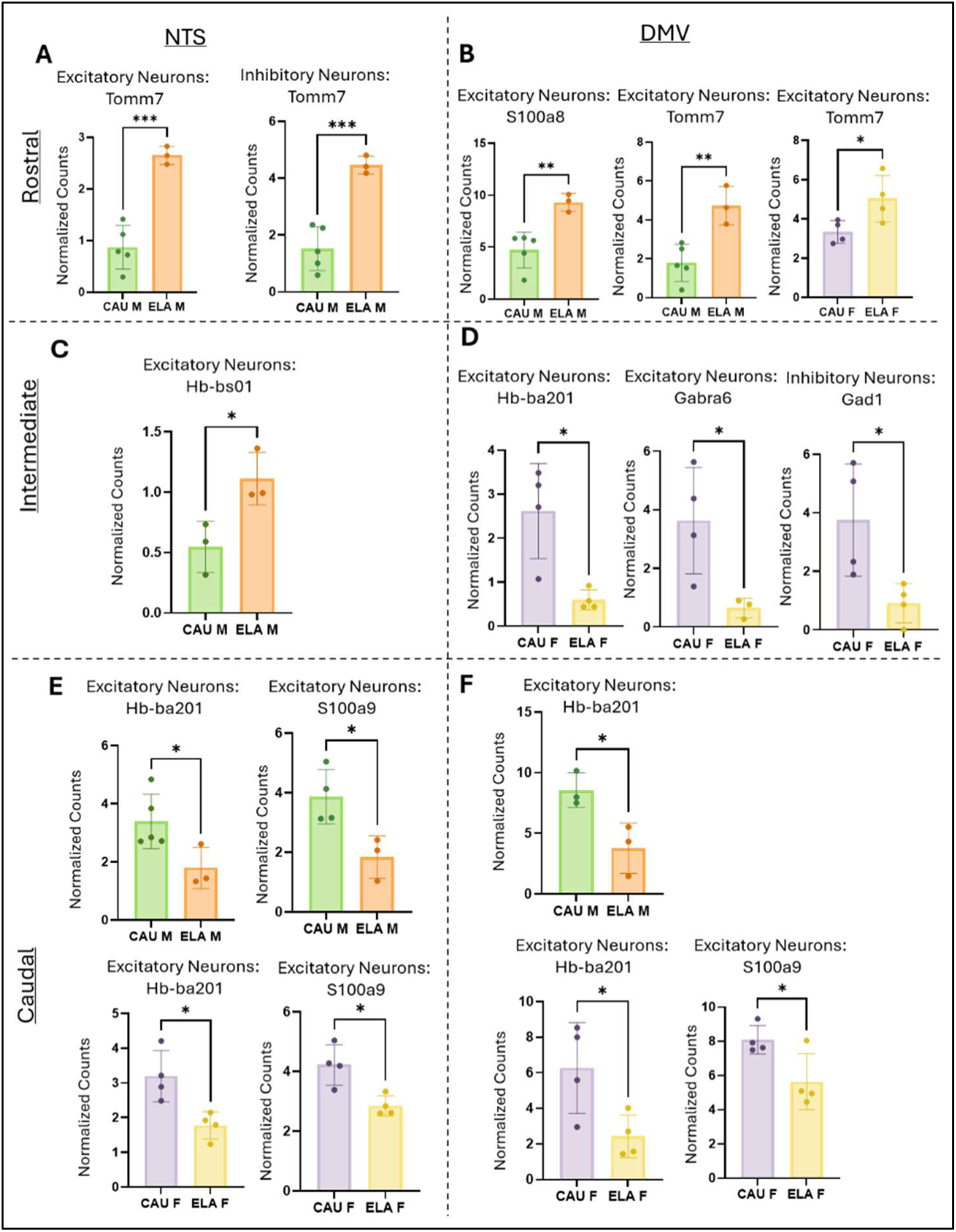
RNA expression of DEGs along the rostral-caudal axis of the vagal medulla. A) The rostral NTS displayed increases in Tomm7 in excitatory (t(6)= 6.759, p= 0.0005, Cohen’s d= 4.936) and inhibitory neurons (t(6)= 6.147, p= 0.0008, Cohen’s d= 4.489) of ELA males. B) Excitatory neurons of the rostral DMV displayed increases in S100a8 (t(6)= 4.189, p= 0.0058, Cohen’s d= 3.059) and Tomm7 (t(6)= 4.165, p= 0.0059, Cohen’s d= 3.041) in ELA males and S100a8 in ELA females (t(6)= 2.573, p= 0.0422, Cohen’s d= 1.819). C) The intermediate NTS displayed increased Hb-bs01 in excitatory neurons of ELA males (t(4)= 3.232, p= 0.0319, Cohen’s d= 2.639). D) In the intermediate DMV, ELA females displayed decreased Hb-ba201 (t(6)= 3.663, p= 0.0105, Cohen’s d= -2.590) and Gabra6 in excitatory neurons (t(5)= 2.749, p= 0.04, Cohen’s d= -2.099), and decreased Gad1 in inhibitory neurons (t(6)= 2.793, p= 0.0315, Cohen’s d= -1.975) E) Excitatory neurons in the caudal NTS downregulated Hb-ba201 in both males (t(5)= 2.731, p= 0.0412, Cohen’s d= -2.086) and females (t(4)= 3.424, p=0.0217, Cohen’s d= -2.421). Excitatory neurons also downregulated S100a9 in males (t(5)= 3.155, p=0.0252, Cohen’s d= -2.410) and females (NTS t(6)= 3.649, p= 0.0107, Cohen’s d= -2.580). F) Excitatory neurons in the DMV downregulated Hb-ba201 in both males (DMV t(4)= 3.289, p= 0.0303, Cohen’s d=-2.685) and females (DMV t(6)= 2.718, p= 0.0347, Cohen’s d= -1.922). Excitatory neurons of the caudal DMV also downregulated S100a9 in females (t(6)= 2.687, p= 0.0362, Cohen’s d= - 1.900).

### snRNAseq at P120 reveals enduring sex-dependent mitochondrial responses to ELA in vagal neurons

To identify long-term changes due to ELA, we performed snRNAseq between the ages of P120-P150, a timepoint at which significant dysfunction of HRV metrics were observed in females. We sequenced ∼828,000 cells within the vagal medulla, and all appropriate cell-types were detected (Fig. 5A-B, Supplemental Fig. 5A-B). In contrast to P9, P120 females displayed the highest number of DEGs (Fig. 5C). Similar patterns in nuclear-encoded mitochondrial genes seen at P9 were observed at P120 (Fig. 5D-E). However, there were fewer differentially expressed mitochondrial transcripts compared to P9, and the expression patterns were more similar between males and females at this timepoint (Fig. 5E). IPA and DAVID analyses of subclustered excitatory and inhibitory neurons (Supplemental Fig. 5C-D) revealed that the same patterns identified at P9 persisted into adulthood. Females displayed enrichment and upregulation of pathways contributing to mitochondrial dysfunction (Supplemental Table 8 IPA) and males showing increased pathways related to buffering oxidative stress (Supplemental Table 8 DAVID). At P120, females displayed altered GABAergic receptor signaling at the pathway level (Supplemental Table 8 IPA). Females still displayed enrichment of amidation pathways as seen at P9 (Supplemental Table 8 DAVID). These changes identified by IPA and DAVID were once again specific to excitatory neurons (Supplemental Tables 9-12).

### HiPlex RNAScope reveals region-specific, topographic differences in ELA responsive DEGs

The vagal complex is highly heterogeneous with many neuron types that are organized topographically, reflecting their distinct projections and visceral functions^97^. Further, these regions contain evident neuronal-type differences along the rostro-caudal axis of each nucleus^13, 19, 24–26, 97–104^. In our snRNAseq, we were unable to clearly distinguish between sensory and motor nuclei or rostro-caudal cell types using clustering or gene expression patterns, warranting a more detailed spatial investigation of our identified DEGs. We therefore performed Hiplex RNAScope, which utilizes *in situ* hybridization to provide spatial resolution of DEGs, at P120. We focused on high-confidence DEGs with enduring transcriptomic changes into adulthood based on snRNAseq data. We probed for marker genes for distinct nuclei (*Phox2b*, *Chat*), inhibitory interneurons (*Gad1*), and genes identified by IPA in the GABA receptor signaling pathway (*Gabra6*)^94^, oxidative stress buffering pathways (*S100a8*, *S100a9*, *Hb-ba201*, *Hb-bs01*)^92, 105–107^ and the mitochondrial dysfunction pathway (*Tomm7*)^108, 109^. Sections through the rostral (Supplemental Fig. 6A), intermediate (Supplemental Fig. 6B), and caudal (Supplemental Fig. 6C) vagal complex as well as the compact (Supplemental Fig. 6D) and loose (Supplemental Fig. 6E) nAmb were analyzed. There were striking rostro-caudal differences in gene expression patterns in CAU mice (Supplemental Fig. 6A’-E’). Consistent with this discovery, differential expression patterns in response to ELA also exhibited rostro-caudal topography. These results are summarized across regions in Supplemental Table 13.

### Rostral NTS, DMV, and Compact nAmb

Sex differences identified in snRNAseq appeared at the rostral level of the NTS (Fig. 6A) and DMV (Fig. 6B). The NTS displayed increases in *Tomm7* in excitatory and inhibitory neurons of ELA males (Fig. 6A), with no changes in females (Supplemental Fig 7A). Studies have shown that *Tomm7* aids in the mitochondrial stress response and contributes to complex I function^108, 109^. The rostral NTS comprises the first synapse of the central taste neuroaxis and projects directly to the DMV to facilitate vago-vagal reflex arcs^26, 97, 103, 104, 110^, and our data suggest that these neurons in male ELA mice have increased capacity to respond to cellular stress. Excitatory neurons in the rostral DMV upregulated *Tomm7* following ELA in both sexes (Fig. 6B). Males also displayed increases in *S100a8* (Fig. 6B) with no changes in females (Supplemental Fig. 7B). This suggests that, unlike the NTS, both male and female GI projecting motor neurons have increased capacity to respond to cellular stress in response to ELA. However, only males may able to effectively buffer oxidative stress due to upregulation of *S100a8*. There were no significant DEGs in the compact nAmb, which contains cardiopulmonary projecting motor neurons^97, 111^.

### Intermediate NTS, DMV, and Loose nAmb

The intermediate level of the NTS (Fig. 6C) and DMV (Fig 6D) also displayed sex differences detected in our snRNAseq. The intermediate NTS displayed increases in *Hb-bs01* in excitatory neurons of male mice that experienced ELA (Fig.6C), with no changes in females (Supplemental Fig. 8A). Thus, males may have increased capacity to buffer oxidative stress in sensory neurons that receive the vast majority of inputs from the GI system^26, 97, 98, 103, 110^. In the intermediate DMV, ELA females displayed decreases of *Hb-ba201* and *Gabra6* in excitatory neurons, and decreased *Gad1* expression in inhibitory neurons (Fig. 6D). Males displayed no changes in *Hb-ba201* (Supplemental Fig. 8B). Interestingly, males displayed trending decreases in *Gad1* and trending increases in *Gabra6* (Supplemental Fig. 8C-D), potentially representing a compensatory mechanism. Most motor neurons within the intermediate DMV project to the GI system; however, this level also is the only location in the DMV containing cardiac-projecting neurons that control ventricular contractility^112, 113^. The loose nAmb, comprising cardiovascular projecting neurons that are part of the baroreceptor reflex^111^, displayed increases in *Gabra6* that reached significance only in females (Supplemental Fig. 9A).

### Caudal NTS and DMV

Interestingly, the caudal portion of the vagal medulla displayed no sex differences in DGE in response to ELA. Excitatory neurons in the caudal NTS (Fig. 6E) and DMV (Fig. 6F) downregulated *Hb-ba201* in both sexes and *S100a9* in females (Fig. 6E). Males also downregulated *S100a9* in the NTS (Fig. 6E) but this change was trending in the DMV (Supplemental Fig. 10A). The caudal NTS contains sensory neurons that receive inputs from the GI, cardiovascular, and pulmonary systems^26, 97, 98, 103, 110^. The caudal DMV contains motor neurons that mainly promote gastric relaxation^24, 98^. These data suggest that sensory and motor neurons in the caudal vagal complex are vulnerable to oxidative stress following ELA in both sexes.

## Discussion

The current study provides a new circuit and molecular perspective on the impact of ELA across the lifespan on functions directly related to autonomic responses to developmental stress. ELA studies in rodents have focused mostly on forebrain structures, but not on direct mediators of brain-body interactions. Here, we undertook functional and molecular analyses of the vagal system components that directly engage peripheral functioning. Two complementary methods were applied over developmental time to determine whether early changes lead to enduring disturbances or resilience.

### Heart Rate Variability

We employed a noninvasive measurement of HRV and autonomic flexibility longitudinally from early postnatal development into young adulthood, to our knowledge, for the first time in mice. All HRV metrics displayed general agreement, with some differences in statistical significance for typical development and following ELA. Like humans^11, 114–117^, HRV metrics in mice increase steadily across development. Strikingly, CAU females display differences in acclimated HRV/rMSSD versus stressed HRV/rMSSD pre-pubertally (P21), whereas these differences do not emerge until post-puberty (P50-P55) in CAU males (Fig. 1A). This suggests that for acute, mild stress acclimation, the female mouse autonomic nervous system matures earlier than that of males.

ELA mice exhibited different developmental trajectories across all HRV metrics compared to CAU. Unlike CAU females, P9 ELA females already displayed increased acclimated HRV and rMSSD compared to the stressed measures, perhaps reflecting precocious maturation of the autonomic system. By P21, acclimated metrics decreased followed by a flat developmental trajectory into adulthood. Importantly, there was no impact of age on HRV or rMSSD development in ELA females, suggesting either precocious maturation or a loss of plasticity in these circuits. RespHRV showed a different pattern, with a nonsignificant increase in acclimated signal compared to the stress signal at P9, and no differences at P21 and P35. During post-pubertal adolescence (P50-P55), there was a large increase in acclimated respHRV, although the variance was high with only some mice displayed this response. However, this was not sustained, such that by adulthood, there is again no difference in respHRV between stressed and acclimated recordings, indicating respHRV is unable to return to homeostatic baseline. We propose that the physiological activity patterns in ELA females reflect an allostatic change, whereby the autonomic nervous system initially increases its activity to adapt to ELA, but eventually fails when this response cannot be sustained^118, 119^. This pattern is similar to changes in mitochondrial energetics in the hippocampus using the same LBN paradigm^73, 74^.

ELA males displayed flattened developmental curves for HRV and rMSSD compared to CAU. RespHRV, however, followed a similar trajectory to CAU. rMSSD and respHRV adaptability showed no effects of rearing, with ELA measurements not displaying statistical differences from CAU measurements at all ages. HRV adaptability did display significant disruptions due to ELA, with patterns similar to females. For all three HRV metrics, ELA males displayed earlier maturation (P35) than their CAU counterparts (P50-P55), also similar to results in females. Each HRV measure represents different components of physiological responses to stress, reflecting the heterogeneity of results in the field that use different, single measures. Here, data from two of the three HRV metrics in male mice are consistent with autonomic resilience to the lasting effects of ELA.

There is much debate regarding the interpretation of HRV metrics. Our rationale for analyzing multiple HRV measures was to probe the adaptability of the autonomic nervous system in response to a mild, time-restricted, developmental stressor that has been utilized widely^69^. HRV is calculated by the ECGenie as the variability in time between beats measures in beats per minutes, and rMSSD is the square root of the mean of the squared differences between adjacent R-R intervals measured in milliseconds^120^. HRV likely represents a broader physiological metric that may be more appropriate for longer-term recordings^60, 80^. It has been suggested that rMSSD is less influenced by respiration than other measures and is a more appropriate measure for short-term recordings on the millisecond scale, particularly in rodents which have high heart rates^60, 121–123^. Thus, measuring HRV alone in beats per minute may mask subtle short-term changes in heart rate in rodents^60, 80^. RespHRV, historically referred to as HF-HRV or RSA, used to be considered a pure measure of vagal activity^42, 124, 125^. However, neurophysiological evidence shows multiple contributors, including respiration, sympathetic tone, mechanical effects, top-down signaling from higher order brain regions, and peripheral signals in combination with a major influence of parasympathetic activity from the vagus nerve^34, 35, 37, 38, 51^. Because respHRV represents the contribution and coordination of multiple systems that work collaboratively to restore body homeostasis, it is suggested that respHRV represents a measure of allostasis^38^. Applying this concept to the current study, ELA disrupts the development of allostasis, or the body’s ability to flexibly respond to mild stressors in female mice.

### Transcriptomics

There are intriguing molecular changes in the neurons comprising central vagal brainstem circuits that align in several ways with the HRV measures and previous studies of the hippocampus^73, 74^ . First, we observed striking sex differences in responses to ELA, evident immediately after the LBN period and enduring through adulthood. DEGs were enriched in genes encoding proteins involved in mitochondrial function, oxidative stress production, oxidative stress buffering, and inhibitory neurotransmitter receptors in excitatory vagal neurons. Consistent with HRV metric changes at P9 and P120, males displayed molecular changes that reflect resiliency, whereas females displayed maladaptive molecular changes. We reported similar sex-specific changes in the hippocampus using this LBN model^73, 74^. In that study, ELA increased mitochondrial proteins for complex I-IV, OXPHOS capacity, and complex I activity in male and female juvenile mice. In adulthood, ELA females displayed reduced proteins for complex I-IV, lower OXPHOS capacity and complex I activity, with no differences between male CAU and ELA mice. In fact, there was increased expression of the antioxidant protein Sod1 in the ELA males during adulthood, suggesting sustained adaptive capacity of the mitochondria to buffer oxidative stress in the hippocampus^73^. The present transcriptome data of vagal neurons aligns with this pattern, although we did not assess a juvenile timepoint. There is a wealth of evidence in both rodent and human literature that mitochondria exhibit enduring structural and functional changes to chronic stress exposure, with reduced ability for energy production^118, 126^. Further, the human literature involving ELA, ACEs, and chronic mental and physical health conditions has a large body of evidence that mitochondrial health is impacted by accumulating oxidative stress^127–133^.

The snRNAseq data also revealed a consistent alteration of GABAA receptor subunits in excitatory neurons. The potential impact of these findings is less clear, as genes encoding GABAA receptor subunits respond to ELA differently, with some increasing and others decreasing following ELA. There is accumulating evidence that, similar to stress, perinatal high-fat diet impacts the motor neuron function within the DMV and inhibitory signaling between the NTS and DMV^134, 135^. Several studies have found changes to the excitatory/inhibitory balance and maturation of GABAA receptor subunits within DMV excitatory motor neurons during development after exposure to a perinatal high fat diet, which are then maintained into adulthood^134–137^. Our results build on this evidence that inhibitory signaling both within the DMV and between the NTS and DMV is sensitive to environmental insults early in life. We further found a significant downregulation of *Gad1* in neurons in the intermediate DMV of ELA females. We suggest that the combined downregulation of *Gad1* and *Gabra6* in these neurons may contribute to dysfunctional signaling of vagal motor neurons in ELA females. ELA males showed mixed changes in different postsynaptic GABAA receptor subunits and *Gad1*, which may be compensatory within these neurons. Direct measures of the functional state of these neurons by electrophysiology or calcium imaging will help draw clearer conclusions.

### Spatial Gene Expression

Our broad evaluation of the genome revealed sex-and age-dependent adaptations in response to ELA. The importance of additional detailed regional mapping of sequencing data cannot be overstated, as meaningful distinctions may be masked within large datasets of heterogeneous brain regions. Spatial transcriptomics is improving but is expensive and currently offers more limited gene sets compared to single-cell sequencing. Therefore, we complemented our snRNAseq analyses with a more targeted spatial approach, multiplex *in situ* hybridization.

The expression of genes selected from the snRNAseq data mapped differently along the rostro-caudal axis of the medulla in CAU mice. This paralleled the distribution of ELA-induced gene expression changes, revealing the heterogeneity in transcriptomic adaptations to ELA by neurons even within the same nucleus. Of particular interest, sex differences in ELA-induced DGE were restricted to the rostral and intermediate DMV and NTS and were transcript-specific. The intermediate regions of these nuclei comprise the largest portion of the vagal medulla and thus, are the likely drivers of the sex differences observed in the sequencing data. Further, the intermediate DMV is comprised of motor neurons that project to either the gut or directly to the left ventricle of the heart to control contractility^24, 112, 113^, providing a strong link to our functional data. The nAmb also showed DGE, including downregulation of *Gabra6* in the loose portion of the nucleus in females only. The nAmb contains several populations of cardiovascular and cardiopulmonary neurons that project directly to the sinoatrial node of the heart to control beat-to-beat timing, as well as the baroreceptor reflex and the dive reflex coordinating respiration and heart rate^111^. A recent study identified a neuronal subtype exclusive to the loose nAmb termed “ambiguus cardiovascular” that selectively innervate a subset of cardiac parasympathetic ganglion neurons that slow heart rate and atrioventricular node conduction in response to increased blood pressure^111^, a classical parasympathetic response to stress. Taken together, we observed adaptive DGE in males towards buffering oxidative stress in one population of motor neurons that contribute to ventricular contractility, and thus, HRV measures. In contrast, females displayed maladaptive changes in transcripts important for inhibitory neuronal signaling in the same motor neuron population. Females also displayed maladaptive changes in motor neurons of the loose nAmb which directly engage cardiac ganglion to slow heart rate following stress^111^. There are many other cell types within these subnuclei that likely contribute to DGE due to ELA, and future studies can test functional effects of ELA on the GI system in particular, which comprises a large portion of NTS and DMV neurons within the regions analyzed here and is also an important component of the autonomic stress response. Our spatial data provide a strong foundation for addressing future questions regarding the mechanistic connection between ELA-induced changes in gene expression and vagal circuit function.

### Sex Differences

Sex differences in the LBN model have been shown in multiple brain systems and behaviors^66, 68, 70–74, 138^, including those reported here. ELA-induced sex differences could arise due to a combination of factors. First, our CAU dataset revealed sex differences in the typical development of HRV metrics. Specifically, females have mature autonomic responses before puberty, closer to the occurrence of the stressor, while the male responses do not mature until after puberty. This suggests that the male system has 1) a longer time to adapt to the single stressor that occurred early, and 2) the ability to leverage the plasticity that occurs during puberty to aid in early stress adaptation. Second, the LBN model changes baseline corticosterone levels in rodents in a sex-and developmentally-dependent manner^138–140^. By weaning, female rodents exhibited increased basal levels of plasma corticosterone compared to CAU, whereas levels decreased in males^138–140^. These studies further showed that the LBN paradigm influences maturational timing of early neurodevelopmental milestones in both sexes, but, consistent with our HRV data, only females displayed anxiety-like behavior in adults^138^. This is consistent with the present data revealing that LBN induces precocious maturation of autonomic circuits (a shift in females: P21 → P9; a shift in males: P50 → P35). Third, though yet to be examined in the mouse LBN model, CAU female rats preferentially lick and groom the anogenital region of male pups^141^. More recent findings studied maternal behavior of rats exposed to insufficient nesting material and found that female pups received higher levels of a variety of adverse care activities compared to male pups when the dam returned to the nest^142^. Although speculative, a combination of typical maturational differences, baseline differences in corticosterone, and differences in maternal care could contribute collectively to sex differences in the LBN mouse model^66, 68, 70–74, 138^. Future studies will test whether male and female mouse pups receive differential maternal care in the LBN model.

### Conclusions

The developmental perspective central to our study design revealed longitudinal changes that preceded adult outcomes. It also provided new information on cell-type and topographical regional heterogeneity of responses to ELA. These findings further emphasize sex differences that exist early, prepubertally, in neural responses to stress. This study also contributes to accumulating evidence of the role of mitochondria and oxidative stress in biological responses to ELA, reported in multiple rodent models and most recently in human infants^128, 133^. Altered inhibitory signaling within the DMV appears to be a consistent response to early life environmental experience^134–137^.

In summary, our data provide the following evidence for how vagal circuitry responds to ELA in a sex-dependent manner (Fig. 7): A female mouse experiences ELA followed by a mild stressor, acutely increasing heart rate and respiration (HRV). As time passes, rather than returning to a healthy homeostatic baseline, allostatic load occurs due to molecular changes, including altered inhibitory signaling, oxidative stress, and mitochondrial dysfunction. Thus, autonomic regulation (HRV) remains low and heart rate remains high, leading to chronic stress within the cardiovascular system with the potential for long-term effects on both the brain and the body. Conversely, male mice that experience ELA followed by a mild stressor have cellular changes that retain balanced inhibitory signaling, buffering of oxidative stress, and protection of mitochondrial function. This facilitates the return of heart rate and respiration to homeostasis, potentially contributing to autonomic stress resilience. Our focus here is on changes within neurons located at intermediate levels of the vagal medulla and the loose nAmb, as they relate to the production of respHRV and display molecular sex differences in the ELA response. However, transcript-specific changes observed within the rostral and caudal vagal medulla follow a different model, related to different organ systems, some of which do not display sex differences. ELA results in a variety of chronic health conditions that affect different organ systems, with potentially shared or diverging etiology depending on the vagal circuitry involved. Pursuing detailed understandings from a developmental perspective of complex circuitry can greatly contribute to personalized medicine and healthcare approaches that do not use a one-size-fits-all approach to treat chronic stress.

**Figure 7.**
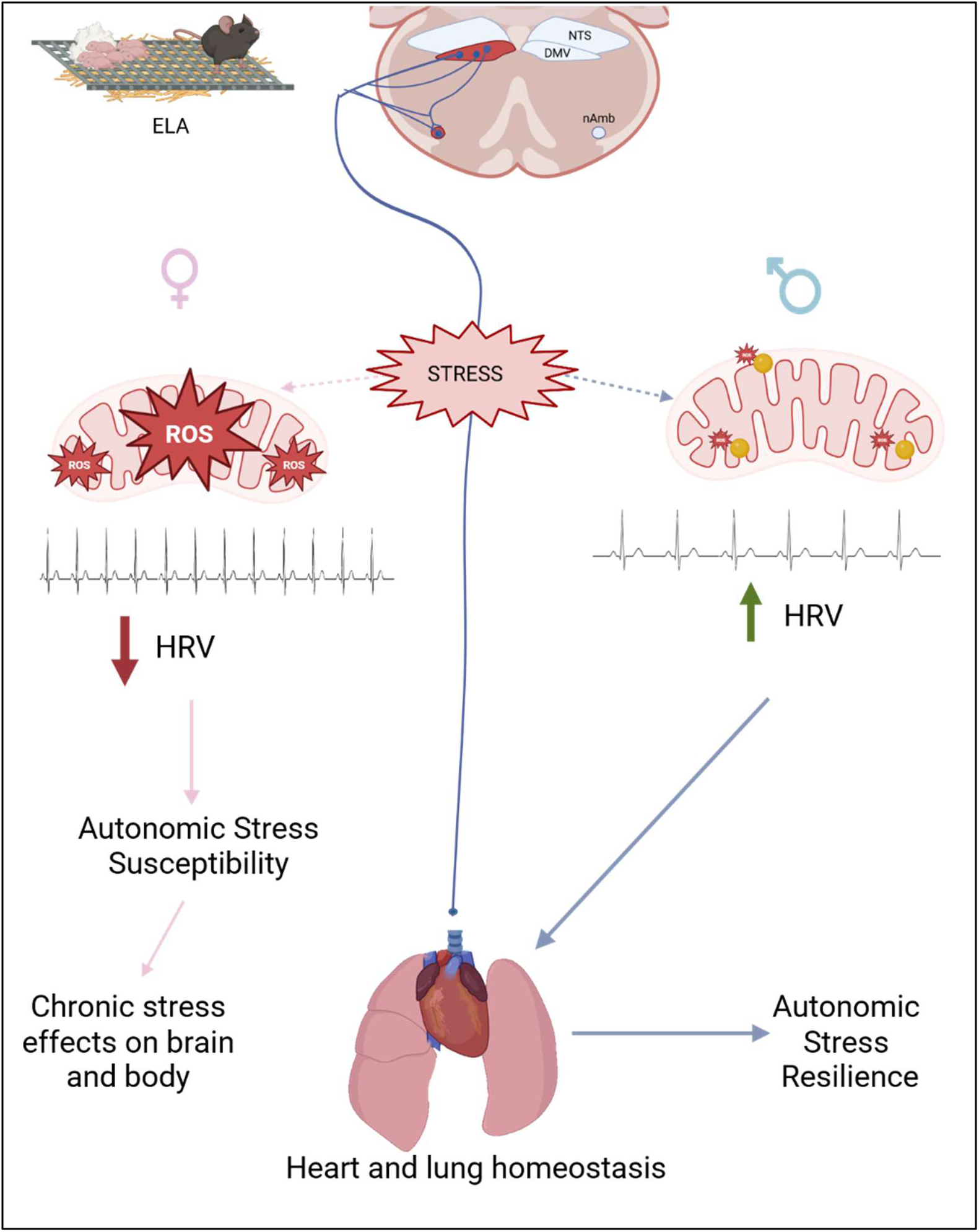
A schematic summarizing our findings for how ELA-induced molecular changes impact the motor neurons responsible for generating the autonomic stress response. Under baseline conditions, motor neurons in the DMV and nAmb exhibit tonic inhibitory control over heart rate. When a male or female mouse experiences a mild, tolerable stressor, vagal motor neurons are inhibited and the sympathetic nervous system increases heart rate and respiration. Even if a male mouse has a history of ELA, these neurons are able to buffer oxidative stress and balance inhibitory signaling, allowing for re-engagement of vagal neurons to slow heart-rate, increasing HRV and bringing the body back to homeostasis. This is part of a typical, healthy stress response that can lead to resilience within autonomic and forebrain circuits. A female mouse with a history of ELA have motor neurons with maladaptive molecular changes, resulting in mitochondrial dysfunction, oxidative cellular damage, and dysregulated inhibitory signaling in vagal motor neurons responsible for HRV. So, even in the absence of the stressor, heart rate remains high and HRV remains low, potentially contributing to autonomic stress susceptibility.

### Author Contributions

SP conceptualized experiments; performed HRV paradigm set-up, recordings, and analysis; snRNAseq sample preparation and analysis; and HiPlex RnaScope tissue preparation and statistical analysis. SP also wrote the first draft of this manuscript. SK performed all HiPlex RnaScope experiments and area fraction generation. SK also wrote the HiPlex RnaScope methods. MH assisted in HRV analysis, tissue processing, and LBN paradigm maintenance. PL conceptualized experiments, data analysis strategies and supervised all studies. PL, MH, and SK edited all drafts of this manuscript.

## Acknowledgements

Simms/Mann Chair in Developmental Neurogenetics, WM Keck Provost Professor of Neurogenetics, CHLA The Saban Research Institute (TSRI) Developmental Neuroscience and Neurogenetics Program. Image acquisition was performed at the TSRI Cellular Imaging Core and the single nucleus sequencing was performed at the TSRI Spatial Biology and Genomics Core. This work was funded by a Research Career Development Fellowship Award granted to SP by TSRI and an R01 awarded to PL from the NIDDK (R01 DK126085).

## Supplementary Figures

**Supplemental Figure 1.**
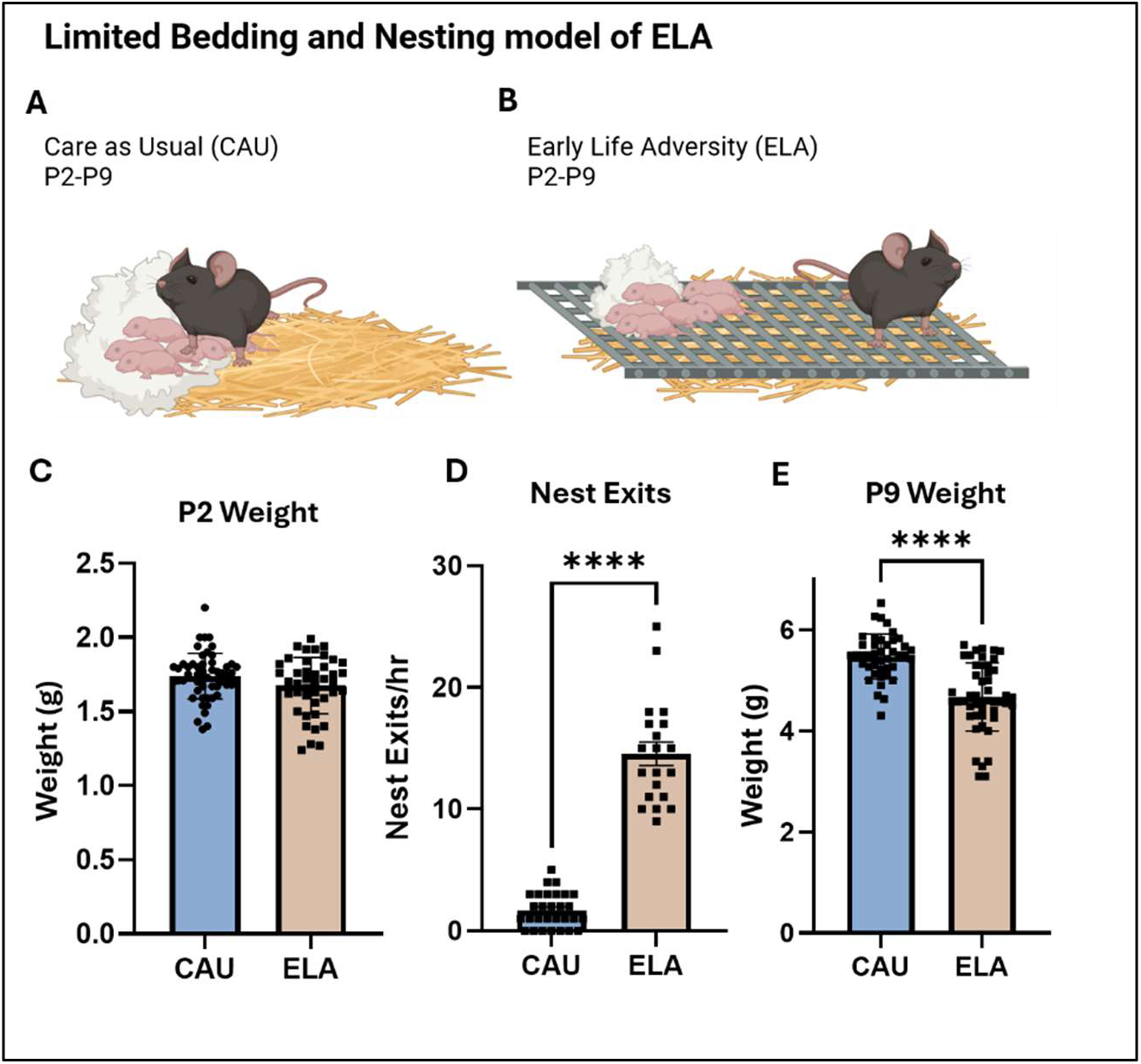
Limited bedding and nesting model of ELA. A) CAU dams receive 100g of bedding and 2.7 grams of nesting material from P2-P9 following litter culling to 3 males and 3 females. B) ELA dams receive 50 g of bedding, 1.35 g of nesting, and placement of a wire grid from P2-P9. C) There are no differences in pup weight when they begin the LBN paradigm (t(94)= 1.8000; p= 0.0751, Cohen’s d= -0.3703). D) Nest exits are significantly increased in the ELA condition compared to CAU (t(27)= 12.44, p< 0.0001, Cohen’s d= 4.762). E) At the end of the LBN period, ELA pups weigh significantly less than CAU pups (t(111)= 4.570, p< 0.0001, Cohen’s d= -0.8607), suggesting impacted maternal care.

**Supplemental Figure 2.**
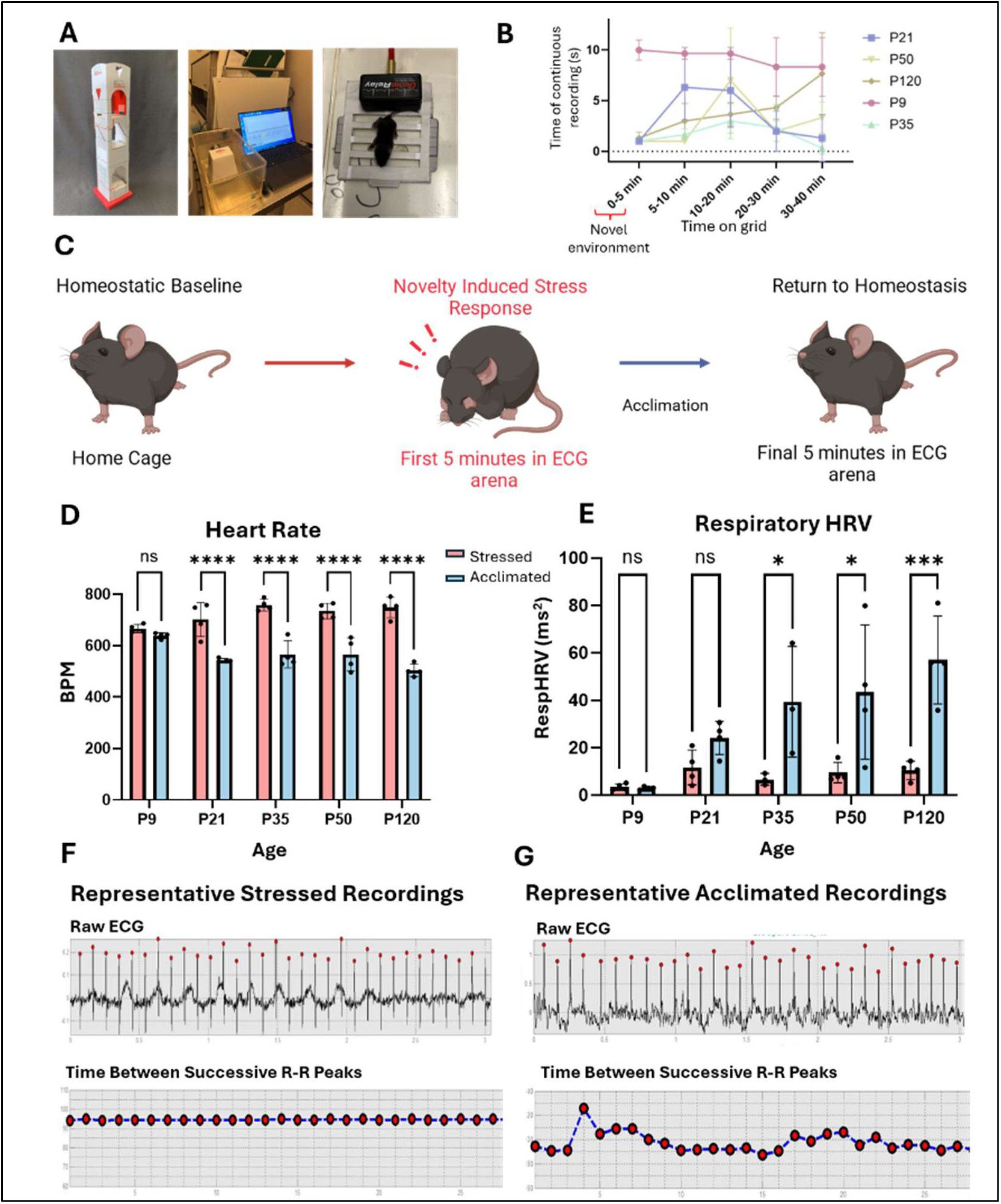
Adapted ECGenie Paradigm. A) Original ECGenie paradigm and our adapted paradigm. B) Continuous recording times as a function of time spent in the arena. Each age had optimal acclimation times. C) Paradigm to produce an HRV response. D) A two-way ANOVA showed a main effect of acclimation (F(1,6)= 130.6, p< 0.0001) and an interaction effect between age and acclimation (F(2,24)= 8.914, p= 0.0001) on heart rate. Post-hoc tests revealed that ages P21 and greater showed a robust decrease in heart rate (P9 t(30)= 0.9812, padj= 0.3343; P21 t(30)=5.698, padj <0.0001 ; P35 t(30) = 6.914, padj <0.0001; P50 t(30)= 6.067, padj < 0.0001; P120 t(30)= 8.804, padj <0.0001) during the acclimated portion of the task. E) A two-way ANOVA showed main effects of age (F(4,14)= 5.479, p= 0.0072), acclimation (F(1,14)= 35.90, p<0.001), and an interaction effect (F(4,14)= 4.269, p= 0.083) on respHRV. Post-hoc tests revealed increased respHRV during the final five minutes of recording at P35 and older (P9 t(14)= 0.05166, p= 0.9595; P21 t(14)= 1.370, p= 0.1924; P35 t(14)= 3.132, padj= 0.0074; P50 t(14)= 3.755, padj= 0.0021; P120 t(14)= 5.148, padj= 0.0001) F) ECG trace and R-R intervals at P120 during the first 5 minutes of recording. G) ECG trace and R-R intervals at P120 during the final 5 minutes.

**Supplemental Figure 3.**
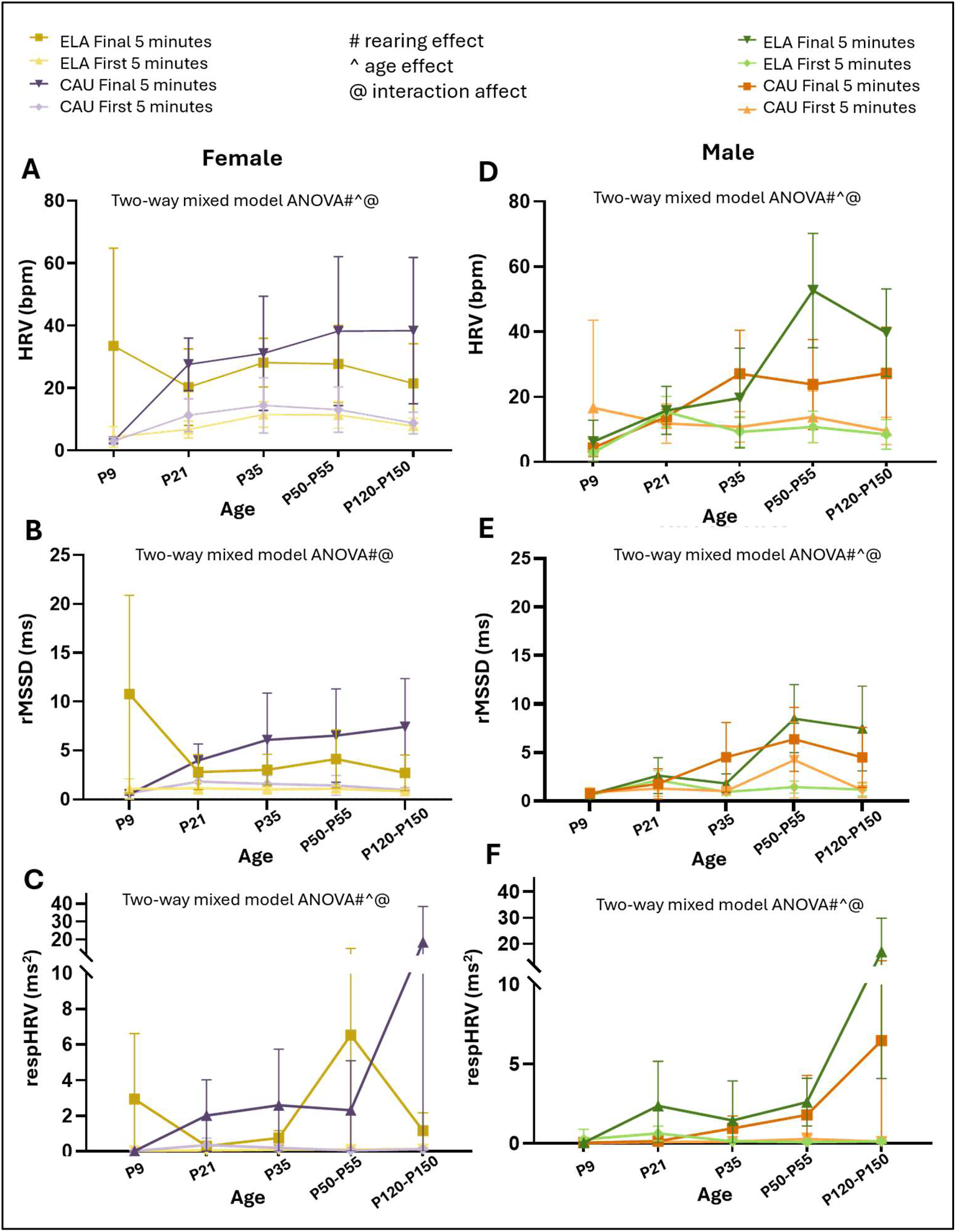
Comparison of all HRV metrics between CAU and ELA across development. All tests used two-way mixed model ANOVAs testing for effects of age and rearing on HRV metrics. Bonferroni post-host testing was used to detect individual differences and none reached statistical significance. A) HRV females displayed significant age (F (2.945, 97.18) = 3.654; p= 0.0157), rearing (F (3, 132) = 23.22; p< 0.0001), and interaction effects (F (12, 132) = 2.647; p= 0.0033). B) rMSSD females displayed significant rearing (F (3, 124) = 16.55; p< 0.0001) and an interaction effect between age and rearing (F (12, 124) = 4.531; p< 0.0001). C) respHRV females displayed significant age (F (1.501, 36.02) = 3.978; p= 0.0381), rearing (F (3, 28) = 7.028; p= 0.0011), and interaction effects (F (12, 96) = 4.861; p< 0.0001). D) Male HRV displayed significant effects of age (F (3.089, 78.01) = 11.09; p< 0.0001), rearing (F (3, 30) = 15.53; p< 0.0001), and an interaction effect (F (9.268, 78.01) = 5.781; p< 0.0001). E) Male rMSSD displayed significant effects of age (F (2.679, 84.39) = 15.94; p< 0.0001), rearing (F (3, 126) = 14.58; p< 0.0001), and an interaction effect (F (12, 126) = 3.991; p< 0.0001). F) Male respHRV displayed significant effects of age (F (1.165, 28.83) = 15.00; p= 0.0003), rearing (F (3, 30) = 8.819; p= 0.0002), and an interaction effect (F (12, 99) = 7.013; p< 0.0001).

**Supplemental Figure 4.**
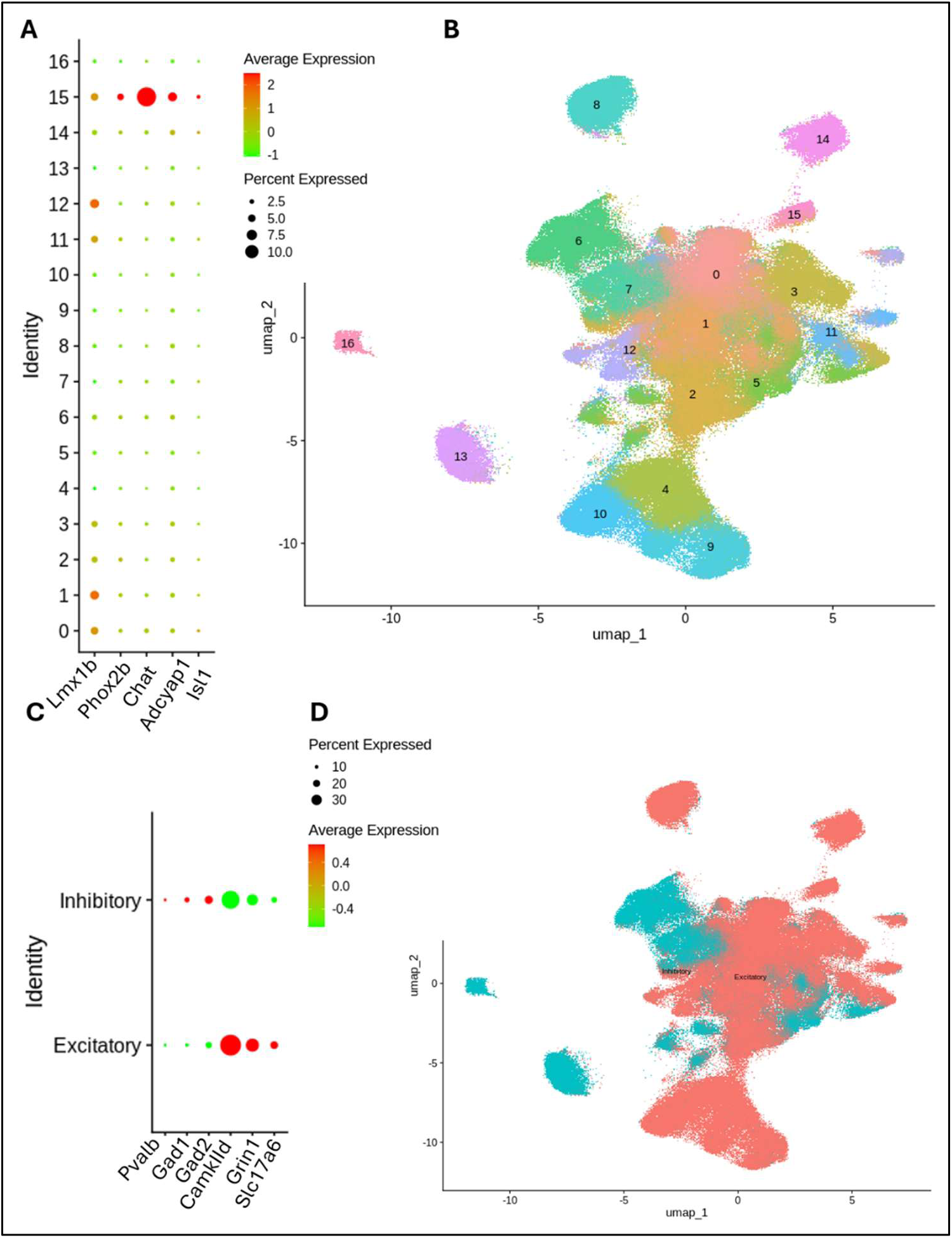
P9 snRNAseq neuronal subclusters. A) Distribution of NTS (Phox2b + Lmx1b), DMV (Phox2b + Chat + Adcyap), nAmb (Phox2b + Chat + Isl1) marker genes among neuronal clusters. B) UMAP display of neuronal clusters. C) Inhibitory (Pvalb, Gad1, Gad2) and excitatory (CamkII2d, Grin1, Slc17a6) neuron markers among neuronal subclusters. D) UMAP display of cluster identities.

**Supplemental Figure 5.**
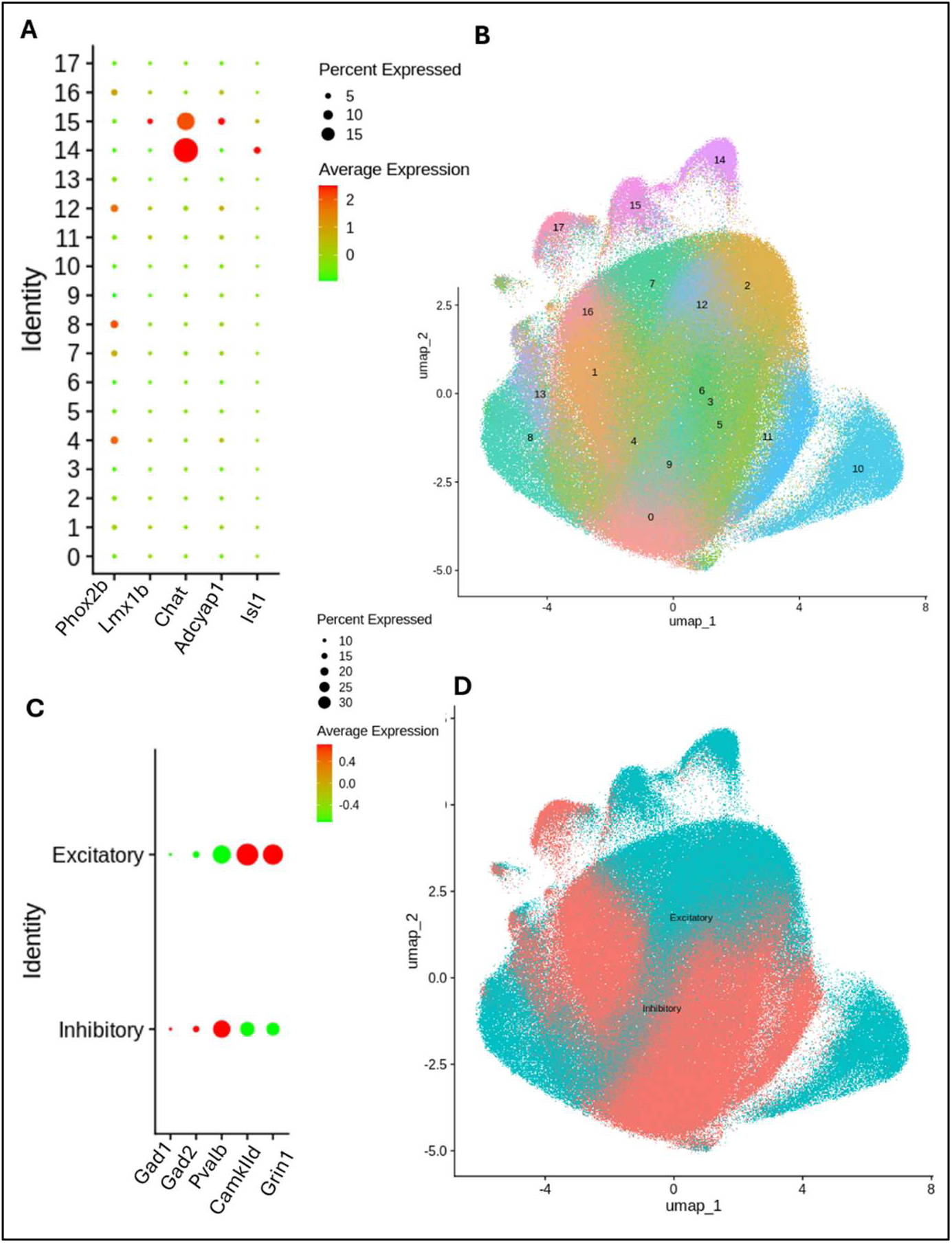
P120 snRNAseq neuronal subclusters. A) Distribution of NTS (Phox2b + Lmx1b), DMV (Phox2b + Chat + Adcyap), nAmb (Phox2b + Chat + Isl1) marker genes among neuronal clusters. B) UMAP display of neuronal clusters. C) Inhibitory (Pvalb, Gad1, Gad2) and excitatory (CamkII2d, Grin1, Slc17a6) neuron markers among neuronal subclusters. D) UMAP display of cluster identities.

**Supplemental Figure 6.**
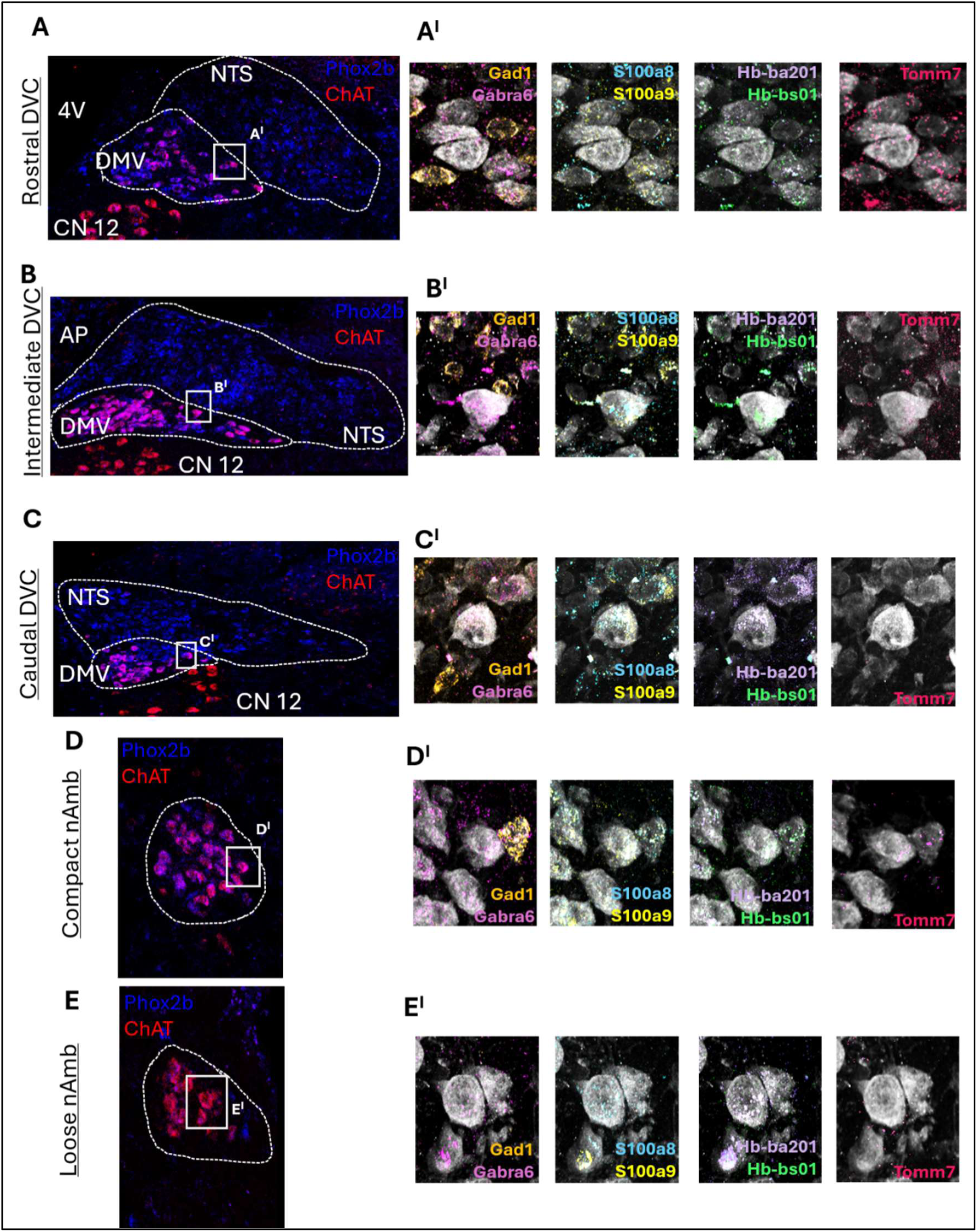
Expression of high-confidence DEGs using in situ hybridization along the rostro-caudal axis of the vagal medulla. Differential Phox2b and Chat expression was used to identify the NTS (Phox2b + Chat-) and the DMV and nAmb (Phox2b+ Chat+) (A-E). High magnification images show differential expression of genes along the rostro-caudal axis of neurons within our regions of interest (A’-E’)

**Supplemental Figure 7.**
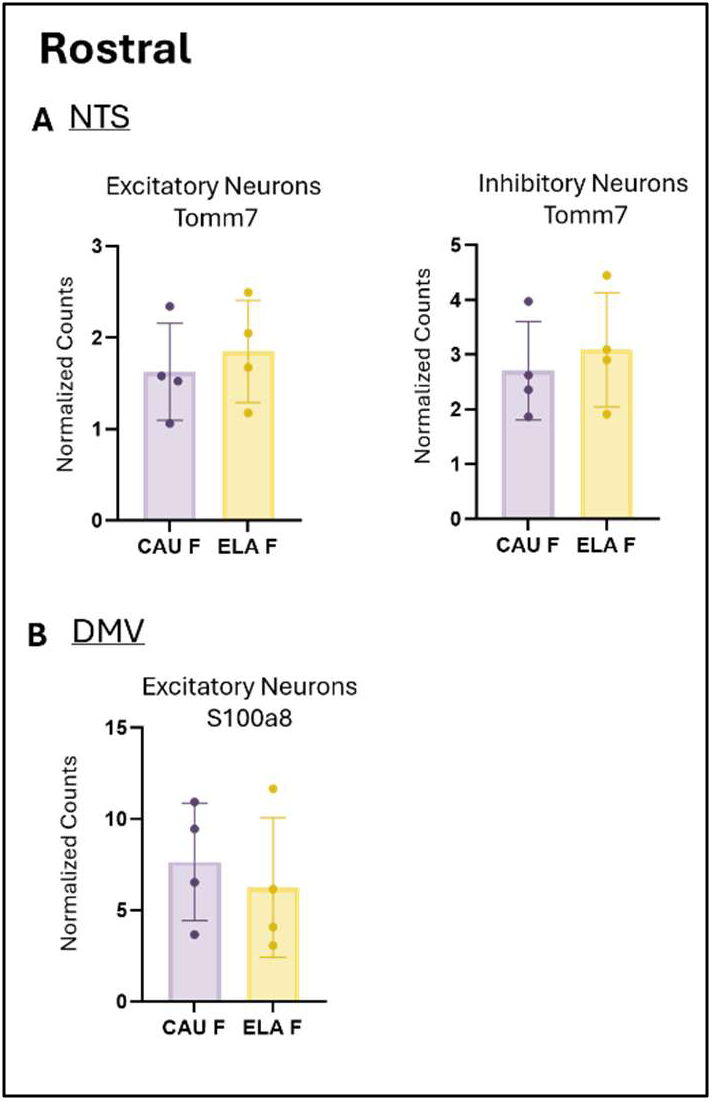
Nonsignificant hiplex results for the rostral vagal medulla. A) Rostral NTS displays no significant differences in Tomm7 expression in excitatory neurons (t(6)= 0.5729, p= 0.58729, Cohen’s d = 0.4051) or inhibitory neurons (t(6)= 0.5583, p= 0.5968, Cohen’s d=.3958) of female mice. B) Rostral DMV displays no significant differences in S100a8 expression in excitatory neurons (t(6)= 0.5617, p= 0.5947, Cohen’s d = -0.3972) of female mice.

**Supplemental Figure 8.**
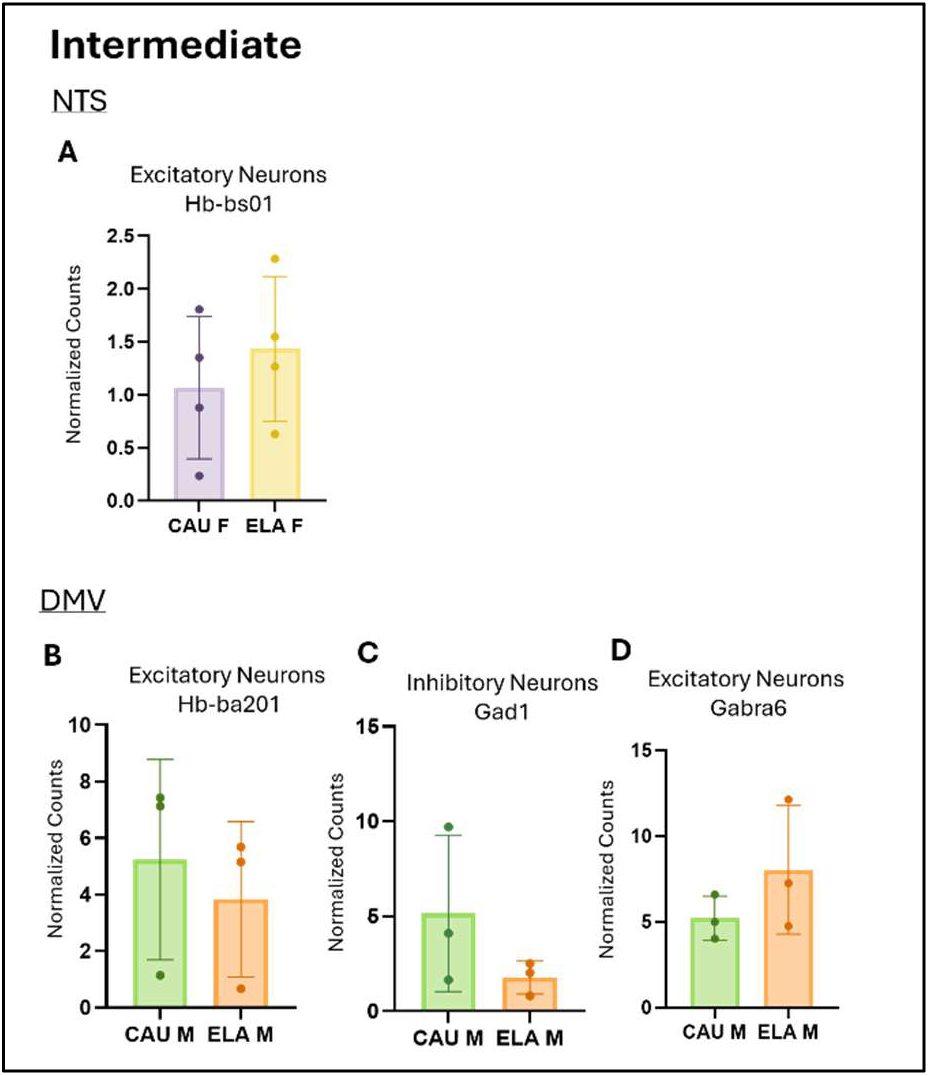
Nonsignificant hiplex results for the intermediate vagal medulla. **A**) Intermediate NTS displays no significant differences in Hb-bs01 expression in excitatory neurons (t(6)= 0.7596, p= 0.4762, Cohen’s d = 0.5) of female mice. B) Intermediate DMV displays no significant differences in Hb-ba expression in excitatory neurons (t(4)= 0.5393, p= 0.6183, Cohen’s d=-0.4404) of male mice. C) There were trending decreased in Gad1 expression in inhibitory neurons of ELA male mice (t(4)= 1.387, p= 0.2378, d = -1.132). D) There were trending increases in Gabra6 expression in excitatory neurons of ELA male mice (t(4)= 1.239, p= 0.2831, d= 1.012)

**Supplemental Figure 9.**
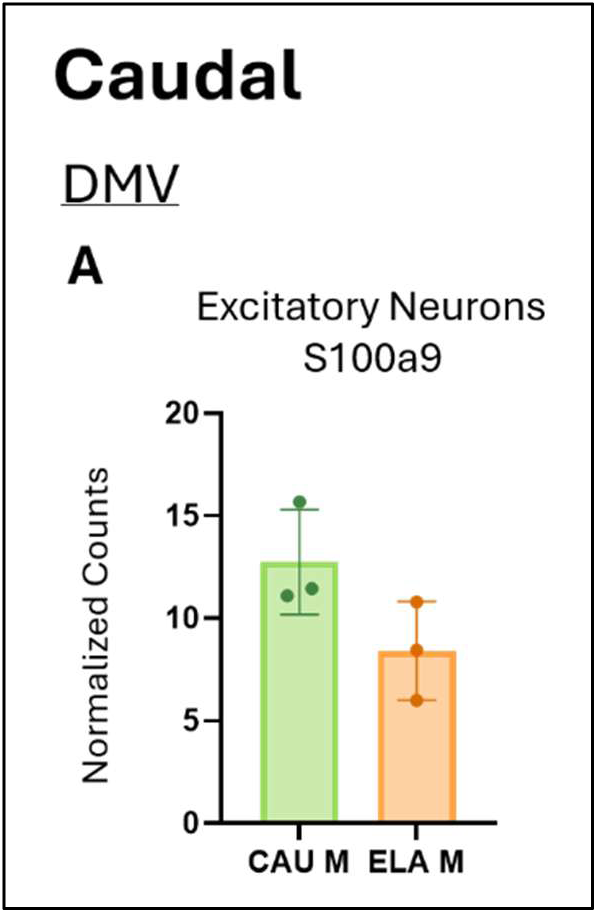
Nonsignificant hiplex results for the caudal DMV. A) S100a9 displayed trending decreases in S100a9 expression in ELA male mice (t(4)= 2.140, p= 0.0991, Cohen’s d= -1.747).

**Supplemental Figure 10.**
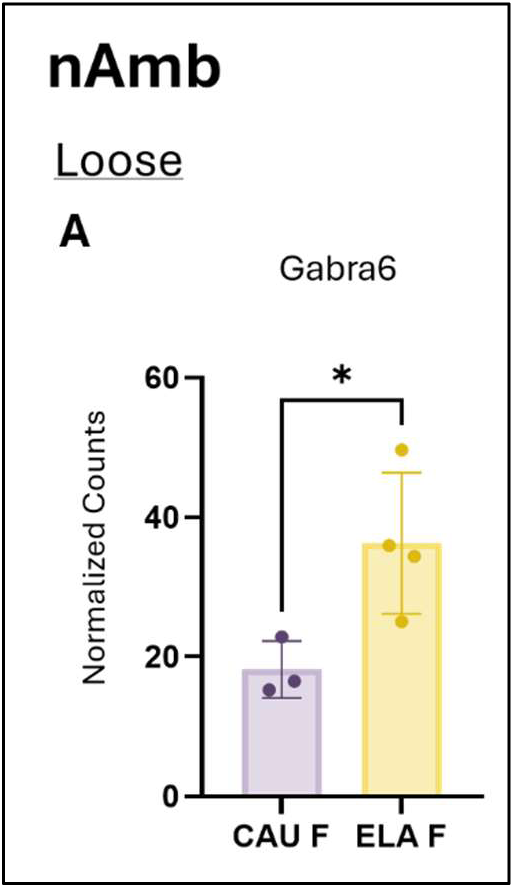
Expression of Gabra6 in the loose nAmb. A) There was increased expression of Gabra6 in neurons of the loose nAmb in ELA female mice (t(5)= 2.866, p= 0.0351, Cohen’s d= 2.189).

