## Supplemental Table 13 for "Early Life Adversity Produces Enduring Molecular and Functional Disruption of Developing Vagal Circuits"

| Brain Region | Vagal Subtype | Inputs/Outputs | Neuronal Subtype | Sex Effect? | DEGs | Pathways |
|---|---|---|---|---|---|---|
| Rostral NTS (transition zone) | Sensory | Direct relay of interoceptive signals to the DMV | Excitatory, Inhibitory | YES, males only | Tomm7 | Mitochondrial function |
| Intermediate NTS | Sensory | Gastrointestinal, cardiovascular, pulmonary | Excitatory | No | Hbb-bs01 | Oxidative stress |
| Caudal NTS | Sensory | Gastrointestinal, cardiovascular, pulmonary | Excitatory, Inhibitory | No | S100a9, Hbb-ba201 | Oxidative stress |
| Rostral DMV | Motor | Gastrointestinal (contraction), pancreas, liver | Excitatory, Inhibitory | No | Tomm7, S100a8 | Mitochondrial function, oxidative stress |
| Intermediate DMV | Motor | Cardiac inhibition, stomach, duodenom | Excitatory, Inhibitory | YES | Hbb-ba201, Gad1, Gabra6 | Oxidative stress, inhibitory signaling |
| Caudal DMV | Motor | Stomach, lower GI trac (gastric relaxation) | Excitatory | No | S100a9, Hb-ba201 | Oxidative stress |
| Compact nAmb | Motor | Cardiopulmonary, esophagus (swallowing) | None | No | None | None |
| Semi-Compact/Loose nAmb | Motor | Cardiovascular, soft palate, pharynx, larynx (speech) | Inhibitory | No | Gabra6 | Inhibitory signaling |
