## Supplemental Table 1 for "Early Life Adversity Produces Enduring Molecular and Functional Disruption of Developing Vagal Circuits"

© 2000-2025 QIAGEN. All rights reserved.

**P9 EXCITATORY FEMALE NEURONS**

| Ingenuity Canonical Pathways | -log(p-value) | Ratio | z-score | Molecules |
|---|---|---|---|---|
| G alpha (q) signalling events | 4.84 | 0.0402 | -2.646 | CASR,GCG,GNG3,NPFFR2,NTS,TAC1,UTS2B |
| Cardiac conduction | 3.5 | 0.0385 | 0 | CASQ2,CORIN,FXYD7,ITPR1,NPPC |
| Extra-nuclear estrogen signaling | 3.43 | 0.0533 | -2 | FOS,GNG3,HRAS,HSPB1 |
| GDNF Family Ligand-Receptor Interactions | 3.41 | 0.0526 | #NUM! | FOS,HRAS,ITPR1,NRTN |
| GPCR-Mediated Integration of Enteroendocrine Signaling Exemplified by an L Cell | 3.41 | 0.0526 | 0 | CALCA,GAL,GCG,ITPR1 |
| Respiratory electron transport | 3.16 | 0.0455 | -2 | COX5B,COX8A,NDUFA13,NDUFS7 |
| Class A/1 (Rhodopsin-like receptors) | 3.1 | 0.0211 | -2.646 | GAL,NPFFR2,NPY,NTS,PENK,TAC1,UTS2B |
| Complex IV assembly | 2.87 | 0.0625 | #NUM! | COX5B,COX8A,SCO2 |
| Oxidative Phosphorylation | 2.83 | 0.037 | -2 | COX5B,COX8A,NDUFA13,NDUFS7 |
| Huntington's Disease Signaling | 2.71 | 0.0209 | #NUM! | CTSD,GNG3,HRAS,ITPR1,PENK,PSMB5 |

| | | | | |
|---|---|---|---|---|
| EGF Signaling | 2.68 | 0.0536 | #NUM! | FOS,HRAS,ITPR1 |
| GPCR-Mediated Nutrient Sensing in Enteroendocrine Cells | 2.68 | 0.0336 | -1 | CASR,GCG,GNG3,ITPR1 |
| Gustation Pathway | 2.56 | 0.0237 | 0.447 | CASR,GABRA6,GCG,ITPR1,LIPC |
| IL-6 Signaling | 2.55 | 0.031 | -2 | CEBPB,FOS,HRAS,HSPB1 |
| Thyroid Cancer Signaling | 2.27 | 0.0385 | #NUM! | FOS,HRAS,NTF3 |
| Neurotrophin/TRK Signaling | 2.27 | 0.0385 | #NUM! | FOS,HRAS,NTF3 |
| Wound Healing Signaling Pathway | 2.26 | 0.0201 | -2.236 | CALCA,CEBPB,FOS,HRAS,TAC1 |
| Incretin synthesis, secretion, and inactivation | 2.25 | 0.08 | #NUM! | GCG,ISL1 |
| Signaling by NTRK2 (TRKB) | 2.25 | 0.08 | #NUM! | HRAS,NTF3 |
| JAK/STAT Signaling | 2.2 | 0.0361 | #NUM! | CEBPB,FOS,HRAS |
| G alpha (i) signalling events | 2.2 | 0.0194 | -2.236 | CASR,GAL,GNG3,NPY,PENK |
| TP53 Regulates Metabolic Genes | 2.17 | 0.0353 | #NUM! | COX5B,COX8A,SCO2 |
| CXCR4 Signaling | 2.15 | 0.0238 | #NUM! | FOS,GNG3,HRAS,ITPR1 |
| Signaling by NTRK3 (TRKC) | 2.09 | 0.0667 | #NUM! | HRAS,NTF3 |
| Ceramide Signaling | 2.09 | 0.033 | #NUM! | CTSD,FOS,HRAS |
| Thyroid Hormone Biosynthesis | 2.05 | 0.5 | #NUM! | CTSD |
| Opioid Signaling Pathway | 2.04 | 0.0178 | -1 | FOS,GNG3,HRAS,ITPR1,PENK |
| Class B/2 (Secretin family receptors) | 2.04 | 0.0316 | #NUM! | CALCA,GCG,GNG3 |

| | | | | |
|---|---|---|---|---|
| Erythropoietin Signaling Pathway | 2.03 | 0.0221 | #NUM! | FOS,HBA2,HRAS,ITPR1 |
| Prolactin Signaling | 2.03 | 0.0312 | #NUM! | CEBPB,FOS,HRAS |
| Oxytocin Signaling Pathway | 2.02 | 0.0175 | -1.342 | FOS,GNG3,HRAS,HSPB1,ITPR1 |
| IL-17 Signaling | 1.97 | 0.0212 | -2 | CEBPB,FOS,HRAS,TRAF5 |
| Neuropathic Pain Signaling in Dorsal Horn Neurons | 1.97 | 0.0297 | #NUM! | FOS,ITPR1,TAC1 |
| GNRH Signaling | 1.94 | 0.0207 | #NUM! | FOS,GNG3,HRAS,ITPR1 |
| Signaling by VEGF | 1.9 | 0.028 | #NUM! | HRAS,HSPB1,ITPR1 |
| Integration of energy metabolism | 1.89 | 0.0278 | #NUM! | GCG,GNG3,ITPR1 |
| PPAR Signaling | 1.89 | 0.0278 | #NUM! | FOS,HRAS,SCAND1 |
| Adrenomedullin signaling pathway | 1.87 | 0.0198 | -1 | CEBPB,FOS,HRAS,ITPR1 |
| Metabolism of vitamin K | 1.87 | 0.333 | #NUM! | VKORC1 |
| NADH Repair | 1.87 | 0.333 | #NUM! | GAPDH |
| α-Adrenergic Signaling | 1.86 | 0.027 | #NUM! | GNG3,HRAS,ITPR1 |
| Fc epsilon receptor (FCERI) signaling | 1.85 | 0.0194 | #NUM! | FOS,HRAS,ITPR1,PSMB5 |
| April Mediated Signaling | 1.81 | 0.0476 | #NUM! | FOS,TRAF5 |
| Cholecystokinin/Gastrin-mediated Signaling | 1.79 | 0.0256 | #NUM! | FOS,HRAS,ITPR1 |
| B Cell Activating Factor Signaling | 1.79 | 0.0465 | #NUM! | FOS,TRAF5 |
| Signaling by Insulin receptor | 1.75 | 0.0444 | #NUM! | CTSD,HRAS |
| Renin-Angiotensin Signaling | 1.75 | 0.0246 | #NUM! | FOS,HRAS,ITPR1 |
| Catecholamine Biosynthesis | 1.75 | 0.25 | #NUM! | DBH |

| | | | | |
|---|---|---|---|---|
| Role of Macrophages, Fibroblasts and Endothelial Cells in Rheumatoid Arthritis | 1.73 | 0.0148 | -1.342 | CEBPB,FOS,HRAS,TRAF5,WIF1 |
| HER-2 Signaling in Breast Cancer | 1.71 | 0.0176 | 0 | COX5B,COX8A,FOS,HRAS |
| Mitochondrial Dysfunction | 1.69 | 0.0145 | 1.342 | COX5B,COX8A,ITPR1,NDUFA13,NDUFS7 |
| Antimicrobial peptides | 1.68 | 0.0408 | #NUM! | ATOX1,CLU |
| G Beta Gamma Signaling | 1.67 | 0.0231 | #NUM! | GNG3,HRAS,ITPR1 |
| UVC-Induced MAPK Signaling | 1.65 | 0.0392 | #NUM! | FOS,HRAS |
| fMLP Signaling in Neutrophils | 1.65 | 0.0226 | #NUM! | GNG3,HRAS,ITPR1 |
| GABA Receptor Signaling | 1.65 | 0.0226 | #NUM! | GABRA6,GNG3,ITPR1 |
| P2Y Purinergic Receptor Signaling Pathway | 1.64 | 0.0224 | #NUM! | FOS,GNG3,HRAS |
| CCR3 Signaling in Eosinophils | 1.62 | 0.0221 | #NUM! | GNG3,HRAS,ITPR1 |
| S100 Family Signaling Pathway | 1.59 | 0.0102 | -1.414 | BSG,CASR,CTSD,FOS,ITPR1,NPFFR2,NTF3,S100A10 |
| GABAergic Receptor Signaling Pathway (Enhanced) | 1.59 | 0.0214 | #NUM! | GABRA6,GNG3,ITPR1 |
| CD27 Signaling in Lymphocytes | 1.58 | 0.0357 | #NUM! | FOS,TRAF5 |
| Selenocysteine Biosynthesis II (Archaea and Eukaryotes) | 1.57 | 0.167 | #NUM! | SEPHS2 |
| G alpha (s) signalling events | 1.55 | 0.0207 | #NUM! | CALCA,GCG,GNG3 |
| C-type lectin receptors (CLRs) | 1.55 | 0.0207 | #NUM! | HRAS,ITPR1,PSMB5 |

| | | | | |
|---|---|---|---|---|
| Apelin Endothelial Signaling Pathway | 1.55 | 0.0207 | #NUM! | FOS,GNG3,HRAS |
| Cytoprotection by HMOX1 | 1.55 | 0.0345 | #NUM! | COX5B,COX8A |
| GABA receptor activation | 1.52 | 0.0333 | #NUM! | GABRA6,GNG3 |
| RAF/MAP kinase cascade | 1.51 | 0.0153 | #NUM! | HRAS,NEFL,NRTN,PSMB5 |
| NCAM signaling for neurite out-growth | 1.48 | 0.0317 | #NUM! | HRAS,NRTN |
| IL-2 Signaling | 1.48 | 0.0317 | #NUM! | FOS,HRAS |
| Thrombopoietin Signaling | 1.47 | 0.0312 | #NUM! | FOS,HRAS |
| Circadian Rhythm Signaling | 1.47 | 0.0148 | #NUM! | GNG3,HRAS,ITPR1,PTGDS |
| Mitochondrial protein import | 1.46 | 0.0308 | #NUM! | TIMM10B,TIMM13 |
| Role of JAK2 in Hormone-like Cytokine Signaling | 1.46 | 0.0308 | #NUM! | FOS,NPY |
| CD40 Signaling | 1.43 | 0.0299 | #NUM! | FOS,TRAF5 |
| Complex I biogenesis | 1.42 | 0.0294 | #NUM! | NDUFA13,NDUFS7 |
| Estrogen Receptor Signaling | 1.41 | 0.0122 | #NUM! | FOS,GNG3,HRAS,NDUFA13,NDUFS7 |
| Eicosanoid Signaling | 1.41 | 0.0142 | -2 | GNG3,HRAS,HSPB1,PTGDS |
| Mitochondrial Division Signaling Pathway | 1.4 | 0.0181 | #NUM! | HRAS,ITPR1,MUL1 |
| Signaling by the B Cell Receptor (BCR) | 1.38 | 0.0176 | #NUM! | HRAS,ITPR1,PSMB5 |
| Dopamine Receptor Signaling | 1.38 | 0.0278 | #NUM! | CALY,SLC18A3 |
| Ephrin B Signaling | 1.37 | 0.0274 | #NUM! | GNG3,HRAS |
| Senescence-Associated Secretory Phenotype (SASP) | 1.36 | 0.027 | #NUM! | CEBPB,FOS |
| ERK5 Signaling | 1.36 | 0.027 | #NUM! | FOS,HRAS |

| | | | | |
|---|---|---|---|---|
| Formation of the posterior neural plate | 1.35 | 0.1 | #NUM! | POU3F1 |
| Embryonic Stem Cell Differentiation into Cardiac Lineages | 1.35 | 0.1 | #NUM! | ISL1 |
| Androgen Signaling | 1.35 | 0.0171 | #NUM! | GNG3,ITPR1,TAF7 |
| Granzyme A Signaling | 1.34 | 0.0267 | #NUM! | NDUFA13,NDUFS7 |
| Antiproliferative Role of Somatostatin Receptor 2 | 1.32 | 0.026 | #NUM! | GNG3,HRAS |
| Formation of the anterior neural plate | 1.31 | 0.0909 | #NUM! | POU3F1 |
| Prostanoid Biosynthesis | 1.31 | 0.0909 | #NUM! | PTGDS |
| VDR/RXR Activation | 1.31 | 0.0256 | #NUM! | CASR,CEBPB |
| Renal Cell Carcinoma Signaling | 1.31 | 0.0256 | #NUM! | FOS,HRAS |
| Aryl Hydrocarbon Receptor Signaling | 1.3 | 0.0164 | #NUM! | CTSD,FOS,HSPB1 |
| IL-17A Signaling in Fibroblasts | 1.29 | 0.025 | #NUM! | CEBPB,FOS |
| IL-3 Signaling | 1.28 | 0.0247 | #NUM! | FOS,HRAS |
| Chemokine Signaling | 1.28 | 0.0247 | #NUM! | FOS,HRAS |
| Estrogen-Dependent Breast Cancer Signaling | 1.27 | 0.0241 | #NUM! | FOS,HRAS |
| Transcriptional regulation of white adipocyte differentiation | 1.26 | 0.0238 | #NUM! | CEBPB,MED29 |
| Acute Phase Response Signaling | 1.25 | 0.0157 | #NUM! | CEBPB,FOS,HRAS |
| Activation of NMDA receptors and postsynaptic events | 1.25 | 0.0235 | #NUM! | HRAS,NEFL |
| LPS-stimulated MAPK Signaling | 1.25 | 0.0235 | #NUM! | FOS,HRAS |
| Dissolution of Fibrin Clot | 1.24 | 0.0769 | #NUM! | S100A10 |
| Transcriptional regulation of testis differentiation | 1.24 | 0.0769 | #NUM! | PTGDS |
| Platelet homeostasis | 1.24 | 0.0233 | #NUM! | GNG3,ITPR1 |
| Glucose metabolism | 1.24 | 0.0233 | #NUM! | GAPDH,PGP |
| VEGF Family Ligand-Receptor Interactions | 1.24 | 0.0233 | #NUM! | FOS,HRAS |
| FGF Signaling | 1.24 | 0.0233 | #NUM! | HRAS,ITPR1 |

| | | | | |
|---|---|---|---|---|
| Endothelin-1 Signaling | 1.24 | 0.0155 | #NUM! | FOS,HRAS,ITPR1 |
| PDGF Signaling | 1.23 | 0.023 | #NUM! | FOS,HRAS |
| Synaptic Long Term Depression | 1.22 | 0.0152 | #NUM! | HRAS,ITPR1,PPP1R17 |
| Regulation of mRNA stability by proteins that bind AU-rich elements | 1.21 | 0.0225 | #NUM! | HSPB1,PSMB5 |
| Myelination Signaling Pathway | 1.21 | 0.0122 | -1 | FOS,HRAS,NTF3,POU3F1 |
| Opioid Signalling | 1.2 | 0.0222 | #NUM! | GNG3,ITPR1 |
| Oxytocin in Brain Signaling Pathway | 1.2 | 0.0149 | #NUM! | GNG3,HRAS,ITPR1 |
| MAPK6/MAPK4 signaling | 1.19 | 0.0217 | #NUM! | HSPB1,PSMB5 |
| Sheddase Signaling Pathway | 1.19 | 0.0147 | #NUM! | BSG,CEBPB,FOS |
| Synthesis of Prostaglandins (PG) and Thromboxanes (TX) | 1.18 | 0.0667 | #NUM! | PTGDS |
| Response of EIF2AK1 (HRI) to heme deficiency | 1.18 | 0.0667 | #NUM! | CEBPB |
| Beta-catenin independent WNT signaling | 1.18 | 0.0146 | #NUM! | GNG3,ITPR1,PSMB5 |
| RANK Signaling in Osteoclasts | 1.18 | 0.0215 | #NUM! | FOS,TRAF5 |
| ERBB Signaling | 1.18 | 0.0215 | #NUM! | FOS,HRAS |
| IL-8 Signaling | 1.17 | 0.0144 | #NUM! | FOS,GNG3,HRAS |
| Extrinsic Prothrombin Activation Pathway | 1.16 | 0.0625 | #NUM! | F13B |
| Non-Small Cell Lung Cancer Signaling | 1.16 | 0.0208 | #NUM! | HRAS,ITPR1 |
| TGF-β Signaling | 1.15 | 0.0206 | #NUM! | FOS,HRAS |
| p53 Signaling | 1.14 | 0.0204 | #NUM! | RPRM,SCO2 |
| Mitochondrial protein degradation | 1.14 | 0.0204 | #NUM! | COX5B,NDUFA13 |
| IL-1 Signaling | 1.14 | 0.0204 | #NUM! | FOS,GNG3 |
| UVA-Induced MAPK Signaling | 1.14 | 0.0204 | #NUM! | FOS,HRAS |

| | | | | |
|---|---|---|---|---|
| Metabolism of Angiotensinogen to Angiotensins | 1.13 | 0.0588 | #NUM! | CTSD |
| Signaling by Type 1 Insulin-like Growth Factor 1 Receptor (IGF1R) | 1.13 | 0.0588 | #NUM! | HRAS |
| Sumoylation Pathway | 1.12 | 0.0198 | #NUM! | ARHGDIG,FOS |
| ERK/MAPK Signaling | 1.11 | 0.0136 | #NUM! | FOS,HRAS,HSPB1 |
| Metabolism of amine-derived hormones | 1.11 | 0.0556 | #NUM! | DBH |
| Gastrin-CREB signalling pathway via PKC and MAPK | 1.11 | 0.0556 | #NUM! | HRAS |
| Response of EIF2AK4 (GCN2) to amino acid deficiency | 1.1 | 0.0194 | #NUM! | CEBPB,FAU |
| Thrombin Signaling | 1.1 | 0.0135 | #NUM! | GNG3,HRAS,ITPR1 |
| IGF-1 Signaling | 1.09 | 0.019 | #NUM! | FOS,HRAS |
| Glycation Signaling Pathway | 1.09 | 0.0133 | #NUM! | FOS,HBA2,HRAS |
| WNK Renal Signaling Pathway | 1.07 | 0.0187 | #NUM! | CASR,NPPC |
| Post-translational protein phosphorylation | 1.07 | 0.0187 | #NUM! | CHGB,PENK |
| Role of Osteoblasts, Osteoclasts and Chondrocytes in Rheumatoid Arthritis | 1.07 | 0.0131 | #NUM! | FOS,TRAF5,WIF1 |
| Role of NFAT in Cardiac Hypertrophy | 1.07 | 0.0131 | #NUM! | GNG3,HRAS,ITPR1 |
| Synthesis, secretion, and deacylation of Ghrelin | 1.06 | 0.05 | #NUM! | GCG |
| EIF2 Signaling | 1.03 | 0.0126 | #NUM! | FAU,HRAS,NKX6-2 |
| Osteoarthritis Pathway | 1.03 | 0.0126 | #NUM! | C1QTNF4,CASR,CEBPB |
| Cardiac Hypertrophy Signaling (Enhanced) | 1.02 | 0.00931 | 0 | GNG3,HRAS,HSPB1,ITPR1,TG |
| Orexin Signaling Pathway | 1.01 | 0.0124 | #NUM! | GCG,GNG3,ITPR1 |

| | | | | |
|---|---|---|---|---|
| IL-12 Signaling and Production in Macrophages | 1.01 | 0.0124 | #NUM! | CEBPB,CLU,FOS |
| Sleep NREM Signaling Pathway | 1.01 | 0.0172 | #NUM! | GABRA6,HRAS |
| Pancreatic Secretion Signaling Pathway | 1.01 | 0.0123 | #NUM! | ARHGDIG,ITPR1,LIPC |
| Amyotrophic Lateral Sclerosis Signaling | 1.01 | 0.0171 | #NUM! | NEFL,PRPH |
| ESR-mediated signaling | 1 | 0.0169 | #NUM! | CTSD,FOS |
| Signaling by ROBO receptors | 0.999 | 0.0122 | #NUM! | FAU,ISL1,PSMB5 |
| Neddylation | 0.999 | 0.0122 | #NUM! | FBXO2,MUL1,PSMB5 |
| MSP-RON Signaling in Macrophages Pathway | 0.99 | 0.0167 | #NUM! | FOS,HRAS |
| RAS processing | 0.988 | 0.0417 | #NUM! | HRAS |
| Synaptic Long Term Potentiation | 0.966 | 0.0161 | #NUM! | HRAS,ITPR1 |
| Regulation of Insulin-like Growth Factor (IGF) transport and uptake by IGFBPs | 0.966 | 0.0161 | #NUM! | CHGB,PENK |
| Sperm Motility | 0.956 | 0.0117 | #NUM! | GNG3,ITPR1,NPPC |
| Signaling by Erythropoietin | 0.955 | 0.0385 | #NUM! | HRAS |
| IL-17A Signaling in Gastric Cells | 0.955 | 0.0385 | #NUM! | FOS |
| Hematoma Resolution Signaling Pathway | 0.949 | 0.0116 | #NUM! | FOS,NDUFA13,NDUFS7 |
| Endocannabinoid Developing Neuron Pathway | 0.944 | 0.0156 | #NUM! | GNG3,HRAS |
| Neutrophil Extracellular Trap Signaling Pathway | 0.943 | 0.00978 | -1 | ITPR1,NDUFA13,NDUFS7, TIMM13 |
| Cardiac Hypertrophy Signaling | 0.942 | 0.0115 | #NUM! | GNG3,HRAS,HSPB1 |
| Effects of PIP2 hydrolysis | 0.939 | 0.037 | #NUM! | ITPR1 |
| ATF4 activates genes in response to endoplasmic reticulum stress | 0.939 | 0.037 | #NUM! | CEBPB |
| Cardiogenesis | 0.939 | 0.037 | #NUM! | ISL1 |

| | | | | |
|---|---|---|---|---|
| Neuroprotective Role of THOP1 in Alzheimer's Disease | 0.933 | 0.0154 | #NUM! | NTS,TAC1 |
| HGF Signaling | 0.927 | 0.0153 | #NUM! | FOS,HRAS |
| 14-3-3-mediated Signaling | 0.922 | 0.0152 | #NUM! | FOS,HRAS |
| EGR2 and SOX10-mediated initiation of Schwann cell myelination | 0.91 | 0.0345 | #NUM! | POU3F1 |
| Glycolysis I | 0.91 | 0.0345 | #NUM! | GAPDH |
| Colorectal Cancer Metastasis Signaling | 0.906 | 0.0111 | #NUM! | FOS,GNG3,HRAS |
| Activation of kainate receptors upon glutamate binding | 0.897 | 0.0333 | #NUM! | GNG3 |
| FOXO-mediated transcription of oxidative stress, metabolic and neuronal genes | 0.897 | 0.0333 | #NUM! | NPY |
| Gluconeogenesis I | 0.897 | 0.0333 | #NUM! | GAPDH |
| Ferroptosis Signaling Pathway | 0.895 | 0.0146 | #NUM! | HRAS,HSPB1 |
| Adipogenesis pathway | 0.89 | 0.0145 | #NUM! | CEBPB,DLK1 |
| STAT3 Pathway | 0.885 | 0.0144 | #NUM! | HRAS,NDUFA13 |
| White Adipose Tissue Browning Pathway | 0.885 | 0.0144 | #NUM! | CEBPB,ITPR1 |
| G-protein beta:gamma signalling | 0.87 | 0.0312 | #NUM! | GNG3 |
| Thrombin signalling through proteinase activated receptors (PARs) | 0.87 | 0.0312 | #NUM! | GNG3 |
| Glucocorticoid Receptor Signaling | 0.868 | 0.00833 | #NUM! | FOS,HRAS,NDUFA13,NDUFS7,TAF7 |
| DHCR24 Signaling Pathway | 0.865 | 0.014 | #NUM! | CLU,HRAS |
| MSP-RON Signaling in Cancer Cells Pathway | 0.865 | 0.014 | #NUM! | FOS,HRAS |
| Role of Chondrocytes in Rheumatoid Arthritis Signaling Pathway | 0.86 | 0.0139 | #NUM! | CEBPB,FOS |
| CLEAR Signaling Pathway | 0.859 | 0.0105 | #NUM! | CTSD,HRAS,ITPR1 |
| Signal amplification | 0.858 | 0.0303 | #NUM! | GNG3 |

| | | | | |
|---|---|---|---|---|
| MAPK targets/ Nuclear events mediated by MAP kinases | 0.858 | 0.0303 | #NUM! | FOS |
| TNFR2 Signaling | 0.858 | 0.0303 | #NUM! | FOS |
| Selenoamino acid metabolism | 0.851 | 0.0137 | #NUM! | FAU,SEPHS2 |
| Gαi Signaling | 0.846 | 0.0136 | #NUM! | GNG3,HRAS |
| CREB Signaling in Neurons | 0.835 | 0.00812 | -1 | CASR,GNG3,HRAS,ITPR1, NPFFR2 |
| Transcriptional Regulation by NPAS4 | 0.834 | 0.0286 | #NUM! | FOS |
| Coagulation System | 0.834 | 0.0286 | #NUM! | F13B |
| Endocannabinoid Neuronal Synapse Pathway | 0.832 | 0.0133 | #NUM! | GNG3,ITPR1 |
| KEAP1-NFE2L2 pathway | 0.823 | 0.0132 | #NUM! | MUL1,PSMB5 |
| Type II Diabetes Mellitus Signaling | 0.823 | 0.0132 | #NUM! | CEBPB,ITPR1 |
| Transcriptional regulation by the AP-2 (TFAP2) family of transcription factors | 0.823 | 0.0278 | #NUM! | CITED4 |
| Sirtuin Signaling Pathway | 0.821 | 0.0101 | #NUM! | NDUFA13,NDUFS7,TIMM13 |
| Detoxification of Reactive Oxygen Species | 0.812 | 0.027 | #NUM! | ATOX1 |
| Corticotropin Releasing Hormone Signaling | 0.81 | 0.0129 | #NUM! | FOS,ITPR1 |
| Relaxin Signaling | 0.805 | 0.0128 | #NUM! | FOS,GNG3 |
| Formation of Fibrin Clot (Clotting Cascade) | 0.791 | 0.0256 | #NUM! | F13B |
| NGF-stimulated transcription | 0.791 | 0.0256 | #NUM! | FOS |
| FLT3 Signaling | 0.791 | 0.0256 | #NUM! | HRAS |
| HMGB1 Signaling | 0.788 | 0.0125 | #NUM! | FOS,HRAS |
| Serotonin Receptor Signaling | 0.787 | 0.00851 | -1 | F13B,GNG3,HRAS,SLC18A 3 |
| Neurotransmitter release cycle | 0.781 | 0.025 | #NUM! | SLC18A3 |

| | | | | |
|---|---|---|---|---|
| Class C/3 (Metabotropic glutamate/pheromone receptors) | 0.781 | 0.025 | #NUM! | CASR |
| Signaling by FGFR3 | 0.781 | 0.025 | #NUM! | HRAS |
| Transcriptional Regulation by VENTX | 0.781 | 0.025 | #NUM! | CEBPB |
| Transcriptional Regulatory Network in Embryonic Stem Cells | 0.771 | 0.0122 | #NUM! | HRAS,ISL1 |
| DAG and IP3 signaling | 0.771 | 0.0244 | #NUM! | ITPR1 |
| Signaling by FGFR4 | 0.771 | 0.0244 | #NUM! | HRAS |
| RET signaling | 0.771 | 0.0244 | #NUM! | NRTN |
| Parkinson's Signaling Pathway | 0.766 | 0.00949 | #NUM! | ITPR1,NDUFA13,NDUFS7 |
| Synaptogenesis Signaling Pathway | 0.766 | 0.00949 | #NUM! | HRAS,ITPR1,SNCG |
| Role of Osteoclasts in Rheumatoid Arthritis Signaling Pathway | 0.763 | 0.00946 | #NUM! | FOS,HRAS,TRAF5 |
| Pyrimidine Ribonucleotides Interconversion | 0.762 | 0.0238 | #NUM! | PGP |
| Signaling by SCF-KIT | 0.753 | 0.0233 | #NUM! | HRAS |
| DAP12 interactions | 0.753 | 0.0233 | #NUM! | HRAS |
| Vasopressin regulates renal water homeostasis via Aquaporins | 0.753 | 0.0233 | #NUM! | GNG3 |
| Oncostatin M Signaling | 0.753 | 0.0233 | #NUM! | HRAS |
| Intrinsic Prothrombin Activation Pathway | 0.753 | 0.0233 | #NUM! | F13B |
| Antigen Presentation Pathway | 0.753 | 0.0233 | #NUM! | PSMB5 |
| Gαq Signaling | 0.751 | 0.0118 | #NUM! | GNG3,ITPR1 |
| Aldosterone Signaling in Epithelial Cells | 0.744 | 0.0117 | #NUM! | HSPB1,ITPR1 |
| Glioblastoma Multiforme Signaling | 0.744 | 0.0117 | #NUM! | HRAS,ITPR1 |
| Assembly and cell surface presentation of NMDA receptors | 0.743 | 0.0227 | #NUM! | NEFL |
| Aspirin ADME | 0.743 | 0.0227 | #NUM! | BSG |
| Carboxyterminal post-translational modifications of tubulin | 0.735 | 0.0222 | #NUM! | TPGS1 |
| Pyrimidine Ribonucleotides De Novo Biosynthesis | 0.735 | 0.0222 | #NUM! | PGP |

| | | | | |
|---|---|---|---|---|
| Neuroinflammation Signaling Pathway | 0.728 | 0.00909 | #NUM! | FOS,GABRA6,NTF3 |
| MIF Regulation of Innate Immunity | 0.726 | 0.0217 | #NUM! | FOS |
| Dopamine-DARPP32 Feedback in cAMP Signaling | 0.717 | 0.0112 | #NUM! | CALY,ITPR1 |
| Tumor Microenvironment Pathway | 0.71 | 0.0111 | #NUM! | FOS,HRAS |
| G alpha (z) signalling events | 0.709 | 0.0208 | #NUM! | GNG3 |
| iNOS Signaling | 0.701 | 0.0204 | #NUM! | FOS |
| Apelin Muscle Signaling Pathway | 0.701 | 0.0204 | #NUM! | GNG3 |
| Ion channel transport | 0.699 | 0.0109 | #NUM! | CASQ2,FXYD7 |
| Signaling by ERBB2 | 0.694 | 0.02 | #NUM! | HRAS |
| Melanoma Signaling | 0.694 | 0.02 | #NUM! | HRAS |
| nNOS Signaling in Skeletal Muscle Cells | 0.694 | 0.02 | #NUM! | ITPR1 |
| Transcriptional regulation of granulopoiesis | 0.686 | 0.0196 | #NUM! | CEBPB |
| Assembly of RNA Polymerase II Complex | 0.686 | 0.0196 | #NUM! | TAF7 |
| Cell Cycle: G2/M DNA Damage Checkpoint Regulation | 0.686 | 0.0196 | #NUM! | RPRM |
| Regulation of eIF4 and p70S6K Signaling | 0.681 | 0.0106 | #NUM! | FAU,HRAS |
| Signaling by FGFR1 | 0.678 | 0.0192 | #NUM! | HRAS |
| TNFR1 Signaling | 0.678 | 0.0192 | #NUM! | FOS |
| Regulation of Apoptosis | 0.671 | 0.0189 | #NUM! | PSMB5 |
| Signaling by EGFR | 0.671 | 0.0189 | #NUM! | HRAS |
| Macrophage Alternative Activation Signaling Pathway | 0.668 | 0.0104 | #NUM! | CEBPB,FOS |
| Production of Nitric Oxide and Reactive Oxygen Species in Macrophages | 0.668 | 0.0104 | #NUM! | CLU,FOS |
| Retinol Biosynthesis | 0.664 | 0.0185 | #NUM! | LIPC |
| UVB-Induced MAPK Signaling | 0.664 | 0.0185 | #NUM! | FOS |
| Regulation of the Epithelial Mesenchymal Transition by Growth Factors Pathway | 0.661 | 0.0103 | #NUM! | FOS,HRAS |
| G-Protein Coupled Receptor Signaling | 0.658 | 0.00701 | -2.236 | CASR,FOS,GNG3,HRAS,NPFFR2 |

| | | | | |
|---|---|---|---|---|
| Meiotic synapsis | 0.657 | 0.0182 | #NUM! | SYCP1 |
| Signaling by PTK6 | 0.657 | 0.0182 | #NUM! | HRAS |
| Somitogenesis | 0.657 | 0.0182 | #NUM! | PSMB5 |
| Lymphotoxin β Receptor Signaling | 0.657 | 0.0182 | #NUM! | TRAF5 |
| CSDE1 Signaling Pathway | 0.65 | 0.0179 | #NUM! | FOS |
| CNTF Signaling | 0.65 | 0.0179 | #NUM! | HRAS |
| FAT10 Signaling Pathway | 0.65 | 0.0179 | #NUM! | PSMB5 |
| Acetylcholine Receptor Signaling Pathway | 0.648 | 0.0101 | #NUM! | ITPR1,SLC18A3 |
| Adrenergic Receptor Signaling Pathway (Enhanced) | 0.648 | 0.0101 | #NUM! | DBH,ITPR1 |
| TCF dependent signaling in response to WNT | 0.648 | 0.0101 | #NUM! | PSMB5,WIF1 |
| Human Embryonic Stem Cell Pluripotency | 0.642 | 0.01 | #NUM! | HRAS,NTF3 |
| Pulmonary Healing Signaling Pathway | 0.639 | 0.00995 | #NUM! | DLK1,HRAS |
| Signaling by PDGF | 0.637 | 0.0172 | #NUM! | HRAS |
| Cancer Drug Resistance by Drug Efflux | 0.637 | 0.0172 | #NUM! | HRAS |
| Ephrin Receptor Signaling | 0.636 | 0.0099 | #NUM! | GNG3,HRAS |
| Cachexia Signaling Pathway | 0.636 | 0.00815 | #NUM! | CEBPB,NPY,PSMB5 |
| Signaling by ERBB4 | 0.63 | 0.0169 | #NUM! | HRAS |
| Metabolism of polyamines | 0.63 | 0.0169 | #NUM! | PSMB5 |
| Polyamine Regulation in Colon Cancer | 0.63 | 0.0169 | #NUM! | FOS |
| NIK-->noncanonical NF-kB signaling | 0.624 | 0.0167 | #NUM! | PSMB5 |
| Endometrial Cancer Signaling | 0.624 | 0.0167 | #NUM! | HRAS |
| Role of Tissue Factor in Cancer | 0.617 | 0.00962 | #NUM! | FOS,HRAS |
| Thyroid Hormone Metabolism II (via Conjugation and/or Degradation) | 0.611 | 0.0161 | #NUM! | B4GAT1 |
| Coronavirus Pathogenesis Pathway | 0.608 | 0.00948 | #NUM! | FAU,FOS |
| Activin Inhibin Signaling Pathway | 0.605 | 0.00943 | #NUM! | CEBPB,FOS |
| Triacylglycerol Degradation | 0.605 | 0.0159 | #NUM! | LIPC |
| Cell surface interactions at the vascular wall | 0.6 | 0.00935 | #NUM! | BSG,HRAS |

| | | | | |
|---|---|---|---|---|
| mTOR Signaling | 0.6 | 0.00935 | #NUM! | FAU,HRAS |
| Collagen degradation | 0.599 | 0.0156 | #NUM! | CTSD |
| ERB2-ERBB3 Signaling | 0.599 | 0.0156 | #NUM! | HRAS |
| Hedgehog ligand biogenesis | 0.594 | 0.0154 | #NUM! | PSMB5 |
| Calcium Signaling | 0.588 | 0.00917 | #NUM! | CASQ2,ITPR1 |
| Autophagy | 0.585 | 0.00913 | #NUM! | FOS,GCG |
| RHOGDI Signaling | 0.583 | 0.00909 | #NUM! | ARHGDIG,GNG3 |
| ERBB4 Signaling | 0.582 | 0.0149 | #NUM! | HRAS |
| SPINK1 General Cancer Pathway | 0.582 | 0.0149 | #NUM! | HRAS |
| TNFR2 non-canonical NF-kB pathway | 0.577 | 0.0147 | #NUM! | PSMB5 |
| Agrin Interactions at Neuromuscular Junction | 0.577 | 0.0147 | #NUM! | HRAS |
| Phospholipases | 0.577 | 0.0147 | #NUM! | LIPC |
| Glutamate Receptor Signaling | 0.577 | 0.0147 | #NUM! | GNG3 |
| Protein Kinase A Signaling | 0.577 | 0.00758 | #NUM! | GNG3,ITPR1,PGP |
| Role of JAK1 and JAK3 in γc Cytokine Signaling | 0.571 | 0.0145 | #NUM! | HRAS |
| TP53 Regulates Transcription of DNA Repair Genes | 0.566 | 0.0143 | #NUM! | FOS |
| GM-CSF Signaling | 0.566 | 0.0143 | #NUM! | HRAS |
| Glioma Invasiveness Signaling | 0.561 | 0.0141 | #NUM! | HRAS |
| Regulation of RUNX2 expression and activity | 0.556 | 0.0139 | #NUM! | PSMB5 |
| Signaling by FGFR2 | 0.551 | 0.0137 | #NUM! | HRAS |
| Growth Hormone Signaling | 0.551 | 0.0137 | #NUM! | FOS |
| Neurovascular Coupling Signaling Pathway | 0.551 | 0.00862 | #NUM! | GABRA6,ITPR1 |
| Nicotine Degradation III | 0.541 | 0.0133 | #NUM! | B4GAT1 |
| Sertoli Cell-Germ Cell Junction Signaling Pathway (Enhanced) | 0.54 | 0.00847 | #NUM! | FOS,HRAS |
| Hepatic Fibrosis Signaling Pathway | 0.538 | 0.00721 | #NUM! | CEBPB,FOS,HRAS |
| Cellular response to hypoxia | 0.536 | 0.0132 | #NUM! | PSMB5 |
| Plasma lipoprotein assembly, remodeling, and clearance | 0.536 | 0.0132 | #NUM! | LIPC |

| | | | | |
|---|---|---|---|---|
| Macropinocytosis Signaling | 0.536 | 0.0132 | #NUM! | HRAS |
| Breast Cancer Regulation by Stathmin1 | 0.534 | 0.0066 | #NUM! | CASR,GNG3,HRAS,NPFFR2 |
| Signaling by NOTCH1 | 0.531 | 0.013 | #NUM! | DLK1 |
| SUMOylation of DNA damage response and repair proteins | 0.531 | 0.013 | #NUM! | NSMCE3 |
| Angiopoietin Signaling | 0.531 | 0.013 | #NUM! | HRAS |
| Leptin Signaling in Obesity | 0.531 | 0.013 | #NUM! | NPY |
| Toll-like Receptor Signaling | 0.531 | 0.013 | #NUM! | FOS |
| Role of Osteoblasts in Rheumatoid Arthritis Signaling Pathway | 0.53 | 0.00833 | #NUM! | CTSD,WIF1 |
| NF-κB Activation by Viruses | 0.526 | 0.0128 | #NUM! | HRAS |
| Maturity Onset Diabetes of Young (MODY) Signaling | 0.526 | 0.0128 | #NUM! | GAPDH |
| Signaling by MET | 0.522 | 0.0127 | #NUM! | HRAS |
| G alpha (12/13) signalling events | 0.517 | 0.0125 | #NUM! | GNG3 |
| Degradation of the extracellular matrix | 0.513 | 0.0123 | #NUM! | BSG |
| Signaling by NTRK1 (TRKA) | 0.513 | 0.0123 | #NUM! | HRAS |
| Role of JAK family kinases in IL-6-type Cytokine Signaling | 0.513 | 0.0123 | #NUM! | FOS |
| Cyclophilin Signaling Pathway | 0.509 | 0.00803 | #NUM! | BSG,TIMM13 |
| FLT3 Signaling in Hematopoietic Progenitor Cells | 0.508 | 0.0122 | #NUM! | HRAS |
| RAR Activation | 0.508 | 0.00693 | #NUM! | DLK1,FOS,SCAND1 |
| Transport of bile salts and organic acids, metal ions and amine compounds | 0.5 | 0.0119 | #NUM! | BSG |
| Role of MAPK Signaling in the Pathogenesis of Influenza | 0.5 | 0.0119 | #NUM! | HRAS |
| PEDF Signaling | 0.5 | 0.0119 | #NUM! | HRAS |
| Integrin cell surface interactions | 0.496 | 0.0118 | #NUM! | BSG |
| Melatonin Degradation I | 0.496 | 0.0118 | #NUM! | B4GAT1 |

| | | | | |
|---|---|---|---|---|
| Nicotine Degradation II | 0.496 | 0.0118 | #NUM! | B4GAT1 |
| BAG2 Signaling Pathway | 0.491 | 0.0116 | #NUM! | PSMB5 |
| ABRA Signaling Pathway | 0.483 | 0.0114 | #NUM! | FOS |
| Oxidative Stress Induced Senescence | 0.483 | 0.0114 | #NUM! | FOS |
| Regulation of mitotic cell cycle | 0.483 | 0.0114 | #NUM! | PSMB5 |
| Amyloid fiber formation | 0.483 | 0.0114 | #NUM! | CALCA |
| Regulation of Cellular Mechanics by Calpain Protease | 0.479 | 0.0112 | #NUM! | HRAS |
| Ribosomal Quality Control Signaling Pathway | 0.476 | 0.00758 | #NUM! | FAU,PSMB5 |
| Tuberculosis Latent Signaling Pathway | 0.475 | 0.0111 | #NUM! | TRAF5 |
| BMP signaling pathway | 0.475 | 0.0111 | #NUM! | HRAS |
| Deubiquitination | 0.473 | 0.00755 | #NUM! | MUL1,PSMB5 |
| Degradation of beta-catenin by the destruction complex | 0.471 | 0.011 | #NUM! | PSMB5 |
| Actin Nucleation by ARP-WASP Complex | 0.471 | 0.011 | #NUM! | HRAS |
| Superpathway of Melatonin Degradation | 0.471 | 0.011 | #NUM! | B4GAT1 |
| Signaling by Rho Family GTPases | 0.469 | 0.00749 | #NUM! | FOS,GNG3 |
| Acute Myeloid Leukemia Signaling | 0.468 | 0.0109 | #NUM! | HRAS |
| Unfolded protein response | 0.468 | 0.0109 | #NUM! | CEBPB |
| EPH-Ephrin signaling | 0.464 | 0.0108 | #NUM! | HRAS |
| Regulation of TP53 Activity through Phosphorylation | 0.464 | 0.0108 | #NUM! | TAF7 |
| Post-translational modification: synthesis of GPI-anchored proteins | 0.464 | 0.0108 | #NUM! | LY6H |
| NRF2-mediated Oxidative Stress Response | 0.463 | 0.00741 | #NUM! | FOS,HRAS |
| Molecular Mechanisms of Cancer | 0.458 | 0.00581 | -2.236 | CASR,FOS,GNG3,HRAS,NPFFR2 |
| Serotonin Degradation | 0.456 | 0.0105 | #NUM! | B4GAT1 |
| Transcriptional regulation by RUNX3 | 0.453 | 0.0104 | #NUM! | PSMB5 |
| Death Receptor Signaling | 0.449 | 0.0103 | #NUM! | HSPB1 |
| VEGF Signaling | 0.449 | 0.0103 | #NUM! | HRAS |

| | | | | |
|---|---|---|---|---|
| DNA Methylation and Transcriptional Repression Signaling | 0.446 | 0.0102 | #NUM! | CEBPB |
| Small Cell Lung Cancer Signaling | 0.446 | 0.0102 | #NUM! | TRAF5 |
| Protein Ubiquitination Pathway | 0.445 | 0.00717 | #NUM! | HSPB1,PSMB5 |
| Insulin Secretion Signaling Pathway | 0.445 | 0.00717 | #NUM! | GCG,ITPR1 |
| Protein folding | 0.442 | 0.0101 | #NUM! | GNG3 |
| Melanocyte Development and Pigmentation Signaling | 0.442 | 0.0101 | #NUM! | HRAS |
| Chronic Myeloid Leukemia Signaling | 0.441 | 0.00712 | #NUM! | FOS,HRAS |
| S Phase | 0.439 | 0.01 | #NUM! | PSMB5 |
| Apelin Cardiomyocyte Signaling Pathway | 0.439 | 0.01 | #NUM! | ITPR1 |
| Cellular response to heat stress | 0.435 | 0.0099 | #NUM! | HSPB1 |
| Potassium Channels | 0.428 | 0.00971 | #NUM! | GNG3 |
| ABC-family proteins mediated transport | 0.428 | 0.00971 | #NUM! | PSMB5 |
| GPER1 signaling | 0.428 | 0.00971 | #NUM! | GNG3 |
| Mouse Embryonic Stem Cell Pluripotency | 0.428 | 0.00971 | #NUM! | HRAS |
| Cellular Effects of Sildenafil (Viagra) | 0.428 | 0.00584 | 0 | CASR,GNG3,NPFFR2,NPPC |
| DNA Replication Pre-Initiation | 0.425 | 0.00962 | #NUM! | PSMB5 |
| Cargo recognition for clathrin-mediated endocytosis | 0.422 | 0.00952 | #NUM! | SLC18A3 |
| Sleep REM Signaling Pathway | 0.419 | 0.00943 | #NUM! | FOS |
| CDK5 Signaling | 0.419 | 0.00943 | #NUM! | HRAS |
| Paxillin Signaling | 0.419 | 0.00943 | #NUM! | HRAS |
| Apoptosis Signaling | 0.419 | 0.00943 | #NUM! | HRAS |
| Telomerase Signaling | 0.412 | 0.00926 | #NUM! | HRAS |
| Sphingolipid metabolism | 0.406 | 0.00909 | #NUM! | B3GALT4 |
| CCR5 Signaling in Macrophages | 0.404 | 0.00599 | #NUM! | FOS,GNG3,ITPR1 |
| Senescence Pathway | 0.404 | 0.00664 | #NUM! | CEBPB,HRAS |
| Phagosome Formation | 0.404 | 0.00567 | -1 | CASR,HRAS,ITPR1,NPFFR2 |
| Interleukin-4 and Interleukin-13 signaling | 0.403 | 0.00901 | #NUM! | FOS |

| | | | | |
|---|---|---|---|---|
| O-linked glycosylation | 0.397 | 0.00885 | #NUM! | B4GAT1 |
| Hedgehog 'off' state | 0.394 | 0.00877 | #NUM! | PSMB5 |
| Airway Pathology in Chronic Obstructive Pulmonary Disease | 0.394 | 0.00877 | #NUM! | PTGDS |
| Role of MAPK Signaling in Promoting the Pathogenesis of Influenza | 0.394 | 0.00877 | #NUM! | HRAS |
| Prostate Cancer Signaling | 0.388 | 0.00862 | #NUM! | HRAS |
| Nonsense-Mediated Decay (NMD) | 0.386 | 0.00855 | #NUM! | FAU |
| Axonal Guidance Signaling | 0.384 | 0.0058 | #NUM! | GNG3,HRAS,NTF3 |
| Neuregulin Signaling | 0.383 | 0.00847 | #NUM! | HRAS |
| PAK Signaling | 0.383 | 0.00847 | #NUM! | HRAS |
| Regulation of lipid metabolism by PPARalpha | 0.38 | 0.0084 | #NUM! | MED29 |
| Bladder Cancer Signaling | 0.38 | 0.0084 | #NUM! | HRAS |
| Fc Epsilon RI Signaling | 0.377 | 0.00833 | #NUM! | HRAS |
| Virus Entry via Endocytic Pathways | 0.377 | 0.00833 | #NUM! | HRAS |
| Nitric Oxide Signaling in the Cardiovascular System | 0.377 | 0.00833 | #NUM! | ITPR1 |
| NGF Signaling | 0.375 | 0.00826 | #NUM! | HRAS |
| Eukaryotic Translation Elongation | 0.372 | 0.0082 | #NUM! | FAU |
| Synthesis of DNA | 0.372 | 0.0082 | #NUM! | PSMB5 |
| Eukaryotic Translation Termination | 0.372 | 0.0082 | #NUM! | FAU |
| Role of NANOG in Mammalian Embryonic Stem Cell Pluripotency | 0.369 | 0.00813 | #NUM! | HRAS |
| p38 MAPK Signaling | 0.367 | 0.00806 | #NUM! | HSPB1 |
| Pulmonary Fibrosis Idiopathic Signaling Pathway | 0.363 | 0.00613 | #NUM! | FOS,HRAS |
| Glutaminergic Receptor Signaling Pathway (Enhanced) | 0.363 | 0.00613 | #NUM! | GABRA6,ITPR1 |
| TCR signaling | 0.361 | 0.00794 | #NUM! | PSMB5 |
| Gαs Signaling | 0.359 | 0.00787 | #NUM! | GNG3 |
| Glycosaminoglycan metabolism | 0.356 | 0.00781 | #NUM! | B4GAT1 |
| MHC class II antigen presentation | 0.356 | 0.00781 | #NUM! | CTSD |
| ROBO SLIT Signaling Pathway | 0.356 | 0.00781 | #NUM! | FOS |

| | | | | |
|---|---|---|---|---|
| Glioma Signaling | 0.356 | 0.00781 | #NUM! | HRAS |
| GP6 Signaling Pathway | 0.356 | 0.00781 | #NUM! | ITPR1 |
| Xenobiotic Metabolism Signaling | 0.355 | 0.00604 | #NUM! | HRAS,SCAND1 |
| Interleukin-1 family signaling | 0.354 | 0.00775 | #NUM! | PSMB5 |
| Clathrin-mediated endocytosis | 0.354 | 0.00775 | #NUM! | SLC18A3 |
| Insulin Receptor Signaling | 0.354 | 0.00775 | #NUM! | HRAS |
| LXR/RXR Activation | 0.351 | 0.00769 | #NUM! | CLU |
| Gap Junction Signaling | 0.349 | 0.00597 | #NUM! | HRAS,ITPR1 |
| Mitotic G1 phase and G1/S transition | 0.347 | 0.00758 | #NUM! | PSMB5 |
| Response to elevated platelet cytosolic Ca2+ | 0.347 | 0.00758 | #NUM! | CLU |
| Gα12/13 Signaling | 0.342 | 0.00746 | #NUM! | HRAS |
| Complement cascade | 0.337 | 0.00735 | #NUM! | CLU |
| Atherosclerosis Signaling | 0.337 | 0.00735 | #NUM! | CLU |
| RAC Signaling | 0.335 | 0.0073 | #NUM! | HRAS |
| PKCθ Signaling in T Lymphocytes | 0.333 | 0.00536 | #NUM! | FOS,HRAS,ITPR1 |
| SNARE Signaling Pathway | 0.333 | 0.00725 | #NUM! | SNCG |
| Role of PKR in Interferon Induction and Antiviral Response | 0.333 | 0.00725 | #NUM! | FOS |
| Hereditary Breast Cancer Signaling | 0.333 | 0.00725 | #NUM! | HRAS |
| Ephrin A Signaling | 0.328 | 0.00714 | #NUM! | HRAS |
| Signaling by NOTCH4 | 0.326 | 0.00709 | #NUM! | PSMB5 |
| SRP-dependent cotranslational protein targeting to membrane | 0.324 | 0.00704 | #NUM! | FAU |
| Hedgehog 'on' state | 0.319 | 0.00694 | #NUM! | PSMB5 |
| Bone Mineralization Signaling Pathway | 0.319 | 0.00694 | #NUM! | CASR |
| PIP3 activates AKT signaling | 0.317 | 0.0069 | #NUM! | NTF3 |
| Iron homeostasis signaling pathway | 0.317 | 0.0069 | #NUM! | HBA2 |
| RNA Polymerase II Transcription | 0.307 | 0.00667 | #NUM! | TAF7 |
| PTEN Regulation | 0.307 | 0.00667 | #NUM! | PSMB5 |
| Eukaryotic Translation Initiation | 0.307 | 0.00667 | #NUM! | FAU |
| Dilated Cardiomyopathy Signaling Pathway | 0.305 | 0.00662 | #NUM! | ITPR1 |
| NAD Signaling Pathway | 0.305 | 0.00662 | #NUM! | CEBPB |

| | | | | |
|---|---|---|---|---|
| WNT/SHH Axonal Guidance Signaling Pathway | 0.305 | 0.00662 | #NUM! | ITPR1 |
| PTEN Signaling | 0.305 | 0.00662 | #NUM! | HRAS |
| Transcriptional regulation by RUNX1 | 0.303 | 0.00658 | #NUM! | PSMB5 |
| PI3K Signaling in B Lymphocytes | 0.3 | 0.00507 | #NUM! | FOS,HRAS,ITPR1 |
| IL-10 Signaling | 0.297 | 0.00645 | #NUM! | FOS |
| Epithelial Adherens Junction Signaling | 0.291 | 0.00633 | #NUM! | HRAS |
| Microautophagy Signaling Pathway | 0.289 | 0.00629 | #NUM! | PSMB5 |
| Ovarian Cancer Signaling | 0.289 | 0.00629 | #NUM! | HRAS |
| Class I MHC mediated antigen processing and presentation | 0.289 | 0.00525 | #NUM! | FBXO2,PSMB5 |
| Preeclampsia Signaling Pathway | 0.286 | 0.00621 | #NUM! | FOS |
| eNOS Signaling | 0.286 | 0.00621 | #NUM! | ITPR1 |
| Inhibition of ARE-Mediated mRNA Degradation Pathway | 0.284 | 0.00617 | #NUM! | PSMB5 |
| Necroptosis Signaling Pathway | 0.282 | 0.00613 | #NUM! | TIMM13 |
| Xenobiotic Metabolism General Signaling Pathway | 0.282 | 0.00613 | #NUM! | HRAS |
| Fcgamma receptor (FCGR) dependent phagocytosis | 0.28 | 0.0061 | #NUM! | ITPR1 |
| HOTAIR Regulatory Pathway | 0.278 | 0.00606 | #NUM! | WIF1 |
| Coronavirus Replication Pathway | 0.277 | 0.00602 | #NUM! | FAU |
| Cardiac β-adrenergic Signaling | 0.27 | 0.00588 | #NUM! | GNG3 |
| Germ Cell-Sertoli Cell Junction Signaling | 0.268 | 0.00585 | #NUM! | HRAS |
| Phagosome Maturation | 0.268 | 0.00585 | #NUM! | CTSD |
| WNT/β-catenin Signaling | 0.265 | 0.00578 | #NUM! | WIF1 |
| Ribonucleotide Reductase Signaling Pathway | 0.263 | 0.00575 | #NUM! | FOS |
| Netrin Signaling | 0.261 | 0.00571 | #NUM! | ITPR1 |
| Systemic Lupus Erythematosus in T Cell Signaling Pathway | 0.252 | 0.00465 | #NUM! | FOS,HRAS,ITPR1 |
| Tight Junction Signaling | 0.252 | 0.00552 | #NUM! | FOS |

| | | | | |
|---|---|---|---|---|
| Major pathway of rRNA processing in the nucleolus and cytosol | 0.244 | 0.00538 | #NUM! | FAU |
| MicroRNA Biogenesis Signaling Pathway | 0.243 | 0.00535 | #NUM! | HRAS |
| Hepatitis B Chronic Liver Pathogenesis Signaling Pathway | 0.241 | 0.00532 | #NUM! | HRAS |
| NOD1/2 Signaling Pathway | 0.234 | 0.00518 | #NUM! | FOS |
| Regulation of the Epithelial-Mesenchymal Transition Pathway | 0.233 | 0.00515 | #NUM! | HRAS |
| IL-33 Signaling Pathway | 0.231 | 0.00513 | #NUM! | FOS |
| FXR/RXR Activation | 0.231 | 0.00513 | #NUM! | LIPC |
| CDX Gastrointestinal Cancer Signaling Pathway | 0.228 | 0.00508 | #NUM! | FOS |
| ILK Signaling | 0.228 | 0.00508 | #NUM! | FOS |
| PPARα/RXRα Activation | 0.227 | 0.00505 | #NUM! | HRAS |
| Mitotic G2-G2/M phases | 0.224 | 0.005 | #NUM! | PSMB5 |
| PI3K/AKT Signaling | 0.224 | 0.005 | #NUM! | HRAS |
| ID1 Signaling Pathway | 0.222 | 0.00495 | #NUM! | HRAS |
| Natural Killer Cell Signaling | 0.22 | 0.00493 | #NUM! | HRAS |
| Clathrin-mediated Endocytosis Signaling | 0.213 | 0.00478 | #NUM! | CLU |
| HIF1α Signaling | 0.211 | 0.00476 | #NUM! | HRAS |
| Integrin Signaling | 0.211 | 0.00476 | #NUM! | HRAS |
| Xenobiotic Metabolism PXR Signaling Pathway | 0.198 | 0.00452 | #NUM! | SCAND1 |
| Pathogen Induced Cytokine Storm Signaling Pathway | 0 | 0.00262 | #NUM! | FOS |
| Actin Cytoskeleton Signaling | 0 | 0.00413 | #NUM! | HRAS |
| Chaperone Mediated Autophagy Signaling Pathway | 0 | 0.00157 | #NUM! | PSMB5 |
| LPS/IL-1 Mediated Inhibition of RXR Function | 0 | 0.00336 | #NUM! | LIPC |
| Mitotic Metaphase and Anaphase | 0 | 0.00424 | #NUM! | PSMB5 |
| Neutrophil degranulation | 0 | 0.0021 | #NUM! | CTSD |
| Cell Cycle Checkpoints | 0 | 0.00368 | #NUM! | PSMB5 |
| RHO GTPase cycle | 0 | 0.00222 | #NUM! | ARHGDIG |
| Hepatic Cholestasis | 0 | 0.00444 | #NUM! | GCG |
| Autism Signaling Pathway | 0 | 0.00324 | #NUM! | HRAS |

| | | | | |
|---|---|---|---|---|
| NAFLD Signaling Pathway | 0 | 0.00446 | #NUM! | FOS |
| BBSome Signaling Pathway | 0 | 0.00406 | #NUM! | CASR,NPFFR2 |
| Cohesin Chromatin Regulation Pathway | 0 | 0.0038 | #NUM! | MED29 |
| Lung Ionic Balance Signaling Pathway | 0 | 0.00383 | #NUM! | CASR,NPFFR2 |
| TRIM21 Intracellular Antibody Signaling Pathway | 0 | 0.00353 | #NUM! | FOS,PSMB5 |
| ID3 Signaling Pathway | 0 | 0.00103 | #NUM! | HRAS |
| Tuberculosis Active Signaling Pathway | 0 | 0.00408 | #NUM! | FOS |
| Role of NFAT in Regulation of the Immune Response | 0 | 0.00382 | #NUM! | FOS,GNG3,HRAS,ITPR1 |
| FcγRIIB Signaling in B Lymphocytes | 0 | 0.00375 | #NUM! | HRAS,ITPR1 |
| Calcium-induced T Lymphocyte Apoptosis | 0 | 0.00215 | #NUM! | ITPR1 |
| CTLA4 Signaling in Cytotoxic T Lymphocytes | 0 | 0.00328 | #NUM! | FOS,HRAS |
| CD28 Signaling in T Helper Cells | 0 | 0.00383 | #NUM! | FOS,ITPR1 |
| IL-15 Signaling | 0 | 0.0019 | #NUM! | HRAS |
| Docosahexaenoic Acid (DHA) Signaling | 0 | 0.00398 | #NUM! | ITPR1 |
| ICOS-ICOSL Signaling in T Helper Cells | 0 | 0.00195 | #NUM! | ITPR1 |
| Autoimmune Thyroid Disease Signaling | 0 | 0.00216 | #NUM! | TG |
| p70S6K Signaling | 0 | 0.00172 | #NUM! | HRAS |
| G Protein Signaling Mediated by Tubby | 0 | 0.00213 | #NUM! | GNG3 |
| Systemic Lupus Erythematosus Signaling | 0 | 0.00187 | #NUM! | FOS,HRAS |
| CDC42 Signaling | 0 | 0.00173 | #NUM! | FOS |
| FAK Signaling | 0 | 0.00386 | -2 | CASR,FOS,HRAS,NPFFR2 |
| AMPK Signaling | 0 | 0.0041 | #NUM! | GNG3 |
| Phospholipase C Signaling | 0 | 0.00267 | #NUM! | GNG3,HRAS,ITPR1 |
| Regulation of IL-2 Expression in Activated and Anergic T Lymphocytes | 0 | 0.00428 | #NUM! | FOS,HRAS |
| OX40 Signaling Pathway | 0 | 0.00207 | #NUM! | TRAF5 |

| | | | | |
|---|---|---|---|---|
| Sertoli Cell-Sertoli Cell Junction Signaling | 0 | 0.00405 | #NUM! | HRAS |
| TEC Kinase Signaling | 0 | 0.00346 | #NUM! | FOS,GNG3 |
| SAPK/JNK Signaling | 0 | 0.00398 | #NUM! | GNG3,HRAS |
| IL-4 Signaling | 0 | 0.00174 | #NUM! | HRAS |
| B Cell Receptor Signaling | 0 | 0.00157 | #NUM! | HRAS |
| NF-κB Signaling | 0 | 0.0035 | #NUM! | HRAS,TRAF5 |
| T Cell Receptor Signaling | 0 | 0.00322 | #NUM! | FOS,HRAS |
| T Cell Exhaustion Signaling Pathway | 0 | 0.00353 | #NUM! | FOS,HRAS |
| Systemic Lupus Erythematosus in B Cell Signaling Pathway | 0 | 0.00412 | #NUM! | FOS,HRAS,TRAF5 |
| Xenobiotic Metabolism CAR Signaling Pathway | 0 | 0.00429 | #NUM! | SCAND1 |
