## Supplemental Table 2 for "Early Life Adversity Produces Enduring Molecular and Functional Disruption of Developing Vagal Circuits"

© 2000-2025 QIAGEN. All rights reserved.

**P9 EXCITATORY MALE NEURONS**

| Ingenuity Canonical Pathways | -log(p-value) | Ratio | z-score | Molecules |
|---|---|---|---|---|
| MyD88 cascade initiated on plasma membrane | 4.26 | 0.16 | 0 | PELI2,UBA52,UBB,UBE2V1 |
| Hematoma Resolution Signaling Pathway | 4.11 | 0.0386 | -1 | C7,CCL17,F2,FTH1,FTL,IL2,MC4R,NDUFA4L2,SOCS3,TUBA1A |
| Formation of the anterior neural plate | 4.03 | 0.273 | #NUM! | OTX2,PAX6,ZIC2 |
| MyD88 dependent cascade initiated on endosome | 3.88 | 0.129 | 0 | PELI2,UBA52,UBB,UBE2V1 |
| Iron uptake and transport | 3.81 | 0.0833 | 1.342 | FTH1,FTL,TF,UBA52,UBB |
| C-type lectin receptors (CLRs) | 3.62 | 0.0483 | 0.378 | CCL17,HRAS,ITPR1,UBA52,UBB,UBE2M,UBE2V1 |
| Signaling by FGFR3 | 3.45 | 0.1 | 1 | FGF20,HRAS,UBA52,UBB |
| Signaling by FGFR2 | 3.41 | 0.0685 | 1.342 | FGF20,HRAS,POLR2L,UBA52,UBB |
| Signaling by FGFR4 | 3.4 | 0.0976 | 1 | FGF20,HRAS,UBA52,UBB |
| MyD88:MAL(TIRAP) cascade initiated on plasma membrane | 3.36 | 0.0952 | 0 | PELI2,UBA52,UBB,UBE2V1 |
| Aggrephagy | 3.29 | 0.0909 | 0 | TUBA1A,UBA52,UBB,UBE2V1 |
| Signaling by NOTCH3 | 3.07 | 0.08 | #NUM! | HES5,PTCRA,UBA52,UBB |

| | | | | |
|---|---|---|---|---|
| Signaling by FGFR1 | 3.01 | 0.0769 | 1 | FGF20,HRAS,UBA52,UBB |
| RAS processing | 2.97 | 0.125 | #NUM! | HRAS,UBA52,UBB |
| Signaling by TGF-beta Receptor Complex | 2.95 | 0.0741 | 1 | LTBP2,UBA52,UBB,UBE2M |
| Signaling by PTK6 | 2.92 | 0.0727 | 1 | HRAS,SOCS3,UBA52,UBB |
| NLR signaling pathways | 2.89 | 0.0714 | 0 | NLRC4,UBA52,UBB,UBE2V1 |
| Interleukin-3, Interleukin-5 and GM-CSF signaling | 2.89 | 0.0714 | 1 | IL2,UBA52,UBB,VAV1 |
| Signaling by NTRK3 (TRKC) | 2.69 | 0.1 | #NUM! | BAX,HRAS,NTF3 |
| Signaling by CSF3 (G-CSF) | 2.69 | 0.1 | #NUM! | SOCS3,UBA52,UBB |
| Glutathione Redox Reactions I | 2.69 | 0.1 | #NUM! | GPX1,GPX4,Gstp1 (includes others) |
| NRF2-mediated Oxidative Stress Response | 2.66 | 0.0296 | 2 | ACTA1,FTH1,FTL,Gstm6,Gstp1 (includes others),HRAS,JUND,UBB |
| Endosomal Sorting Complex Required For Transport (ESCRT) | 2.65 | 0.0968 | #NUM! | UBA52,UBB,VPS25 |
| Toll Like Receptor 3 (TLR3) Cascade | 2.57 | 0.0909 | #NUM! | UBA52,UBB,UBE2V1 |
| Signaling by NOTCH2 | 2.53 | 0.0882 | #NUM! | HES5,UBA52,UBB |
| RUNX1 and FOXP3 control the development of regulatory T lymphocytes (Tregs) | 2.51 | 0.2 | #NUM! | CTLA4,IL2 |

| | | | | |
|---|---|---|---|---|
| Huntington's Disease Signaling | 2.49 | 0.0279 | -0.447 | BAX,BET1L,HRAS,ITPR1,NEUROD1,POLR2L,UBA52,UBB |
| Alpha-protein kinase 1 signaling pathway | 2.43 | 0.182 | #NUM! | UBA52,UBB |
| Pexophagy | 2.43 | 0.182 | #NUM! | UBA52,UBB |
| MyD88-independent TLR4 cascade | 2.39 | 0.0789 | #NUM! | UBA52,UBB,UBE2V1 |
| MAP kinase activation | 2.36 | 0.0769 | #NUM! | UBA52,UBB,UBE2V1 |
| FLT3 Signaling | 2.36 | 0.0769 | #NUM! | HRAS,UBA52,UBB |
| Erythropoietin Signaling Pathway | 2.35 | 0.0331 | 0 | CCND2,HBA2,Hbb-bs/Hbb-bt,HRAS,IL2,ITPR1 |
| Mitophagy | 2.33 | 0.075 | #NUM! | UBA52,UBB,UBE2V1 |
| Aryl Hydrocarbon Receptor Signaling | 2.32 | 0.0328 | -2 | BAX,CCND2,Gstm6,Gstp1 (includes others),MCM7,NFIB |
| Mitochondrial iron-sulfur cluster biogenesis | 2.28 | 0.154 | #NUM! | GLRX5,ISCA1 |
| Regulation of beta-cell development | 2.27 | 0.0714 | #NUM! | INSM1,NEUROD1,PAX6 |
| GABA Receptor Signaling | 2.26 | 0.0376 | #NUM! | GABRA6,GABRD,ITPR1,UBA52,UBB |
| TAK1-dependent IKK and NF-kappa-B activation | 2.21 | 0.0682 | #NUM! | UBA52,UBB,UBE2V1 |

| | | | | |
|---|---|---|---|---|
| Ferroptosis Signaling Pathway | 2.2 | 0.0365 | #NUM! | FTH1,FTL,GPX4,HRAS,TF |
| Hereditary Breast Cancer Signaling | 2.19 | 0.0362 | 0 | CCND2,HRAS,POLR2L,UBA52,UBB |
| DNA Damage Bypass | 2.11 | 0.0625 | #NUM! | UBA52,UBB,USP43 |
| Extrinsic Prothrombin Activation Pathway | 2.1 | 0.125 | #NUM! | F13A1,F2 |
| Iron homeostasis signaling pathway | 2.1 | 0.0345 | #NUM! | FTH1,FTL,HBA2,Hbb-bs/Hbb-bt,TF |
| Regulation of TP53 Activity through Phosphorylation | 2.09 | 0.043 | 1 | MAPKAPK5,TAF9,UBA52,UBB |
| TP53 Regulates Transcription of Cell Cycle Genes | 2.08 | 0.0612 | #NUM! | BAX,CNOT3,RGCC |
| Fc epsilon receptor (FCERI) signaling | 2.08 | 0.0291 | 0 | HRAS,ITPR1,UBA52,UBB,UBE2V1,VAV1 |
| Signaling by ERBB2 | 2.06 | 0.06 | #NUM! | HRAS,UBA52,UBB |
| Clathrin-mediated Endocytosis Signaling | 2.05 | 0.0287 | #NUM! | ACTA1,F2,FGF20,TF,UBA52,UBB |
| Transcriptional regulation by RUNX2 | 2.04 | 0.0588 | #NUM! | BAX,BGLAP,SATB2 |
| Gustation Pathway | 2.03 | 0.0284 | -2.236 | ABHD11,GABRA6,GABRD,GCG,ITPR1,PKD2L1 |
| Signaling by EGFR | 1.99 | 0.0566 | #NUM! | HRAS,UBA52,UBB |
| Regulation of TP53 Activity through Methylation | 1.96 | 0.105 | #NUM! | UBA52,UBB |

| | | | | |
|---|---|---|---|---|
| Deadenylation-dependent mRNA decay | 1.92 | 0.0536 | #NUM! | CNOT3,LSM2,LSM4 |
| Maturation of TCA enzymes and regulation of TCA cycle | 1.91 | 0.1 | #NUM! | ISCA1,SDHAF1 |
| TNF signaling | 1.9 | 0.0526 | #NUM! | SMPD3,UBA52,UBB |
| Signaling by ERBB4 | 1.86 | 0.0508 | #NUM! | HRAS,UBA52,UBB |
| Polyamine Regulation in Colon Cancer | 1.86 | 0.0508 | #NUM! | AKT1S1,EIF5A,OAZ1 |
| NIK-->noncanonical NF-kB signaling | 1.84 | 0.05 | #NUM! | UBA52,UBB,UBE2M |
| Chaperone Mediated Autophagy | 1.83 | 0.0909 | #NUM! | UBA52,UBB |
| Gamma carboxylation, hypusinylation, hydroxylation, and arylsulfatase activation | 1.82 | 0.0492 | #NUM! | BGLAP,EIF5A,F2 |
| Signaling by the B Cell Receptor (BCR) | 1.82 | 0.0294 | 1.342 | HRAS,ITPR1,UBA52,UBB,VAV1 |
| Cytosolic sensors of pathogen-associated DNA | 1.77 | 0.0469 | #NUM! | POLR2L,UBA52,UBB |
| Actin Cytoskeleton Signaling | 1.76 | 0.0248 | 2.236 | ACTA1,F2,FGF20,HRAS,PFN1,VAV1 |
| Parkinson's Signaling Pathway | 1.73 | 0.0222 | -1.633 | ACTA1,IL2,ITPR1,NDUFA4L2,TUBA1A,UBA52,UBB |
| Incretin synthesis, secretion, and inactivation | 1.73 | 0.08 | #NUM! | GCG,PAX6 |
| Glycogen metabolism | 1.73 | 0.08 | #NUM! | UBA52,UBB |
| Signaling by NTRK2 (TRKB) | 1.73 | 0.08 | #NUM! | HRAS,NTF3 |

| | | | | |
|---|---|---|---|---|
| Ion channel transport | 1.7 | 0.0273 | 0.447 | ANO2,ATP8B4,SLN,UBA52,UBB |
| Signaling by Erythropoietin | 1.69 | 0.0769 | #NUM! | HRAS,VAV1 |
| Role of JAK1 and JAK3 in γc Cytokine Signaling | 1.68 | 0.0435 | #NUM! | HRAS,IL2,SOCS3 |
| Acute Phase Response Signaling | 1.63 | 0.0262 | #NUM! | F2,FTL,HRAS,SOCS3,TF |
| Ephrin B Signaling | 1.62 | 0.0411 | #NUM! | ACP1,HRAS,VAV1 |
| Interleukin-1 family signaling | 1.62 | 0.031 | #NUM! | PELI2,UBA52,UBB,UBE2V1 |
| RAF/MAP kinase cascade | 1.61 | 0.0229 | -1 | DUSP5,FGF20,HRAS,IL2,UBA52,UBB |
| Glycolysis I | 1.6 | 0.069 | #NUM! | ENO1,Eno1b |
| Metabolism of vitamin K | 1.6 | 0.333 | #NUM! | VKORC1 |
| Spermine Biosynthesis | 1.6 | 0.333 | #NUM! | Amd2 |
| Hypusine Biosynthesis | 1.6 | 0.333 | #NUM! | EIF5A |
| Spermidine Biosynthesis I | 1.6 | 0.333 | #NUM! | Amd2 |
| trans-Golgi Network Vesicle Budding | 1.59 | 0.04 | #NUM! | BLOC1S1,FTH1,FTL |
| Mitotic G1 phase and G1/S transition | 1.58 | 0.0303 | #NUM! | CCND2,TOP2A,UBA52,UBB |
| 14-3-3-mediated Signaling | 1.58 | 0.0303 | #NUM! | AKT1S1,BAX,HRAS,TUBA1A |
| Protein ubiquitination | 1.58 | 0.0667 | #NUM! | UBA52,UBB |
| Regulation of CDH11 Expression and Function | 1.58 | 0.0667 | #NUM! | BHLHE22,PRDM8 |
| Gluconeogenesis I | 1.58 | 0.0667 | #NUM! | ENO1,Eno1b |

| | | | | |
|---|---|---|---|---|
| Plasma lipoprotein assembly, remodeling, and clearance | 1.57 | 0.0395 | #NUM! | PCSK9,UBA52,UBB |
| Interferon alpha/beta signaling | 1.57 | 0.0395 | #NUM! | SOCS3,UBA52,UBB |
| GPCR-Mediated Integration of Enteroendocrine Signaling Exemplified by an L Cell | 1.57 | 0.0395 | #NUM! | GAL,GCG,ITPR1 |
| Signaling by NOTCH1 | 1.56 | 0.039 | #NUM! | HES5,UBA52,UBB |
| Signaling by CSF1 (M-CSF) in myeloid cells | 1.55 | 0.0645 | #NUM! | UBA52,UBB |
| Renal Cell Carcinoma Signaling | 1.54 | 0.0385 | #NUM! | HRAS,UBA52,UBB |
| Signaling by MET | 1.53 | 0.038 | #NUM! | HRAS,UBA52,UBB |
| RIPK1-mediated regulated necrosis | 1.52 | 0.0625 | #NUM! | UBA52,UBB |
| Beta-catenin independent WNT signaling | 1.51 | 0.0244 | #NUM! | ITPR1,NME7,PFN1,UBA52,UBB |
| GABAergic Receptor Signaling Pathway (Enhanced) | 1.5 | 0.0286 | -1 | GABRA6,GABRD,ITPR1,KCNJ6 |
| Protein Ubiquitination Pathway | 1.5 | 0.0215 | #NUM! | UBA52,UBB,UBE2M,UBE2V1,USP43,Usp9y |
| Signaling by NOTCH4 | 1.49 | 0.0284 | 1 | HES5,NME7,UBA52,UBB |
| Late endosomal microautophagy | 1.48 | 0.0588 | #NUM! | UBA52,UBB |
| Activation of anterior HOX genes in hindbrain during early embryogenesis | 1.46 | 0.0357 | #NUM! | MAFB,PAX6,POLR2L |
| PIP3 activates AKT signaling | 1.46 | 0.0276 | 0 | AKT1S1,FGF20,NTF3,VAV1 |
| Oncogene Induced Senescence | 1.45 | 0.0571 | #NUM! | UBA52,UBB |

| | | | | |
|---|---|---|---|---|
| Coagulation System | 1.45 | 0.0571 | #NUM! | F13A1,F2 |
| FGF Signaling | 1.44 | 0.0349 | #NUM! | FGF20,HRAS,ITPR1 |
| Oxidative Stress Induced Senescence | 1.41 | 0.0341 | #NUM! | MAPKAPK5,UBA52,UBB |
| NAP1L1 Transcription Regulation Signaling Pathway | 1.4 | 0.0337 | #NUM! | BAX,PRDM8,TUBA1A |
| Fanconi Anemia Pathway | 1.39 | 0.0526 | #NUM! | UBA52,UBB |
| Regulation of TP53 Expression and Degradation | 1.39 | 0.0526 | #NUM! | UBA52,UBB |
| Reelin signalling pathway | 1.38 | 0.2 | #NUM! | RELN |
| CMP-N-acetylneuraminate Biosynthesis I (Eukaryotes) | 1.38 | 0.2 | #NUM! | NANP |
| Formation of Fibrin Clot (Clotting Cascade) | 1.37 | 0.0513 | #NUM! | F13A1,F2 |
| NGF-stimulated transcription | 1.37 | 0.0513 | #NUM! | CDK5R2,JUND |
| MAPK6/MAPK4 signaling | 1.36 | 0.0326 | #NUM! | MAPKAPK5,UBA52,UBB |
| p75 NTR receptor-mediated signalling | 1.34 | 0.0253 | 1 | NME7,UBA52,UBB,VAV1 |
| Apelin Adipocyte Signaling Pathway | 1.34 | 0.0319 | #NUM! | GPX1,GPX4,Gstp1 (includes others) |
| RNA Polymerase III Transcription | 1.33 | 0.0488 | #NUM! | NFIB,POLR2L |
| Mechanisms of Viral Exit from Host Cells | 1.33 | 0.0488 | #NUM! | ACTA1,VPS25 |
| Transcriptional regulation by RUNX3 | 1.32 | 0.0312 | #NUM! | TEAD2,UBA52,UBB |
| Non-Small Cell Lung Cancer Signaling | 1.32 | 0.0312 | #NUM! | CCND2,HRAS,ITPR1 |
| Pyrimidine Ribonucleotides Interconversion | 1.31 | 0.0476 | #NUM! | NME7,PGP |

| | | | | |
|---|---|---|---|---|
| Tryptophan Degradation to 2-amino-3-carboxymuconate Semialdehyde | 1.3 | 0.167 | #NUM! | KMO |
| Xenobiotic Metabolism General Signaling Pathway | 1.3 | 0.0245 | #NUM! | FTL,Gstm6,Gstp1 (includes others),HRAS |
| Signaling by SCF-KIT | 1.29 | 0.0465 | #NUM! | HRAS,VAV1 |
| Intrinsic Prothrombin Activation Pathway | 1.29 | 0.0465 | #NUM! | F13A1,F2 |
| Elastic fibre formation | 1.27 | 0.0455 | #NUM! | LTBP2,MFAP4 |
| TP53 Regulates Transcription of Cell Death Genes | 1.27 | 0.0455 | #NUM! | BAX,NLRC4 |
| Arachidonic acid metabolism | 1.25 | 0.0444 | #NUM! | GPX1,GPX4 |
| Pyrimidine Ribonucleotides De Novo Biosynthesis | 1.25 | 0.0444 | #NUM! | NME7,PGP |
| Pancreatic Secretion Signaling Pathway | 1.25 | 0.0206 | -1 | ABHD11,AKT1S1,BET1L,CADPS2,ITPR1 |
| DNA Replication Pre-Initiation | 1.23 | 0.0288 | #NUM! | MCM7,UBA52,UBB |
| Neddylation | 1.23 | 0.0203 | 0.447 | COMMD5,SOCS3,UBA52,UBB,UBE2M |
| WNT/β-catenin Signaling | 1.22 | 0.0231 | #NUM! | SOX8,UBA52,UBB,WIF1 |
| Cargo recognition for clathrin-mediated endocytosis | 1.22 | 0.0286 | #NUM! | TF,UBA52,UBB |
| Myelination Signaling Pathway | 1.22 | 0.0183 | 0.816 | AKT1S1,HES5,HRAS,MPZ,NTF3,SOX8 |
| Signaling by VEGF | 1.2 | 0.028 | #NUM! | HRAS,ITPR1,VAV1 |

| | | | | |
|---|---|---|---|---|
| Class A/1 (Rhodopsin-like receptors) | 1.2 | 0.0181 | -0.816 | CCL17,F2,GAL,MC4R,NPFF,SSTR1 |
| Pyruvate metabolism | 1.19 | 0.0408 | #NUM! | UBA52,UBB |
| Nucleotide Excision Repair | 1.18 | 0.0275 | #NUM! | POLR2L,UBA52,UBB |
| Sphingomyelin Metabolism | 1.18 | 0.125 | #NUM! | SMPD3 |
| Transcriptional activity of SMAD2/SMAD3:SMAD4 heterotrimer | 1.16 | 0.0392 | #NUM! | UBA52,UBB |
| Transcriptional and post-translational regulation of MITF-M expression and activity | 1.16 | 0.0392 | #NUM! | MC4R,ZIC1 |
| Assembly of RNA Polymerase II Complex | 1.16 | 0.0392 | #NUM! | POLR2L,TAF9 |
| Glutathione-mediated Detoxification | 1.16 | 0.0392 | #NUM! | Gstm6,Gstp1 (includes others) |
| UVC-Induced MAPK Signaling | 1.16 | 0.0392 | #NUM! | HRAS,SMPD3 |
| Hedgehog 'off' state | 1.14 | 0.0263 | #NUM! | TUBA1A,UBA52,UBB |
| Mitochondrial Dysfunction | 1.13 | 0.0174 | -0.447 | BAX,GPX1,GPX4,Gstp1 (includes others),ITPR1,NDUFA4L2 |
| Regulation of Apoptosis | 1.13 | 0.0377 | #NUM! | UBA52,UBB |
| Sleep NREM Signaling Pathway | 1.12 | 0.0259 | #NUM! | GABRA6,GABRD,HRAS |
| IL-17 Signaling | 1.11 | 0.0212 | -1 | CCL17,HRAS,IL2,TRAF5 |
| Intrinsic Pathway for Apoptosis | 1.1 | 0.0364 | #NUM! | BAX,NMT1 |
| Formation of the posterior neural plate | 1.09 | 0.1 | #NUM! | OTX2 |

| | | | | |
|---|---|---|---|---|
| Metabolism of non-coding RNA | 1.09 | 0.0357 | #NUM! | GEMIN4, GEMIN7 |
| E3 ubiquitin ligases ubiquitinate target proteins | 1.09 | 0.0357 | #NUM! | UBA52,UBB |
| EGF Signaling | 1.09 | 0.0357 | #NUM! | HRAS,ITPR1 |
| Leukocyte Extravasation Signaling | 1.08 | 0.0206 | #NUM! | ACTA1,CD99,CLDN18,VAV1 |
| NGF Signaling | 1.08 | 0.0248 | #NUM! | BAX,HRAS,SMPD3 |
| Cell Cycle Control of Chromosomal Replication | 1.08 | 0.0351 | #NUM! | MCM7,TOP2A |
| FXR/RXR Activation | 1.07 | 0.0205 | #NUM! | Gstm6,Gstp1 (includes others),IL2,SOCS3 |
| Synthesis of DNA | 1.07 | 0.0246 | #NUM! | MCM7,UBA52,UBB |
| Circadian Clock | 1.06 | 0.0345 | #NUM! | UBA52,UBB |
| Regulation of Insulin-like Growth Factor (IGF) transport and uptake by IGFBPs | 1.05 | 0.0242 | #NUM! | F2,PCSK9,TF |
| DNA Double Strand Break Response | 1.05 | 0.0339 | #NUM! | UBA52,UBB |
| NAD biosynthesis II (from tryptophan) | 1.05 | 0.0909 | #NUM! | KMO |
| NAD Phosphorylation and Dephosphorylation | 1.05 | 0.0909 | #NUM! | ACP1 |
| Asparagine N-linked glycosylation | 1.05 | 0.024 | #NUM! | DPM1,UBA52,UBB |
| Th1 Pathway | 1.05 | 0.024 | #NUM! | IL2,SOCS3,VAV1 |
| Mitotic G2-G2/M phases | 1.04 | 0.02 | 0 | NME7,TUBA1A,UBA52,UBB |
| TCR signaling | 1.04 | 0.0238 | #NUM! | UBA52,UBB,UBE2V1 |
| GABA receptor activation | 1.04 | 0.0333 | #NUM! | GABRA6, KCNJ6 |

| | | | | |
|---|---|---|---|---|
| PCP (Planar Cell Polarity) Pathway | 1.04 | 0.0333 | #NUM! | JUND,PFN1 |
| ID1 Signaling Pathway | 1.03 | 0.0198 | 1 | BAX,BHLHE22,HRAS,STMN3 |
| Endocannabinoid Developing Neuron Pathway | 1.02 | 0.0234 | #NUM! | AKT1S1,HRAS,PAX6 |
| Clathrin-mediated endocytosis | 1.02 | 0.0233 | #NUM! | TF,UBA52,UBB |
| Signaling by Leptin | 1.01 | 0.0833 | #NUM! | SOCS3 |
| Hematopoiesis from Multipotent Stem Cells | 1.01 | 0.0833 | #NUM! | IL2 |
| G Beta Gamma Signaling | 1.01 | 0.0231 | #NUM! | HRAS,ITPR1,KCNJ6 |
| IL-2 Signaling | 1 | 0.0317 | #NUM! | HRAS,IL2 |
| Costimulation by the CD28 family | 1 | 0.0229 | #NUM! | CTLA4,NME7,VAV1 |
| Response to elevated platelet cytosolic Ca2+ | 0.993 | 0.0227 | #NUM! | F13A1,PFN1,TF |
| Peroxisomal protein import | 0.991 | 0.0312 | #NUM! | UBA52,UBB |
| Platelet Aggregation (Plug Formation) | 0.981 | 0.0769 | #NUM! | F2 |
| Dolichyl-diphosphooligosaccharide Biosynthesis | 0.981 | 0.0769 | #NUM! | DPM1 |
| Hedgehog ligand biogenesis | 0.979 | 0.0308 | #NUM! | UBA52,UBB |
| Role of JAK2 in Hormone-like Cytokine Signaling | 0.979 | 0.0308 | #NUM! | CCND2,SOCS3 |
| Gα12/13 Signaling | 0.978 | 0.0224 | #NUM! | F2,HRAS,VAV1 |
| Pathogen Induced Cytokine Storm Signaling Pathway | 0.972 | 0.0157 | -1.342 | CCL17,FTH1,FTL,IL2,NLRC4,SOCS3 |
| NoRC negatively regulates rRNA expression | 0.957 | 0.0299 | #NUM! | POLR2L,SAP18 |
| Tryptophan catabolism | 0.95 | 0.0714 | #NUM! | KMO |
| Formation of axial mesoderm | 0.95 | 0.0714 | #NUM! | TEAD2 |
| TNFR2 non-canonical NF-kB pathway | 0.947 | 0.0294 | #NUM! | UBA52,UBB |

| | | | | |
|---|---|---|---|---|
| Agrin Interactions at Neuromuscular Junction | 0.947 | 0.0294 | #NUM! | ACTA1,HRAS |
| Multiple Sclerosis Signaling Pathway | 0.937 | 0.0183 | 1 | BAX,C7,CTLA4,IL2 |
| Ephrin A Signaling | 0.936 | 0.0214 | #NUM! | ACP1,HRAS,VAV1 |
| Th2 Pathway | 0.929 | 0.0213 | #NUM! | IL2,SOCS3,VAV1 |
| YAP1- and WWTR1 (TAZ)-stimulated gene expression | 0.922 | 0.0667 | #NUM! | TEAD2 |
| Remodeling of Epithelial Adherens Junctions | 0.916 | 0.0282 | #NUM! | ACTA1,TUBA1A |
| NAFLD Signaling Pathway | 0.911 | 0.0179 | 0 | AKT1S1,BAX,IL2,SOCS3 |
| Agranulocyte Adhesion and Diapedesis | 0.911 | 0.0179 | #NUM! | ACTA1,CCL17,CD99,CLDN18 |
| Hedgehog 'on' state | 0.909 | 0.0208 | #NUM! | NME7,UBA52,UBB |
| Regulation of RUNX2 expression and activity | 0.906 | 0.0278 | #NUM! | UBA52,UBB |
| G alpha (s) signalling events | 0.903 | 0.0207 | #NUM! | GCG,GPR150,MC4R |
| Endocannabinoid Cancer Inhibition Pathway | 0.89 | 0.0204 | #NUM! | AKT1S1,CCND2,SMPD3 |
| ISG15 antiviral mechanism | 0.887 | 0.027 | #NUM! | UBA52,UBB |
| Senescence-Associated Secretory Phenotype (SASP) | 0.887 | 0.027 | #NUM! | UBA52,UBB |
| Synaptogenesis Signaling Pathway | 0.883 | 0.0158 | -0.447 | BET1L,HRAS,ITPR1,RAB3A,RELN |
| Signaling by Type 1 Insulin-like Growth Factor 1 Receptor (IGF1R) | 0.871 | 0.0588 | #NUM! | HRAS |
| FOXO-mediated transcription of cell cycle genes | 0.871 | 0.0588 | #NUM! | BTG1 |
| Specification of the neural plate border | 0.871 | 0.0588 | #NUM! | ZIC1 |

| | | | | |
|---|---|---|---|---|
| RAN Signaling | 0.871 | 0.0588 | #NUM! | RCC1 |
| Neurovascular Coupling Signaling Pathway | 0.871 | 0.0172 | -1 | GABRA6, GABRD,ITPR1,KCNJ6 |
| RNA Polymerase II Transcription | 0.871 | 0.02 | #NUM! | LSM10,POLR2L,TAF9 |
| Cellular response to hypoxia | 0.868 | 0.0263 | #NUM! | UBA52,UBB |
| GDNF Family Ligand-Receptor Interactions | 0.868 | 0.0263 | #NUM! | HRAS,ITPR1 |
| Dilated Cardiomyopathy Signaling Pathway | 0.865 | 0.0199 | #NUM! | ACTA1,BAX,ITPR1 |
| Toll-like Receptor Signaling | 0.859 | 0.026 | #NUM! | UBA52,UBB |
| Transcriptional regulation by RUNX1 | 0.859 | 0.0197 | #NUM! | SOCS3,UBA52,UBB |
| KEAP1-NFE2L2 pathway | 0.859 | 0.0197 | #NUM! | NME7,UBA52,UBB |
| Type II Diabetes Mellitus Signaling | 0.859 | 0.0197 | #NUM! | ITPR1,SMPD3,SOCS3 |
| Mitotic Metaphase and Anaphase | 0.852 | 0.0169 | 1 | RCC1,TUBA1A,UBA52,UBB |
| VDR/RXR Activation | 0.85 | 0.0256 | #NUM! | BGLAP,IL2 |
| Thyroid Cancer Signaling | 0.85 | 0.0256 | #NUM! | HRAS,NTF3 |
| Neurotrophin/TRK Signaling | 0.85 | 0.0256 | #NUM! | HRAS,NTF3 |
| Hypoxia Signaling in the Cardiovascular System | 0.85 | 0.0256 | #NUM! | UBE2M,UBE2V1 |
| Gastrin-CREB signalling pathway via PKC and MAPK | 0.848 | 0.0556 | #NUM! | HRAS |
| Formation of the nephric duct | 0.848 | 0.0556 | #NUM! | WFDC2 |
| Glutaminergic Receptor Signaling Pathway (Enhanced) | 0.842 | 0.0153 | 0.447 | AKT1S1, BET1L,GABRA6,GABRD,ITPR1 |
| GABA synthesis, release, reuptake and degradation | 0.827 | 0.0526 | #NUM! | RAB3A |

| | | | | |
|---|---|---|---|---|
| Bile Acid Biosynthesis, Neutral Pathway | 0.827 | 0.0526 | #NUM! | AKR1D1 |
| Orexin Signaling Pathway | 0.824 | 0.0165 | 1 | AKT1S1,GCG,ITPR1,KCNJ6 |
| DDX58/IFIH1-mediated induction of interferon-alpha/beta | 0.824 | 0.0247 | #NUM! | UBA52,UBB |
| Axonal Guidance Signaling | 0.823 | 0.0135 | #NUM! | ADAM2,ADAMTS18,HRAS,NTF3,NTN4,PFN1,TUBA1A |
| Gap Junction Signaling | 0.808 | 0.0149 | 1 | ACTA1,BGLAP,HRAS,ITPR1,TUBA1A |
| RNA polymerase II transcribes snRNA genes | 0.807 | 0.0241 | #NUM! | POLR2L,TAF9 |
| JAK/STAT Signaling | 0.807 | 0.0241 | #NUM! | HRAS,SOCS3 |
| Synthesis, secretion, and deacylation of Ghrelin | 0.806 | 0.05 | #NUM! | GCG |
| Sertoli Cell-Sertoli Cell Junction Signaling | 0.802 | 0.0162 | 1 | ACTA1,CLDN18,HRAS,TUBA1A |
| Role of Macrophages, Fibroblasts and Endothelial Cells in Rheumatoid Arthritis | 0.801 | 0.0148 | 0.447 | HRAS,IL16,SOCS3,TRAF5,WIF1 |
| Role of MAPK Signaling in the Pathogenesis of Influenza | 0.799 | 0.0238 | #NUM! | BAX,HRAS |
| Activation of NMDA receptors and postsynaptic events | 0.791 | 0.0235 | #NUM! | HRAS,TUBA1A |
| Glucose metabolism | 0.783 | 0.0233 | #NUM! | ENO1,PGP |
| Mitochondrial Division Signaling Pathway | 0.779 | 0.0181 | #NUM! | BAX,HRAS,ITPR1 |
| PDGF Signaling | 0.775 | 0.023 | #NUM! | ACP1,HRAS |

| | | | | |
|---|---|---|---|---|
| ABRA Signaling Pathway | 0.768 | 0.0227 | #NUM! | ACTA1,JUND |
| Regulation of mitotic cell cycle | 0.768 | 0.0227 | #NUM! | UBA52,UBB |
| Amyloid fiber formation | 0.768 | 0.0227 | #NUM! | UBA52,UBB |
| Regulation of mRNA stability by proteins that bind AU-rich elements | 0.76 | 0.0225 | #NUM! | UBA52,UBB |
| Regulation of Cellular Mechanics by Calpain Protease | 0.76 | 0.0225 | #NUM! | CCND2,HRAS |
| Germ Cell-Sertoli Cell Junction Signaling | 0.753 | 0.0175 | #NUM! | ACTA1,HRAS,TUBA1A |
| Glioblastoma Multiforme Signaling | 0.753 | 0.0175 | #NUM! | CCND2,HRAS,ITPR1 |
| Phagosome Maturation | 0.753 | 0.0175 | #NUM! | BET1L,Dynlt1b (includes others),TUBA1A |
| BMP signaling pathway | 0.752 | 0.0222 | #NUM! | HRAS,SOSTDC1 |
| RAF-independent MAPK1/3 activation | 0.751 | 0.0435 | #NUM! | DUSP5 |
| Azathioprine ADME | 0.751 | 0.0435 | #NUM! | VAV1 |
| NAD Salvage Pathway II | 0.751 | 0.0435 | #NUM! | ACP1 |
| Degradation of beta-catenin by the destruction complex | 0.745 | 0.022 | #NUM! | UBA52,UBB |
| Ceramide Signaling | 0.745 | 0.022 | #NUM! | HRAS,SMPD3 |
| NER (Nucleotide Excision Repair, Enhanced Pathway) | 0.745 | 0.022 | #NUM! | POLR2L,TOP2A |
| Cell junction organization | 0.738 | 0.0217 | #NUM! | CLDN18,KRT14 |
| G alpha (q) signalling events | 0.737 | 0.0172 | #NUM! | F2,GCG,NPFF |
| Other interleukin signaling | 0.734 | 0.0417 | #NUM! | IL16 |
| Interleukin-6 family signaling | 0.734 | 0.0417 | #NUM! | SOCS3 |
| Regulation of RUNX1 Expression and Activity | 0.734 | 0.0417 | #NUM! | CCND2 |
| IL-22 Signaling | 0.734 | 0.0417 | #NUM! | SOCS3 |

| | | | | |
|---|---|---|---|---|
| Tumoricidal Function of Hepatic Natural Killer Cells | 0.734 | 0.0417 | #NUM! | BAX |
| Tryptophan Degradation III (Eukaryotic) | 0.734 | 0.0417 | #NUM! | KMO |
| Inflammasome pathway | 0.734 | 0.0417 | #NUM! | NLRC4 |
| Androgen Signaling | 0.732 | 0.0171 | #NUM! | ITPR1,POLR2L,TAF9 |
| Netrin Signaling | 0.732 | 0.0171 | #NUM! | BET1L,DRAXIN,ITPR1 |
| Post-translational modification: synthesis of GPI-anchored proteins | 0.731 | 0.0215 | #NUM! | DPM1,LY6G6D |
| Fcγ Receptor-mediated Phagocytosis in Macrophages and Monocytes | 0.731 | 0.0215 | #NUM! | ACTA1,VAV1 |
| Th1 and Th2 Activation Pathway | 0.727 | 0.017 | #NUM! | IL2,SOCS3,VAV1 |
| Heparan Sulfate Biosynthesis (Late Stages) | 0.724 | 0.0213 | #NUM! | AARSD1,GLCE |
| Growth hormone receptor signaling | 0.718 | 0.04 | #NUM! | SOCS3 |
| Prolactin Signaling | 0.71 | 0.0208 | #NUM! | HRAS,SOCS3 |
| Tumor Microenvironment Pathway | 0.708 | 0.0167 | #NUM! | CTLA4,FGF20,HRAS |
| Tight Junction Signaling | 0.703 | 0.0166 | #NUM! | ACTA1,BET1L,CLDN18 |
| VEGF Signaling | 0.703 | 0.0206 | #NUM! | ACTA1,HRAS |
| Fertilization | 0.703 | 0.0385 | #NUM! | ADAM2 |
| Interleukin-20 family signaling | 0.703 | 0.0385 | #NUM! | SOCS3 |
| Pyrimidine Deoxyribonucleotides De Novo Biosynthesis I | 0.703 | 0.0385 | #NUM! | NME7 |
| p53 Signaling | 0.696 | 0.0204 | #NUM! | BAX,CCND2 |
| Small Cell Lung Cancer Signaling | 0.696 | 0.0204 | #NUM! | CCND2,TRAF5 |

| | | | | |
|---|---|---|---|---|
| Crosstalk between Dendritic Cells and Natural Killer Cells | 0.696 | 0.0204 | #NUM! | ACTA1,IL2 |
| UVA-Induced MAPK Signaling | 0.696 | 0.0204 | #NUM! | HRAS,SMPD3 |
| Effects of PIP2 hydrolysis | 0.688 | 0.037 | #NUM! | ITPR1 |
| Pyroptosis | 0.688 | 0.037 | #NUM! | BAX |
| Cardiogenesis | 0.688 | 0.037 | #NUM! | TBX20 |
| S Phase | 0.683 | 0.02 | #NUM! | UBA52,UBB |
| Serotonin Receptor Signaling | 0.68 | 0.0128 | 0 | F13A1,HRAS,Htr5b,IL2,KCNJ6,RAB3A |
| Cellular response to heat stress | 0.677 | 0.0198 | #NUM! | AKT1S1,UBB |
| Heparan Sulfate Biosynthesis | 0.677 | 0.0198 | #NUM! | AARSD1,GLCE |
| MicroRNA Biogenesis Signaling Pathway | 0.676 | 0.016 | #NUM! | HOPX,HRAS,POLR2L |
| Insulin Secretion Signaling Pathway | 0.674 | 0.0143 | -2 | BET1L,GCG,ITPR1,NEUROD1 |
| MTOR signalling | 0.674 | 0.0357 | #NUM! | AKT1S1 |
| Pyroptosis Signaling Pathway | 0.67 | 0.0196 | #NUM! | BAX,NLRC4 |
| Gene Silencing by RNA | 0.67 | 0.0196 | #NUM! | DDX4,POLR2L |
| COPI-mediated anterograde transport | 0.67 | 0.0196 | #NUM! | BET1L,TUBA1A |
| Chronic Myeloid Leukemia Signaling | 0.667 | 0.0142 | 0 | AKT1S1,CCND2,HRAS,RCC1 |
| D-myo-inositol (1,4,5,6)-Tetrakisphosphate Biosynthesis | 0.667 | 0.0159 | #NUM! | ACP1,DUSP5,SOCS3 |
| D-myo-inositol (3,4,5,6)-tetrakisphosphate Biosynthesis | 0.667 | 0.0159 | #NUM! | ACP1,DUSP5,SOCS3 |
| Potassium Channels | 0.664 | 0.0194 | #NUM! | KCNG1,KCNJ6 |
| ABC-family proteins mediated transport | 0.664 | 0.0194 | #NUM! | UBA52,UBB |

| | | | | |
|---|---|---|---|---|
| Neutrophil degranulation | 0.661 | 0.0126 | 0 | ATP8B4, FTH1,FTL,PKP1,RAB3A,RNASET2 |
| Processing of Capped Intronless Pre-mRNA | 0.661 | 0.0345 | #NUM! | LSM10 |
| EGR2 and SOX10-mediated initiation of Schwann cell myelination | 0.661 | 0.0345 | #NUM! | MPZ |
| Class I MHC mediated antigen processing and presentation | 0.654 | 0.0131 | 0.447 | SOCS3,UBA52,UBB,UBE2M,UBE2V1 |
| Macrophage Alternative Activation Signaling Pathway | 0.654 | 0.0156 | #NUM! | AKT1S1, CCL17,SOCS3 |
| CLEAR Signaling Pathway | 0.653 | 0.014 | -1 | AKT1S1, BLOC1S1,HRAS,ITPR1 |
| IGF-1 Signaling | 0.652 | 0.019 | #NUM! | HRAS,SOCS3 |
| mRNA Capping | 0.648 | 0.0333 | #NUM! | POLR2L |
| Paxillin Signaling | 0.646 | 0.0189 | #NUM! | ACTA1,HRAS |
| Apoptosis Signaling | 0.646 | 0.0189 | #NUM! | BAX,HRAS |
| Processing of Capped Intron-Containing Pre-mRNA | 0.643 | 0.0139 | 2 | LSM2,LSM4,POLR2L,SAP18 |
| Post-translational protein phosphorylation | 0.64 | 0.0187 | #NUM! | PCSK9,TF |
| Transcriptional regulation of brown and beige adipocyte differentiation | 0.635 | 0.0323 | #NUM! | EBF2 |
| Integration of energy metabolism | 0.634 | 0.0185 | #NUM! | GCG,ITPR1 |
| Telomerase Signaling | 0.634 | 0.0185 | #NUM! | HRAS,IL2 |
| Xenobiotic Metabolism AHR Signaling Pathway | 0.628 | 0.0183 | #NUM! | Gstm6,Gstp1 (includes others) |
| Synaptic Long Term Depression | 0.628 | 0.0152 | #NUM! | HRAS,ITPR1,PPP1R17 |

| | | | | |
|---|---|---|---|---|
| TCF dependent signaling in response to WNT | 0.628 | 0.0152 | #NUM! | UBA52,UBB,WIF1 |
| Thrombin signalling through proteinase activated receptors (PARs) | 0.623 | 0.0312 | #NUM! | F2 |
| RHO GTPases activate IQGAPs | 0.623 | 0.0312 | #NUM! | TUBA1A |
| Chromatin modifications during the maternal to zygotic transition (MZT) | 0.623 | 0.0312 | #NUM! | METTL23 |
| 3-phosphoinositide Degradation | 0.62 | 0.015 | #NUM! | ACP1,DUSP5,SOCS3 |
| Interleukin-4 and Interleukin-13 signaling | 0.617 | 0.018 | #NUM! | F13A1,SOCS3 |
| Regulation of Actin-based Motility by Rho | 0.617 | 0.018 | #NUM! | ACTA1,PFN1 |
| α-Adrenergic Signaling | 0.617 | 0.018 | #NUM! | HRAS,ITPR1 |
| Sirtuin Signaling Pathway | 0.613 | 0.0135 | #NUM! | BAX,NDUFA4L2,TUBA1A,ZIC2 |
| Adrenomedullin signaling pathway | 0.612 | 0.0149 | #NUM! | BAX,HRAS,ITPR1 |
| Sialic acid metabolism | 0.612 | 0.0303 | #NUM! | NANP |
| Natural Killer Cell Signaling | 0.608 | 0.0148 | #NUM! | HRAS,IL2,VAV1 |
| Glucocorticoid Receptor Signaling | 0.608 | 0.0117 | #NUM! | BGLAP,HRAS,IL2,KRT14,NDUFA4L2,POLR2L,TAF9 |
| O-linked glycosylation | 0.606 | 0.0177 | #NUM! | ADAMTS18,GALNT15 |
| D-myo-inositol-5-phosphate Metabolism | 0.604 | 0.0147 | #NUM! | ACP1,DUSP5,SOCS3 |
| Granulocyte Adhesion and Diapedesis | 0.604 | 0.0147 | #NUM! | CCL17,CD99,CLDN18 |
| Binding and Uptake of Ligands by Scavenger Receptors | 0.6 | 0.0175 | #NUM! | FTH1,FTL |

| | | | | |
|---|---|---|---|---|
| HDR through Homologous Recombination (HRR) or Single Strand Annealing (SSA) | 0.6 | 0.0175 | #NUM! | UBA52,UBB |
| Airway Pathology in Chronic Obstructive Pulmonary Disease | 0.6 | 0.0175 | #NUM! | FGF20,IL2 |
| Role of MAPK Signaling in Promoting the Pathogenesis of Influenza | 0.6 | 0.0175 | #NUM! | BAX,HRAS |
| Breast Cancer Regulation by Stathmin1 | 0.595 | 0.0116 | 0.447 | CCND2,GPR150,HRAS,MC4R,Qrfpr,SSTR1,TUBA1A |
| GPVI-mediated activation cascade | 0.59 | 0.0286 | #NUM! | VAV1 |
| Activation of the pre-replicative complex | 0.59 | 0.0286 | #NUM! | MCM7 |
| IL-9 Signaling | 0.59 | 0.0286 | #NUM! | SOCS3 |
| IL-8 Signaling | 0.588 | 0.0144 | #NUM! | BAX,CCND2,HRAS |
| Amyotrophic Lateral Sclerosis Signaling | 0.584 | 0.0171 | #NUM! | BAX,GPX1 |
| Cholecystokinin/Gastrin-mediated Signaling | 0.584 | 0.0171 | #NUM! | HRAS,ITPR1 |
| Integrin Signaling | 0.581 | 0.0143 | #NUM! | ACTA1,HRAS,PFN1 |
| ESR-mediated signaling | 0.579 | 0.0169 | #NUM! | JUND,POLR2L |
| Striated Muscle Contraction | 0.579 | 0.0278 | #NUM! | ACTA1 |
| Autism Signaling Pathway | 0.575 | 0.0129 | 1 | AKT1S1,HRAS,IL2,RELN |
| Bladder Cancer Signaling | 0.574 | 0.0168 | #NUM! | FGF20,HRAS |
| GPCR-Mediated Nutrient Sensing in Enteroendocrine Cells | 0.574 | 0.0168 | #NUM! | GCG,ITPR1 |
| CREB Signaling in Neurons | 0.573 | 0.0114 | -0.447 | GPR150,HRAS,ITPR1,MC4R,POLR2L,Qrfpr,SSTR1 |

| | | | | |
|---|---|---|---|---|
| Fc Epsilon RI Signaling | 0.569 | 0.0167 | #NUM! | HRAS,VAV1 |
| Virus Entry via Endocytic Pathways | 0.569 | 0.0167 | #NUM! | ACTA1,HRAS |
| MSP-RON Signaling in Macrophages Pathway | 0.569 | 0.0167 | #NUM! | HRAS,SOCS3 |
| Detoxification of Reactive Oxygen Species | 0.569 | 0.027 | #NUM! | GPX1 |
| Replacement of protamines by nucleosomes in the male pronucleus | 0.569 | 0.027 | #NUM! | METTL23 |
| Nucleotide Excision Repair Pathway | 0.569 | 0.027 | #NUM! | POLR2L |
| Interferon Signaling | 0.569 | 0.027 | #NUM! | BAX |
| Cell surface interactions at the vascular wall | 0.566 | 0.014 | #NUM! | CD99,F2,HRAS |
| 3-phosphoinositide Biosynthesis | 0.562 | 0.014 | #NUM! | ACP1,DUSP5,SOCS3 |
| Cell Cycle Regulation by BTG Family Proteins | 0.559 | 0.0263 | #NUM! | BTG1 |
| Notch Signaling | 0.559 | 0.0263 | #NUM! | HES5 |
| Renin-Angiotensin Signaling | 0.559 | 0.0164 | #NUM! | HRAS,ITPR1 |
| Keratinization | 0.559 | 0.0139 | #NUM! | KRT14,PKP1,PKP3 |
| RHOA Signaling | 0.554 | 0.0163 | #NUM! | ACTA1,PFN1 |
| Complement System | 0.55 | 0.0256 | #NUM! | C7 |
| Synaptic Long Term Potentiation | 0.549 | 0.0161 | #NUM! | HRAS,ITPR1 |
| Nuclear Cytoskeleton Signaling Pathway | 0.541 | 0.0136 | #NUM! | ACTA1,TUBA1A,ZIC2 |
| Neurotransmitter release cycle | 0.54 | 0.025 | #NUM! | RAB3A |
| Gαs Signaling | 0.535 | 0.0157 | #NUM! | Htr5b,MC4R |
| Thrombin Signaling | 0.534 | 0.0135 | #NUM! | F2,HRAS,ITPR1 |
| DAG and IP3 signaling | 0.531 | 0.0244 | #NUM! | ITPR1 |
| Glioma Signaling | 0.53 | 0.0156 | #NUM! | CCND2,HRAS |
| GP6 Signaling Pathway | 0.53 | 0.0156 | #NUM! | ITPR1,VAV1 |

| | | | | |
|---|---|---|---|---|
| Glycation Signaling Pathway | 0.527 | 0.0133 | #NUM! | HBA2,HRAS,IL2 |
| Insulin Receptor Signaling | 0.526 | 0.0155 | #NUM! | HRAS,SOCS3 |
| IL-6 Signaling | 0.526 | 0.0155 | #NUM! | HRAS,SOCS3 |
| Pulmonary Fibrosis Idiopathic Signaling Pathway | 0.526 | 0.0123 | #NUM! | ACTA1,BAX,F2,HRAS |
| April Mediated Signaling | 0.523 | 0.0238 | #NUM! | TRAF5 |
| Cardiac conduction | 0.521 | 0.0154 | #NUM! | ITPR1,SLN |
| Neuroinflammation Signaling Pathway | 0.515 | 0.0121 | #NUM! | GABRA6,GABRD,KCNJ6,NTF3 |
| Role of Osteoblasts, Osteoclasts and Chondrocytes in Rheumatoid Arthritis | 0.514 | 0.0131 | #NUM! | BGLAP,TRAF5,WIF1 |
| DAP12 interactions | 0.514 | 0.0233 | #NUM! | HRAS |
| B Cell Activating Factor Signaling | 0.514 | 0.0233 | #NUM! | TRAF5 |
| Oncostatin M Signaling | 0.514 | 0.0233 | #NUM! | HRAS |
| Xenobiotic Metabolism Signaling | 0.512 | 0.0121 | #NUM! | FTL,Gstm6,Gstp1 (includes others),HRAS |
| fMLP Signaling in Neutrophils | 0.508 | 0.015 | #NUM! | HRAS,ITPR1 |
| PI3K Cascade | 0.506 | 0.0227 | #NUM! | FGF20 |
| Netrin-1 signaling | 0.506 | 0.0227 | #NUM! | NTN4 |
| Assembly and cell surface presentation of NMDA receptors | 0.506 | 0.0227 | #NUM! | TUBA1A |
| IL-27 Signaling Pathway | 0.504 | 0.0149 | #NUM! | IL2,SOCS3 |
| Signaling by Insulin receptor | 0.498 | 0.0222 | #NUM! | HRAS |
| Bile acid and bile salt metabolism | 0.498 | 0.0222 | #NUM! | AKR1D1 |
| Carboxyterminal post-translational modifications of tubulin | 0.498 | 0.0222 | #NUM! | TUBA1A |
| Complement cascade | 0.495 | 0.0147 | #NUM! | C7,F2 |
| CCR3 Signaling in Eosinophils | 0.495 | 0.0147 | #NUM! | HRAS,ITPR1 |

| | | | | |
|---|---|---|---|---|
| Sertoli Cell-Germ Cell Junction Signaling Pathway (Enhanced) | 0.492 | 0.0127 | #NUM! | ACTA1,CLDN18,HRAS |
| SUMOylation of DNA replication proteins | 0.49 | 0.0217 | #NUM! | TOP2A |
| M-decay: degradation of maternal mRNAs by maternally stored factors | 0.49 | 0.0217 | #NUM! | CNOT3 |
| IL-23 Signaling Pathway | 0.49 | 0.0217 | #NUM! | SOCS3 |
| SNARE Signaling Pathway | 0.487 | 0.0145 | #NUM! | BET1L,RAB3A |
| CGAS-STING Signaling Pathway | 0.487 | 0.0145 | #NUM! | BAX,IL2 |
| EIF2 Signaling | 0.486 | 0.0126 | #NUM! | ACTA1,HRAS,UBA52 |
| STAT3 Pathway | 0.483 | 0.0144 | #NUM! | HRAS,SOCS3 |
| Interleukin-2 family signaling | 0.482 | 0.0213 | #NUM! | IL2 |
| Role of Osteoblasts in Rheumatoid Arthritis Signaling Pathway | 0.48 | 0.0125 | #NUM! | BGLAP,IL2,WIF1 |
| RHO GTPases Activate Formins | 0.479 | 0.0143 | #NUM! | PFN1,TUBA1A |
| Superpathway of Inositol Phosphate Compounds | 0.477 | 0.0124 | #NUM! | ACP1,DUSP5,SOCS3 |
| Antimicrobial peptides | 0.468 | 0.0204 | #NUM! | BPIFB6 |
| iNOS Signaling | 0.468 | 0.0204 | #NUM! | HMGA1 |
| DHCR24 Signaling Pathway | 0.467 | 0.014 | #NUM! | HRAS,TF |
| S100 Family Signaling Pathway | 0.465 | 0.0102 | -0.378 | GPR150,ITPR1,MC4R,NTF3,Qrfpr,S100A16,S100A3,SSTR1 |
| Bone Mineralization Signaling Pathway | 0.463 | 0.0139 | #NUM! | FGF20,SMPD3 |
| Signaling by ROBO receptors | 0.462 | 0.0122 | #NUM! | PFN1,UBA52,UBB |
| Melanoma Signaling | 0.461 | 0.02 | #NUM! | HRAS |
| MYC Mediated Apoptosis Signaling | 0.461 | 0.02 | #NUM! | BAX |
| nNOS Signaling in Skeletal Muscle Cells | 0.461 | 0.02 | #NUM! | ITPR1 |

| | | | | |
|---|---|---|---|---|
| Gap junction trafficking and regulation | 0.454 | 0.0196 | #NUM! | TUBA1A |
| Cell Cycle: G2/M DNA Damage Checkpoint Regulation | 0.454 | 0.0196 | #NUM! | TOP2A |
| Wound Healing Signaling Pathway | 0.453 | 0.012 | #NUM! | F2,HRAS, IL2 |
| Docosahexaenoic Acid (DHA) Signaling | 0.448 | 0.012 | #NUM! | AKT1S1, BAX,ITPR 1 |
| B-WICH complex positively regulates rRNA expression | 0.447 | 0.0192 | #NUM! | POLR2L |
| Apoptotic execution phase | 0.447 | 0.0192 | #NUM! | PKP1 |
| PTEN Regulation | 0.44 | 0.0133 | #NUM! | UBA52,U BB |
| Endocannabinoid Neuronal Synapse Pathway | 0.44 | 0.0133 | #NUM! | ITPR1,KC NJ6 |
| G alpha (i) signalling events | 0.428 | 0.0116 | #NUM! | GAL,PCP 2,SSTR1 |
| Lymphotoxin β Receptor Signaling | 0.428 | 0.0182 | #NUM! | TRAF5 |
| Role of Cytokines in Mediating Communication between Immune Cells | 0.428 | 0.0182 | #NUM! | IL2 |
| Role of Pattern Recognition Receptors in Recognition of Bacteria and Viruses | 0.426 | 0.013 | #NUM! | IL2,NLRC 4 |
| IL-10 Signaling | 0.423 | 0.0129 | #NUM! | MAFB,SO CS3 |
| Corticotropin Releasing Hormone Signaling | 0.423 | 0.0129 | #NUM! | ITPR1,JU ND |
| DNA Damage/Telomere Stress Induced Senescence | 0.422 | 0.0179 | #NUM! | HMGA1 |
| CD27 Signaling in Lymphocytes | 0.422 | 0.0179 | #NUM! | TRAF5 |
| CNTF Signaling | 0.422 | 0.0179 | #NUM! | HRAS |
| HSP90 chaperone cycle for steroid hormone receptors in the presence of ligand | 0.416 | 0.0175 | #NUM! | TUBA1A |

| | | | | |
|---|---|---|---|---|
| Cohesin Chromatin Regulation Pathway | 0.415 | 0.0114 | #NUM! | FAT2,PCDHA12,POLR2L |
| Phagosome Formation | 0.41 | 0.00992 | 0 | GPR150,HRAS,ITPR1,MC4R,Qrfpr,SSTR1,VAV1 |
| Signaling by PDGF | 0.41 | 0.0172 | #NUM! | HRAS |
| Cancer Drug Resistance by Drug Efflux | 0.41 | 0.0172 | #NUM! | HRAS |
| HMGB1 Signaling | 0.406 | 0.0125 | #NUM! | HRAS,IL2 |
| Metabolism of polyamines | 0.404 | 0.0169 | #NUM! | OAZ1 |
| Endometrial Cancer Signaling | 0.398 | 0.0167 | #NUM! | HRAS |
| GADD45 Signaling | 0.398 | 0.0167 | #NUM! | CCND2 |
| Circadian Rhythm Signaling | 0.398 | 0.0111 | #NUM! | AKT1S1,HRAS,ITPR1 |
| Fcgamma receptor (FCGR) dependent phagocytosis | 0.393 | 0.0122 | #NUM! | ITPR1,VAV1 |
| Transcriptional Regulatory Network in Embryonic Stem Cells | 0.393 | 0.0122 | #NUM! | HRAS,PAX6 |
| Visual phototransduction | 0.393 | 0.0164 | #NUM! | NMT1 |
| Cell Cycle Checkpoints | 0.393 | 0.011 | #NUM! | MCM7,UBA52,UBB |
| Kinesins | 0.387 | 0.0161 | #NUM! | TUBA1A |
| MSP-RON Signaling Pathway | 0.387 | 0.0161 | #NUM! | ACTA1 |
| Coronavirus Replication Pathway | 0.387 | 0.012 | #NUM! | TUBA1A,UBA52 |
| NCAM signaling for neurite out-growth | 0.382 | 0.0159 | #NUM! | HRAS |
| Triacylglycerol Degradation | 0.382 | 0.0159 | #NUM! | AARSD1 |
| CXCR4 Signaling | 0.38 | 0.0119 | #NUM! | HRAS,ITPR1 |
| UFMylation Signaling Pathway | 0.377 | 0.0156 | #NUM! | GPX4 |
| Thrombopoietin Signaling | 0.377 | 0.0156 | #NUM! | HRAS |
| ERB2-ERBB3 Signaling | 0.377 | 0.0156 | #NUM! | HRAS |

| | | | | |
|---|---|---|---|---|
| Mitochondrial protein import | 0.372 | 0.0154 | #NUM! | TIMM10B |
| RAB geranylgeranylation | 0.372 | 0.0154 | #NUM! | RAB3A |
| SPINK1 Pancreatic Cancer Pathway | 0.372 | 0.0154 | #NUM! | CELA1 |
| Opioid Signaling Pathway | 0.371 | 0.0107 | #NUM! | HRAS,ITPR1,KCNJ6 |
| Induction of Apoptosis by HIV1 | 0.367 | 0.0152 | #NUM! | BAX |
| T Cell Receptor Signaling | 0.366 | 0.00965 | 1.633 | AKT1S1,CTLA4,DUSP5,HRAS,IL2,VAV1 |
| Protein Kinase A Signaling | 0.365 | 0.0101 | #NUM! | ACP1,DUSP5,ITPR1,PGP |
| Abacavir ADME | 0.362 | 0.0149 | #NUM! | NME7 |
| Ribavirin ADME | 0.362 | 0.0149 | #NUM! | NME7 |
| CD40 Signaling | 0.362 | 0.0149 | #NUM! | TRAF5 |
| ERBB4 Signaling | 0.362 | 0.0149 | #NUM! | HRAS |
| SPINK1 General Cancer Pathway | 0.362 | 0.0149 | #NUM! | HRAS |
| Sensory processing of sound by inner hair cells of the cochlea | 0.353 | 0.0145 | #NUM! | RAB3A |
| Cell Cycle: G1/S Checkpoint Regulation | 0.353 | 0.0145 | #NUM! | CCND2 |
| Molecular Mechanisms of Cancer | 0.353 | 0.00929 | -0.378 | BAX,CCND2,F2,GPR150,HRAS,MC4R,Qrfpr,SSTR1 |
| Dopamine-DARPP32 Feedback in cAMP Signaling | 0.351 | 0.0112 | #NUM! | ITPR1,KCNJ6 |
| TP53 Regulates Transcription of DNA Repair Genes | 0.348 | 0.0143 | #NUM! | POLR2L |
| GM-CSF Signaling | 0.348 | 0.0143 | #NUM! | HRAS |
| SUMOylation of chromatin organization proteins | 0.344 | 0.0141 | #NUM! | SATB2 |
| Glioma Invasiveness Signaling | 0.344 | 0.0141 | #NUM! | HRAS |
| Translocation of SLC2A4 (GLUT4) to the plasma membrane | 0.339 | 0.0139 | #NUM! | TUBA1A |

| | | | | |
|---|---|---|---|---|
| Growth Hormone Signaling | 0.335 | 0.0137 | #NUM! | SOCS3 |
| Olfactory Signaling Pathway | 0.331 | 0.0135 | #NUM! | ANO2 |
| RNA Polymerase I Transcription | 0.331 | 0.0135 | #NUM! | POLR2L |
| ERK5 Signaling | 0.331 | 0.0135 | #NUM! | HRAS |
| Major pathway of rRNA processing in the nucleolus and cytosol | 0.33 | 0.0108 | #NUM! | RPP25,UBA52 |
| Hepatic Fibrosis Signaling Pathway | 0.329 | 0.00962 | #NUM! | FTH1,FTL,HRAS,TF |
| Senescence Pathway | 0.329 | 0.00997 | #NUM! | CCND2,HRAS,MAPKAPK5 |
| Extra-nuclear estrogen signaling | 0.326 | 0.0133 | #NUM! | HRAS |
| Granzyme A Signaling | 0.326 | 0.0133 | #NUM! | NDUFA4L2 |
| Macrophage Classical Activation Signaling Pathway | 0.325 | 0.0106 | #NUM! | IL2,SOCS3 |
| Hepatitis B Chronic Liver Pathogenesis Signaling Pathway | 0.325 | 0.0106 | #NUM! | HRAS,SOCS3 |
| Macropinocytosis Signaling | 0.322 | 0.0132 | #NUM! | HRAS |
| Angiopoietin Signaling | 0.318 | 0.013 | #NUM! | HRAS |
| Antiproliferative Role of Somatostatin Receptor 2 | 0.318 | 0.013 | #NUM! | HRAS |
| Leptin Signaling in Obesity | 0.318 | 0.013 | #NUM! | SOCS3 |
| HIPPO signaling | 0.318 | 0.013 | #NUM! | TEAD2 |
| PKR-mediated signaling | 0.314 | 0.0128 | #NUM! | TUBA1A |
| NF-κB Activation by Viruses | 0.314 | 0.0128 | #NUM! | HRAS |
| Maturity Onset Diabetes of Young (MODY) Signaling | 0.314 | 0.0128 | #NUM! | NEUROD1 |
| GNRH Signaling | 0.312 | 0.0104 | #NUM! | HRAS,ITPR1 |
| Caveolar-mediated Endocytosis Signaling | 0.31 | 0.0127 | #NUM! | ACTA1 |
| TREM1 Signaling | 0.31 | 0.0127 | #NUM! | NLRC4 |
| IL-7 Signaling Pathway | 0.31 | 0.0127 | #NUM! | BAX |

| | | | | |
|---|---|---|---|---|
| Role of MAPK Signaling in Inhibiting the Pathogenesis of Influenza | 0.31 | 0.0127 | #NUM! | BAX |
| Endothelin-1 Signaling | 0.31 | 0.0103 | #NUM! | HRAS,ITPR1 |
| Regulation of the Epithelial-Mesenchymal Transition Pathway | 0.31 | 0.0103 | #NUM! | FGF20,HRAS |
| Regulation of the Epithelial Mesenchymal Transition by Growth Factors Pathway | 0.31 | 0.0103 | #NUM! | FGF20,HRAS |
| IL-33 Signaling Pathway | 0.308 | 0.0103 | #NUM! | CCL17,CELA1 |
| G alpha (12/13) signalling events | 0.307 | 0.0125 | #NUM! | VAV1 |
| Degradation of the extracellular matrix | 0.303 | 0.0123 | #NUM! | ADAMTS18 |
| Signaling by NTRK1 (TRKA) | 0.303 | 0.0123 | #NUM! | HRAS |
| IL-3 Signaling | 0.303 | 0.0123 | #NUM! | HRAS |
| Role of JAK family kinases in IL-6-type Cytokine Signaling | 0.303 | 0.0123 | #NUM! | SOCS3 |
| Chemokine Signaling | 0.303 | 0.0123 | #NUM! | HRAS |
| Adrenergic Receptor Signaling Pathway (Enhanced) | 0.301 | 0.0101 | #NUM! | IL2,ITPR1 |
| FLT3 Signaling in Hematopoietic Progenitor Cells | 0.299 | 0.0122 | #NUM! | HRAS |
| Role of Osteoclasts in Rheumatoid Arthritis Signaling Pathway | 0.298 | 0.00946 | #NUM! | ADAM2,HRAS,TRAF5 |
| Human Embryonic Stem Cell Pluripotency | 0.296 | 0.01 | #NUM! | HRAS,NTF3 |
| Estrogen-Dependent Breast Cancer Signaling | 0.295 | 0.012 | #NUM! | HRAS |
| Oxytocin in Brain Signaling Pathway | 0.294 | 0.00995 | #NUM! | HRAS,ITPR1 |
| Pulmonary Healing Signaling Pathway | 0.294 | 0.00995 | #NUM! | CELA1,HRAS |
| PEDF Signaling | 0.292 | 0.0119 | #NUM! | HRAS |
| Ephrin Receptor Signaling | 0.291 | 0.0099 | #NUM! | ACP1,HRAS |
| LPS-stimulated MAPK Signaling | 0.288 | 0.0118 | #NUM! | HRAS |

| | | | | |
|---|---|---|---|---|
| Intra-Golgi and retrograde Golgi-to-ER traffic | 0.287 | 0.0098 | #NUM! | BET1L,TUBA1A |
| Platelet homeostasis | 0.285 | 0.0116 | #NUM! | ITPR1 |
| Cyclins and Cell Cycle Regulation | 0.285 | 0.0116 | #NUM! | CCND2 |
| VEGF Family Ligand-Receptor Interactions | 0.285 | 0.0116 | #NUM! | HRAS |
| Mitotic Prometaphase | 0.285 | 0.00976 | #NUM! | NME7,TUBA1A |
| Role of Tissue Factor in Cancer | 0.278 | 0.00962 | #NUM! | F2,HRAS |
| Telomere Maintenance | 0.275 | 0.0112 | #NUM! | POLR2L |
| HIF1α Signaling | 0.274 | 0.00952 | #NUM! | HRAS,TF |
| Opioid Signalling | 0.272 | 0.0111 | #NUM! | ITPR1 |
| RAB GEFs exchange GTP for GDP on RABs | 0.272 | 0.0111 | #NUM! | RAB3A |
| Tuberculosis Latent Signaling Pathway | 0.272 | 0.0111 | #NUM! | TRAF5 |
| Activin Inhibin Signaling Pathway | 0.27 | 0.00943 | #NUM! | CCND2,PAX6 |
| Actin Nucleation by ARP-WASP Complex | 0.268 | 0.011 | #NUM! | HRAS |
| mTOR Signaling | 0.266 | 0.00935 | #NUM! | AKT1S1, HRAS |
| Acute Myeloid Leukemia Signaling | 0.265 | 0.0109 | #NUM! | HRAS |
| EPH-Ephrin signaling | 0.262 | 0.0108 | #NUM! | HRAS |
| RANK Signaling in Osteoclasts | 0.262 | 0.0108 | #NUM! | TRAF5 |
| ERBB Signaling | 0.262 | 0.0108 | #NUM! | HRAS |
| Calcium Signaling | 0.258 | 0.00917 | #NUM! | ACTA1,ITPR1 |
| Immunogenic Cell Death Signaling Pathway | 0.256 | 0.0105 | #NUM! | BAX |
| Class B/2 (Secretin family receptors) | 0.256 | 0.0105 | #NUM! | GCG |
| Autophagy | 0.256 | 0.00913 | #NUM! | AKT1S1, GCG |
| ERK/MAPK Signaling | 0.254 | 0.00909 | #NUM! | HRAS,MAPKAPK5 |
| Xenobiotic Metabolism PXR Signaling Pathway | 0.252 | 0.00905 | #NUM! | Gstm6,Gstp1 (includes others) |
| Mitochondrial translation | 0.25 | 0.0103 | #NUM! | MRPL53 |

| | | | | |
|---|---|---|---|---|
| Death Receptor Signaling | 0.25 | 0.0103 | #NUM! | ACTA1 |
| TGF-β Signaling | 0.25 | 0.0103 | #NUM! | HRAS |
| Interferon gamma signaling | 0.247 | 0.0102 | #NUM! | SOCS3 |
| DNA Methylation and Transcriptional Repression Signaling | 0.247 | 0.0102 | #NUM! | SAP18 |
| Hepatic Cholestasis | 0.245 | 0.00889 | #NUM! | GCG,IL2 |
| Protein folding | 0.245 | 0.0101 | #NUM! | TUBA1A |
| Melanocyte Development and Pigmentation Signaling | 0.245 | 0.0101 | #NUM! | HRAS |
| Salvage Pathways of Pyrimidine Ribonucleotides | 0.242 | 0.01 | #NUM! | NME7 |
| Apelin Cardiomyocyte Signaling Pathway | 0.242 | 0.01 | #NUM! | ITPR1 |
| HER-2 Signaling in Breast Cancer | 0.241 | 0.00881 | #NUM! | AKT1S1, HRAS |
| Neuropathic Pain Signaling in Dorsal Horn Neurons | 0.239 | 0.0099 | #NUM! | ITPR1 |
| Sumoylation Pathway | 0.239 | 0.0099 | #NUM! | RCC1 |
| Role of NFAT in Cardiac Hypertrophy | 0.237 | 0.00873 | #NUM! | HRAS,ITPR1 |
| mRNA 3 Prime End Processing Signaling Pathway | 0.236 | 0.0098 | #NUM! | CNOT3 |
| Response of EIF2AK4 (GCN2) to amino acid deficiency | 0.234 | 0.00971 | #NUM! | UBA52 |
| GPER1 signaling | 0.234 | 0.00971 | #NUM! | NME7 |
| Mouse Embryonic Stem Cell Pluripotency | 0.234 | 0.00971 | #NUM! | HRAS |
| Xenobiotic Metabolism CAR Signaling Pathway | 0.23 | 0.00858 | #NUM! | Gstm6,Gstp1 (includes others) |
| Sleep REM Signaling Pathway | 0.226 | 0.00943 | #NUM! | IL2 |
| CDK5 Signaling | 0.226 | 0.00943 | #NUM! | HRAS |
| Extracellular matrix organization | 0.223 | 0.00935 | #NUM! | NTN4 |
| PPAR Signaling | 0.221 | 0.00926 | #NUM! | HRAS |
| Sphingolipid metabolism | 0.216 | 0.00909 | #NUM! | SMPD3 |

| | | | | |
|---|---|---|---|---|
| PD-1, PD-L1 cancer immunotherapy pathway | 0.207 | 0.00877 | #NUM! | IL2 |
| Prostate Cancer Signaling | 0.202 | 0.00862 | #NUM! | HRAS |
| Nonsense-Mediated Decay (NMD) | 0.2 | 0.00855 | #NUM! | UBA52 |
| Cerebral Malformation Signaling Pathway | 0.2 | 0.00855 | #NUM! | F2 |
| Neuregulin Signaling | 0.198 | 0.00847 | #NUM! | HRAS |
| PAK Signaling | 0.198 | 0.00847 | #NUM! | HRAS |
| NAD Signaling Pathway | 0 | 0.00662 | #NUM! | POLR2L |
| Oxytocin Signaling Pathway | 0 | 0.00702 | #NUM! | HRAS,ITPR1 |
| IL-13 Signaling Pathway | 0 | 0.00813 | #NUM! | SOCS3 |
| CDX Gastrointestinal Cancer Signaling Pathway | 0 | 0.00508 | #NUM! | IL2 |
| Role of Chondrocytes in Rheumatoid Arthritis Signaling Pathway | 0 | 0.00694 | #NUM! | AKT1S1 |
| Ribonucleotide Reductase Signaling Pathway | 0 | 0.00575 | #NUM! | AKT1S1 |
| Neutrophil Extracellular Trap Signaling Pathway | 0 | 0.00489 | #NUM! | ITPR1,NDUFA4L2 |
| Chaperone Mediated Autophagy Signaling Pathway | 0 | 0.00157 | #NUM! | GPX4 |
| NOD1/2 Signaling Pathway | 0 | 0.00518 | #NUM! | IL2 |
| Acetylcholine Receptor Signaling Pathway | 0 | 0.00505 | #NUM! | ITPR1 |
| Cachexia Signaling Pathway | 0 | 0.00815 | #NUM! | IL2,MC4R,SOCS3 |
| Microautophagy Signaling Pathway | 0 | 0.00629 | #NUM! | VPS25 |
| PPARα/RXRα Activation | 0 | 0.00505 | #NUM! | HRAS |
| LPS/IL-1 Mediated Inhibition of RXR Function | 0 | 0.00671 | #NUM! | Gstm6,Gstp1 (includes others) |
| Eukaryotic Translation Elongation | 0 | 0.0082 | #NUM! | UBA52 |

| | | | | |
|---|---|---|---|---|
| Glycosaminoglycan metabolism | 0 | 0.00781 | #NUM! | GLCE |
| SRP-dependent cotranslational protein targeting to membrane | 0 | 0.00704 | #NUM! | UBA52 |
| Immunoregulatory interactions between a Lymphoid and a non-Lymphoid cell | 0 | 0.00465 | #NUM! | CD99 |
| MHC class II antigen presentation | 0 | 0.00781 | #NUM! | TUBA1A |
| Selenoamino acid metabolism | 0 | 0.00685 | #NUM! | UBA52 |
| L1CAM interactions | 0 | 0.00806 | #NUM! | TUBA1A |
| Cilium Assembly | 0 | 0.0049 | #NUM! | TUBA1A |
| Deubiquitination | 0 | 0.00755 | #NUM! | UBA52,UBB |
| Eukaryotic Translation Initiation | 0 | 0.00667 | #NUM! | UBA52 |
| Eukaryotic Translation Termination | 0 | 0.0082 | #NUM! | UBA52 |
| RHO GTPase cycle | 0 | 0.00444 | #NUM! | DDX4,VAV1 |
| LXR/RXR Activation | 0 | 0.00769 | #NUM! | TF |
| Hepatic Fibrosis / Hepatic Stellate Cell Activation | 0 | 0.00521 | #NUM! | BAX |
| ROBO SLIT Signaling Pathway | 0 | 0.00781 | #NUM! | PAX6 |
| BBSome Signaling Pathway | 0 | 0.00811 | #NUM! | GPR150, MC4R,Qrfpr,SSTR1 |
| HEY1 Signaling Pathway | 0 | 0.00621 | #NUM! | BAX |
| WNT/SHH Axonal Guidance Signaling Pathway | 0 | 0.00662 | #NUM! | ITPR1 |
| Histone Modification Signaling Pathway | 0 | 0.00667 | #NUM! | PRDM8,ZIC2 |
| Ribosomal Quality Control Signaling Pathway | 0 | 0.00379 | #NUM! | UBA52 |
| Chromatin organization | 0 | 0.00784 | #NUM! | SAP18,TAF9 |
| Cyclophilin Signaling Pathway | 0 | 0.00803 | #NUM! | F2,IL2 |
| Irritable Bowel Syndrome Signaling Pathway | 0 | 0.00346 | #NUM! | IL2 |

| | | | | |
|---|---|---|---|---|
| Lung Ionic Balance Signaling Pathway | 0 | 0.00766 | #NUM! | GPR150, MC4R,Qrfpr,SSTR1 |
| TRIM21 Intracellular Antibody Signaling Pathway | 0 | 0.00353 | #NUM! | UBA52,UBB |
| ID3 Signaling Pathway | 0 | 0.00103 | #NUM! | HRAS |
| Preeclampsia Signaling Pathway | 0 | 0.00621 | #NUM! | AKT1S1 |
| Tuberculosis Active Signaling Pathway | 0 | 0.00408 | #NUM! | BAX |
| TR/RXR Activation | 0 | 0.00763 | #NUM! | ENO1 |
| RAR Activation | 0 | 0.00693 | #NUM! | IL2,PAX6, SOCS3 |
| IL-12 Signaling and Production in Macrophages | 0 | 0.00826 | #NUM! | SOCS3,ZIC2 |
| Role of PKR in Interferon Induction and Antiviral Response | 0 | 0.00725 | #NUM! | BAX |
| Role of NFAT in Regulation of the Immune Response | 0 | 0.00191 | #NUM! | HRAS,ITPR1 |
| FcγRIIB Signaling in B Lymphocytes | 0 | 0.00375 | #NUM! | HRAS,ITPR1 |
| CCR5 Signaling in Macrophages | 0 | 0.002 | #NUM! | ITPR1 |
| Calcium-induced T Lymphocyte Apoptosis | 0 | 0.00215 | #NUM! | ITPR1 |
| CTLA4 Signaling in Cytotoxic T Lymphocytes | 0 | 0.00656 | -1 | CTLA4,HRAS,IL2, VAV1 |
| T Helper Cell Differentiation | 0 | 0.00211 | #NUM! | IL2 |
| CD28 Signaling in T Helper Cells | 0 | 0.00766 | 0 | CTLA4,IL2,ITPR1, VAV1 |
| IL-15 Signaling | 0 | 0.0038 | #NUM! | AKT1S1, HRAS |
| Reelin Signaling in Neurons | 0 | 0.00725 | #NUM! | RELN |
| Cellular Effects of Sildenafil (Viagra) | 0 | 0.0073 | 1 | ACTA1,GPR150,MC4R,Qrfpr,SSTR1 |
| Cardiac Hypertrophy Signaling | 0 | 0.00383 | #NUM! | HRAS |

| | | | | |
|---|---|---|---|---|
| ICOS-ICOSL Signaling in T Helper Cells | 0 | 0.00586 | #NUM! | IL2,ITPR1,VAV1 |
| HGF Signaling | 0 | 0.00763 | #NUM! | HRAS |
| Aldosterone Signaling in Epithelial Cells | 0 | 0.00585 | #NUM! | ITPR1 |
| Role of NANOG in Mammalian Embryonic Stem Cell Pluripotency | 0 | 0.00813 | #NUM! | HRAS |
| Type I Diabetes Mellitus Signaling | 0 | 0.00391 | #NUM! | IL2,SOCS3 |
| Allograft Rejection Signaling | 0 | 0.00203 | #NUM! | IL2 |
| Autoimmune Thyroid Disease Signaling | 0 | 0.00216 | #NUM! | IL2 |
| Graft-versus-Host Disease Signaling | 0 | 0.00223 | #NUM! | IL2 |
| p70S6K Signaling | 0 | 0.00344 | #NUM! | F2,HRAS |
| Colorectal Cancer Metastasis Signaling | 0 | 0.00738 | #NUM! | BAX,HRAS |
| G Protein Signaling Mediated by Tubby | 0 | 0.00213 | #NUM! | VAV1 |
| Communication between Innate and Adaptive Immune Cells | 0 | 0.00106 | #NUM! | IL2 |
| Sphingosine-1-phosphate Signaling | 0 | 0.0084 | #NUM! | SMPD3 |
| Systemic Lupus Erythematosus Signaling | 0 | 0.00562 | #NUM! | C7,HRAS,IL2,LSM10,LSM2,LSM4 |
| CDC42 Signaling | 0 | 0.00173 | #NUM! | VAV1 |
| ILK Signaling | 0 | 0.00508 | #NUM! | ACTA1 |
| FAK Signaling | 0 | 0.00676 | 0.816 | GPR150,HRAS,MAPKAPK5,MC4R,Qrfpr,SOCS3,SSTR1 |
| AMPK Signaling | 0 | 0.0082 | #NUM! | AKT1S1,RAB3A |
| RAC Signaling | 0 | 0.0073 | #NUM! | HRAS |
| Phospholipase C Signaling | 0 | 0.00178 | #NUM! | HRAS,ITPR1 |
| Ovarian Cancer Signaling | 0 | 0.00629 | #NUM! | HRAS |
| Altered T Cell and B Cell Signaling in Rheumatoid Arthritis | 0 | 0.00106 | #NUM! | IL2 |

| | | | | |
|---|---|---|---|---|
| Regulation of eIF4 and p70S6K Signaling | 0 | 0.00532 | #NUM! | HRAS |
| Neuroprotective Role of THOP1 in Alzheimer's Disease | 0 | 0.00769 | #NUM! | RELN |
| Regulation of IL-2 Expression in Activated and Anergic T Lymphocytes | 0 | 0.00642 | #NUM! | HRAS,IL2,VAV1 |
| PKCθ Signaling in T Lymphocytes | 0 | 0.00714 | #NUM! | HRAS,IL2,ITPR1,VAV1 |
| Antiproliferative Role of TOB in T Cell Signaling | 0 | 0.00235 | #NUM! | IL2 |
| OX40 Signaling Pathway | 0 | 0.00415 | #NUM! | IL2,TRAF5 |
| PI3K Signaling in B Lymphocytes | 0 | 0.00507 | #NUM! | HRAS,ITPR1,VAV1 |
| P2Y Purinergic Receptor Signaling Pathway | 0 | 0.00746 | #NUM! | HRAS |
| Signaling by Rho Family GTPases | 0 | 0.00375 | #NUM! | ACTA1 |
| RHOGDI Signaling | 0 | 0.00455 | #NUM! | ACTA1 |
| Hematopoiesis from Pluripotent Stem Cells | 0 | 0.00223 | #NUM! | IL2 |
| eNOS Signaling | 0 | 0.00621 | #NUM! | ITPR1 |
| Epithelial Adherens Junction Signaling | 0 | 0.00633 | #NUM! | HRAS |
| Gαi Signaling | 0 | 0.0068 | #NUM! | HRAS |
| Gαq Signaling | 0 | 0.00592 | #NUM! | ITPR1 |
| Sperm Motility | 0 | 0.00389 | #NUM! | ITPR1 |
| TEC Kinase Signaling | 0 | 0.00346 | #NUM! | ACTA1,VAV1 |
| Adipogenesis pathway | 0 | 0.00725 | #NUM! | SAP18 |
| Estrogen Receptor Signaling | 0 | 0.00487 | #NUM! | HRAS,NDUFA4L2 |
| SAPK/JNK Signaling | 0 | 0.00199 | #NUM! | HRAS |
| PI3K/AKT Signaling | 0 | 0.005 | #NUM! | HRAS |
| PTEN Signaling | 0 | 0.00662 | #NUM! | HRAS |
| Nitric Oxide Signaling in the Cardiovascular System | 0 | 0.00833 | #NUM! | ITPR1 |
| IL-4 Signaling | 0 | 0.00348 | #NUM! | AKT1S1, HRAS |
| B Cell Receptor Signaling | 0 | 0.00313 | #NUM! | HRAS,VAV1 |
| cAMP-mediated signaling | 0 | 0.0083 | #NUM! | Htr5b,MC4R |

| | | | | |
|---|---|---|---|---|
| p38 MAPK Signaling | 0 | 0.00806 | #NUM! | MAPKAPK5 |
| NF-κB Signaling | 0 | 0.00525 | #NUM! | HRAS,TRAF5,UBE2V1 |
| G-Protein Coupled Receptor Signaling | 0 | 0.00842 | 0.447 | GPR150,HRAS,Htr5b,MC4R,Qrfpr,SSTR1 |
| Osteoarthritis Pathway | 0 | 0.0042 | #NUM! | BGLAP |
| Th17 Activation Pathway | 0 | 0.00204 | #NUM! | SOCS3 |
| Apelin Endothelial Signaling Pathway | 0 | 0.0069 | #NUM! | HRAS |
| Cardiac Hypertrophy Signaling (Enhanced) | 0 | 0.00745 | -1 | FGF20,HRAS,IL2,ITPR1 |
| T Cell Exhaustion Signaling Pathway | 0 | 0.00353 | #NUM! | CTLA4,HRAS |
| Systemic Lupus Erythematosus in T Cell Signaling Pathway | 0 | 0.00775 | 0.447 | AKT1S1,HRAS,IL2,ITPR1,VAV1 |
| Systemic Lupus Erythematosus in B Cell Signaling Pathway | 0 | 0.00687 | -0.447 | CCND2,HRAS,IL2,TRAF5,VAV1 |
| White Adipose Tissue Browning Pathway | 0 | 0.00719 | #NUM! | ITPR1 |
| Inhibition of ARE-Mediated mRNA Degradation Pathway | 0 | 0.00617 | #NUM! | CNOT3 |
| HOTAIR Regulatory Pathway | 0 | 0.00606 | #NUM! | WIF1 |
| Coronavirus Pathogenesis Pathway | 0 | 0.00474 | #NUM! | BAX |
| MSP-RON Signaling in Cancer Cells Pathway | 0 | 0.00699 | #NUM! | HRAS |
| Eicosanoid Signaling | 0 | 0.00709 | #NUM! | HRAS,MAPKAPK5 |
