## Supplemental Table 3 for "Early Life Adversity Produces Enduring Molecular and Functional Disruption of Developing Vagal Circuits"

| Category | Term | FDR | PValue | Genes | Fold |
|---|---|---|---|---|---|
| UP_KW_PTM | KW-0027~Amidation | 1.55525E-05 | 8.18553E-07 | 21334, 14419, 54615, 22044, 22353, 20604, | 21.15748 |
| UP_KW_PTM | KW-0165~Cleavage on pair of basic residues | 0.001401382 | 0.000147514 | 21334, 14061, 14419, 14560, 54615, 22044, | 5.05665 |
| GOTERM_CC_DIRECT | GO:0005576~extracellular region | 0.001626818 | 1.03988E-05 | 20202, 14061, 21334, 269855, 14560, 22264, 22044, 22353, 18591, 58991, 20604, 76293, 16170, 73690, 14419, | 2.855875 |
| GOTERM_CC_DIRECT | GO:0005615~extracellular space | 0.001626818 | 1.37284E-05 | 13166, 20295, 18591, 58991, 319433, 16170, 14419, 56410, 54615, 54612, 12159, 100503605, 11568, 69511, 20202, 21334, | 2.525142 |
| UP_KW_MOLECULAR_FUNCTION | KW-9996~Developmental protein | 0.005249052 | 0.000238593 | 15285, 104382, 65100, 23937, 74469, 269855, 18012, 18591, 15425, 60364, 15414, 14369, | 2.594183 |
| UP_KW_MOLECULAR_FUNCTION | KW-0217~Developmental protein | 0.005249052 | 0.000238593 | 15285, 104382, 65100, 23937, 74469, 269855, 18012, 18591, 15425, 60364, 15414, 14369, | 2.594183 |
| KEGG_PATHWAY | mmu04080:Neuroactive ligand-receptor interaction | 0.006536389 | 3.63133E-05 | 14061, 21334, 22044, 22353, 58991, 20604, 14419, 13609, 54615, | 4.293421 |
| UP_KW_CELLULAR_COMPONENT | KW-0964~Secreted | 0.010629281 | 0.000561774 | 20295, 22353, 18591, 58991, 16170, 14419, 56410, 54615, 13024, 54612, 12159, 11568, 20202, 21334, 20201, | 2.10421 |
| UP_KW_CELLULAR_COMPONENT | KW-0539~Nucleus | 0.010629281 | 0.000708619 | 104382, 68992, 18012, 15425, 66373, 12236, 66491, 23937, 22289, 54324, 12608, 60364, 18508, 13498, 18227, 666528, 18988, 68240, 15285, 15361, 65100, 14870, 15364, 21681, 107071, 98267, 20957, 109095, 16170, 22138, 60406, 80981, 78784, | 1.498985 |

| | | | | | |
|---|---|---|---|---|---|
| UP_KW_DOMAIN | KW-0371~Homeobox | 0.014950429 | 0.000747521 | 15285, 104382, 15425, 15414, 15402, 18508, | 5.297735 |
| GOTERM_MF_DIRECT | GO:1990837~sequence-specific double-stranded DNA binding | 0.021336804 | 6.23883E-05 | 15285, 14281, 104382, 68992, 65100, 18012, 12608, 15414, 15402, | 4.228882 |
| GOTERM_CC_DIRECT | GO:0005634~nucleus | 0.024294514 | 0.000307525 | 104382, 68992, 18012, 15425, 66373, 12236, 66491, 23937, 22289, 54324, 12608, 60364, 18508, 13498, 18227, 666528, 18988, 15285, 15361, 65100, 14870, 15364, 21681, 107071, 98267, 109054, 109095, 16170, 22138, 60406, 80981, 78784, 434484, | 1.595639 |
| GOTERM_CC_DIRECT | GO:0005737~cytoplasm | 0.028036582 | 0.000473191 | 107971, 242785, 18012, 14693, 58991, 57267, 13609, 69684, 66373, 12236, 23937, 269855, 232174, 54324, 12608, 18508, 11431, 13498, 18227, 58804, 15361, 14870, 21681, 109054, 18591, 21648, 16170, 270150, 22138, 80981, 78784, 434484, 66395, 11568, 623503, 17750, 21973, 20202, 14281, | 1.548289 |
| UP_KW_BIOLOGICAL_PROCESS | KW-0145~Chemotaxis | 0.033915704 | 0.000584754 | 20202, 20201, 68992, 20295, 13609, 16170 | 8.704969 |
| GOTERM_MF_DIRECT | GO:0000979~RNA polymerase II core promoter sequence- | 0.057041082 | 0.000400366 | 15361, 14281, 12608, 18508 | 27.47522 |
| GOTERM_MF_DIRECT | GO:0005184~neuropeptide hormone activity | 0.057041082 | 0.00050036 | 14419, 54615, 22353, 623503 | 25.51271 |
| GOTERM_BP_DIRECT | GO:0009952~anterior/posterior pattern | 0.060330528 | 8.40007E-05 | 68713, 18012, 15425, 15414, 15402, 18508, | 9.768701 |
| GOTERM_BP_DIRECT | GO:0030182~neuron differentiation | 0.060330528 | 9.63746E-05 | 15285, 14281, 104382, 12608, 13609, 18508, | 6.304701 |
| GOTERM_CC_DIRECT | GO:0001673~male germ cell nucleus | 0.06871784 | 0.001449743 | 15361, 21973, 74469, 15364, 20957 | 10.24302 |

| | | | | | |
|---|---|---|---|---|---|
| INTERPRO | IPR017970:Homeobox_CS | 0.0767414 | 0.000177341 | 15285, 104382, 15425, 15414, 15402, 18508, | 6.802858 |
| INTERPRO | IPR020479:HD_metazoa | 0.0767414 | 0.000295159 | 15285, 104382, 15425, 15414, 15402, 13799 | 10.26036 |
| UP_KW_MOLECULAR_FUNCTION | KW-0238~DNA-binding | 0.086737723 | 0.005913936 | 15285, 15361, 21973, 14281, 104382, 68992, 65100, 75316, 15331, 15364, 18012, 20957, 60406, 12608, 15425, | 1.824667 |
| UP_KW_MOLECULAR_FUNCTION | KW-0372~Hormone | 0.088953223 | 0.008086657 | 14419, 22044, 22353, 20604, 58991 | 6.24592 |
| UP_KW_MOLECULAR_FUNCTION | KW-0527~Neuropeptide | 0.090500479 | 0.011899886 | 21334, 14419, 54615 | 17.80087 |
| UP_KW_MOLECULAR_FUNCTION | KW-0339~Growth factor | 0.090500479 | 0.012340974 | 14560, 12587, 18591, 16183, 12159 | 5.51965 |
| GOTERM_MF_DIRECT | GO:0003680~minor groove of adenine- | 0.115004349 | 0.00134508 | 15361, 15364, 18508 | 53.57669 |
| INTERPRO | IPR050296:Antp_home | 0.144122804 | 0.000831478 | 15425, 15414, 15402 | 66.69231 |
| UP_KW_BIOLOGICAL_PROCESS | KW-0804~Transcription | 0.148436348 | 0.005334145 | 15285, 15361, 104382, 68992, 15364, 18012, 109095, 16170, 60406, 15425, 15402, 11568, 66491, 74469, 75316, 240514, 12608, 12388, | 1.705347 |
| UP_KW_BIOLOGICAL_PROCESS | KW-0805~Transcription regulation | 0.148436348 | 0.007677742 | 15285, 15361, 104382, 68992, 15364, 18012, 109095, 16170, 60406, 15425, 15402, 11568, 74469, 75316, 240514, | 1.685305 |
| INTERPRO | IPR009057:Homeodomain-like_sf | 0.173066017 | 0.001475834 | 15285, 104382, 15425, 15414, 15402, 18508, | 4.1559 |
| INTERPRO | IPR001356:HD | 0.173066017 | 0.001664096 | 15285, 104382, 15425, 15414, 15402, 18508, | 4.662633 |
| GOTERM_BP_DIRECT | GO:0006366~transcription by RNA polymerase | 0.184018758 | 0.00044094 | 14281, 18012, 12608, 18508, 18227, 16183, | 5.883225 |
| GOTERM_MF_DIRECT | GO:0003677~DNA binding | 0.253169165 | 0.003753895 | 15285, 15361, 21973, 14281, 104382, 68992, 65100, 75316, 18012, 60406, 12608, 15425, | 2.172028 |
| GOTERM_MF_DIRECT | GO:0008301~DNA binding, bending | 0.253169165 | 0.004441564 | 15361, 21973, 15364 | 29.76483 |
| INTERPRO | IPR017995:Homeobox_antennapedia | 0.303247295 | 0.003499007 | 15425, 15414, 15402 | 33.34615 |

| | | | | | |
|---|---|---|---|---|---|
| GOTERM_MF_DIRECT | GO:0000978~RNA polymerase II cis-regulatory region | 0.306322008 | 0.006269749 | 15361, 14281, 68992, 65100, 18012, 22289, 12608, 15425, 434484, | 2.274053 |
| UP_KW_MOLECULAR_FUNCTION | KW-0202~Cytokine | 0.316126789 | 0.050292898 | 14560, 20295, 16183, 12159, 16170 | 3.560174 |
| GOTERM_BP_DIRECT | GO:2000379~positive regulation of reactive oxygen species | 0.36634242 | 0.001170423 | 14061, 13078, 13197, 18591 | 19.16023 |
| GOTERM_BP_DIRECT | GO:0043065~positive regulation of apoptotic | 0.373268934 | 0.001933756 | 21973, 14419, 13078, 18012, 13197, 15402, | 3.997718 |
| GOTERM_BP_DIRECT | GO:0010628~positive regulation of gene expression | 0.373268934 | 0.002180825 | 68992, 232174, 15364, 12608, 15402, 18508, 18591, 22289, 12159, | 3.24385 |
| GOTERM_BP_DIRECT | GO:0048333~mesodermal cell differentiation | 0.373268934 | 0.002341997 | 15364, 22289, 12159 | 40.89973 |
| GOTERM_BP_DIRECT | GO:0045944~positive regulation of transcription by RNA polymerase II | 0.373268934 | 0.002385105 | 15361, 21973, 14281, 104382, 15364, 18012, 22289, 16183, 14419, 12608, 13609, 15402, | 2.350192 |
| GOTERM_MF_DIRECT | GO:0001664~G protein-coupled receptor | 0.379600014 | 0.009822672 | 110257, 13609, 54615, 58991 | 9.042479 |
| GOTERM_MF_DIR | GO:0008083~growth | 0.379600014 | 0.010223369 | 14560, 12587, 18591, | 5.913542 |
| GOTERM_MF_DIRECT | GO:0097493~structural molecule activity | 0.379600014 | 0.011099416 | 66395, 22138 | 178.589 |
| GOTERM_BP_DIRECT | GO:0045444~fat cell differentiation | 0.381509449 | 0.00274248 | 14560, 15364, 12608, 18227, 623503 | 8.603502 |
| GOTERM_BP_DIRECT | GO:0001709~cell fate determination | 0.385398321 | 0.004023797 | 104382, 232174, 18508 | 31.27626 |
| GOTERM_BP_DIRECT | GO:0008380~RNA splicing | 0.385398321 | 0.004062878 | 78784, 69878, 21681, 56009, 66373, 109095 | 5.717166 |
| GOTERM_BP_DIRECT | GO:0007417~central nervous system | 0.385398321 | 0.004071964 | 15285, 65100, 232174, 18012, 12667 | 7.705745 |
| GOTERM_BP_DIRECT | GO:1902895~positive regulation of miRNA | 0.385398321 | 0.004265114 | 14281, 18508, 18591, 12159 | 12.22291 |
| GOTERM_BP_DIRECT | GO:0045893~positive regulation of DNA-templated transcription | 0.385398321 | 0.00482682 | 15361, 14281, 15364, 18012, 12608, 14369, 18508, 18591, 13498, | 2.716201 |
| GOTERM_BP_DIRECT | GO:0000122~negative regulation of transcription by RNA | 0.385398321 | 0.004886209 | 15285, 15361, 15331, 15364, 13197, 60406, 22097, 12608, 18508, | 2.444581 |
| GOTERM_BP_DIRECT | GO:0009611~response to wounding | 0.385398321 | 0.005467711 | 15361, 14061, 381232, 18508, 18591 | 7.089286 |
| GOTERM_BP_DIRECT | GO:0030595~leukocyte chemotaxis | 0.385398321 | 0.00555984 | 20202, 13609, 16170 | 26.58482 |

| | | | | | |
|---|---|---|---|---|---|
| GOTERM_BP_DIRECT | GO:0048706~embryonic skeletal system | 0.385398321 | 0.005619848 | 15425, 15414, 15402, 12159 | 11.07701 |
| GOTERM_BP_DIRECT | GO:0021549~cerebellum development | 0.385398321 | 0.005867819 | 319352, 14560, 18012, 18508 | 10.90659 |
| GOTERM_BP_DIRECT | GO:0010467~gene expression | 0.385398321 | 0.007108625 | 15361, 319352, 18508, 18591, 18227, 16183, | 3.589512 |
| GOTERM_BP_DIR | GO:0045471~response | 0.385398321 | 0.007330498 | 14281, 14870, 22044, | 6.515888 |
| GOTERM_BP_DIRECT | GO:0008285~negative regulation of cell | 0.385398321 | 0.007432805 | 15361, 14419, 68713, 13078, 27206, 54612, | 3.562455 |
| GOTERM_BP_DIRECT | GO:0055085~transmembrane transport | 0.385398321 | 0.007461637 | 108115, 68267, 55961, 244562, 20538, 67473 | 4.946013 |
| GOTERM_BP_DIRECT | GO:0030902~hindbrain development | 0.385398321 | 0.007958719 | 18012, 18508, 13799 | 22.15402 |
| GOTERM_BP_DIRECT | GO:0003016~respiratory system process | 0.385398321 | 0.007958719 | 15361, 15402, 22289 | 22.15402 |
| GOTERM_BP_DIRECT | GO:0008283~cell population proliferation | 0.385398321 | 0.008143517 | 15361, 23937, 15364, 13609, 18508, 12667, | 4.014968 |
| GOTERM_BP_DIRECT | GO:0007399~nervous system development | 0.385398321 | 0.008394032 | 14281, 104382, 14419, 14588, 18508, 21648, | 3.989148 |
| GOTERM_BP_DIRECT | GO:0060441~epithelial tube branching involved in lung morphogenesis | 0.385398321 | 0.008619132 | 15364, 15402, 12159 | 21.26786 |
| GOTERM_BP_DIRECT | GO:0051412~response to corticosterone | 0.392632914 | 0.010010778 | 14281, 15496, 22044 | 19.69246 |
| GOTERM_BP_DIR | GO:0009749~response | 0.392632914 | 0.010035346 | 15361, 18012, 22044, | 8.973779 |
| GOTERM_BP_DIR | GO:0008542~visual | 0.392632914 | 0.010035346 | 319352, 13166, 58991, | 8.973779 |
| GOTERM_BP_DIRECT | GO:0045669~positive regulation of osteoblast | 0.392632914 | 0.010035346 | 68713, 14560, 12608, 12159 | 8.973779 |
| GOTERM_BP_DIRECT | GO:0001938~positive regulation of | 0.393966901 | 0.010384112 | 68992, 22353, 18591, 12159 | 8.861607 |
| SMART | SM00389:HOX | 0.407069779 | 0.004473294 | 15285, 104382, 15425, 15414, 15402, 18508, | 3.841011 |
| GOTERM_BP_DIRECT | GO:0061844~antimicrobial humoral immune response mediated by | 0.412084764 | 0.011717388 | 20202, 14061, 15331, 20295, 22353 | 5.680517 |
| GOTERM_BP_DIRECT | GO:0045165~cell fate commitment | 0.412084764 | 0.011849083 | 104382, 18012, 18508, 12159 | 8.439626 |
| GOTERM_BP_DIRECT | GO:0043010~camera-type eye development | 0.412084764 | 0.011849083 | 23937, 18012, 18508, 12159 | 8.439626 |
| GOTERM_CC_DIRECT | GO:1990660~calprotectin complex | 0.434380863 | 0.010996984 | 20202, 20201 | 180.2771 |
| GOTERM_BP_DIRECT | GO:0021542~dentate gyrus development | 0.442243934 | 0.013069509 | 319352, 18012, 12227 | 17.1515 |

| | | | | | |
|---|---|---|---|---|---|
| UP_KW_BIOLOGICAL_PROCESS | KW-0892~Osteogenesis | 0.442274344 | 0.030501679 | 68713, 14560, 12159 | 10.82239 |
| GOTERM_BP_DIRECT | GO:0021983~pituitary gland development | 0.457637856 | 0.013889967 | 15364, 18508, 12159 | 16.61551 |
| GOTERM_BP_DIRECT | GO:0050729~positive regulation of | 0.471431722 | 0.014685173 | 20202, 20201, 12608, 16170 | 7.790424 |
| COG_ONTOLOGY | Defense mechanisms | 0.487901488 | 0.095402131 | 244562, 67473 | 18.41228 |
| GOTERM_BP_DIRECT | GO:0007214~gamma-aminobutyric acid signaling pathway | 0.488152274 | 0.015595919 | 319352, 14399, 14403 | 15.63813 |
| GOTERM_MF_DIRECT | GO:0001228~DNA-binding transcription activator activity, RNA | 0.495904125 | 0.015950133 | 14281, 104382, 68992, 18012, 12608, 15402, 18508, 18227, 18988 | 2.780797 |
| KEGG_PATHWAY | mmu04657:IL-17 signaling pathway | 0.499474302 | 0.005549714 | 20202, 14281, 20201, 12608, 20295 | 6.924873 |
| UP_KW_PTM | KW-0488~Methylation | 0.501327894 | 0.079157036 | 66497, 15361, 20202, 21681, 107071, 22289, 22138, 12608, 22186, | 1.681779 |
| GOTERM_BP_DIRECT | GO:0021986~habenula development | 0.502635884 | 0.016732558 | 18508, 18227 | 118.1548 |
| GOTERM_BP_DIRECT | GO:0035264~multicellular organism growth | 0.502635884 | 0.016861587 | 15361, 15364, 15402, 22289, 13498 | 5.092878 |
| GOTERM_BP_DIRECT | GO:0030900~forebrain development | 0.506497086 | 0.017395667 | 65100, 18508, 12667, 12159 | 7.308542 |
| GOTERM_BP_DIRECT | GO:0032967~positive regulation of collagen | 0.521101496 | 0.018313471 | 14061, 18591, 12159 | 14.37017 |
| UP_SEQ_FEATURE | DNA_BIND:Homeobox | 0.528205894 | 0.001147823 | 15285, 104382, 15425, 15414, 15402, 18508, | 4.973445 |
| UP_SEQ_FEATURE | DOMAIN:Homeobox | 0.528205894 | 0.001274321 | 15285, 104382, 15425, 15402, 18508, 18988, | 5.840526 |
| GOTERM_BP_DIR | GO:0006284~base- | 0.535872092 | 0.019260578 | 15361, 15364, 68240 | 13.99201 |
| GOTERM_BP_DIR | GO:0006935~chemotax | 0.546166528 | 0.021415166 | 68992, 20295, 13609, | 6.751701 |
| GOTERM_BP_DIRECT | GO:0006805~xenobiotic metabolic process | 0.546166528 | 0.021950313 | 14870, 13078, 232174, 27973 | 6.688005 |
| GOTERM_BP_DIRECT | GO:0045664~regulation of neuron | 0.546166528 | 0.02222209 | 18012, 18508, 13498 | 12.96821 |
| GOTERM_BP_DIRECT | GO:0070488~neutrophil aggregation | 0.546166528 | 0.022247998 | 20202, 20201 | 88.61607 |
| GOTERM_BP_DIRECT | GO:0051464~positive regulation of cortisol | 0.546166528 | 0.022247998 | 14419, 58991 | 88.61607 |
| GOTERM_BP_DIRECT | GO:0042305~specification of segmental identity, mandibular | 0.546166528 | 0.022247998 | 12667, 12159 | 88.61607 |

| | | | | | |
|---|---|---|---|---|---|
| GOTERM_BP_DIRECT | GO:0043542~endothelial cell migration | 0.559752013 | 0.023248486 | 20202, 20201, 13078 | 12.65944 |
| GOTERM_MF_DIRECT | GO:0046982~protein heterodimerization | 0.58093619 | 0.021587912 | 21973, 18012, 12608, 13197, 18591, 18227, | 3.230291 |
| GOTERM_MF_DIRECT | GO:0000981~DNA-binding transcription factor activity, RNA | 0.58093619 | 0.02208237 | 15285, 14281, 104382, 65100, 18012, 12608, 15425, 15414, 15402, | 2.085945 |
| INTERPRO | IPR001827:Homeobox_Antennapedia_CS | 0.58256565 | 0.00784223 | 15425, 15414, 15402 | 22.23077 |
| GOTERM_BP_DIRECT | GO:0007283~spermatogenesis | 0.589226004 | 0.024943273 | 15361, 78784, 232174, 15364, 237465, 20957, | 2.791057 |
| GOTERM_BP_DIRECT | GO:0001649~osteoblast differentiation | 0.60049703 | 0.025900032 | 14560, 13877, 12667, 12159 | 6.273704 |
| GOTERM_BP_DIRECT | GO:0002793~positive regulation of peptide | 0.609145959 | 0.027732683 | 20202, 20201 | 70.89286 |
| GOTERM_BP_DIRECT | GO:0018119~peptidyl-cysteine S-nitrosylation | 0.609145959 | 0.027732683 | 20202, 20201 | 70.89286 |
| GOTERM_BP_DIR | GO:0002790~peptide | 0.609145959 | 0.027732683 | 20202, 20201 | 70.89286 |
| GOTERM_BP_DIRECT | GO:0009791~post-embryonic development | 0.611252535 | 0.028316811 | 15285, 13498, 18227, 12159 | 6.059219 |
| GOTERM_MF_DIRECT | GO:0003714~transcription corepressor activity | 0.636570078 | 0.026282512 | 15364, 13498, 666528, 12227, 60406 | 4.442511 |
| GOTERM_MF_DIR | GO:0005125~cytokine | 0.636570078 | 0.029194894 | 14560, 20295, 16183, | 4.293004 |
| GOTERM_MF_DIRECT | GO:0001221~transcription coregulator binding | 0.636570078 | 0.030514153 | 15361, 14281, 18508 | 10.93402 |
| GOTERM_MF_DIRECT | GO:0005515~protein binding | 0.636570078 | 0.032199114 | 15361, 15364, 18012, 14693, 20957, 21648, 109095, 16170, 22138, 60406, 54698, 22097, 56410, 15402, 69684, 66373, 12236, 66395, 12159, 21973, 14281, 14061, 269855, 74469, 320051, 13197, 237465, | 1.344941 |
| GOTERM_MF_DIR | GO:0005179~hormone | 0.636570078 | 0.033503688 | 14419, 22353, 20604, | 5.669491 |
| UP_KW_BIOLOGICAL_PROCESS | KW-0769~Symport | 0.660603498 | 0.056948577 | 214111, 68267, 55961, 20538 | 4.524617 |
| GOTERM_MF_DIRECT | GO:0003723~RNA binding | 0.714011749 | 0.039667319 | 15361, 78784, 69878, 21681, 22186, 233871, 18508, 56009, 66373, | 2.185911 |
| GOTERM_BP_DIRECT | GO:0007507~heart development | 0.720969448 | 0.033975397 | 15361, 13411, 18591, 22289, 12159, 22138 | 3.344003 |
| UP_KW_BIOLOGICAL_PROCESS | KW-0508~mRNA splicing | 0.721276711 | 0.074614832 | 78784, 69878, 21681, 56009, 66373, 109095 | 2.634398 |

| | | | | | |
|---|---|---|---|---|---|
| GOTERM_CC_DIRECT | GO:0005622~intracellular anatomical structure | 0.782789787 | 0.023120373 | 18012, 22186, 18508, 16170 | 6.555531 |
| GOTERM_BP_DIRECT | GO:0050672~negative regulation of | 0.805671924 | 0.038610476 | 14419, 16183 | 50.63776 |
| GOTERM_CC_DIRECT | GO:0035985~senescence-associated | 0.807799006 | 0.027267477 | 15361, 15364 | 72.11084 |
| UP_KW_CELLULAR_COMPONENT | KW-0747~Spliceosome | 0.815897241 | 0.081589724 | 69878, 21681, 56009, 66373 | 3.911097 |
| GOTERM_BP_DIRECT | GO:0035914~skeletal muscle cell | 0.825386573 | 0.040814975 | 14281, 12227, 60406 | 9.328008 |
| GOTERM_BP_DIRECT | GO:0006357~regulation of transcription by RNA polymerase II | 0.825386573 | 0.04147096 | 14281, 104382, 68992, 65100, 15331, 12608, 15425, 434484, 15414, 15402, 18508, 54612, | 1.73757 |
| GOTERM_BP_DIRECT | GO:0048704~embryonic skeletal system | 0.825386573 | 0.042120328 | 15414, 15402, 12159 | 9.16718 |
| GOTERM_BP_DIR | GO:0035425~autocrine | 0.825386573 | 0.04400392 | 20202, 20201 | 44.30804 |
| GOTERM_BP_DIRECT | GO:0090402~oncogene-induced cell | 0.825386573 | 0.04400392 | 15361, 15364 | 44.30804 |
| GOTERM_BP_DIRECT | GO:0021978~telencephalon regionalization | 0.825386573 | 0.04400392 | 18508, 12159 | 44.30804 |
| GOTERM_BP_DIRECT | GO:0010468~regulation of gene expression | 0.825386573 | 0.044170048 | 14281, 14061, 57267, 18508, 22289, 18227, | 2.720669 |
| INTERPRO | IPR047575:Sm | 0.831008131 | 0.01278474 | 69878, 233871, 66373 | 17.2906 |
| GOTERM_BP_DIR | GO:0006397~mRNA | 0.834948373 | 0.045348634 | 78784, 21681, 56009, | 3.723364 |
| GOTERM_MF_DIRECT | GO:0051575~5'-deoxyribose-5- | 0.837761324 | 0.04899189 | 15361, 15364 | 39.68643 |
| GOTERM_MF_DIRECT | GO:0043565~sequence-specific DNA binding | 0.839966424 | 0.051576886 | 21973, 14281, 18012, 12608, 18508, 18227, | 2.615319 |
| GOTERM_BP_DIRECT | GO:0006355~regulation of DNA-templated transcription | 0.844394764 | 0.046536133 | 15361, 14281, 75316, 15364, 12608, 14369, 18508, 18227, 666528, | 2.016084 |
| GOTERM_BP_DIRECT | GO:0006406~mRNA export from nucleus | 0.849479785 | 0.047494876 | 21681, 56009, 109095 | 8.575749 |
| GOTERM_BP_DIRECT | GO:0034121~regulation of toll-like receptor | 0.870533003 | 0.049367287 | 20202, 20201 | 39.38492 |

| | | | | | |
|---|---|---|---|---|---|
| UP_KW_DOMAIN | KW-0732~Signal | 0.888752438 | 0.088875244 | 214111, 319352, 20295, 70355, 22353, 18591, 58991, 319433, 14419, 54698, 56410, 54615, 12477, 13024, 14399, 93703, 54612, 100312987, 12159, 11568, 623503, 69511, 245827, 21334, 14061, | 1.235013 |
| GOTERM_BP_DIRECT | GO:0030336~negative regulation of cell | 0.913558613 | 0.052536917 | 68713, 13078, 12667, 60406 | 4.72619 |
| GOTERM_BP_DIR | GO:0001764~neuron | 0.922791786 | 0.054259784 | 15285, 104382, 18508, | 4.664004 |
| GOTERM_BP_DIRECT | GO:0098869~cellular oxidant detoxification | 0.922791786 | 0.054542006 | 20202, 20201, 100503605 | 7.935768 |
| GOTERM_MF_DIRECT | GO:0003906~DNA-(apurinic or apyrimidinic site) | 0.925743944 | 0.05955078 | 15361, 15364 | 32.47072 |
| GOTERM_MF_DIR | GO:0031720~haptoglob | 0.933881011 | 0.069993152 | 110257, 100503605 | 27.47522 |
| GOTERM_MF_DIRECT | GO:0003690~double-stranded DNA binding | 0.933881011 | 0.070583389 | 14281, 18012, 18508, 20957 | 4.177519 |
| GOTERM_MF_DIRECT | GO:0062153~C5-methylcytidine- | 0.933881011 | 0.075171043 | 21681, 56009 | 25.51271 |
| GOTERM_MF_DIRECT | GO:0005344~oxygen carrier activity | 0.933881011 | 0.08544103 | 110257, 100503605 | 22.32362 |
| GOTERM_MF_DIRECT | GO:0003729~mRNA binding | 0.933881011 | 0.089658805 | 78784, 21681, 233871, 56009, 109095 | 2.947012 |
| GOTERM_MF_DIR | GO:0020037~heme | 0.933881011 | 0.093032046 | 110257, 13078, 232174, | 3.701326 |
| GOTERM_MF_DIRECT | GO:0003682~chromatin binding | 0.933881011 | 0.096569453 | 15361, 21973, 14281, 15331, 18012, 12608, | 2.212607 |
| GOTERM_BP_DIRECT | GO:0007435~salivary gland morphogenesis | 0.975656852 | 0.060004455 | 12388, 18508 | 32.22403 |
| GOTERM_BP_DIRECT | GO:1904628~cellular response to phorbol 13- | 0.975656852 | 0.060004455 | 14281, 12227 | 32.22403 |
| GOTERM_BP_DIRECT | GO:0002320~lymphoid progenitor cell | 0.975656852 | 0.060004455 | 15361, 12159 | 32.22403 |
| GOTERM_CC_DIRECT | GO:0090575~RNA polymerase II | 0.995798319 | 0.040376698 | 15361, 14281, 18012, 12608 | 5.263565 |
| GOTERM_CC_DIR | GO:0031982~vesicle | 0.995798319 | 0.066183001 | 54615, 67473, 66395, | 4.292312 |
| GOTERM_CC_DIR | GO:0005833~hemoglob | 0.995798319 | 0.069366354 | 110257, 100503605 | 27.73494 |
| GOTERM_CC_DIR | GO:0034709~methylos | 0.995798319 | 0.074499726 | 69878, 13877 | 25.75387 |
| GOTERM_CC_DIRECT | GO:0031838~haptoglobin-hemoglobin complex | 0.995798319 | 0.074499726 | 110257, 100503605 | 25.75387 |
| GOTERM_CC_DIRECT | GO:0043025~neuronal cell body | 0.995798319 | 0.077487623 | 14419, 13166, 22353, 18591, 20604, 21648, | 2.159007 |

| | | | | | |
|---|---|---|---|---|---|
| GOTERM_CC_DIRECT | GO:0005667~transcription regulator complex | 0.995798319 | 0.093044535 | 14281, 18508, 18227, 666528, 18988 | 2.907695 |
| GOTERM_CC_DIRECT | GO:1902711~GABA-A receptor complex | 0.995798319 | 0.099747465 | 14399, 14403 | 18.97654 |
| GOTERM_BP_DIRECT | GO:0043922~negative regulation by host of | 0.998405104 | 0.065278587 | 15364, 18988 | 29.53869 |
| GOTERM_BP_DIRECT | GO:0032024~positive regulation of insulin | 0.998405104 | 0.066518671 | 14061, 22044, 58991 | 7.089286 |
| GOTERM_BP_DIRECT | GO:0042593~glucose homeostasis | 0.998405104 | 0.069451973 | 15361, 13166, 18012, 18508 | 4.219813 |
| GOTERM_BP_DIRECT | GO:0003007~heart morphogenesis | 0.998405104 | 0.069638104 | 22289, 12159, 22138 | 6.905148 |
| GOTERM_BP_DIRECT | GO:0035881~amacrine cell differentiation | 0.998405104 | 0.070523302 | 104382, 18012 | 27.26648 |
| GOTERM_BP_DIRECT | GO:0060644~mammary gland epithelial cell | 0.998405104 | 0.075738765 | 12608, 15402 | 25.31888 |
| GOTERM_BP_DIRECT | GO:0001568~blood vessel development | 0.998405104 | 0.079273627 | 18508, 18591, 12159 | 6.405981 |
| GOTERM_BP_DIRECT | GO:0048661~positive regulation of smooth | 0.998405104 | 0.082573193 | 13609, 18591, 12159 | 6.255252 |
| GOTERM_BP_DIR | GO:0007611~learning | 0.998405104 | 0.084238654 | 13411, 22353, 18227 | 6.182517 |
| GOTERM_BP_DIRECT | GO:0043508~negative regulation of JUN | 0.998405104 | 0.086082581 | 14870, 54612 | 22.15402 |
| GOTERM_BP_DIRECT | GO:0007406~negative regulation of neuroblast | 0.998405104 | 0.086082581 | 18508, 12227 | 22.15402 |
| GOTERM_BP_DIRECT | GO:0030194~positive regulation of blood | 0.998405104 | 0.086082581 | 20202, 14061 | 22.15402 |
| GOTERM_BP_DIRECT | GO:0045580~regulation of T cell differentiation | 0.998405104 | 0.091211255 | 54698, 232174 | 20.85084 |
| GOTERM_BP_DIRECT | GO:0014002~astrocyte development | 0.998405104 | 0.091211255 | 20202, 20201 | 20.85084 |
| GOTERM_BP_DIRECT | GO:0048712~negative regulation of astrocyte | 0.998405104 | 0.091211255 | 14061, 15364 | 20.85084 |
| GOTERM_BP_DIRECT | GO:0009636~response to toxic substance | 0.998405104 | 0.092716105 | 14281, 14870, 13078 | 5.842818 |
| GOTERM_BP_DIRECT | GO:0001768~establishment of T cell polarity | 0.998405104 | 0.096311319 | 54698, 232174 | 19.69246 |
| GOTERM_BP_DIRECT | GO:0021953~central nervous system neuron | 0.998405104 | 0.096311319 | 15285, 18227 | 19.69246 |
| GOTERM_BP_DIRECT | GO:0032496~response to lipopolysaccharide | 0.998405104 | 0.097205378 | 20202, 14281, 20201, 12608 | 3.635531 |
| UP_SEQ_FEATURE | DISULFID:Interchain | 1 | 0.003765313 | 13166, 14560, 56410, 12477, 18591, 12159 | 5.799826 |

| | | | | | |
|---|---|---|---|---|---|
| UP_SEQ_FEATURE | REGION:Disordered | 1 | 0.005363172 | 214111, 242785, 104382, 68992, 18012, 14693, 99010, 381232, 55961, 15425, 93703, 12159, 108115, 245827, 269855, 237465, 12608, 13411, 60364, 14588, 13498, 666528, 15361, 319352, 68499, 15364, 107071, 18591, 20957, 109095, 16170, 270150, 60406, 15402, 383435, 11568, 14281, 69993, 69878, 14560, 115488471, 16183, 240514, 244562, 12388, 12667, 100034363, 434794, 66497, 107971, 13166, 70355, 27206, 20538, 58991, 228715, 57267, 13609, 100038416, 12236, 320051, 22044, 100862349, 22289, | 1.187117 |
| UP_SEQ_FEATURE | COMPBIAS:Polar residues | 1 | 0.007343125 | 66497, 107971, 214111, 104382, 13166, 18012, 70355, 27206, 99010, 57267, 13609, 93703, 100038416, 12236, 269855, 320051, 100862349, 22289, 100303741, 54324, 12608, 60364, 18508, 13498, 666528, 18988, 15285, 15361, 319352, 65100, 15364, 21681, 98267, 18591, 20957, | 1.345864 |
| UP_SEQ_FEATUR | MOTIF:Antp-type | 1 | 0.009820858 | 15425, 15414, 15402 | 19.81607 |
| UP_SEQ_FEATUR | DOMAIN:Sm | 1 | 0.012337715 | 69878, 233871, 66373 | 17.61429 |
| KEGG_PATHWAY | mmu04950:Maturity onset diabetes of the | 1 | 0.017964085 | 15285, 18012, 18508 | 14.3114 |
| INTERPRO | IPR000116:HMGA | 1 | 0.019022438 | 15361, 15364 | 103.7436 |
| BIOCARTA | m_dbpbPathway:Transcriptional activation of | 1 | 0.039872276 | 14061, 18591 | 45.92857 |
| INTERPRO | IPR025715:FoP_C | 1 | 0.043827863 | 21681, 56009 | 44.46154 |

| | | | | | |
|---|---|---|---|---|---|
| UP_SEQ_FEATURE | COMPBIAS:Pro residues | 1 | 0.0487056 | 319352, 104382, 68992, 65100, 269855, 14693, 22289, 109095, 22138, 78784, 57267, 54324, | 1.65438 |
| SMART | SM01218:FoP_duplicati | 1 | 0.053741775 | 21681, 56009 | 35.94089 |
| UP_SEQ_FEATUR | DNA_BIND:A.T hook 3 | 1 | 0.054984607 | 15361, 15364 | 35.22857 |
| INTERPRO | IPR017956:AT_hook_DN | 1 | 0.055995995 | 15361, 15364 | 34.5812 |
| KEGG_PATHWAY | mmu05200:Pathways in cancer | 1 | 0.057623571 | 14281, 14061, 14870, 14693, 13197, 14369, | 2.115372 |
| UP_SEQ_FEATUR | DNA_BIND:A.T hook 1 | 1 | 0.066789752 | 15361, 15364 | 28.82338 |
| UP_SEQ_FEATUR | DNA_BIND:A.T hook 2 | 1 | 0.066789752 | 15361, 15364 | 28.82338 |
| SMART | SM00384:AT_hook | 1 | 0.068565209 | 15361, 15364 | 27.95402 |
| KEGG_PATHWAY | mmu04060:Cytokine-cytokine receptor | 1 | 0.074523774 | 100041579, 14560, 20295, 16183, 12159, | 2.628625 |
| INTERPRO | IPR050056:Hemoglobin_oxygen_transport | 1 | 0.079872869 | 110257, 100503605 | 23.94083 |
| KEGG_PATHWAY | mmu05217:Basal cell | 1 | 0.083896253 | 13197, 14369, 12159 | 6.133459 |
| INTERPRO | IPR029034:Cystine- | 1 | 0.084918645 | 14560, 18591, 12159 | 6.142713 |
| UP_KW_LIGAND | KW-0349~Heme | 1 | 0.088522429 | 110257, 13078, 232174, | 3.683336 |
| KEGG_PATHWAY | mmu04024:cAMP signaling pathway | 1 | 0.09260797 | 14281, 98932, 22353, 20604, 58991 | 2.875059 |
| UP_SEQ_FEATUR | DOMAIN:Globin | 1 | 0.095665956 | 110257, 100503605 | 19.81607 |
| INTERPRO | IPR012292:Globin/Prot | 1 | 0.097386096 | 110257, 100503605 | 19.45192 |
| INTERPRO | IPR000971:Globin | 1 | 0.097386096 | 110257, 100503605 | 19.45192 |
| INTERPRO | IPR009050:Globin- | 1 | 0.097386096 | 110257, 100503605 | 19.45192 |
