## Supplemental Table 4 for "Early Life Adversity Produces Enduring Molecular and Functional Disruption of Developing Vagal Circuits"

**P9 EXCITATORY NEURONS FEMALE**

| Category | Term | FDR | PValue | Genes | Fold Enrichment |
|---|---|---|---|---|---|
| UP_KW_CELLULAR_COMPONENT | KW-0964~Secreted | 5.19E-07 | 1.78996E-08 | 18159, 72014, 23829, 67445, 67405, 24117, 116903, 224065, 12759, 27220, 14419, 21819, 12310, 12653, 18619, 109648, 69683, 15450, 19215, 21333, 14221, 13033, 18188, 66208, 14060, 18081, 241327, 330483, 30052, 18205, 53419, 14526 | 3.039328 |
| UP_KW_PTM | KW-0165~Cleavage on pair of basic residues | 7.56E-07 | 3.2867E-08 | 18159, 21333, 67405, 116903, 224065, 27220, 14419, 30052, 12310, 12653, 18619, 109648, 18205, 14526 | 7.5360399 |
| UP_KW_MOLECULAR_FUNC | KW-0527~Neurope | 7.6E-06 | 1.65122E-07 | 27220, 14419, 21333, 30052, 18619, 109648 | 45.0257353 |
| GOTERM_CC_DIRECT | GO:0030424~axon | 1.3E-05 | 6.07311E-08 | 68566, 21333, 18039, 13166, 12374, 19132, 18188, 110012, 20618, 20508, 15507, 12310, 18619, 12215, 18205, 68350 | 6.10345443 |
| GOTERM_BP_DIRECT | GO:0007218~neuropeptide signaling | 5.1E-05 | 5.54E-08 | 27220, 14419, 21333, 104443, 30052, 12310, 67405, 18619, 109648 | 17.1977281 |
| UP_KW_PTM | KW-0027~Amidatio | 0.00015 | 1.30752E-05 | 14419, 21333, 12310, 109648, 14526, 116903 | 19.3049853 |
| GOTERM_CC_DIRECT | GO:0005615~extracellular space | 0.000204 | 1.90878E-06 | 18159, 13386, 13166, 15122, 67445, 116903, 12759, 27220, 14419, 21819, 12310, 20194, 12653, 109648, 15450, 209294, 19215, 21333, 14221, 13033, 18188, 66208, 241327, 330483, 30052, 18205, 14526, 14109 | 2.79449058 |
| UP_KW_MOLECULAR_FUNC | KW-0372~Hormone | 0.000824 | 3.58317E-05 | 224065, 18159, 14419, 21819, 12310, 14526, 116903 | 11.0589525 |
| GOTERM_MF_DIRECT | GO:0005179~hormone activity | 0.000928 | 3.2318E-06 | 224065, 18159, 14419, 21819, 12310, 109648, 14526, 116903 | 12.7465791 |
| GOTERM_CC_DIRECT | GO:0005576~extracellular region | 0.001493 | 2.09321E-05 | 18159, 19215, 21333, 72014, 23829, 67405, 14060, 24117, 18081, 116903, 241327, 224065, 12759, 14419, 21819, 12310, 12653, 18619, 53419, 18205, 14526, 69683 | 2.89712651 |

| | | | | | |
|---|---|---|---|---|---|
| GOTERM_MF_DIRECT | GO:0005184~neuropeptide | 0.001538 | 1.07208E-05 | 27220, 18159, 14419, 67405, 109648 | 35.8497537 |
| UP_KW_DOMAIN | KW-0732~Signal | 0.004155 | 0.000259663 | 69742, 18159, 12373, 13386, 12374, 72014, 23829, 67445, 246048, 67405, 24117, 73382, 116903, 224065, 12759, 27220, 330173, 14419, 21819, 12310, 12653, 14399, 18619, 109648, 12215, 69683, 72309, 15450, 23934, 19711, 23936, 19215, 21333, 14221, 13033, 18188, 66208, 27973, 66647, 14060, 241327, 330483, 30052, 18205, 14526, 14227 | 1.58474178 |
| KEGG_PATHWAY | mmu04080:Neuroactive ligand-receptor | 0.006002 | 3.31625E-05 | 224065, 23936, 14419, 21333, 104443, 12310, 67405, 14399, 18619, 109648, 14526, 116903 | 4.70625 |
| GOTERM_CC_DIRECT | GO:0043025~neuronal cell body | 0.028925 | 0.000540661 | 14419, 21333, 75406, 12374, 19132, 13166, 12310, 20618, 67405, 18619, 16438, 68350 | 3.5839521 |
| GOTERM_CC_DIRECT | GO:0043195~terminal bouton | 0.05231 | 0.001222187 | 19132, 13166, 12310, 20508, 109648 | 10.7261649 |
| GOTERM_BP_DIRECT | GO:0007268~chemical | 0.059314 | 0.000128944 | 27220, 21333, 110012, 20618, 20508, 18619, 109648 | 9.03093722 |
| GOTERM_CC_DIRECT | GO:0045202~synapse | 0.060483 | 0.001785732 | 12759, 27220, 23934, 14704, 23936, 21333, 110012, 20618, 14399, 18205, 67184, 16438 | 3.08913548 |
| GOTERM_CC_DIRECT | GO:0043679~axon terminus | 0.060483 | 0.001978403 | 12374, 20618, 20508, 67405, 18619 | 9.41069182 |
| GOTERM_CC_DIRECT | GO:0030141~secretory granule | 0.081999 | 0.003065392 | 27220, 18159, 19711, 14419, 30052, 12653 | 6.10734694 |
| UP_KW_MOLECULAR_FUNC | KW-0143~Chaperon | 0.095698 | 0.006241155 | 12759, 100126824, 15507, 30055, 11927, 14356 | 5.04961517 |
| GOTERM_CC_ | GO:0098981~c | 0.126695 | 0.005328311 | 231290, 18039, 20508 | 27.2054545 |
| UP_KW_CELLULAR_COMPONENT | KW-0968~Cytoplasmic vesicle | 0.129227 | 0.011098296 | 12759, 68566, 19711, 378462, 13166, 20508, 67405, 18619, 12684, 109648, 16438 | 2.52921751 |
| UP_KW_CELLULAR_COMPO | KW-0999~Mitochon | 0.129227 | 0.013368345 | 12859, 100126824, 12868, 75406, 30055, 14356, 67184 | 3.57183789 |
| GOTERM_MF_DIRECT | GO:0048018~receptor ligand | 0.161558 | 0.001997922 | 12759, 21333, 67405, 18619, 14526 | 9.38124396 |

| | | | | | |
|---|---|---|---|---|---|
| GOTERM_MF_DIRECT | GO:0044877~protein-containing | 0.161558 | 0.002251678 | 15461, 12759, 68566, 14281, 18039, 19132, 15122, 21819, 12310, 14399, 16438 | 3.21916156 |
| GOTERM_CC_ | GO:0034466~c | 0.212572 | 0.009933272 | 13166, 18619 | 199.506667 |
| UP_KW_MOLECULAR_FUNC | KW-0529~Neurotra | 0.249626 | 0.021706628 | 27220, 21333 | 90.0514706 |
| UP_KW_PTM | KW-1015~Disulfide bond | 0.286963 | 0.037429986 | 18159, 104443, 13386, 12374, 13166, 246048, 24117, 73382, 116903, 224065, 12759, 27220, 21819, 12310, 12653, 14399, 14356, 18619, 12215, 16438, 23934, 100126824, 23936, 19215, 13033, 18188, 27973, 14060, 241327, 330483, 30055, 18205, 53419 | 1.36682459 |
| GOTERM_CC_DIRECT | GO:0030425~dendrite | 0.301319 | 0.015488384 | 14704, 23936, 15507, 13166, 110012, 14399, 18619, 18205, 16438 | 2.79247278 |
| GOTERM_CC_DIRECT | GO:0005743~mitochondrial inner | 0.321788 | 0.018813231 | 12759, 12859, 100126824, 12868, 75406, 30055, 14356, 67184 | 2.95565432 |
| GOTERM_CC_DIRECT | GO:0031966~mitochondrial | 0.321788 | 0.019547849 | 12759, 12859, 12868, 52668, 67184 | 4.86601626 |
| UP_KW_PTM | KW-0325~Glycoprotein | 0.353221 | 0.061429668 | 69742, 231290, 18039, 12373, 104443, 13386, 12374, 13166, 72014, 246048, 57780, 24117, 73382, 12759, 21819, 12310, 54218, 12653, 14399, 12215, 72309, 15450, 23934, 23936, 19215, 14221, 13033, 67874, 20508, 14060, 18081, 241327, 330483, 12608, 18205, 53419, 13732, 108902 | 1.27238787 |
| GOTERM_CC_DIRECT | GO:0042719~mitochondrial | 0.376788 | 0.024649703 | 30055, 14356 | 79.8026667 |
| GOTERM_CC_DIRECT | GO:0043204~perikaryon | 0.387018 | 0.0271274 | 19711, 15507, 19132, 18619, 109648 | 4.39441997 |
| KEGG_PATHWAY | mmu04725:Cholinergic | 0.406692 | 0.005943011 | 15461, 14704, 14281, 20508, 16438 | 6.76783739 |
| KEGG_PATHWAY | mmu05022:Pathways of neurodegenerat | 0.406692 | 0.011525558 | 15461, 12859, 12868, 18039, 75406, 19132, 19173, 67184, 16438 | 2.86190878 |
| KEGG_PATHWAY | mmu05010:Alzheimer disease | 0.406692 | 0.012079648 | 15461, 12859, 12868, 75406, 19173, 14433, 67184, 16438 | 3.14556555 |

| | | | | | |
|---|---|---|---|---|---|
| KEGG_PATHWAY | mmu05415:Diabetic | 0.406692 | 0.012680346 | 12859, 12868, 75406, 13033, 14433, 67184 | 4.22911866 |
| KEGG_PATHWAY | mmu05208:Chemical | 0.406692 | 0.01569268 | 15461, 12859, 14281, 12868, 75406, 67184 | 4.00750546 |
| KEGG_PATHWAY | mmu04723:Retrograde | 0.406692 | 0.016752112 | 14704, 75406, 14399, 67184, 16438 | 4.99846814 |
| KEGG_PATHWAY | mmu04714:Thermogenesis | 0.406692 | 0.018244004 | 15461, 12859, 12868, 75406, 14526, 67184 | 3.85596113 |
| KEGG_PATHWAY | mmu04932:Non-alcoholic | 0.406692 | 0.02022224 | 12859, 14281, 12868, 75406, 67184 | 4.72077546 |
| BIOCARTA | m_mta3Pathway:Downregulate | 0.445758 | 0.0096006 | 15507, 13033, 14433 | 17.5363636 |
| BIOCARTA | m_il6Pathway:I | 0.445758 | 0.013715635 | 15461, 14281, 12608 | 14.6136364 |
| KEGG_PATHWAY | mmu05020:Prion disease | 0.496875 | 0.030737165 | 12859, 12868, 75406, 19173, 67184, 16438 | 3.36160714 |
| KEGG_PATHWAY | mmu05012:Parkinson disease | 0.496875 | 0.031157257 | 12859, 12868, 75406, 19173, 67184, 16438 | 3.3493385 |
| KEGG_PATHWAY | mmu05014:Amyotrophic | 0.496875 | 0.032941991 | 12859, 12868, 18039, 75406, 19132, 19173, 67184 | 2.855125 |
| GOTERM_BP_ | GO:0032099~n | 0.551307 | 0.001797742 | 27220, 66208, 14526 | 46.7425432 |
| GOTERM_MF_ | GO:0031716~c | 0.566492 | 0.009869206 | 12310, 116903 | 200.758621 |
| GOTERM_CC_ | GO:0034774~s | 0.570242 | 0.043934142 | 13166, 14526 | 44.3348148 |
| GOTERM_CC_ | GO:1904115~a | 0.570242 | 0.045299563 | 68566, 15507, 18039 | 8.80176471 |
| UP_KW_MOLECULAR_FUNC | KW-0838~Vasoactiv | 0.586378 | 0.06373678 | 18159, 67405 | 30.0171569 |
| UP_KW_CELLULAR_COMPONENT | KW-0256~Endoplasmic reticulum | 0.588126 | 0.081120876 | 14281, 19711, 23936, 19215, 66208, 27973, 12759, 230904, 12684, 12215, 69683, 14227, 16438 | 1.6797921 |
| KEGG_PATHWAY | mmu05016:Huntington | 0.606925 | 0.047641033 | 12859, 12868, 75406, 19173, 67184, 16438 | 2.97960633 |
| KEGG_PATHW | mmu04915:Estr | 0.606925 | 0.054995522 | 15461, 14281, 13033, 16438 | 4.56576493 |
| KEGG_PATHW | mmu04210:Apo | 0.606925 | 0.055995414 | 15461, 14281, 13033, 16438 | 4.53194444 |
| KEGG_PATHW | mmu04730:Lon | 0.606925 | 0.056853506 | 15461, 19051, 16438 | 7.64765625 |
| KEGG_PATHW | mmu04728:Do | 0.606925 | 0.057004035 | 68566, 14704, 14281, 16438 | 4.49862132 |
| KEGG_PATHW | mmu00190:Oxi | 0.622856 | 0.062176745 | 12859, 12868, 75406, 67184 | 4.33909574 |
| KEGG_PATHWAY | mmu04270:Vascular smooth | 0.622856 | 0.065382656 | 18159, 12310, 16438, 116903 | 4.24869792 |
| KEGG_PATHWAY | mmu05170:Human | 0.626243 | 0.069198087 | 15461, 14704, 14281, 22033, 16438 | 3.17330135 |
| GOTERM_CC_ | GO:0098982~G | 0.638616 | 0.056452336 | 13033, 14399, 109648, 16438 | 4.58636015 |
| GOTERM_CC_DIRECT | GO:0005783~endoplasmic reticulum | 0.638616 | 0.056699536 | 12759, 14281, 19711, 23936, 13166, 66208, 230904, 27973, 12684, 69683, 14227, 16438 | 1.86309728 |

| | | | | | |
|---|---|---|---|---|---|
| GOTERM_CC_ | GO:0097060~s | 0.640368 | 0.061815148 | 75406, 18081, 16438 | 7.3891358 |
| GOTERM_CC_ | GO:0005883~n | 0.640368 | 0.062839802 | 18039, 19132 | 30.6933333 |
| GOTERM_CC_ | GO:0043209~m | 0.676886 | 0.069586363 | 12859, 18039, 15122, 14433 | 4.20014035 |
| GOTERM_BP_ DIRECT | GO:0043524~n egative | 0.681526 | 0.003546199 | 15461, 12759, 18159, 12608, 16392, 18205 | 5.89954428 |
| GOTERM_BP_ | GO:0014823~re | 0.681526 | 0.004216845 | 14281, 100126824, 18935, 14526 | 12.2758194 |
| GOTERM_BP_ DIRECT | GO:0090280~p ositive | 0.681526 | 0.004717298 | 12374, 16392, 14526 | 28.9358601 |
| GOTERM_BP_ | GO:0048265~re | 0.681526 | 0.00718457 | 21333, 13166, 12310 | 23.3712716 |
| GOTERM_BP_ DIRECT | GO:0061844~a ntimicrobial humoral | 0.681526 | 0.007410184 | 21333, 67405, 14433, 109648, 14109 | 6.49201988 |
| GOTERM_BP_ DIRECT | GO:0031640~ki lling of cells of | 0.681526 | 0.007970836 | 21333, 67405, 14433, 18081 | 9.76149496 |
| GOTERM_BP_ | GO:0008217~re | 0.681526 | 0.008236806 | 224065, 21333, 109648, 53419 | 9.64528669 |
| GOTERM_BP_ DIRECT | GO:0045766~p ositive | 0.681526 | 0.00826139 | 15461, 15507, 16392, 69683, 18081 | 6.29040436 |
| GOTERM_BP_ DIRECT | GO:0097746~bl ood vessel | 0.681526 | 0.008889472 | 224065, 18159, 67405 | 20.9535538 |
| GOTERM_MF_ DIRECT | GO:0061629~R NA polymerase | 0.704645 | 0.017634985 | 14281, 15507, 12608, 52668, 16392 | 5.01896552 |
| GOTERM_MF_ | GO:0031628~o | 0.704645 | 0.019641681 | 104443, 18619 | 100.37931 |
| GOTERM_MF_ | GO:0016531~c | 0.704645 | 0.019641681 | 100126824, 11927 | 100.37931 |
| GOTERM_BP_ DIRECT | GO:0045944~p ositive regulation of transcription by | 0.748159 | 0.010571809 | 15461, 14281, 18991, 70261, 16392, 57756, 24074, 14419, 12608, 18935, 20194, 68040, 18205, 15207 | 2.21194562 |
| GOTERM_BP_ | GO:0007631~fe | 0.750058 | 0.011413921 | 14419, 12310, 109648 | 18.4137291 |
| GOTERM_CC_ | GO:0043292~c | 0.842083 | 0.090504207 | 15507, 66106 | 21.0007018 |
| UP_KW_LIGAN | KW- | 0.849343 | 0.038606504 | 100126824, 13166, 11927 | 9.30853994 |
| GOTERM_BP_ DIRECT | GO:0061077~c haperone- | 0.903608 | 0.01495658 | 12759, 15507, 14227 | 15.99087 |
| GOTERM_BP_ DIRECT | GO:0097193~in trinsic | 0.903608 | 0.015714914 | 15461, 12759, 52668 | 15.5808477 |
| INTERPRO | IPR015476:Calc | 1 | 0.010666829 | 12310, 116903 | 185.441667 |
| UP_SEQ_FEAT | DOMAIN:Calcit | 1 | 0.011053611 | 12310, 116903 | 178.983871 |
| UP_SEQ_FEAT | SITE:Cleavage; | 1 | 0.011053611 | 21333, 67405 | 178.983871 |

| | | | | | |
|---|---|---|---|---|---|
| UP_SEQ_FEATURE | CARBOHYD:N-linked (GlcNAc...) asparagine | 1 | 0.011564821 | 69742, 231290, 12373, 104443, 13386, 12374, 13166, 72014, 246048, 24117, 73382, 12759, 21819, 12310, 54218, 14399, 12215, 72309, 15450, 23934, 19215, 14221, 13033, 67874, 20508, 14060, 18081, 330483, 18205, 53419, 13732, 108902 | 1.53387356 |
| INTERPRO | IPR021117:Calc | 1 | 0.015957857 | 12310, 116903 | 123.627778 |
| INTERPRO | IPR001693:Calc | 1 | 0.015957857 | 12310, 116903 | 123.627778 |
| INTERPRO | IPR018360:Calc | 1 | 0.015957857 | 12310, 116903 | 123.627778 |
| SMART | SM00113:CALC | 1 | 0.016087673 | 12310, 116903 | 121.6 |
| UP_SEQ_FEAT | SITE:Cleavage; | 1 | 0.016534884 | 21333, 67405 | 119.322581 |
| UP_SEQ_FEAT | DOMAIN:Tim10- | 1 | 0.016534884 | 30055, 14356 | 119.322581 |
| INTERPRO | IPR017970:Homeobox_CS | 1 | 0.016904329 | 18991, 18935, 16392, 14912, 15416 | 5.0667122 |
| GOTERM_BP_DIRECT | GO:0006874~intracellular | 1 | 0.020014248 | 18159, 12373, 12374, 12310 | 6.92482121 |
| GOTERM_BP_DIRECT | GO:0045892~negative regulation of | 1 | 0.020938746 | 386655, 24074, 18991, 12608, 12310, 16392, 14912, 67184, 15207 | 2.63814643 |
| INTERPRO | IPR021116:Calc | 1 | 0.021220824 | 12310, 116903 | 92.7208333 |
| GOTERM_BP_DIRECT | GO:0032760~positive | 1 | 0.022320007 | 12759, 15507, 67445, 16392 | 6.64101706 |
| GOTERM_BP_DIRECT | GO:0051480~regulation of | 1 | 0.02416016 | 12310, 16438, 116903 | 12.4010829 |
| GOTERM_BP_ | GO:0019233~s | 1 | 0.026028493 | 21333, 18619, 15416 | 11.9147659 |
| UP_SEQ_FEAT | SITE:Cleavage; | 1 | 0.02740719 | 21333, 109648 | 71.5935484 |
| GOTERM_BP_DIRECT | GO:0000122~negative regulation of | 1 | 0.028219551 | 386655, 24074, 70103, 18991, 12608, 52668, 66647, 16392, 14912, 15207, 15416 | 2.19513421 |
| GOTERM_BP_DIRECT | GO:0006366~transcription by | 1 | 0.031221993 | 14281, 24074, 12608, 16392, 15207 | 4.20230333 |
| GOTERM_MF_DIRECT | GO:0005515~protein binding | 1 | 0.03255767 | 15461, 14570, 18039, 13386, 23829, 20618, 22033, 246048, 20957, 16392, 12759, 24074, 18935, 20194, 12653, 14433, 12215, 72309, 16438, 14281, 23936, 21333, 13033, 19132, 70103, 110012, 20508, 66208, 66647, 14060, 57756, 330483, 30052, 12608, 230904, 18205, 14109, 15416 | 1.36790884 |

| | | | | | |
|---|---|---|---|---|---|
| GOTERM_BP_ DIRECT | GO:0008284~p ositive | 1 | 0.032594273 | 15461, 12759, 12374, 22033, 70261, 109648, 16392, 18205 | 2.63053273 |
| GOTERM_BP_ DIRECT | GO:0030182~n euron | 1 | 0.036292894 | 14281, 12608, 18935, 16392, 53419 | 4.00298459 |
| INTERPRO | IPR004217:Tim | 1 | 0.036842835 | 30055, 14356 | 52.9833333 |
| INTERPRO | IPR035427:Tim | 1 | 0.036842835 | 30055, 14356 | 52.9833333 |
| INTERPRO | IPR050405:Inter | 1 | 0.036842835 | 18039, 19132 | 52.9833333 |
| INTERPRO | IPR006821:Inter | 1 | 0.036842835 | 18039, 19132 | 52.9833333 |
| GOTERM_BP_ DIRECT | GO:0060261~p ositive | 1 | 0.037284207 | 67224, 24074, 70103 | 9.80085583 |
| UP_SEQ_FEAT | MOTIF:Twin | 1 | 0.038160268 | 30055, 14356 | 51.1382488 |
| GOTERM_BP_ DIRECT | GO:0099538~s ynaptic | 1 | 0.043280939 | 18619, 109648 | 45.0113379 |
| GOTERM_MF_ | GO:1903136~c | 1 | 0.043655824 | 11927, 20618 | 44.6130268 |
| GOTERM_MF_ | GO:0005507~c | 1 | 0.04477046 | 100126824, 13166, 11927 | 8.85699797 |
| GOTERM_BP_ | GO:0001503~o | 1 | 0.046407151 | 18159, 12374, 13386, 12310 | 4.97057719 |
| GOTERM_MF_ | GO:0001540~a | 1 | 0.047162781 | 12759, 15122, 230904 | 8.60394089 |
| GOTERM_BP_ | GO:0046850~re | 1 | 0.047973575 | 27220, 13386 | 40.5102041 |
| GOTERM_BP_ | GO:0008343~a | 1 | 0.047973575 | 27220, 109648 | 40.5102041 |
| GOTERM_BP_ DIRECT | GO:0045039~pr otein insertion | 1 | 0.047973575 | 30055, 67184 | 40.5102041 |
| GOTERM_MF_ DIRECT | GO:0042802~id entical protein binding | 1 | 0.051494473 | 15450, 14281, 18039, 12374, 22033, 241327, 15507, 330483, 21819, 12608, 12310, 52668, 14433, 12684, 14526, 16438, 68350 | 1.64397714 |
| SMART | SM00389:HOX | 1 | 0.052448127 | 18991, 18935, 16392, 14912, 15416 | 3.48091603 |
| GOTERM_BP_ DIRECT | GO:0032024~p ositive | 1 | 0.052520818 | 12374, 16392, 16438 | 8.10204082 |
| GOTERM_BP_ | GO:0071305~c | 1 | 0.05264335 | 12374, 18619 | 36.8274583 |
| UP_SEQ_FEAT URE | DNA_BIND:Hom eobox | 1 | 0.053673331 | 18991, 18935, 16392, 14912, 15416 | 3.50948767 |
| INTERPRO | IPR001356:HD | 1 | 0.055316364 | 18991, 18935, 16392, 14912, 15416 | 3.47269039 |
| GOTERM_MF_ DIRECT | GO:1990837~s equence- | 1 | 0.055780346 | 14281, 18991, 12608, 18935, 16392, 14912, 15416 | 2.55976383 |
| GOTERM_BP_ DIRECT | GO:0007204~p ositive | 1 | 0.056746152 | 14704, 21333, 12310, 16438 | 4.57742419 |
| UP_KW_BIOLO GICAL_PROCE | KW- 0053~Apoptosi | 1 | 0.057778153 | 22033, 14433, 12684, 67184, 16438, 68350 | 2.79009581 |
| GOTERM_MF_ DIRECT | GO:0005102~si gnaling | 1 | 0.058109365 | 12759, 18159, 23829, 21819, 12310, 24117 | 2.8679803 |

| | | | | | |
|---|---|---|---|---|---|
| GOTERM_MF_ DIRECT | GO:0001664~G protein- | 1 | 0.058479307 | 224065, 15122, 109648 | 7.62374509 |
| GOTERM_BP_ DIRECT | GO:0033138~p ositive | 1 | 0.06149092 | 16392, 18205, 14526 | 7.41040319 |
| GOTERM_BP_ | GO:0016486~p | 1 | 0.061914762 | 30052, 53419 | 31.1616954 |
| INTERPRO | IPR050822:Cer | 1 | 0.062331936 | 67445, 23829 | 30.9069444 |
| UP_SEQ_FEAT | REGION:Linker | 1 | 0.064529818 | 18039, 19132 | 29.8306452 |
| GOTERM_BP_ DIRECT | GO:0097696~c ell surface | 1 | 0.066516618 | 67445, 18188 | 28.9358601 |
| GOTERM_BP_ DIRECT | GO:0048484~e nteric nervous | 1 | 0.066516618 | 18935, 18205 | 28.9358601 |
| GOTERM_BP_ DIRECT | GO:0048812~n euron | 1 | 0.072325474 | 12759, 18039, 18205 | 6.75170068 |
| UP_SEQ_FEAT URE | PROPEP:Remov ed in mature | 1 | 0.073487661 | 15461, 23934, 14704, 23936, 19173 | 3.15112449 |
| GOTERM_BP_ | GO:0009636~re | 1 | 0.073720549 | 14281, 18039, 18619 | 6.67750617 |
| GOTERM_BP_ | GO:0019732~a | 1 | 0.075653176 | 21333, 109648 | 25.3188776 |
| GOTERM_MF_ | GO:0048038~q | 1 | 0.076297262 | 75406, 27973 | 25.0948276 |
| GOTERM_BP_ | GO:0042552~m | 1 | 0.076536555 | 15461, 18991, 18205 | 6.53390388 |
| GOTERM_MF_ | GO:0042277~p | 1 | 0.0791362 | 13033, 104443, 72309 | 6.40719002 |
| GOTERM_BP_ | GO:0097194~e | 1 | 0.080188094 | 13033, 12684 | 23.8295318 |
| GOTERM_BP_ | GO:0021987~c | 1 | 0.082268751 | 14281, 18039, 109648 | 6.26446455 |
| GOTERM_BP_ | GO:0051258~pr | 1 | 0.084700916 | 18039, 12373 | 22.5056689 |
| GOTERM_BP_ | GO:0003407~n | 1 | 0.084700916 | 14281, 12215 | 22.5056689 |
| GOTERM_BP_ DIRECT | GO:0006878~in tracellular | 1 | 0.084700916 | 100126824, 11927 | 22.5056689 |
| GOTERM_BP_ DIRECT | GO:0120162~p ositive | 1 | 0.088128475 | 13166, 12608, 18935 | 6.01636694 |
| UP_SEQ_FEAT | DOMAIN:Home | 1 | 0.088997881 | 18935, 16392, 14912, 15416 | 3.76808149 |
| GOTERM_BP_ DIRECT | GO:0045779~n egative | 1 | 0.089191747 | 27220, 12310 | 21.32116 |
| GOTERM_BP_ | GO:0001878~re | 1 | 0.089191747 | 21333, 109648 | 21.32116 |
| GOTERM_MF_ DIRECT | GO:0043565~s equence- | 1 | 0.089314187 | 14281, 18991, 12608, 16392, 14912, 15416 | 2.51998269 |
| GOTERM_MF_ DIRECT | GO:0030550~a cetylcholine | 1 | 0.089945484 | 23934, 23936 | 21.1324864 |
| GOTERM_MF_ DIRECT | GO:0001227~D NA-binding transcription | 1 | 0.091060519 | 18991, 12608, 14912, 15207, 15416 | 2.92651051 |
| GOTERM_BP_ | GO:0006814~s | 1 | 0.092602703 | 231290, 70261, 57780 | 5.8428179 |

| | | | | | |
|---|---|---|---|---|---|
| UP_KW_LIGAND | KW-0479~Metal-binding | 1 | 0.093681007 | 15450, 67078, 100126824, 75406, 12373, 13166, 12374, 15122, 70103, 66208, 22033, 70261, 20768, 16392, 12859, 57756, 30055, 278304, 11927, 68040, 14356, 108902, 16438, 68350 | 1.23335234 |
| UP_SEQ_FEAT | SITE:Cleavage | 1 | 0.094178547 | 13166, 53419, 241327 | 5.77367326 |
| GOTERM_BP_ | GO:0010737~pr | 1 | 0.098107864 | 14419, 14526 | 19.2905734 |
| GOTERM_BP_DIRECT | GO:0006123~mitochondrial | 1 | 0.098107864 | 12859, 12868 | 19.2905734 |
| GOTERM_BP_ | GO:0042981~re | 1 | 0.098444337 | 12759, 22033, 12684, 18205 | 3.61698251 |
