## Supplemental Table 5 for "Early Life Adversity Produces Enduring Molecular and Functional Disruption of Developing Vagal Circuits"

### Ingenuity Pathway Analysis (IPA)

| Significantly Enriched Pathway | Direction | -log(p-adjusted value) > 1.2 | Sex | Category |
|---|---|---|---|---|
| Respiratory electron transport | ↓ | 3.16 | F | Mitochondrial Dysfunction |
| Oxidative Phosphorylation | ↓ | 2.83 | | |
| Mitochondrial Dysfunction | ↑ | 1.69 | | |
| S100 Family signaling pathway | ↓ | 1.59 | | Oxidative Damage |
| NRF2-mediated oxidative stress response | ↑ | 2.66 | M | Pathways that buffer oxidative stress |
| GABAergic Receptor Signaling (Enhanced) | ↓ | 1.5 | M | Neuronal Signaling |

### DAVID Gene Ontology

| Significantly Enhanced GO Terms | FDR < .05 = significant | Sex | Category |
|---|---|---|---|
| Amidation | 0.15 x 10-3 | F | Oxidative Damage |
