## Supplemental Table 6 for "Early Life Adversity Produces Enduring Molecular and Functional Disruption of Developing Vagal Circuits"

**P9 EXCITATORY NEURONS**

| Gene Name | logFC | logCPM | LR | PValue | FDR | negLogFDR | significant | Sex |
|---|---|---|---|---|---|---|---|---|
| Gabra6 | 0.821787 | 4.364922 | 16.51036 | 4.84E-05 | 0.0223 | 1.65169671 | TRUE | Female |
| Gabrd | 0.644135 | 2.517864 | 7.160979 | 0.007451 | 0.291595 | 0.53522004 | FALSE | Female |
| Gabarapl2 | -0.34138 | 6.561853 | 3.152074 | 0.075831 | 0.650141 | 0.18699245 | FALSE | Female |
| Gabrq | -0.4623 | 1.944675 | 3.584296 | 0.058328 | 0.599058 | 0.22253143 | FALSE | Female |
| Gabre | -0.55853 | 0.206742 | 2.582528 | 0.108049 | 0.720112 | 0.14260012 | FALSE | Female |
| Gabrb2 | 0.252969 | 7.657459 | 1.808657 | 0.178669 | 0.838512 | 0.07649047 | FALSE | Female |
| Gabarap | -0.36859 | 6.662693 | 3.68778 | 0.054813 | 0.585418 | 0.23253421 | FALSE | Female |
| Gabra1 | -0.10862 | 5.908942 | 0.320165 | 0.571508 | 0.999896 | 4.50E-05 | FALSE | Female |
| Gabra5 | 0.014488 | 6.511586 | 0.005792 | 0.939337 | 0.999896 | 4.50E-05 | FALSE | Female |
| Gabrg2 | 0.076676 | 7.04742 | 0.164321 | 0.685209 | 0.999896 | 4.50E-05 | FALSE | Female |
| Gabrr2 | 0.245284 | -0.29171 | 0.399087 | 0.527561 | 0.999896 | 4.50E-05 | FALSE | Female |
| Gabrr3 | -0.28801 | -0.50563 | 0.491434 | 0.483288 | 0.999896 | 4.50E-05 | FALSE | Female |
| Gabra3 | -0.09967 | 6.595699 | 0.274361 | 0.600421 | 0.999896 | 4.50E-05 | FALSE | Female |
| Gabpa | 0.119474 | 3.819405 | 0.34637 | 0.556175 | 0.999896 | 4.50E-05 | FALSE | Female |
| Gabrg3 | -0.1694 | 9.61966 | 0.807425 | 0.368883 | 0.999896 | 4.50E-05 | FALSE | Female |
| Gabrb1 | -0.11333 | 9.318272 | 0.361942 | 0.547429 | 0.999896 | 4.50E-05 | FALSE | Female |
| Gabrr1 | 0.432302 | -0.45679 | 1.180264 | 0.277302 | 0.941238 | 0.02630035 | FALSE | Female |
| Gabrg1 | -0.15561 | 5.400045 | 0.6439 | 0.422302 | 0.999896 | 4.50E-05 | FALSE | Female |
| Gabarapl1 | -0.15031 | 6.074454 | 0.614849 | 0.432968 | 0.999896 | 4.50E-05 | FALSE | Female |
| Gabbr1 | -0.10782 | 7.144138 | 0.322536 | 0.570087 | 0.999896 | 4.50E-05 | FALSE | Female |
| Gabrb3 | 0.082394 | 9.939662 | 0.192814 | 0.660585 | 0.999896 | 4.50E-05 | FALSE | Female |
| Gabpb1 | 0.067246 | 5.106703 | 0.119931 | 0.729109 | 0.999896 | 4.50E-05 | FALSE | Female |
| Gabpb2 | 0.066623 | 4.300986 | 0.111477 | 0.738468 | 0.999896 | 4.50E-05 | FALSE | Female |
| Gabra2 | 0.044196 | 7.814242 | 0.054951 | 0.814661 | 0.999896 | 4.50E-05 | FALSE | Female |
| Gabbr2 | 0.033431 | 8.200956 | 0.031501 | 0.859127 | 0.999896 | 4.50E-05 | FALSE | Female |
| Gabra4 | 0.003541 | 4.925723 | 0.000327 | 0.985567 | 0.999896 | 4.50E-05 | FALSE | Female |

| Gene Name | logFC | logCPM | LR | PValue | FDR | negLogFDR | significant | Sex |
|---|---|---|---|---|---|---|---|---|
| Gabra6 | -1.32526 | 4.364922 | 42.71936 | 6.32E-11 | 4.19E-08 | 7.37727264 | TRUE | Males |
| Gabrd | -1.0255 | 2.517864 | 16.97223 | 3.79E-05 | 0.005073 | 2.29474504 | TRUE | Males |
| Gabarapl2 | 0.433256 | 6.561853 | 6.047876 | 0.013923 | 0.310347 | 0.50815272 | FALSE | Males |
| Gabrq | -0.40589 | 1.944675 | 3.438334 | 0.0637 | 0.63637 | 0.19629051 | FALSE | Males |
| Gabre | -0.51943 | 0.206742 | 2.933676 | 0.08675 | 0.712943 | 0.14694499 | FALSE | Males |
| Gabrb2 | -0.29505 | 7.657459 | 2.840717 | 0.091903 | 0.726087 | 0.1390115 | FALSE | Males |
| Gabarap | 0.267856 | 6.662693 | 2.321157 | 0.127625 | 0.811406 | 0.09076191 | FALSE | Males |
| Gabra1 | -0.26514 | 5.908942 | 2.232728 | 0.135115 | 0.825119 | 0.08348337 | FALSE | Males |
| Gabra5 | 0.219901 | 6.511586 | 1.561172 | 0.211494 | 0.93732 | 0.02811192 | FALSE | Males |
| Gabrg2 | -0.21715 | 7.04742 | 1.5315 | 0.215887 | 0.942322 | 0.02580083 | FALSE | Males |
| Gabrr2 | -0.46248 | -0.29171 | 1.516771 | 0.218109 | 0.944697 | 0.02470725 | FALSE | Males |
| Gabrr3 | -0.41468 | -0.50563 | 1.281371 | 0.257644 | 0.979086 | 0.00917925 | FALSE | Males |
| Gabra3 | -0.19831 | 6.595699 | 1.271757 | 0.259437 | 0.980472 | 0.00856492 | FALSE | Males |
| Gabpa | -0.19782 | 3.819405 | 1.096989 | 0.294928 | 0.999937 | 2.75E-05 | FALSE | Males |

| | | | | | | | | |
|---|---|---|---|---|---|---|---|---|
| Gabrg3 | -0.14724 | 9.61966 | 0.716016 | 0.397454 | 0.999937 | 2.75E-05 | FALSE | Males |
| Gabrb1 | -0.12879 | 9.318272 | 0.547458 | 0.459358 | 0.999937 | 2.75E-05 | FALSE | Males |
