## Supplemental Table 7 for "Early Life Adversity Produces Enduring Molecular and Functional Disruption of Developing Vagal Circuits"

### Ingenuity Pathway Analysis (IPA)

| Significantly Enriched Pathway | Direction | -log(p-adjusted value) > 1.2 | Sex | Category |
|---|---|---|---|---|
| Mitochondrial Dysfunction | ↑ | 1.5 | F | Mitochondrial Dysfunction |
| GABAergic Receptor Signaling | ↓ | 2.24 | F | Neuronal Signaling |

### DAVID Gene Ontology

| Significantly Enhanced GO Terms | FDR < .05 = significant | Sex | Category |
|---|---|---|---|
| Amidation | 1.38 x 10 -5 | F | Oxidative Damage |
| Hemoglobin complex | 2.74 x 10 -5 | M | Pathways that buffer |
| Haptoglobin-hemoglobin complex | 2.74 x 10 -5 | | |
| Hemoglobin oxygen transport | 4.23 x 10 -5 | | |
| Oxygen Carrier Activity | 6.93 x 10 -5 | | |
| Oxygen Binding | 0.02 x 10 -2 | | |
| Cellular Oxidant Detoxification | 0.05 x 10 -2 | | |
| Peroxidase Activity | 0.05 x 10 -2 | | |
