## Supplemental Table 9 for "Early Life Adversity Produces Enduring Molecular and Functional Disruption of Developing Vagal Circuits"

© 2000-2025 QIAGEN. All rights reserved.

**P120 EXCITATORY NEURONS FEMALE**

| Ingenuity Canonical Pathways | -log(p-value) | Ratio | z-score | Molecules |
|---|---|---|---|---|
| Synthesis, secretion, and deacylation of Ghrelin | 5.01 | 0.2 | -1 | GCG,GH1,SPCS3,UCN |
| ROBO SLIT Signaling Pathway | 4.5 | 0.0547 | 0.816 | ARF5,ARF6,CTNND1,FOS,Myl12a,MYL9,PAX6 |
| Formation of the anterior neural plate | 4.29 | 0.273 | #NUM! | OTX2,PAX6,ZIC2 |
| Class B/2 (Secretin family receptors) | 4.28 | 0.0632 | 0 | CALCA,CRH,FZD7,GCG,GNB2,UCN |
| Amyloid fiber formation | 3.43 | 0.0568 | -1.342 | CALB1,CALCA,PRL,TTR,UBA52 |
| RNA Polymerase II Transcription | 3.2 | 0.04 | -2.449 | ALYREF,CSTF2T,POLR2K,POLR2L,SNRPF,TAF7L |
| Incretin synthesis, secretion, and inactivation | 3.17 | 0.12 | #NUM! | GCG,PAX6,SPCS3 |

| | | | | |
|---|---|---|---|---|
| Processing of Capped Intron-Containing Pre-mRNA | 3.03 | 0.0278 | -2.828 | ALYREF,CSTF2T,LSM5,PCBP1,PCBP2,POLR2K,POLR2L,SNRPF |
| Regulation of CDH19 Expression and Function | 3.01 | 0.286 | #NUM! | CTNND1,JUP |
| Nucleotide Excision Repair | 3.01 | 0.0459 | -2.236 | POLR2K,POLR2L,RPA3,SUMO2,UBA52 |
| Transcriptional Regulatory Network in Embryonic Stem Cells | 3 | 0.0366 | 0.816 | CALB1,EOMES,FOXD3,FZD7,GJD2,PAX6 |
| Role of JAK2 in Hormone-like Cytokine Signaling | 2.97 | 0.0615 | #NUM! | FOS,GH1,NPY,PRL |
| Signaling by FGFR2 | 2.78 | 0.0548 | -2 | FRS3,POLR2K,POLR2L,UBA52 |
| Senescence-Associated Secretory Phenotype (SASP) | 2.76 | 0.0541 | -2 | CDK6,CEBPB,FOS,UBA52 |
| VDR/RXR Activation | 2.67 | 0.0513 | #NUM! | CALB1,CASR,CEBPB,GADD45A |

| | | | | |
|---|---|---|---|---|
| Nucleotide Excision Repair Pathway | 2.67 | 0.0811 | #NUM! | POLR2K,POLR2L,RPA3 |
| Hereditary Breast Cancer Signaling | 2.56 | 0.0362 | 1 | CDK6,GADD45A,POLR2K,POLR2L,UBA52 |
| Activation of anterior HOX genes in hindbrain during early embryogenesis | 2.56 | 0.0476 | -2 | HOXD1,PAX6,POLR2K,POLR2L |
| Cell junction organization | 2.41 | 0.0435 | -1 | CDH3,CLDN2,CTNND1,JUP |
| Prolactin Signaling | 2.35 | 0.0417 | #NUM! | CEBPB,FOS,KCNMB1,PRL |
| Response of EIF2AK1 (HRI) to heme deficiency | 2.33 | 0.133 | #NUM! | CEBPB,EIF2S3 |
| DNA Methylation and Transcriptional Repression Signaling | 2.32 | 0.0408 | #NUM! | CDK6,CEBPB,GADD45A,GLI1 |
| Assembly of RNA Polymerase II Complex | 2.27 | 0.0588 | #NUM! | POLR2K,POLR2L,TAF7L |

| | | | | |
|---|---|---|---|---|
| B-WICH complex positively regulates rRNA expression | 2.25 | 0.0577 | #NUM! | POLR2K,POLR2L,TAF1D |
| Prolactin receptor signaling | 2.22 | 0.118 | #NUM! | GH1,PRL |
| G alpha (q) signalling events | 2.13 | 0.0287 | 0.447 | CASR,GCG,GNB2,TAC3,TRH |
| Class A/1 (Rhodopsin-like receptors) | 2.07 | 0.0211 | 1.134 | GAL,GPR55,NPY,PRLH,S1PR1,TAC3,TRH |
| Tight Junction Signaling | 2.06 | 0.0276 | #NUM! | CLDN2,CNKSR3,CSTF2T,FOS,MYL9 |
| Ion channel transport | 2.04 | 0.0273 | -1.342 | ANO2,ATP6V1B1,CASQ2,SLN,UBA52 |
| G alpha (i) signalling events | 2.03 | 0.0233 | 0.816 | CASR,GAL,GNB2,GPR55,NPY,PCP2 |
| Cytosolic sensors of pathogen-associated DNA | 2 | 0.0469 | #NUM! | POLR2K,POLR2L,UBA52 |
| NoRC negatively regulates rRNA expression | 1.95 | 0.0448 | #NUM! | POLR2K,POLR2L,TAF1D |

| | | | | |
|---|---|---|---|---|
| TP53 Regulates Transcription of DNA Repair Genes | 1.9 | 0.0429 | #NUM! | FOS,POLR2K,POLR2L |
| Signaling by NTRK2 (TRKB) | 1.89 | 0.08 | #NUM! | FRS3,NTF3 |
| Growth hormone receptor signaling | 1.89 | 0.08 | #NUM! | GH1,PRL |
| Insulin Secretion Signaling Pathway | 1.88 | 0.0215 | -1.342 | EIF2S3,GCG,GH1,NEUROD1,PRL,SPCS3 |
| Chronic Myeloid Leukemia Signaling | 1.86 | 0.0214 | -0.816 | CDK6,FOS,FZD7,GLI1,PCBP2,RCC1 |
| GABA Receptor Signaling | 1.86 | 0.0301 | #NUM! | GABRA6,GABRD,GNB2,UBA52 |
| Growth Hormone Signaling | 1.85 | 0.0411 | #NUM! | FOS,GH1,PRL |
| RNA Polymerase I Transcription | 1.83 | 0.0405 | #NUM! | POLR2K,POLR2L,TAF1D |
| Cardiogenesis | 1.83 | 0.0741 | #NUM! | EOMES,TBX20 |
| Integrin Signaling | 1.81 | 0.0238 | 1 | ARF5,ARF6,ITGAM,Myl12a,MYL9 |

| | | | | |
|---|---|---|---|---|
| GPCR-Mediated Integration of Enteroendocrine Signaling Exemplified by an L Cell | 1.8 | 0.0395 | #NUM! | CALCA,GAL, GCG |
| Molecular Mechanisms of Cancer | 1.77 | 0.0139 | -1.732 | CASR,CDK6, CTNND1,FOS,FZD7,GLI1, GNB2,GPR55,GPR88,GPRC5C,ITGAM, S1PR1 |
| Processing of Capped Intronless Pre-mRNA | 1.77 | 0.069 | #NUM! | CSTF2T,SNRPF |
| Bone Mineralization Signaling Pathway | 1.74 | 0.0278 | -2 | CALB1,CASR,FZD7,PRL |
| mRNA Capping | 1.74 | 0.0667 | #NUM! | POLR2K,POLR2L |
| Regulation of CDH11 Expression and Function | 1.74 | 0.0667 | #NUM! | CTNND1,JUP |
| Glutathione Redox Reactions I | 1.74 | 0.0667 | #NUM! | Gstp1 (includes others),MGST3 |
| G alpha (s) signalling events | 1.73 | 0.0276 | 0 | CALCA,CRH, GCG,GNB2 |
| HER-2 Signaling in Breast Cancer | 1.68 | 0.022 | 1.342 | ARF5,ARF6, COX7C,Cox8b,FOS |

| | | | | |
|---|---|---|---|---|
| Corticotropin Releasing Hormone Signaling | 1.64 | 0.0258 | 0 | CRH,FOS,GLI1,UCN |
| Cargo concentration in the ER | 1.64 | 0.0588 | #NUM! | CNIH3,FOLR1 |
| Oxidative Stress Induced Senescence | 1.63 | 0.0341 | #NUM! | CDK6,FOS,UBA52 |
| Telomere Maintenance | 1.62 | 0.0337 | #NUM! | POLR2K,POLR2L,RPA3 |
| Oncogene Induced Senescence | 1.61 | 0.0571 | #NUM! | CDK6,UBA52 |
| Osteoarthritis Pathway | 1.6 | 0.021 | #NUM! | CASR,CEBPB,FZD7,GLI1,ITGAM |
| NER (Nucleotide Excision Repair, Enhanced Pathway) | 1.6 | 0.033 | #NUM! | POLR2K,POLR2L,RPA3 |
| Preeclampsia Signaling Pathway | 1.59 | 0.0248 | 0 | FOS,FZD7,KCNMB1,MYL9 |
| Regulation of TP53 Activity through Phosphorylation | 1.57 | 0.0323 | #NUM! | RPA3,TAF7L,UBA52 |
| IL-12 Signaling and Production in Macrophages | 1.57 | 0.0207 | 0 | CEBPB,FOS,ITGAM,PRL,ZIC2 |

| | | | | |
|---|---|---|---|---|
| G-Protein Coupled Receptor Signaling | 1.57 | 0.014 | -1.265 | CASR,FOS,FZD7,GNB2,GPR55,GPR88,GPRC5C,Htr5b,MYL9,S1PR1 |
| CREB Signaling in Neurons | 1.55 | 0.0146 | -0.816 | CASR,FZD7,GNB2,GPR55,GPR88,GPRC5C,POLR2K,POLR2L,S1PR1 |
| Fanconi Anemia Pathway | 1.55 | 0.0526 | #NUM! | RPA3,UBA52 |
| Mitophagy | 1.51 | 0.05 | #NUM! | TOMM7,UBA52 |
| Signaling by FGFR3 | 1.51 | 0.05 | #NUM! | FRS3,UBA52 |
| WNT/β-catenin Signaling | 1.49 | 0.0231 | #NUM! | CDH3,FZD7,SFRP5,UBA52 |
| Signaling by FGFR4 | 1.49 | 0.0488 | #NUM! | FRS3,UBA52 |
| RNA Polymerase III Transcription | 1.49 | 0.0488 | #NUM! | POLR2K,POLR2L |
| Sumoylation Pathway | 1.48 | 0.0297 | #NUM! | FOS,RCC1,SUMO2 |
| Androgen Signaling | 1.47 | 0.0229 | #NUM! | GNB2,POLR2K,POLR2L,TAF7L |
| mRNA 3 Prime End Processing Signaling Pathway | 1.47 | 0.0294 | #NUM! | CSTF2T,PCBP1,PCBP2 |

| | | | | |
|---|---|---|---|---|
| Transmission across Electrical Synapses | 1.47 | 0.2 | #NUM! | GJD2 |
| Regulation of beta-cell development | 1.47 | 0.0476 | #NUM! | NEUROD1,PAX6 |
| Potassium Channels | 1.46 | 0.0291 | #NUM! | GNB2,KCNG1,KCNMB1 |
| Response of EIF2AK4 (GCN2) to amino acid deficiency | 1.46 | 0.0291 | #NUM! | CEBPB,EIF2S3,UBA52 |
| Vasopressin regulates renal water homeostasis via Aquaporins | 1.45 | 0.0465 | #NUM! | AQP1,GNB2 |
| Signaling by Rho Family GTPases | 1.41 | 0.0187 | #NUM! | CDH3,FOS,GNB2,ITGAM,MYL9 |
| Aryl Hydrocarbon Receptor Signaling | 1.41 | 0.0219 | #NUM! | CDK6,FOS,Gstp1 (includes others),MGST3 |
| Interleukin-4 and Interleukin-13 signaling | 1.38 | 0.027 | #NUM! | FOS,ITGAM,S1PR1 |
| DNA Damage Bypass | 1.36 | 0.0417 | #NUM! | RPA3,UBA52 |

| | | | | |
|---|---|---|---|---|
| HDR through Homologous Recombination (HRR) or Single Strand Annealing (SSA) | 1.35 | 0.0263 | #NUM! | RPA3,SUMO2,UBA52 |
| iNOS Signaling | 1.34 | 0.0408 | #NUM! | FOS,HMGA1 |
| Erythrocytes take up oxygen and release carbon dioxide | 1.32 | 0.143 | #NUM! | AQP1 |
| Epithelial-Mesenchymal Transition (EMT) during gastrulation | 1.32 | 0.143 | #NUM! | EOMES |
| Cerebral Malformation Signaling Pathway | 1.32 | 0.0256 | #NUM! | JUP,Myl12a,MYL9 |
| Transcriptional and post-translational regulation of MITF-M expression and activity | 1.31 | 0.0392 | #NUM! | FOXD3,KIT |
| Glutathione-mediated Detoxification | 1.31 | 0.0392 | #NUM! | Gstp1 (includes others),MGST3 |

| | | | | |
|---|---|---|---|---|
| ESR-mediated signaling | 1.31 | 0.0254 | #NUM! | FOS,POLR2K,POLR2L |
| Huntington's Disease Signaling | 1.3 | 0.0174 | #NUM! | GNB2,NEUROD1,POLR2K,POLR2L,UBA52 |
| Signaling by FGFR1 | 1.3 | 0.0385 | #NUM! | FRS3,UBA52 |
| IL-15 Production | 1.28 | 0.0246 | #NUM! | KIT,ROS1,STYK1 |
| Beta-catenin independent WNT signaling | 1.26 | 0.0195 | #NUM! | FZD7,GNB2,STK17B,UBA52 |
| BBSome Signaling Pathway | 1.25 | 0.0142 | -0.378 | CASR,FZD7,GLI1,GPR55,GPR88,GPRC5C,S1PR1 |
| IL-8 Signaling | 1.25 | 0.0192 | #NUM! | FOS,GNB2,ITGAM,MYL9 |
| Phagosome Formation | 1.24 | 0.0127 | -1 | ARF6,CASR,FZD7,GPR55,GPR88,GPRC5C,ITGAM,MYL9,S1PR1 |
| Gustation Pathway | 1.23 | 0.019 | 0 | CASR,GABRA6,GABRD,GCG |
| Cell Cycle Control of Chromosomal Replication | 1.23 | 0.0351 | #NUM! | CDK6,RPA3 |
| Activin Inhibin Signaling Pathway | 1.22 | 0.0189 | #NUM! | CEBPB,FOS,PAX6,PRL |

| | | | | |
|---|---|---|---|---|
| Breast Cancer Regulation by Stathmin1 | 1.22 | 0.0132 | -1.134 | CASR,CDK6,FZD7,GNB2,GPR55,GPR88,GPRC5C,S1PR1 |
| Cellular response to mitochondrial stress | 1.22 | 0.111 | #NUM! | EIF2S3 |
| Malate-aspartate shuttle | 1.22 | 0.111 | #NUM! | SLC25A22 |
| RUNX1 regulates megakaryocyte differentiation and platelet function | 1.21 | 0.0345 | #NUM! | MYL9,NR4A3 |
| TR/RXR Activation | 1.2 | 0.0229 | #NUM! | GH1,NRGN,TRH |
| Iron uptake and transport | 1.19 | 0.0333 | #NUM! | ATP6V1B1,UBA52 |
| GABA receptor activation | 1.19 | 0.0333 | #NUM! | GABRA6,GNB2 |
| RHOGDI Signaling | 1.17 | 0.0182 | #NUM! | CDH3,GNB2,ITGAM,MYL9 |
| Neurotransmitter clearance | 1.17 | 0.1 | #NUM! | SLC6A4 |
| POU5F1 (OCT4), SOX2, NANOG repress genes related to differentiation | 1.17 | 0.1 | #NUM! | EOMES |

| | | | | |
|---|---|---|---|---|
| RUNX1 and FOXP3 control the development of regulatory T lymphocytes (Tregs) | 1.17 | 0.1 | #NUM! | TNFRSF18 |
| Formation of the posterior neural plate | 1.17 | 0.1 | #NUM! | OTX2 |
| Axonal Guidance Signaling | 1.16 | 0.0135 | #NUM! | ADAM18,FZD7,GLI1,GNB2,ITGAM,MYL9,NTF3 |
| Hepatic Fibrosis Signaling Pathway | 1.16 | 0.0144 | 0 | CEBPB,FOS,FZD7,GLI1,ITGAM,MYL9 |
| Lung Ionic Balance Signaling Pathway | 1.15 | 0.0134 | -0.378 | CASR,FZD7,GPR55,GPR88,GPRC5C,KCNMB1,S1PR1 |
| GABAergic Receptor Signaling Pathway (Enhanced) | 1.13 | 0.0214 | #NUM! | GABRA6,GABRD,GNB2 |
| Ephrin A Signaling | 1.13 | 0.0214 | #NUM! | ARF6,CHN2,MYL9 |
| Erythrocytes take up carbon dioxide and release oxygen | 1.13 | 0.0909 | #NUM! | AQP1 |

| | | | | |
|---|---|---|---|---|
| Processing and activation of SUMO | 1.13 | 0.0909 | #NUM! | SUMO2 |
| Metallothioneins bind metals | 1.13 | 0.0909 | #NUM! | MT2A |
| Alpha-protein kinase 1 signaling pathway | 1.13 | 0.0909 | #NUM! | UBA52 |
| Pexophagy | 1.13 | 0.0909 | #NUM! | UBA52 |
| Myelination Signaling Pathway | 1.11 | 0.0152 | -1.342 | ARF6,FOS,FZD7,GLI1,NTF3 |
| Hedgehog 'on' state | 1.11 | 0.0208 | #NUM! | GLI1,STK17B,UBA52 |
| Ribavirin ADME | 1.1 | 0.0299 | #NUM! | SLC28A3,STK17B |
| Iron homeostasis signaling pathway | 1.1 | 0.0207 | #NUM! | ATP6V1B1,HBA2,PCBP1 |
| Neuroinflammation Signaling Pathway | 1.1 | 0.0152 | #NUM! | CALB1,FOS,GABRA6,GABRD,NTF3 |
| Vitamin D (calciferol) metabolism | 1.1 | 0.0833 | #NUM! | SUMO2 |
| Specification of primordial germ cells | 1.1 | 0.0833 | #NUM! | EOMES |
| Assembly of RNA Polymerase I Complex | 1.1 | 0.0833 | #NUM! | TAF1D |

| | | | | |
|---|---|---|---|---|
| Sertoli Cell-Germ Cell Junction Signaling Pathway (Enhanced) | 1.09 | 0.0169 | 1 | CLDN2,CTNND1,FOS,JUP |
| Sensory processing of sound by inner hair cells of the cochlea | 1.08 | 0.029 | #NUM! | KCNMB1,OTOF |
| Role of Macrophages, Fibroblasts and Endothelial Cells in Rheumatoid Arthritis | 1.07 | 0.0148 | -1.342 | CEBPB,FOS,FZD7,IL16,SFRP5 |
| POU5F1 (OCT4), SOX2, NANOG activate genes related to proliferation | 1.06 | 0.0769 | #NUM! | FOXD3 |
| Passive transport by Aquaporins | 1.06 | 0.0769 | #NUM! | AQP1 |
| SUMOylation of immune response proteins | 1.06 | 0.0769 | #NUM! | SUMO2 |
| Cleavage and Polyadenylation of Pre-mRNA | 1.06 | 0.0769 | #NUM! | CSTF2T |

| | | | | |
|---|---|---|---|---|
| Basal Cell Carcinoma Signaling | 1.06 | 0.0282 | #NUM! | FZD7,GLI1 |
| Remodeling of Epithelial Adherens Junctions | 1.06 | 0.0282 | #NUM! | ARF6,CTNND1 |
| NAD Signaling Pathway | 1.06 | 0.0199 | #NUM! | CEBPB,POLR2K,POLR2L |
| WNT/SHH Axonal Guidance Signaling Pathway | 1.06 | 0.0199 | #NUM! | ARF6,FZD7,GLI1 |
| Tuberculosis Active Signaling Pathway | 1.04 | 0.0163 | 0 | ATP6V1B1,FOS,GADD45A,ITGAM |
| COPII-mediated vesicle transport | 1.04 | 0.0274 | #NUM! | CNIH3,FOLR1 |
| Mitochondrial Dysfunction | 1.04 | 0.0145 | 2 | COX7C,Cox8b,Gstp1 (includes others),MGST3,TOMM7 |
| Regulation of TP53 Activity through Association with Co-factors | 1.03 | 0.0714 | #NUM! | POU4F1 |
| Extra-nuclear estrogen signaling | 1.02 | 0.0267 | #NUM! | FOS,GNB2 |
| Toll-like Receptor Signaling | 1 | 0.026 | #NUM! | FOS,UBA52 |
| PKR-mediated signaling | 0.993 | 0.0256 | #NUM! | EIF2S3,SUMO2 |

| | | | | |
|---|---|---|---|---|
| Renal Cell Carcinoma Signaling | 0.993 | 0.0256 | #NUM! | FOS,UBA52 |
| Thyroid Cancer Signaling | 0.993 | 0.0256 | #NUM! | FOS,NTF3 |
| Neurotrophin/TRK Signaling | 0.993 | 0.0256 | #NUM! | FOS,NTF3 |
| Sperm Motility | 0.985 | 0.0156 | #NUM! | GNB2,KIT,ROS1,STYK1 |
| Signaling by MET | 0.983 | 0.0253 | #NUM! | ARF6,UBA52 |
| Cellular Effects of Sildenafil (Viagra) | 0.983 | 0.0117 | 0.707 | CASR,FZD7,GNB2,GPR55,GPR88,GPRC5C,MYL9,S1PR1 |
| Mismatch Repair | 0.978 | 0.0625 | #NUM! | RPA3 |
| Leukotriene Biosynthesis | 0.978 | 0.0625 | #NUM! | MGST3 |
| IL-17A Signaling in Fibroblasts | 0.974 | 0.025 | #NUM! | CEBPB,FOS |
| DDX58/IFIH1-mediated induction of interferon-alpha/beta | 0.965 | 0.0247 | #NUM! | PCBP2,UBA52 |
| RAF/MAP kinase cascade | 0.962 | 0.0153 | #NUM! | DUSP5,FRS3,KIT,UBA52 |
| CXCR4 Signaling | 0.954 | 0.0179 | #NUM! | FOS,GNB2,MYL9 |

| | | | | |
|---|---|---|---|---|
| FOXO-mediated transcription of cell cycle genes | 0.953 | 0.0588 | #NUM! | GADD45A |
| Germ layer formation at gastrulation | 0.953 | 0.0588 | #NUM! | EOMES |
| RAN Signaling | 0.953 | 0.0588 | #NUM! | RCC1 |
| RNA polymerase II transcribes snRNA genes | 0.948 | 0.0241 | #NUM! | POLR2K,POLR2L |
| JAK/STAT Signaling | 0.948 | 0.0241 | #NUM! | CEBPB,FOS |
| Cachexia Signaling Pathway | 0.947 | 0.0136 | -1.342 | CEBPB,CRH,EIF2S3,NPY,S1PR1 |
| Formation of definitive endoderm | 0.93 | 0.0556 | #NUM! | EOMES |
| NRF2-mediated Oxidative Stress Response | 0.928 | 0.0148 | #NUM! | Cyp2j11/Cyp2j8,FOS,Gstp1 (includes others),MGST3 |
| Platelet homeostasis | 0.923 | 0.0233 | #NUM! | GNB2,KCNMB1 |
| Regulation of the Epithelial Mesenchymal Transition in Development Pathway | 0.923 | 0.0233 | #NUM! | FZD7,GLI1 |

| | | | | |
|---|---|---|---|---|
| Ribonucleotide Reductase Signaling Pathway | 0.92 | 0.0172 | #NUM! | CDH3,CDK6, FOS |
| Unfolded Protein Response (UPR) | 0.908 | 0.0526 | #NUM! | EIF2S3 |
| RHO GTPases activate CIT | 0.908 | 0.0526 | #NUM! | MYL9 |
| RHO GTPases Activate ROCKs | 0.908 | 0.0526 | #NUM! | MYL9 |
| Regulation of TP53 Activity through Methylation | 0.908 | 0.0526 | #NUM! | UBA52 |
| ABRA Signaling Pathway | 0.906 | 0.0227 | #NUM! | FOS,MYL9 |
| Regulation of Cellular Mechanics by Calpain Protease | 0.898 | 0.0225 | #NUM! | CDK6,ITGAM |
| BMP signaling pathway | 0.89 | 0.0222 | #NUM! | CHRD,SOST DC1 |
| SUMOylation of transcription factors | 0.887 | 0.05 | #NUM! | SUMO2 |
| Ceramide Signaling | 0.883 | 0.022 | #NUM! | FOS,S1PR1 |
| Acute Myeloid Leukemia Signaling | 0.875 | 0.0217 | #NUM! | JUP,KIT |
| RHO GTPases activate PAKs | 0.867 | 0.0476 | #NUM! | MYL9 |

| | | | | |
|---|---|---|---|---|
| Apelin Adipocyte Signaling Pathway | 0.86 | 0.0213 | #NUM! | Gstp1 (includes others),MGST3 |
| Insertion of tail-anchored proteins into the endoplasmic reticulum membrane | 0.849 | 0.0455 | #NUM! | OTOF |
| Chaperone Mediated Autophagy | 0.849 | 0.0455 | #NUM! | UBA52 |
| Acute Phase Response Signaling | 0.834 | 0.0157 | #NUM! | CEBPB,FOS,TTR |
| IL-1 Signaling | 0.831 | 0.0204 | #NUM! | FOS,GNB2 |
| RAF-independent MAPK1/3 activation | 0.831 | 0.0435 | #NUM! | DUSP5 |
| Azathioprine ADME | 0.831 | 0.0435 | #NUM! | SLC28A3 |
| Protein folding | 0.824 | 0.0202 | #NUM! | GNB2,PFDN4 |
| Leukocyte Extravasation Signaling | 0.82 | 0.0155 | #NUM! | CLDN2,CTNND1,ITGAM |
| Other interleukin signaling | 0.814 | 0.0417 | #NUM! | IL16 |
| Regulation of RUNX1 Expression and Activity | 0.814 | 0.0417 | #NUM! | CDK6 |

| | | | | |
|---|---|---|---|---|
| RAS processing | 0.814 | 0.0417 | #NUM! | UBA52 |
| Gene Silencing by RNA | 0.804 | 0.0196 | #NUM! | POLR2K,POLR2L |
| COPI-mediated anterograde transport | 0.804 | 0.0196 | #NUM! | ARF5,FOLR1 |
| MyD88 cascade initiated on plasma membrane | 0.798 | 0.04 | #NUM! | UBA52 |
| Glycogen metabolism | 0.798 | 0.04 | #NUM! | UBA52 |
| ABC-family proteins mediated transport | 0.797 | 0.0194 | #NUM! | EIF2S3,UBA52 |
| GPER1 signaling | 0.797 | 0.0194 | #NUM! | GNB2,STK17B |
| Mouse Embryonic Stem Cell Pluripotency | 0.797 | 0.0194 | #NUM! | FOXD3,FZD7 |
| Human Embryonic Stem Cell Pluripotency | 0.792 | 0.015 | #NUM! | FOXD3,FZD7,NTF3 |
| IL-17A Signaling in Gastric Cells | 0.782 | 0.0385 | #NUM! | FOS |
| Paxillin Signaling | 0.778 | 0.0189 | #NUM! | ARF6,ITGAM |
| Extracellular matrix organization | 0.771 | 0.0187 | #NUM! | MATN4,TTR |
| Signaling by VEGF | 0.771 | 0.0187 | #NUM! | CTNND1,JUP |

| | | | | |
|---|---|---|---|---|
| ATF4 activates genes in response to endoplasmic reticulum stress | 0.767 | 0.037 | #NUM! | CEBPB |
| Integration of energy metabolism | 0.765 | 0.0185 | #NUM! | GCG,GNB2 |
| Oxidative Phosphorylation | 0.765 | 0.0185 | #NUM! | COX7C,Cox8b |
| Xenobiotic Metabolism AHR Signaling Pathway | 0.759 | 0.0183 | #NUM! | Gstp1 (includes others),MGST3 |
| S100 Family Signaling Pathway | 0.755 | 0.0102 | -1.414 | CASR,FOS,FZD7,GPR55,GPR88,GPRC5C,NTF3,S1PR1 |
| RHO GTPases activate PKNs | 0.753 | 0.0357 | #NUM! | MYL9 |
| Regulation of Actin-based Motility by Rho | 0.747 | 0.018 | #NUM! | ITGAM,MYL9 |
| SUMOylation of intracellular receptors | 0.739 | 0.0345 | #NUM! | SUMO2 |
| SIRT1 negatively regulates rRNA expression | 0.739 | 0.0345 | #NUM! | TAF1D |

| | | | | |
|---|---|---|---|---|
| TNFs bind their physiological receptors | 0.739 | 0.0345 | #NUM! | TNFRSF18 |
| Hedgehog 'off' state | 0.729 | 0.0175 | #NUM! | GLI1,UBA52 |
| Activation of kainate receptors upon glutamate binding | 0.726 | 0.0333 | #NUM! | GNB2 |
| Protein ubiquitination | 0.726 | 0.0333 | #NUM! | UBA52 |
| Signaling by NTRK3 (TRKC) | 0.726 | 0.0333 | #NUM! | NTF3 |
| FOXO-mediated transcription of oxidative stress, metabolic and neuronal genes | 0.726 | 0.0333 | #NUM! | NPY |
| Signaling by CSF3 (G-CSF) | 0.726 | 0.0333 | #NUM! | UBA52 |
| Sleep NREM Signaling Pathway | 0.718 | 0.0172 | #NUM! | GABRA6,GABRD |
| Calcium Signaling | 0.716 | 0.0138 | #NUM! | CASQ2,MYL9,TNNT2 |
| Endosomal Sorting Complex Required For Transport (ESCRT) | 0.713 | 0.0323 | #NUM! | UBA52 |

| | | | | |
|---|---|---|---|---|
| Signaling by CSF1 (M-CSF) in myeloid cells | 0.713 | 0.0323 | #NUM! | UBA52 |
| MyD88 dependent cascade initiated on endosome | 0.713 | 0.0323 | #NUM! | UBA52 |
| Sonic Hedgehog Signaling | 0.713 | 0.0323 | #NUM! | GLI1 |
| Nonsense-Mediated Decay (NMD) | 0.712 | 0.0171 | #NUM! | SMG8,UBA52 |
| Cholecystokinin/Gastrin-mediated Signaling | 0.712 | 0.0171 | #NUM! | FOS,GH1 |
| PAK Signaling | 0.706 | 0.0169 | #NUM! | ITGAM,MYL9 |
| Nuclear Cytoskeleton Signaling Pathway | 0.705 | 0.0136 | #NUM! | CDH3,ITGAM,ZIC2 |
| GPCR-Mediated Nutrient Sensing in Enteroendocrine Cells | 0.701 | 0.0168 | #NUM! | CASR,GCG |
| Toll-like Receptor Cascades | 0.701 | 0.0312 | #NUM! | ITGAM |
| G-protein beta:gamma signalling | 0.701 | 0.0312 | #NUM! | GNB2 |

| | | | | |
|---|---|---|---|---|
| Thrombin signalling through proteinase activated receptors (PARs) | 0.701 | 0.0312 | #NUM! | GNB2 |
| RIPK1-mediated regulated necrosis | 0.701 | 0.0312 | #NUM! | UBA52 |
| MSP-RON Signaling in Macrophages Pathway | 0.695 | 0.0167 | #NUM! | FOS,ITGAM |
| Glycation Signaling Pathway | 0.689 | 0.0133 | #NUM! | FOS,HBA2,TTR |
| Toll Like Receptor 3 (TLR3) Cascade | 0.689 | 0.0303 | #NUM! | UBA52 |
| Signal amplification | 0.689 | 0.0303 | #NUM! | GNB2 |
| MAPK targets/ Nuclear events mediated by MAP kinases | 0.689 | 0.0303 | #NUM! | FOS |
| Sialic acid metabolism | 0.689 | 0.0303 | #NUM! | NEU4 |
| TNFR2 Signaling | 0.689 | 0.0303 | #NUM! | FOS |
| Synthesis of DNA | 0.685 | 0.0164 | #NUM! | RPA3,UBA52 |

| | | | | |
|---|---|---|---|---|
| Role of NANOG in Mammalian Embryonic Stem Cell Pluripotency | 0.679 | 0.0163 | #NUM! | FOXD3,FZD7 |
| Signaling by NOTCH2 | 0.677 | 0.0294 | #NUM! | UBA52 |
| Late endosomal microautophagy | 0.677 | 0.0294 | #NUM! | UBA52 |
| Role of Osteoblasts, Osteoclasts and Chondrocytes in Rheumatoid Arthritis | 0.674 | 0.0131 | #NUM! | FOS,FZD7,SFRP5 |
| SUMOylation of SUMOylation proteins | 0.666 | 0.0286 | #NUM! | SUMO2 |
| Activation of the pre-replicative complex | 0.666 | 0.0286 | #NUM! | RPA3 |
| Transcriptional Regulation by NPAS4 | 0.666 | 0.0286 | #NUM! | FOS |
| Neurovascular Coupling Signaling Pathway | 0.663 | 0.0129 | #NUM! | GABRA6,GABRD,KCNMB1 |
| Gαs Signaling | 0.659 | 0.0157 | #NUM! | GNB2,Htr5b |

| | | | | |
|---|---|---|---|---|
| Striated Muscle Contraction | 0.655 | 0.0278 | #NUM! | TNNT2 |
| Transcriptional regulation by the AP-2 (TFAP2) family of transcription factors | 0.655 | 0.0278 | #NUM! | KIT |
| Endocannabinoid Developing Neuron Pathway | 0.654 | 0.0156 | #NUM! | GNB2,PAX6 |
| Clathrin-mediated endocytosis | 0.649 | 0.0155 | #NUM! | ARF6,UBA52 |
| IL-6 Signaling | 0.649 | 0.0155 | #NUM! | CEBPB,FOS |
| Detoxification of Reactive Oxygen Species | 0.645 | 0.027 | #NUM! | PRDX3 |
| Cardiac conduction | 0.644 | 0.0154 | #NUM! | CASQ2,SLN |
| HGF Signaling | 0.639 | 0.0153 | #NUM! | FOS,ITGAM |
| MyD88-independent TLR4 cascade | 0.635 | 0.0263 | #NUM! | UBA52 |
| Regulation of TP53 Expression and Degradation | 0.635 | 0.0263 | #NUM! | UBA52 |
| Resolution of Abasic Sites (AP sites) | 0.635 | 0.0263 | #NUM! | RPA3 |

| | | | | |
|---|---|---|---|---|
| Mitotic G1 phase and G1/S transition | 0.634 | 0.0152 | #NUM! | CDK6,UBA52 |
| Orexin Signaling Pathway | 0.629 | 0.0124 | #NUM! | ATP6V1B1,GCG,GNB2 |
| Gα12/13 Signaling | 0.625 | 0.0149 | #NUM! | CDH3,MYL9 |
| P2Y Purinergic Receptor Signaling Pathway | 0.625 | 0.0149 | #NUM! | FOS,GNB2 |
| MAP kinase activation | 0.625 | 0.0256 | #NUM! | UBA52 |
| NGF-stimulated transcription | 0.625 | 0.0256 | #NUM! | FOS |
| FLT3 Signaling | 0.625 | 0.0256 | #NUM! | UBA52 |
| Complement System | 0.625 | 0.0256 | #NUM! | ITGAM |
| Class C/3 (Metabotropic glutamate/pheromone receptors) | 0.615 | 0.025 | #NUM! | CASR |
| Transcriptional Regulation by VENTX | 0.615 | 0.025 | #NUM! | CEBPB |
| Sertoli Cell-Sertoli Cell Junction Signaling | 0.612 | 0.0121 | #NUM! | CDH3,CLDN2,JUP |
| Ferroptosis Signaling Pathway | 0.611 | 0.0146 | #NUM! | ARF5,ARF6 |

| | | | | |
|---|---|---|---|---|
| Adipogenesis pathway | 0.607 | 0.0145 | #NUM! | CEBPB,FZD7 |
| Wound Healing Signaling Pathway | 0.605 | 0.012 | #NUM! | CALCA,CEBPB,FOS |
| MyD88:MAL(TIRAP) cascade initiated on plasma membrane | 0.597 | 0.0238 | #NUM! | UBA52 |
| April Mediated Signaling | 0.597 | 0.0238 | #NUM! | FOS |
| Signaling by NOTCH4 | 0.593 | 0.0142 | #NUM! | STK17B,UBA52 |
| SRP-dependent cotranslational protein targeting to membrane | 0.589 | 0.0141 | #NUM! | SPCS3,UBA52 |
| Signaling by SCF-KIT | 0.588 | 0.0233 | #NUM! | KIT |
| Transport of vitamins, nucleosides, and related molecules | 0.588 | 0.0233 | #NUM! | SLC28A3 |
| Smooth Muscle Contraction | 0.588 | 0.0233 | #NUM! | MYL9 |
| B Cell Activating Factor Signaling | 0.588 | 0.0233 | #NUM! | FOS |

| | | | | |
|---|---|---|---|---|
| Oncostatin M Signaling | 0.588 | 0.0233 | #NUM! | MT2A |
| Role of Chondrocytes in Rheumatoid Arthritis Signaling Pathway | 0.58 | 0.0139 | #NUM! | CEBPB,FOS |
| SUMOylation of transcription cofactors | 0.579 | 0.0227 | #NUM! | SUMO2 |
| TAK1-dependent IKK and NF-kappa-B activation | 0.579 | 0.0227 | #NUM! | UBA52 |
| TBC/RABGAPs | 0.579 | 0.0227 | #NUM! | ARF6 |
| Aggrephagy | 0.579 | 0.0227 | #NUM! | UBA52 |
| Retinoid metabolism and transport | 0.579 | 0.0227 | #NUM! | TTR |
| PIP3 activates AKT signaling | 0.576 | 0.0138 | #NUM! | KIT,NTF3 |
| Apelin Endothelial Signaling Pathway | 0.576 | 0.0138 | #NUM! | FOS,GNB2 |
| Hematoma Resolution Signaling Pathway | 0.574 | 0.0116 | #NUM! | FOS,ITGAM,S1PR1 |
| Signaling by Insulin receptor | 0.571 | 0.0222 | #NUM! | ATP6V1B1 |

| | | | | |
|---|---|---|---|---|
| Carboxyterminal post-translational modifications of tubulin | 0.571 | 0.0222 | #NUM! | AGBL2 |
| Gαi Signaling | 0.568 | 0.0136 | #NUM! | GNB2,S1PR1 |
| Semaphorin Neuronal Repulsive Signaling Pathway | 0.568 | 0.0136 | #NUM! | ITGAM,MYL9 |
| SUMOylation of DNA replication proteins | 0.563 | 0.0217 | #NUM! | SUMO2 |
| MIF Regulation of Innate Immunity | 0.563 | 0.0217 | #NUM! | FOS |
| Role of OCT4 in Mammalian Embryonic Stem Cell Pluripotency | 0.563 | 0.0217 | #NUM! | FOXD3 |
| Cohesin Chromatin Regulation Pathway | 0.562 | 0.0114 | #NUM! | FAT2,POLR2K,POLR2L |
| PTEN Regulation | 0.556 | 0.0133 | #NUM! | ATN1,UBA52 |
| Eukaryotic Translation Initiation | 0.556 | 0.0133 | #NUM! | EIF2S3,UBA52 |

| | | | | |
|---|---|---|---|---|
| SUMOylation of RNA binding proteins | 0.555 | 0.0213 | #NUM! | SUMO2 |
| Dilated Cardiomyopathy Signaling Pathway | 0.552 | 0.0132 | #NUM! | MYL9,TNNT2 |
| PTEN Signaling | 0.552 | 0.0132 | #NUM! | CNKSR3,ITGAM |
| KEAP1-NFE2L2 pathway | 0.548 | 0.0132 | #NUM! | STK17B,UBA52 |
| G alpha (z) signalling events | 0.547 | 0.0208 | #NUM! | GNB2 |
| Complex IV assembly | 0.547 | 0.0208 | #NUM! | COX7C |
| TP53 Regulates Transcription of Cell Cycle Genes | 0.54 | 0.0204 | #NUM! | GADD45A |
| Pyruvate metabolism | 0.54 | 0.0204 | #NUM! | UBA52 |
| Apelin Muscle Signaling Pathway | 0.54 | 0.0204 | #NUM! | GNB2 |
| Colorectal Cancer Metastasis Signaling | 0.539 | 0.0111 | #NUM! | FOS,FZD7,GNB2 |
| FAK Signaling | 0.537 | 0.00869 | -1.667 | CASR,FOS,FZD7,GPR55,GPR88,GPRC5C,ITGAM,KIT,S1PR1 |
| Signaling by ERBB2 | 0.532 | 0.02 | #NUM! | UBA52 |

| | | | | |
|---|---|---|---|---|
| Signaling by NOTCH3 | 0.532 | 0.02 | #NUM! | UBA52 |
| Meiotic recombination | 0.532 | 0.02 | #NUM! | RPA3 |
| Relaxin Signaling | 0.532 | 0.0128 | #NUM! | FOS,GNB2 |
| Gap junction trafficking and regulation | 0.525 | 0.0196 | #NUM! | GJD2 |
| Transcriptional activity of SMAD2/SMAD3: SMAD4 heterotrimer | 0.525 | 0.0196 | #NUM! | UBA52 |
| Transcriptional regulation of granulopoiesis | 0.525 | 0.0196 | #NUM! | CEBPB |
| UVC-Induced MAPK Signaling | 0.525 | 0.0196 | #NUM! | FOS |
| Cell Cycle: G2/M DNA Damage Checkpoint Regulation | 0.525 | 0.0196 | #NUM! | GADD45A |
| p75 NTR receptor-mediated signalling | 0.525 | 0.0127 | #NUM! | STK17B,UBA52 |
| TNFR1 Signaling | 0.518 | 0.0192 | #NUM! | FOS |
| Regulation of Apoptosis | 0.511 | 0.0189 | #NUM! | UBA52 |
| Signaling by EGFR | 0.511 | 0.0189 | #NUM! | UBA52 |

| | | | | |
|---|---|---|---|---|
| Xenobiotic Metabolism General Signaling Pathway | 0.507 | 0.0123 | #NUM! | Gstp1 (includes others),MGST3 |
| Signaling by TGF-beta Receptor Complex | 0.505 | 0.0185 | #NUM! | UBA52 |
| UVB-Induced MAPK Signaling | 0.505 | 0.0185 | #NUM! | FOS |
| Oxytocin Signaling Pathway | 0.501 | 0.0105 | #NUM! | FOS,GNB2,MYL9 |
| Meiotic synapsis | 0.498 | 0.0182 | #NUM! | SYCP1 |
| Signaling by PTK6 | 0.498 | 0.0182 | #NUM! | UBA52 |
| Amino acids regulate mTORC1 | 0.498 | 0.0182 | #NUM! | ATP6V1B1 |
| Sensory processing of sound by outer hair cells of the cochlea | 0.498 | 0.0182 | #NUM! | KCNMB1 |
| CSDE1 Signaling Pathway | 0.492 | 0.0179 | #NUM! | FOS |
| NLR signaling pathways | 0.492 | 0.0179 | #NUM! | UBA52 |
| Metabolism of non-coding RNA | 0.492 | 0.0179 | #NUM! | SNRPF |
| DNA Damage/Telomere Stress Induced Senescence | 0.492 | 0.0179 | #NUM! | HMGA1 |

| | | | | |
|---|---|---|---|---|
| Deadenylation-dependent mRNA decay | 0.492 | 0.0179 | #NUM! | LSM5 |
| Interleukin-3, Interleukin-5 and GM-CSF signaling | 0.492 | 0.0179 | #NUM! | UBA52 |
| E3 ubiquitin ligases ubiquitinate target proteins | 0.492 | 0.0179 | #NUM! | UBA52 |
| CD27 Signaling in Lymphocytes | 0.492 | 0.0179 | #NUM! | FOS |
| EGF Signaling | 0.492 | 0.0179 | #NUM! | FOS |
| Neurexins and neuroligins | 0.485 | 0.0175 | #NUM! | APBA3 |
| TNF signaling | 0.485 | 0.0175 | #NUM! | UBA52 |
| Folate Signaling Pathway | 0.485 | 0.0175 | #NUM! | FOLR1 |
| Germ Cell-Sertoli Cell Junction Signaling | 0.479 | 0.0117 | #NUM! | CTNND1,JUP |
| Glioblastoma Multiforme Signaling | 0.479 | 0.0117 | #NUM! | CDK6,FZD7 |
| Phagosome Maturation | 0.479 | 0.0117 | #NUM! | ATP6V1B1,Dynlt1b (includes others) |
| Circadian Clock | 0.479 | 0.0172 | #NUM! | UBA52 |

| | | | | |
|---|---|---|---|---|
| Cytoprotection by HMOX1 | 0.479 | 0.0172 | #NUM! | COX7C |
| Signaling by ERBB4 | 0.473 | 0.0169 | #NUM! | UBA52 |
| DNA Double Strand Break Response | 0.473 | 0.0169 | #NUM! | UBA52 |
| Polyamine Regulation in Colon Cancer | 0.473 | 0.0169 | #NUM! | FOS |
| Sirtuin Signaling Pathway | 0.471 | 0.0101 | #NUM! | GADD45A,TOMM7,ZIC2 |
| LPS/IL-1 Mediated Inhibition of RXR Function | 0.468 | 0.0101 | #NUM! | Cyp2j11/Cyp2j8,Gstp1 (includes others),MGST3 |
| NIK-->noncanonical NF-kB signaling | 0.467 | 0.0167 | #NUM! | UBA52 |
| GADD45 Signaling | 0.467 | 0.0167 | #NUM! | GADD45A |
| PCP (Planar Cell Polarity) Pathway | 0.467 | 0.0167 | #NUM! | FZD7 |
| Visual phototransduction | 0.462 | 0.0164 | #NUM! | TTR |
| Senescence Pathway | 0.461 | 0.00997 | #NUM! | CDK6,CEBPB,GADD45A |
| MSP-RON Signaling Pathway | 0.456 | 0.0161 | #NUM! | ITGAM |

| | | | | |
|---|---|---|---|---|
| Transcriptional Regulation by MECP2 | 0.45 | 0.0159 | #NUM! | CRH |
| IL-2 Signaling | 0.45 | 0.0159 | #NUM! | FOS |
| Erythropoietin Signaling Pathway | 0.448 | 0.011 | #NUM! | FOS,HBA2 |
| Semaphorin interactions | 0.445 | 0.0156 | #NUM! | MYL9 |
| Peroxisomal protein import | 0.445 | 0.0156 | #NUM! | UBA52 |
| Thrombopoietin Signaling | 0.445 | 0.0156 | #NUM! | FOS |
| Autism Signaling Pathway | 0.443 | 0.00971 | #NUM! | FOLR1,FZD7, GLI1 |
| Mitochondrial protein import | 0.44 | 0.0154 | #NUM! | TOMM7 |
| Hedgehog ligand biogenesis | 0.44 | 0.0154 | #NUM! | UBA52 |
| Pyridoxal 5'-phosphate Salvage Pathway | 0.434 | 0.0152 | #NUM! | CDK6 |
| Oxidative Ethanol Degradation III | 0.434 | 0.0152 | #NUM! | Cyp2j11/Cyp2 j8 |
| WNT/Ca+ pathway | 0.434 | 0.0152 | #NUM! | FZD7 |
| MicroRNA Biogenesis Signaling Pathway | 0.43 | 0.0107 | #NUM! | POLR2K,POL R2L |

| | | | | |
|---|---|---|---|---|
| Abacavir ADME | 0.429 | 0.0149 | #NUM! | STK17B |
| CD40 Signaling | 0.429 | 0.0149 | #NUM! | FOS |
| SPINK1 General Cancer Pathway | 0.429 | 0.0149 | #NUM! | MT2A |
| Regulation of eIF4 and p70S6K Signaling | 0.427 | 0.0106 | #NUM! | EIF2S3,ITGAM |
| Role of Osteoclasts in Rheumatoid Arthritis Signaling Pathway | 0.425 | 0.00946 | #NUM! | ADAM18,FOS,SFRP5 |
| IL-17 Signaling | 0.424 | 0.0106 | #NUM! | CEBPB,FOS |
| TNFR2 non-canonical NF-kB pathway | 0.424 | 0.0147 | #NUM! | UBA52 |
| Glutamate Receptor Signaling | 0.424 | 0.0147 | #NUM! | GNB2 |
| RHO GTPase cycle | 0.422 | 0.00889 | -2 | CHN2,FRS3,JUP,KCTD13 |
| Cell Cycle: G1/S Checkpoint Regulation | 0.419 | 0.0145 | #NUM! | CDK6 |
| Macrophage Alternative Activation Signaling Pathway | 0.416 | 0.0104 | #NUM! | CEBPB,FOS |
| GNRH Signaling | 0.413 | 0.0104 | #NUM! | FOS,GNB2 |

| | | | | |
|---|---|---|---|---|
| SUMOylation of chromatin organization proteins | 0.41 | 0.0141 | #NUM! | SUMO2 |
| IL-33 Signaling Pathway | 0.408 | 0.0103 | #NUM! | FOS,KIT |
| FXR/RXR Activation | 0.408 | 0.0103 | #NUM! | Gstp1 (includes others),MGST3 |
| Pulmonary Fibrosis Idiopathic Signaling Pathway | 0.406 | 0.0092 | #NUM! | FOS,FZD7,GLI1 |
| Regulation of RUNX2 expression and activity | 0.405 | 0.0139 | #NUM! | UBA52 |
| Dopamine Receptor Signaling | 0.405 | 0.0139 | #NUM! | PRL |
| CDX Gastrointestinal Cancer Signaling Pathway | 0.402 | 0.0102 | #NUM! | FOS,FZD7 |
| ILK Signaling | 0.402 | 0.0102 | #NUM! | FOS,MYL9 |
| Ephrin B Signaling | 0.4 | 0.0137 | #NUM! | GNB2 |
| Synaptic Long Term Depression | 0.4 | 0.0101 | #NUM! | CRH,PPP1R17 |
| Glucocorticoid Receptor Signaling | 0.397 | 0.00833 | #NUM! | FOS,POLR2K,POLR2L,PRL,TAF7L |

| | | | | |
|---|---|---|---|---|
| ISG15 antiviral mechanism | 0.396 | 0.0135 | #NUM! | UBA52 |
| Olfactory Signaling Pathway | 0.396 | 0.0135 | #NUM! | ANO2 |
| ERK5 Signaling | 0.396 | 0.0135 | #NUM! | FOS |
| Adrenomedullin signaling pathway | 0.389 | 0.0099 | #NUM! | CEBPB,FOS |
| Ephrin Receptor Signaling | 0.389 | 0.0099 | #NUM! | GNB2,ITGAM |
| Cellular response to hypoxia | 0.387 | 0.0132 | #NUM! | UBA52 |
| Plasma lipoprotein assembly, remodeling, and clearance | 0.387 | 0.0132 | #NUM! | UBA52 |
| Interferon alpha/beta signaling | 0.387 | 0.0132 | #NUM! | UBA52 |
| GDNF Family Ligand-Receptor Interactions | 0.387 | 0.0132 | #NUM! | FOS |
| Macropinocytosis Signaling | 0.387 | 0.0132 | #NUM! | ARF6 |
| Sheddase Signaling Pathway | 0.384 | 0.0098 | #NUM! | CEBPB,FOS |
| Granulocyte Adhesion and Diapedesis | 0.384 | 0.0098 | #NUM! | CLDN2,ITGAM |

| | | | | |
|---|---|---|---|---|
| Signaling by NOTCH1 | 0.383 | 0.013 | #NUM! | UBA52 |
| SUMOylation of DNA damage response and repair proteins | 0.383 | 0.013 | #NUM! | SUMO2 |
| Antiproliferative Role of Somatostatin Receptor 2 | 0.383 | 0.013 | #NUM! | GNB2 |
| Leptin Signaling in Obesity | 0.383 | 0.013 | #NUM! | NPY |
| Fc epsilon receptor (FCERI) signaling | 0.379 | 0.00971 | #NUM! | FOS,UBA52 |
| Maturity Onset Diabetes of Young (MODY) Signaling | 0.378 | 0.0128 | #NUM! | NEUROD1 |
| Role of WNT/GSK-3β Signaling in the Pathogenesis of Influenza | 0.378 | 0.0128 | #NUM! | FZD7 |
| Caveolar-mediated Endocytosis Signaling | 0.374 | 0.0127 | #NUM! | ITGAM |
| Clathrin-mediated Endocytosis Signaling | 0.372 | 0.00957 | #NUM! | ARF6,UBA52 |

| | | | | |
|---|---|---|---|---|
| G alpha (12/13) signalling events | 0.37 | 0.0125 | #NUM! | GNB2 |
| Role of BRCA1 in DNA Damage Response | 0.37 | 0.0125 | #NUM! | GADD45A |
| IL-3 Signaling | 0.366 | 0.0123 | #NUM! | FOS |
| Role of JAK family kinases in IL-6-type Cytokine Signaling | 0.366 | 0.0123 | #NUM! | FOS |
| Chemokine Signaling | 0.366 | 0.0123 | #NUM! | FOS |
| BEX2 Signaling Pathway | 0.362 | 0.0122 | #NUM! | NHLH2 |
| Estrogen-Dependent Breast Cancer Signaling | 0.358 | 0.012 | #NUM! | FOS |
| Immunoregulatory interactions between a Lymphoid and a non-Lymphoid cell | 0.357 | 0.0093 | #NUM! | CRTAM,PIANP |
| PI Metabolism | 0.354 | 0.0119 | #NUM! | TNFAIP8L3 |
| Transcriptional regulation of white adipocyte differentiation | 0.354 | 0.0119 | #NUM! | CEBPB |

| | | | | |
|---|---|---|---|---|
| Integrin cell surface interactions | 0.35 | 0.0118 | #NUM! | ITGAM |
| Activation of NMDA receptors and postsynaptic events | 0.35 | 0.0118 | #NUM! | NRGN |
| TP53 Regulates Metabolic Genes | 0.35 | 0.0118 | #NUM! | COX7C |
| LPS-stimulated MAPK Signaling | 0.35 | 0.0118 | #NUM! | FOS |
| Autophagy | 0.348 | 0.00913 | #NUM! | FOS,GCG |
| Cyclins and Cell Cycle Regulation | 0.347 | 0.0116 | #NUM! | CDK6 |
| VEGF Family Ligand-Receptor Interactions | 0.347 | 0.0116 | #NUM! | FOS |
| ERK/MAPK Signaling | 0.346 | 0.00909 | #NUM! | FOS,ITGAM |
| Xenobiotic Metabolism PXR Signaling Pathway | 0.344 | 0.00905 | #NUM! | Gstp1 (includes others),MGST3 |
| PDGF Signaling | 0.343 | 0.0115 | #NUM! | FOS |
| Regulation of mitotic cell cycle | 0.339 | 0.0114 | #NUM! | UBA52 |
| Respiratory electron transport | 0.339 | 0.0114 | #NUM! | COX7C |

| | | | | |
|---|---|---|---|---|
| Thrombin Signaling | 0.339 | 0.00897 | #NUM! | GNB2,MYL9 |
| Agranulocyte Adhesion and Diapedesis | 0.337 | 0.00893 | #NUM! | CLDN2,MYL9 |
| Regulation of mRNA stability by proteins that bind AU-rich elements | 0.336 | 0.0112 | #NUM! | UBA52 |
| Opioid Signalling | 0.332 | 0.0111 | #NUM! | GNB2 |
| Degradation of beta-catenin by the destruction complex | 0.329 | 0.011 | #NUM! | UBA52 |
| Actin Nucleation by ARP-WASP Complex | 0.329 | 0.011 | #NUM! | ITGAM |
| MAPK6/MAPK4 signaling | 0.326 | 0.0109 | #NUM! | UBA52 |
| Role of Hypercytokinemia/hyperchemokinemia in the Pathogenesis of Influenza | 0.326 | 0.0109 | #NUM! | S1PR1 |
| Unfolded protein response | 0.326 | 0.0109 | #NUM! | CEBPB |
| EPH-Ephrin signaling | 0.322 | 0.0108 | #NUM! | MYL9 |

| | | | | |
|---|---|---|---|---|
| Fcγ Receptor-mediated Phagocytosis in Macrophages and Monocytes | 0.322 | 0.0108 | #NUM! | ARF6 |
| RANK Signaling in Osteoclasts | 0.322 | 0.0108 | #NUM! | FOS |
| ERBB Signaling | 0.322 | 0.0108 | #NUM! | FOS |
| Xenobiotic Metabolism CAR Signaling Pathway | 0.318 | 0.00858 | #NUM! | Gstp1 (includes others),MGST3 |
| Transcriptional regulation by RUNX3 | 0.312 | 0.0104 | #NUM! | UBA52 |
| Non-Small Cell Lung Cancer Signaling | 0.312 | 0.0104 | #NUM! | CDK6 |
| Mitotic Metaphase and Anaphase | 0.312 | 0.00847 | #NUM! | RCC1,UBA52 |
| Mitochondrial translation | 0.309 | 0.0103 | #NUM! | MRPL53 |
| TGF-β Signaling | 0.309 | 0.0103 | #NUM! | FOS |
| VEGF Signaling | 0.309 | 0.0103 | #NUM! | EIF2S3 |
| EIF2 Signaling | 0.308 | 0.0084 | #NUM! | EIF2S3,UBA52 |
| p53 Signaling | 0.306 | 0.0102 | #NUM! | GADD45A |
| Interferon gamma signaling | 0.306 | 0.0102 | #NUM! | MT2A |

| | | | | |
|---|---|---|---|---|
| Small Cell Lung Cancer Signaling | 0.306 | 0.0102 | #NUM! | CDK6 |
| UVA-Induced MAPK Signaling | 0.306 | 0.0102 | #NUM! | FOS |
| Role of Osteoblasts in Rheumatoid Arthritis Signaling Pathway | 0.305 | 0.00833 | #NUM! | FZD7,SFRP5 |
| Melanocyte Development and Pigmentation Signaling | 0.303 | 0.0101 | #NUM! | KIT |
| cAMP-mediated signaling | 0.303 | 0.0083 | #NUM! | Htr5b,S1PR1 |
| Actin Cytoskeleton Signaling | 0.301 | 0.00826 | #NUM! | ITGAM,MYL9 |
| S Phase | 0.3 | 0.01 | #NUM! | UBA52 |
| ATM Signaling | 0.3 | 0.01 | #NUM! | GADD45A |
| Salvage Pathways of Pyrimidine Ribonucleotides | 0.3 | 0.01 | #NUM! | CDK6 |
| Apelin Cardiomyocyte Signaling Pathway | 0.3 | 0.01 | #NUM! | MYL9 |
| Pancreatic Secretion Signaling Pathway | 0.299 | 0.00823 | #NUM! | AQP1,CA8 |

| | | | | |
|---|---|---|---|---|
| Cellular response to heat stress | 0.297 | 0.0099 | #NUM! | RPA3 |
| Neuropathic Pain Signaling in Dorsal Horn Neurons | 0.297 | 0.0099 | #NUM! | FOS |
| Neddylation | 0.293 | 0.00813 | #NUM! | FBXO27,UBA52 |
| DNA Replication Pre-Initiation | 0.288 | 0.00962 | #NUM! | UBA52 |
| Regulation of endogenous retroelements | 0.288 | 0.00962 | #NUM! | SUMO2 |
| Cyclophilin Signaling Pathway | 0.288 | 0.00803 | #NUM! | PRL,TOMM7 |
| Protein Kinase A Signaling | 0.286 | 0.00758 | #NUM! | DUSP5,GNB2,MYL9 |
| Cargo recognition for clathrin-mediated endocytosis | 0.286 | 0.00952 | #NUM! | UBA52 |
| IGF-1 Signaling | 0.286 | 0.00952 | #NUM! | FOS |
| Sleep REM Signaling Pathway | 0.283 | 0.00943 | #NUM! | FOS |
| WNK Renal Signaling Pathway | 0.28 | 0.00935 | #NUM! | CASR |

| | | | | |
|---|---|---|---|---|
| Transport of inorganic cations/anions and amino acids/oligopeptides | 0.277 | 0.00926 | #NUM! | SLC25A22 |
| PPAR Signaling | 0.277 | 0.00926 | #NUM! | FOS |
| Phase II - Conjugation of compounds | 0.275 | 0.00917 | #NUM! | MGST3 |
| Sphingolipid metabolism | 0.272 | 0.00909 | #NUM! | NEU4 |
| α-Adrenergic Signaling | 0.269 | 0.00901 | #NUM! | GNB2 |
| Cardiac Hypertrophy Signaling | 0.267 | 0.00766 | #NUM! | GNB2,MYL9 |
| Estrogen Receptor Signaling | 0.265 | 0.0073 | #NUM! | FOS,GNB2,MYL9 |
| Role of MAPK Signaling in Promoting the Pathogenesis of Influenza | 0.262 | 0.00877 | #NUM! | ATP6V1B1 |
| Neuregulin Signaling | 0.252 | 0.00847 | #NUM! | ITGAM |
| Sphingosine-1-phosphate Signaling | 0.249 | 0.0084 | #NUM! | S1PR1 |
| Cell Cycle Checkpoints | 0.249 | 0.00735 | #NUM! | RPA3,UBA52 |

| | | | | |
|---|---|---|---|---|
| Virus Entry via Endocytic Pathways | 0.247 | 0.00833 | #NUM! | FOLR1 |
| Eukaryotic Translation Elongation | 0.243 | 0.0082 | #NUM! | UBA52 |
| Eukaryotic Translation Termination | 0.243 | 0.0082 | #NUM! | UBA52 |
| Renin-Angiotensin Signaling | 0.243 | 0.0082 | #NUM! | FOS |
| IL-13 Signaling Pathway | 0.24 | 0.00813 | #NUM! | SERPINE3 |
| RHOA Signaling | 0.24 | 0.00813 | #NUM! | MYL9 |
| RAR Activation | 0.238 | 0.00693 | #NUM! | FOS,KIT,PAX6 |
| Opioid Signaling Pathway | 0.236 | 0.00712 | #NUM! | FOS,GNB2 |
| Asparagine N-linked glycosylation | 0.236 | 0.008 | #NUM! | UBA52 |
| Eicosanoid Signaling | 0.235 | 0.00709 | #NUM! | GNB2,KCNMB1 |
| TCR signaling | 0.234 | 0.00794 | #NUM! | UBA52 |
| Glioma Signaling | 0.229 | 0.00781 | #NUM! | CDK6 |
| Interleukin-1 family signaling | 0.227 | 0.00775 | #NUM! | UBA52 |
| LXR/RXR Activation | 0.225 | 0.00769 | #NUM! | TTR |
| G Beta Gamma Signaling | 0.225 | 0.00769 | #NUM! | GNB2 |

| | | | | |
|---|---|---|---|---|
| Costimulation by the CD28 family | 0.223 | 0.00763 | #NUM! | STK17B |
| 14-3-3-mediated Signaling | 0.221 | 0.00758 | #NUM! | FOS |
| fMLP Signaling in Neutrophils | 0.219 | 0.00752 | #NUM! | GNB2 |
| CCR3 Signaling in Eosinophils | 0.213 | 0.00735 | #NUM! | GNB2 |
| RAC Signaling | 0.211 | 0.0073 | #NUM! | ITGAM |
| SNARE Signaling Pathway | 0.21 | 0.00725 | #NUM! | MYL9 |
| CGAS-STING Signaling Pathway | 0.21 | 0.00725 | #NUM! | ATP6V1B1 |
| Role of PKR in Interferon Induction and Antiviral Response | 0.21 | 0.00725 | #NUM! | FOS |
| White Adipose Tissue Browning Pathway | 0.208 | 0.00719 | #NUM! | CEBPB |
| Th2 Pathway | 0.204 | 0.00709 | #NUM! | S1PR1 |
| DHCR24 Signaling Pathway | 0.2 | 0.00699 | #NUM! | TTR |
| MSP-RON Signaling in Cancer Cells Pathway | 0.2 | 0.00699 | #NUM! | FOS |

| | | | | |
|---|---|---|---|---|
| Oxytocin in Brain Signaling Pathway | 0 | 0.00498 | #NUM! | GNB2 |
| Pulmonary Healing Signaling Pathway | 0 | 0.00498 | #NUM! | FZD7 |
| CLEAR Signaling Pathway | 0 | 0.00351 | #NUM! | ATP6V1B1 |
| Pathogen Induced Cytokine Storm Signaling Pathway | 0 | 0.00524 | #NUM! | EOMES,FOS |
| IL-10 Signaling | 0 | 0.00645 | #NUM! | FOS |
| Neutrophil Extracellular Trap Signaling Pathway | 0 | 0.00489 | #NUM! | ITGAM,TOMM7 |
| Circadian Rhythm Signaling | 0 | 0.0037 | #NUM! | GNB2 |
| Chaperone Mediated Autophagy Signaling Pathway | 0 | 0.00157 | #NUM! | ATP6V1B1 |
| NOD1/2 Signaling Pathway | 0 | 0.00518 | #NUM! | FOS |
| Adrenergic Receptor Signaling Pathway (Enhanced) | 0 | 0.00505 | #NUM! | ATP6V1B1 |
| Glutaminergic Receptor Signaling Pathway (Enhanced) | 0 | 0.00613 | #NUM! | GABRA6,GABRD |

| | | | | |
|---|---|---|---|---|
| PPARα/RXRα Activation | 0 | 0.00505 | #NUM! | GH1 |
| TCF dependent signaling in response to WNT | 0 | 0.00505 | #NUM! | UBA52 |
| Cell surface interactions at the vascular wall | 0 | 0.00467 | #NUM! | ITGAM |
| Generic Transcription Pathway | 0 | 0.00233 | #NUM! | NR4A3 |
| Selenoamino acid metabolism | 0 | 0.00685 | #NUM! | UBA52 |
| Signaling by ROBO receptors | 0 | 0.00407 | #NUM! | UBA52 |
| Mitotic G2-G2/M phases | 0 | 0.005 | #NUM! | UBA52 |
| C-type lectin receptors (CLRs) | 0 | 0.0069 | #NUM! | UBA52 |
| Deubiquitination | 0 | 0.00377 | #NUM! | UBA52 |
| Major pathway of rRNA processing in the nucleolus and cytosol | 0 | 0.00538 | #NUM! | UBA52 |
| Neutrophil degranulation | 0 | 0.00629 | #NUM! | ITGAM,JUP,TTR |
| Keratinization | 0 | 0.00463 | #NUM! | JUP |

| | | | | |
|---|---|---|---|---|
| Intra-Golgi and retrograde Golgi-to-ER traffic | 0 | 0.0049 | #NUM! | ARF5 |
| Transcriptional regulation by RUNX1 | 0 | 0.00658 | #NUM! | UBA52 |
| Class I MHC mediated antigen processing and presentation | 0 | 0.00525 | #NUM! | FBXO27,UBA52 |
| Signaling by the B Cell Receptor (BCR) | 0 | 0.00588 | #NUM! | UBA52 |
| Hepatic Cholestasis | 0 | 0.00444 | #NUM! | GCG |
| Hepatic Fibrosis / Hepatic Stellate Cell Activation | 0 | 0.00521 | #NUM! | MYL9 |
| NAFLD Signaling Pathway | 0 | 0.00446 | #NUM! | FOS |
| Histone Modification Signaling Pathway | 0 | 0.00333 | #NUM! | ZIC2 |
| Mitochondrial Division Signaling Pathway | 0 | 0.00602 | #NUM! | S1PR1 |
| Ribosomal Quality Control Signaling Pathway | 0 | 0.00379 | #NUM! | UBA52 |

| | | | | |
|---|---|---|---|---|
| Irritable Bowel Syndrome Signaling Pathway | 0 | 0.00346 | #NUM! | MYL9 |
| TRIM21 Intracellular Antibody Signaling Pathway | 0 | 0.00353 | #NUM! | FOS,UBA52 |
| Role of NFAT in Regulation of the Immune Response | 0 | 0.00191 | #NUM! | FOS,GNB2 |
| CCR5 Signaling in Macrophages | 0 | 0.00399 | #NUM! | FOS,GNB2 |
| CTLA4 Signaling in Cytotoxic T Lymphocytes | 0 | 0.00164 | #NUM! | FOS |
| CD28 Signaling in T Helper Cells | 0 | 0.00192 | #NUM! | FOS |
| Endothelin-1 Signaling | 0 | 0.00515 | #NUM! | FOS |
| Factors Promoting Cardiogenesis in Vertebrates | 0 | 0.00662 | #NUM! | FZD7 |
| Lipid Antigen Presentation by CD1 | 0 | 0.00242 | #NUM! | ARF6 |

| | | | | |
|---|---|---|---|---|
| HMGB1 Signaling | 0 | 0.00625 | #NUM! | FOS |
| Aldosterone Signaling in Epithelial Cells | 0 | 0.00585 | #NUM! | KCNMB1 |
| Type II Diabetes Mellitus Signaling | 0 | 0.00658 | #NUM! | CEBPB |
| Production of Nitric Oxide and Reactive Oxygen Species in Macrophages | 0 | 0.00521 | #NUM! | FOS |
| G Protein Signaling Mediated by Tubby | 0 | 0.00213 | #NUM! | GNB2 |
| Systemic Lupus Erythematosus Signaling | 0 | 0.00281 | #NUM! | FOS,LSM5,SNRPF |
| CDC42 Signaling | 0 | 0.00519 | #NUM! | FOS,ITGAM,MYL9 |
| AMPK Signaling | 0 | 0.0041 | #NUM! | GNB2 |
| Phospholipase C Signaling | 0 | 0.00267 | #NUM! | GNB2,ITGAM,MYL9 |
| Ovarian Cancer Signaling | 0 | 0.00629 | #NUM! | FZD7 |
| Role of NFAT in Cardiac Hypertrophy | 0 | 0.00437 | #NUM! | GNB2 |

| | | | | |
|---|---|---|---|---|
| Regulation of IL-2 Expression in Activated and Anergic T Lymphocytes | 0 | 0.00214 | #NUM! | FOS |
| PKCθ Signaling in T Lymphocytes | 0 | 0.00179 | #NUM! | FOS |
| PI3K Signaling in B Lymphocytes | 0 | 0.00169 | #NUM! | FOS |
| Role of Tissue Factor in Cancer | 0 | 0.00481 | #NUM! | FOS |
| Gap Junction Signaling | 0 | 0.00299 | #NUM! | GJD2 |
| eNOS Signaling | 0 | 0.00621 | #NUM! | AQP1 |
| D-myo-inositol-5-phosphate Metabolism | 0 | 0.0049 | #NUM! | DUSP5 |
| D-myo-inositol (1,4,5,6)-Tetrakisphosphate Biosynthesis | 0 | 0.00529 | #NUM! | DUSP5 |
| Superpathway of Inositol Phosphate Compounds | 0 | 0.00415 | #NUM! | DUSP5 |

| | | | | |
|---|---|---|---|---|
| D-myo-inositol (3,4,5,6)-tetrakisphosphate Biosynthesis | 0 | 0.00529 | #NUM! | DUSP5 |
| 3-phosphoinositide Degradation | 0 | 0.005 | #NUM! | DUSP5 |
| 3-phosphoinositide Biosynthesis | 0 | 0.00465 | #NUM! | DUSP5 |
| Epithelial Adherens Junction Signaling | 0 | 0.00633 | #NUM! | CTNND1 |
| Gαq Signaling | 0 | 0.00592 | #NUM! | GNB2 |
| Regulation of the Epithelial-Mesenchymal Transition Pathway | 0 | 0.00515 | #NUM! | FZD7 |
| TEC Kinase Signaling | 0 | 0.00519 | #NUM! | FOS,GNB2,ITGAM |
| Parkinson's Signaling Pathway | 0 | 0.00633 | #NUM! | ATP6V1B1,UBA52 |
| SAPK/JNK Signaling | 0 | 0.00398 | #NUM! | GADD45A,GNB2 |
| PI3K/AKT Signaling | 0 | 0.005 | #NUM! | ITGAM |
| Cardiac β-adrenergic Signaling | 0 | 0.00588 | #NUM! | GNB2 |
| Protein Ubiquitination Pathway | 0 | 0.00358 | #NUM! | UBA52 |

| | | | | |
|---|---|---|---|---|
| Xenobiotic Metabolism Signaling | 0 | 0.00604 | #NUM! | Gstp1 (includes others),MGST3 |
| Serotonin Receptor Signaling | 0 | 0.00638 | #NUM! | GNB2,Htr5b, SLC6A4 |
| NF-κB Signaling | 0 | 0.00175 | #NUM! | GH1 |
| T Cell Receptor Signaling | 0 | 0.00322 | #NUM! | DUSP5,FOS |
| Th1 and Th2 Activation Pathway | 0 | 0.00568 | #NUM! | S1PR1 |
| Endocannabinoid Neuronal Synapse Pathway | 0 | 0.00667 | #NUM! | GNB2 |
| Endocannabinoid Cancer Inhibition Pathway | 0 | 0.0068 | #NUM! | GPR55 |
| Cardiac Hypertrophy Signaling (Enhanced) | 0 | 0.00559 | #NUM! | FZD7,GNB2,ITGAM |
| T Cell Exhaustion Signaling Pathway | 0 | 0.00353 | #NUM! | EOMES,FOS |
| Synaptogenesis Signaling Pathway | 0 | 0.00633 | #NUM! | CDH3,CTNND1 |
| Systemic Lupus Erythematosus in T Cell Signaling Pathway | 0 | 0.0031 | #NUM! | FOS,GADD45A |

| | | | | |
|---|---|---|---|---|
| Systemic Lupus Erythematosus in B Cell Signaling Pathway | 0 | 0.00137 | #NUM! | FOS |
| Necroptosis Signaling Pathway | 0 | 0.00613 | #NUM! | TOMM7 |
| Regulation of the Epithelial Mesenchymal Transition by Growth Factors Pathway | 0 | 0.00515 | #NUM! | FOS |
| Coronavirus Pathogenesis Pathway | 0 | 0.00474 | #NUM! | FOS |
| Coronavirus Replication Pathway | 0 | 0.00602 | #NUM! | UBA52 |
| Tumor Microenvironment Pathway | 0 | 0.00556 | #NUM! | FOS |
