## Supplemental Table 10 for "Early Life Adversity Produces Enduring Molecular and Functional Disruption of Developing Vagal Circuits"

© 2000-2025 QIAGEN. All rights reserved.

**P120 EXCITATORY NEURONS MALES**

| Ingenuity Canonical Pathways | -log(p-value) | Ratio | z-score | Molecules |
|---|---|---|---|---|
| Class A/1 (Rhodopsin-like receptors) | 4.99 | 0.0271 | -1 | F2,GAL,GHRL,NPFF,PRLH,S1PR1,SST,TAC3,TRH |
| Signaling by FGFR2 | 3.62 | 0.0548 | -2 | FRS3,POLR2K,POLR2L,UBA52 |
| RNA Polymerase II Transcription | 3.39 | 0.0333 | -2.236 | ALYREF,POLR2K,POLR2L,SNRPF,TAF7L |
| Activation of anterior HOX genes in hindbrain during early embryogenesis | 3.38 | 0.0476 | -2 | KDM6A,PAX6,POLR2K,POLR2L |
| Nucleotide Excision Repair Pathway | 3.32 | 0.0811 | #NUM! | POLR2K,POLR2L,RPA3 |
| Fanconi Anemia Pathway | 3.28 | 0.0789 | #NUM! | RPA3,UBA52,UBE2T |
| NER (Nucleotide Excision Repair, Enhanced Pathway) | 3.25 | 0.044 | #NUM! | POLR2K,POLR2L,RPA3,TOP2A |
| G alpha (q) signalling events | 3.1 | 0.0287 | -0.447 | F2,GHRL,NPFF,TAC3,TRH |
| Formation of the anterior neural plate | 3.04 | 0.182 | #NUM! | OTX2,PAX6 |
| Nucleotide Excision Repair | 2.96 | 0.0367 | -2 | POLR2K,POLR2L,RPA3,UBA52 |
| Assembly of RNA Polymerase II Complex | 2.91 | 0.0588 | #NUM! | POLR2K,POLR2L,TAF7L |
| B-WICH complex positively regulates rRNA expression | 2.88 | 0.0577 | #NUM! | POLR2K,POLR2L,TAF1D |
| Response of EIF2AK1 (HRI) to heme deficiency | 2.77 | 0.133 | #NUM! | CEBPB,EIF2S3 |

| | | | | |
|---|---|---|---|---|
| ROBO SLIT Signaling Pathway | 2.7 | 0.0312 | 0 | CTNND1,FOS,MYL9,PAX6 |
| Cytosolic sensors of pathogen-associated DNA | 2.62 | 0.0469 | #NUM! | POLR2K,POLR2L,UBA52 |
| NoRC negatively regulates rRNA expression | 2.57 | 0.0448 | #NUM! | POLR2K,POLR2L,TAF1D |
| Synthesis, secretion, and deacylation of Ghrelin | 2.51 | 0.1 | #NUM! | GHRL,SPCS3 |
| TP53 Regulates Transcription of DNA Repair Genes | 2.51 | 0.0429 | #NUM! | FOS,POLR2K,POLR2L |
| Iron homeostasis signaling pathway | 2.5 | 0.0276 | #NUM! | Atp6ap1l,BMP4,HBA2,Hbb-bs/Hbb-bt |
| Senescence-Associated Secretory Phenotype (SASP) | 2.44 | 0.0405 | #NUM! | CEBPB,FOS,UBA52 |
| RNA Polymerase I Transcription | 2.44 | 0.0405 | #NUM! | POLR2K,POLR2L,TAF1D |
| GPCR-Mediated Integration of Enteroendocrine Signaling Exemplified by an L Cell | 2.41 | 0.0395 | #NUM! | GAL,SST,VIP |
| Incretin synthesis, secretion, and inactivation | 2.32 | 0.08 | #NUM! | PAX6,SPCS3 |
| Telomere Maintenance | 2.22 | 0.0337 | #NUM! | POLR2K,POLR2L,RPA3 |
| Regulation of TP53 Activity through Phosphorylation | 2.17 | 0.0323 | #NUM! | RPA3,TAF7L,UBA52 |
| mRNA Capping | 2.17 | 0.0667 | #NUM! | POLR2K,POLR2L |
| Protein ubiquitination | 2.17 | 0.0667 | #NUM! | UBA52,UBE2T |
| Processing of Capped Intron-Containing Pre-mRNA | 2.16 | 0.0174 | -2.236 | ALYREF,LSM5,POLR2K,POLR2L,SNRPF |
| Acute Phase Response Signaling | 2.08 | 0.0209 | #NUM! | CEBPB,CRABP1,F2,FOS |
| Response of EIF2AK4 (GCN2) to amino acid deficiency | 2.04 | 0.0291 | #NUM! | CEBPB,EIF2S3,UBA52 |
| Mitophagy | 1.92 | 0.05 | #NUM! | TOMM7,UBA52 |

| | | | | |
|---|---|---|---|---|
| Signaling by FGFR3 | 1.92 | 0.05 | #NUM! | FRS3,UBA52 |
| Metabolism of vitamin K | 1.91 | 0.333 | #NUM! | VKORC1 |
| Signaling by FGFR4 | 1.9 | 0.0488 | #NUM! | FRS3,UBA52 |
| RNA Polymerase III Transcription | 1.9 | 0.0488 | #NUM! | POLR2K,POLR2L |
| ESR-mediated signaling | 1.89 | 0.0254 | #NUM! | FOS,POLR2K,POLR2L |
| Regulation of beta-cell development | 1.88 | 0.0476 | #NUM! | NEUROD1,PAX6 |
| Intrinsic Prothrombin Activation Pathway | 1.86 | 0.0465 | #NUM! | F2,KLK12 |
| RHOA Signaling | 1.84 | 0.0244 | #NUM! | CDC42EP5,MYL9,TTN |
| Regulation of Insulin-like Growth Factor (IGF) transport and uptake by IGFBPs | 1.83 | 0.0242 | #NUM! | BMP4,F2,SPP1 |
| Signaling by Retinoic Acid | 1.81 | 0.0435 | #NUM! | CRABP1,CYP26B1 |
| Catecholamine Biosynthesis | 1.78 | 0.25 | #NUM! | DBH |
| Methionine Salvage II (Mammalian) | 1.78 | 0.25 | #NUM! | BHMT2 |
| DNA Damage Bypass | 1.77 | 0.0417 | #NUM! | RPA3,UBA52 |
| iNOS Signaling | 1.76 | 0.0408 | #NUM! | FOS,HMGA1 |
| GABA Receptor Signaling | 1.75 | 0.0226 | #NUM! | GABRA6,GABRD,UBA52 |
| Signaling by FGFR1 | 1.71 | 0.0385 | #NUM! | FRS3,UBA52 |
| Adipogenesis pathway | 1.71 | 0.0217 | #NUM! | BMP4,CEBPB,FZD7 |
| Ephrin A Signaling | 1.69 | 0.0214 | #NUM! | ACP1,ARHGEF5,MYL9 |
| Formation of lateral plate mesoderm | 1.69 | 0.2 | #NUM! | BMP4 |
| Bone Mineralization Signaling Pathway | 1.66 | 0.0208 | #NUM! | FZD7,NR4A2,SPP1 |
| DNA Damage/Telomere Stress Induced Senescence | 1.65 | 0.0357 | #NUM! | HMGA1,HMGA2 |
| Cell Cycle Control of Chromosomal Replication | 1.63 | 0.0351 | #NUM! | RPA3,TOP2A |

| | | | | |
|---|---|---|---|---|
| NAD Signaling Pathway | 1.6 | 0.0199 | #NUM! | CEBPB,POLR2K,POLR2L |
| Signaling by Rho Family GTPases | 1.6 | 0.015 | 1 | ARHGEF5,CDC42EP5,FOS,MYL9 |
| Regulation of CDH19 Expression and Function | 1.54 | 0.143 | #NUM! | CTNND1 |
| Transcriptional Regulatory Network in Embryonic Stem Cells | 1.51 | 0.0183 | #NUM! | BMP4,FZD7,PAX6 |
| Formation of intermediate mesoderm | 1.49 | 0.125 | #NUM! | BMP4 |
| WNT/β-catenin Signaling | 1.45 | 0.0173 | #NUM! | FZD7,SFRP5,UBA52 |
| Androgen Signaling | 1.44 | 0.0171 | #NUM! | POLR2K,POLR2L,TAF7L |
| Cellular response to mitochondrial stress | 1.44 | 0.111 | #NUM! | EIF2S3 |
| Erythropoietin Signaling Pathway | 1.4 | 0.0166 | #NUM! | FOS,HBA2,Hbb-bs/Hbb-bt |
| GDNF Family Ligand-Receptor Interactions | 1.4 | 0.0263 | #NUM! | FOS,GFRA4 |
| Toll-like Receptor Signaling | 1.39 | 0.026 | #NUM! | FOS,UBA52 |
| Formation of the posterior neural plate | 1.39 | 0.1 | #NUM! | OTX2 |
| VDR/RXR Activation | 1.38 | 0.0256 | #NUM! | CEBPB,SPP1 |
| MicroRNA Biogenesis Signaling Pathway | 1.37 | 0.016 | #NUM! | BMP4,POLR2K,POLR2L |
| IL-17A Signaling in Fibroblasts | 1.36 | 0.025 | #NUM! | CEBPB,FOS |
| Role of JAK family kinases in IL-6-type Cytokine Signaling | 1.35 | 0.0247 | #NUM! | FOS,VIP |
| Metallothioneins bind metals | 1.35 | 0.0909 | #NUM! | MT2A |
| Alpha-protein kinase 1 signaling pathway | 1.35 | 0.0909 | #NUM! | UBA52 |
| Pexophagy | 1.35 | 0.0909 | #NUM! | UBA52 |
| NAD Phosphorylation and Dephosphorylation | 1.35 | 0.0909 | #NUM! | ACP1 |
| Glycine Betaine Degradation | 1.35 | 0.0909 | #NUM! | BHMT2 |
| RNA polymerase II transcribes snRNA genes | 1.33 | 0.0241 | #NUM! | POLR2K,POLR2L |
| JAK/STAT Signaling | 1.33 | 0.0241 | #NUM! | CEBPB,FOS |

| | | | | |
|---|---|---|---|---|
| Transport of bile salts and organic acids, metal ions and amine compounds | 1.32 | 0.0238 | #NUM! | SLC13A1, SLC6A2 |
| Specification of primordial germ cells | 1.31 | 0.0833 | #NUM! | BMP4 |
| Assembly of RNA Polymerase I Complex | 1.31 | 0.0833 | #NUM! | TAF1D |
| CDX Gastrointestinal Cancer Signaling Pathway | 1.31 | 0.0152 | #NUM! | BMP4,FOS,FZD7 |
| Adrenergic Receptor Signaling Pathway (Enhanced) | 1.31 | 0.0152 | #NUM! | Atp6ap1l, DBH,SLC6A2 |
| PDGF Signaling | 1.3 | 0.023 | #NUM! | ACP1,FOS |
| ABRA Signaling Pathway | 1.29 | 0.0227 | #NUM! | FOS,MYL9 |
| Oxidative Stress Induced Senescence | 1.29 | 0.0227 | #NUM! | FOS,UBA52 |
| Adrenomedullin signaling pathway | 1.29 | 0.0149 | #NUM! | CEBPB,FOS,TTN |
| Platelet Aggregation (Plug Formation) | 1.28 | 0.0769 | #NUM! | F2 |
| Sheddase Signaling Pathway | 1.28 | 0.0147 | #NUM! | CEBPB,FOS,KLK12 |
| BMP signaling pathway | 1.27 | 0.0222 | #NUM! | BMP4,CHRD |
| Ceramide Signaling | 1.26 | 0.022 | #NUM! | FOS,S1PR1 |
| MAPK6/MAPK4 signaling | 1.25 | 0.0217 | #NUM! | CDC42EP5,UBA52 |
| Activin Inhibin Signaling Pathway | 1.24 | 0.0142 | #NUM! | CEBPB,FOS,PAX6 |
| Class B/2 (Secretin family receptors) | 1.23 | 0.0211 | #NUM! | FZD7,VIP |
| Transcriptional regulation by RUNX3 | 1.22 | 0.0208 | #NUM! | SPP1,UBA52 |
| Prolactin Signaling | 1.22 | 0.0208 | #NUM! | CEBPB,FOS |
| Mitochondrial translation | 1.21 | 0.0206 | #NUM! | MRPL53, TSFM |
| TGF-β Signaling | 1.21 | 0.0206 | #NUM! | BMP4,FOS |
| Interferon gamma signaling | 1.21 | 0.0204 | #NUM! | MT2A,TRIM14 |
| Mismatch Repair | 1.19 | 0.0625 | #NUM! | RPA3 |
| Extrinsic Prothrombin Activation Pathway | 1.19 | 0.0625 | #NUM! | F2 |
| Thrombin Signaling | 1.18 | 0.0135 | #NUM! | ARHGEF5,F2,MYL9 |

| | | | | |
|---|---|---|---|---|
| Gene Silencing by RNA | 1.17 | 0.0196 | #NUM! | POLR2K, POLR2L |
| ABC-family proteins mediated transport | 1.17 | 0.0194 | #NUM! | EIF2S3,UBA52 |
| Mouse Embryonic Stem Cell Pluripotency | 1.17 | 0.0194 | #NUM! | BMP4,FZD7 |
| Germ layer formation at gastrulation | 1.17 | 0.0588 | #NUM! | BMP4 |
| Specification of the neural plate border | 1.17 | 0.0588 | #NUM! | BMP4 |
| Sleep REM Signaling Pathway | 1.15 | 0.0189 | #NUM! | FOS,NR4A2 |
| Metabolism of amine-derived hormones | 1.14 | 0.0556 | #NUM! | DBH |
| Formation of the nephric duct | 1.14 | 0.0556 | #NUM! | BMP4 |
| Post-translational protein phosphorylation | 1.14 | 0.0187 | #NUM! | BMP4,SPP1 |
| G-Protein Coupled Receptor Signaling | 1.12 | 0.00842 | -0.816 | FOS,FZD7,GPRC5C,MYL9,S1PR1,TTN |
| Unfolded Protein Response (UPR) | 1.12 | 0.0526 | #NUM! | EIF2S3 |
| RHO GTPases activate CIT | 1.12 | 0.0526 | #NUM! | MYL9 |
| RHO GTPases Activate ROCKs | 1.12 | 0.0526 | #NUM! | MYL9 |
| Regulation of TP53 Activity through Methylation | 1.12 | 0.0526 | #NUM! | UBA52 |
| Interleukin-4 and Interleukin-13 signaling | 1.11 | 0.018 | #NUM! | FOS,S1PR1 |
| Actin Cytoskeleton Signaling | 1.1 | 0.0124 | #NUM! | F2,MYL9,TTN |
| HDR through Homologous Recombination (HRR) or Single Strand Annealing (SSA) | 1.09 | 0.0175 | #NUM! | RPA3,UBA52 |
| RHO GTPases activate PAKs | 1.08 | 0.0476 | #NUM! | MYL9 |
| Formation of the ureteric bud | 1.08 | 0.0476 | #NUM! | BMP4 |
| Sleep NREM Signaling Pathway | 1.08 | 0.0172 | #NUM! | GABRA6, GABRD |
| Wound Healing Signaling Pathway | 1.07 | 0.012 | #NUM! | CEBPB,F2,FOS |
| Cyclophilin Signaling Pathway | 1.07 | 0.012 | #NUM! | F2,SPP1,TOMM7 |
| Cholecystokinin/Gastrin-mediated Signaling | 1.07 | 0.0171 | #NUM! | FOS,SST |
| Insertion of tail-anchored proteins into the endoplasmic reticulum membrane | 1.06 | 0.0455 | #NUM! | OTOF |
| Chaperone Mediated Autophagy | 1.06 | 0.0455 | #NUM! | UBA52 |

| | | | | |
|---|---|---|---|---|
| Formation of paraxial mesoderm | 1.06 | 0.0455 | #NUM! | BMP4 |
| MSP-RON Signaling in Macrophages Pathway | 1.05 | 0.0167 | #NUM! | FOS,KLK12 |
| Synthesis of DNA | 1.04 | 0.0164 | #NUM! | RPA3,UBA52 |
| NAD Salvage Pathway II | 1.04 | 0.0435 | #NUM! | ACP1 |
| G alpha (i) signalling events | 1.04 | 0.0116 | #NUM! | GAL,PCP2,SST |
| Role of NANOG in Mammalian Embryonic Stem Cell Pluripotency | 1.04 | 0.0163 | #NUM! | BMP4,FZD7 |
| Hematoma Resolution Signaling Pathway | 1.03 | 0.0116 | #NUM! | F2,FOS,S1PR1 |
| Other interleukin signaling | 1.02 | 0.0417 | #NUM! | IL16 |
| RAS processing | 1.02 | 0.0417 | #NUM! | UBA52 |
| RAF/MAP kinase cascade | 1.02 | 0.0115 | #NUM! | FRS3,GFRA4,UBA52 |
| Cohesin Chromatin Regulation Pathway | 1.02 | 0.0114 | #NUM! | FAT2,POLR2K,POLR2L |
| MyD88 cascade initiated on plasma membrane | 1.01 | 0.04 | #NUM! | UBA52 |
| Glycogen metabolism | 1.01 | 0.04 | #NUM! | UBA52 |
| Signaling by NTRK2 (TRKB) | 1.01 | 0.04 | #NUM! | FRS3 |
| IL-6 Signaling | 1 | 0.0155 | #NUM! | CEBPB,FOS |
| Cardiomyocyte Differentiation via BMP Receptors | 0.991 | 0.0385 | #NUM! | BMP4 |
| IL-17A Signaling in Gastric Cells | 0.991 | 0.0385 | #NUM! | FOS |
| Pregnenolone Biosynthesis | 0.991 | 0.0385 | #NUM! | CYP26B1 |
| Mitotic G1 phase and G1/S transition | 0.984 | 0.0152 | #NUM! | TOP2A,UBA52 |
| RAR Activation | 0.978 | 0.00924 | -1 | CRABP1,FOS,HOXA5,PAX6 |
| ATF4 activates genes in response to endoplasmic reticulum stress | 0.975 | 0.037 | #NUM! | CEBPB |
| Histidine Degradation VI | 0.975 | 0.037 | #NUM! | CYP26B1 |
| Gα12/13 Signaling | 0.973 | 0.0149 | #NUM! | F2,MYL9 |
| Insulin Secretion Signaling Pathway | 0.961 | 0.0108 | #NUM! | EIF2S3,NEUROD1,SPCS3 |
| Sulfur amino acid metabolism | 0.96 | 0.0357 | #NUM! | BHMT2 |
| RHO GTPases activate PKNs | 0.96 | 0.0357 | #NUM! | MYL9 |

| | | | | |
|---|---|---|---|---|
| CDP-diacylglycerol Biosynthesis I | 0.96 | 0.0357 | #NUM! | LPCAT4 |
| CREB Signaling in Neurons | 0.949 | 0.00812 | #NUM! | FZD7,GPRC5C,POLR2K,POLR2L,S1PR1 |
| SUMOylation of intracellular receptors | 0.946 | 0.0345 | #NUM! | NR4A2 |
| SIRT1 negatively regulates rRNA expression | 0.946 | 0.0345 | #NUM! | TAF1D |
| Processing of Capped Intronless Pre-mRNA | 0.946 | 0.0345 | #NUM! | SNRPF |
| GABAergic Receptor Signaling Pathway (Enhanced) | 0.942 | 0.0143 | #NUM! | GABRA6,GABRD |
| RHO GTPase cycle | 0.933 | 0.00889 | 0 | ARHGEF5,CDC42EP5,FRS3,KCTD13 |
| Signaling by CSF3 (G-CSF) | 0.932 | 0.0333 | #NUM! | UBA52 |
| Regulation of CDH11 Expression and Function | 0.932 | 0.0333 | #NUM! | CTNND1 |
| Glutathione Redox Reactions I | 0.932 | 0.0333 | #NUM! | Gstp1 (includes others) |
| Phosphatidylglycerol Biosynthesis II (Non-plastidic) | 0.932 | 0.0333 | #NUM! | LPCAT4 |
| SRP-dependent cotranslational protein targeting to membrane | 0.932 | 0.0141 | #NUM! | SPCS3,UBA52 |
| MSP-RON Signaling in Cancer Cells Pathway | 0.927 | 0.014 | #NUM! | FOS,KLK12 |
| Endosomal Sorting Complex Required For Transport (ESCRT) | 0.919 | 0.0323 | #NUM! | UBA52 |
| Signaling by CSF1 (M-CSF) in myeloid cells | 0.919 | 0.0323 | #NUM! | UBA52 |
| MyD88 dependent cascade initiated on endosome | 0.919 | 0.0323 | #NUM! | UBA52 |
| Ubiquinol-10 Biosynthesis (Eukaryotic) | 0.919 | 0.0323 | #NUM! | CYP26B1 |
| Thrombin signalling through proteinase activated receptors (PARs) | 0.906 | 0.0312 | #NUM! | F2 |
| RIPK1-mediated regulated necrosis | 0.906 | 0.0312 | #NUM! | UBA52 |
| Chromatin modifications during the maternal to zygotic transition (MZT) | 0.906 | 0.0312 | #NUM! | KDM6A |
| Toll Like Receptor 3 (TLR3) Cascade | 0.893 | 0.0303 | #NUM! | UBA52 |

| | | | | |
|---|---|---|---|---|
| MAPK targets/ Nuclear events mediated by MAP kinases | 0.893 | 0.0303 | #NUM! | FOS |
| TNFR2 Signaling | 0.893 | 0.0303 | #NUM! | FOS |
| Eukaryotic Translation Initiation Factors Promoting | 0.893 | 0.0133 | #NUM! | EIF2S3,UBA52 |
| Cardiogenesis in Vertebrates | 0.888 | 0.0132 | #NUM! | BMP4,FZD7 |
| Signaling by NOTCH2 | 0.881 | 0.0294 | #NUM! | UBA52 |
| Late endosomal microautophagy | 0.881 | 0.0294 | #NUM! | UBA52 |
| Oncogene Induced Senescence | 0.869 | 0.0286 | #NUM! | UBA52 |
| Activation of the pre-replicative complex | 0.869 | 0.0286 | #NUM! | RPA3 |
| Transcriptional Regulation by NPAS4 | 0.869 | 0.0286 | #NUM! | FOS |
| Coagulation System | 0.869 | 0.0286 | #NUM! | F2 |
| Striated Muscle Contraction | 0.858 | 0.0278 | #NUM! | TTN |
| p75 NTR receptor-mediated signalling | 0.857 | 0.0127 | #NUM! | ARHGEF5,UBA52 |
| Superpathway of Methionine Degradation | 0.847 | 0.027 | #NUM! | BHMT2 |
| MyD88-independent TLR4 cascade | 0.836 | 0.0263 | #NUM! | UBA52 |
| Regulation of TP53 Expression and Degradation | 0.836 | 0.0263 | #NUM! | UBA52 |
| Resolution of Abasic Sites (AP sites) | 0.836 | 0.0263 | #NUM! | RPA3 |
| Formation of Fibrin Clot (Clotting Cascade) | 0.826 | 0.0256 | #NUM! | F2 |
| MAP kinase activation | 0.826 | 0.0256 | #NUM! | UBA52 |
| NGF-stimulated transcription | 0.826 | 0.0256 | #NUM! | FOS |
| FLT3 Signaling | 0.826 | 0.0256 | #NUM! | UBA52 |
| Transcriptional Regulation by VENTX | 0.816 | 0.025 | #NUM! | CEBPB |
| CXCR4 Signaling | 0.814 | 0.0119 | #NUM! | FOS,MYL9 |
| Myelination Signaling Pathway | 0.81 | 0.00915 | #NUM! | BMP4,FOS,FZD7 |
| RET signaling | 0.806 | 0.0244 | #NUM! | GFRA4 |
| Neuroinflammation Signaling Pathway | 0.805 | 0.00909 | #NUM! | FOS,GABRA6,GABRD |
| Phagosome Maturation | 0.802 | 0.0117 | #NUM! | Atp6ap1l, Dynlt1b (includes others) |
| MyD88:MAL(TIRAP) cascade initiated on plasma membrane | 0.796 | 0.0238 | #NUM! | UBA52 |
| Formation of WDR5-containing histone-modifying complexes | 0.796 | 0.0238 | #NUM! | KDM6A |

| | | | | |
|---|---|---|---|---|
| April Mediated Signaling | 0.796 | 0.0238 | #NUM! | FOS |
| Smooth Muscle Contraction | 0.787 | 0.0233 | #NUM! | MYL9 |
| B Cell Activating Factor Signaling | 0.787 | 0.0233 | #NUM! | FOS |
| Oncostatin M Signaling | 0.787 | 0.0233 | #NUM! | MT2A |
| Elastic fibre formation | 0.778 | 0.0227 | #NUM! | BMP4 |
| TAK1-dependent IKK and NF-kappa-B activation | 0.778 | 0.0227 | #NUM! | UBA52 |
| Aggrephagy | 0.778 | 0.0227 | #NUM! | UBA52 |
| Phagosome Formation | 0.773 | 0.00708 | -0.447 | FZD7,GPRC5C,MYL9,S1PR1,TTN |
| Tumor Microenvironment Pathway | 0.767 | 0.0111 | #NUM! | FOS,SPP1 |
| Tight Junction Signaling | 0.764 | 0.011 | #NUM! | FOS,MYL9 |
| SUMOylation of DNA replication proteins | 0.76 | 0.0217 | #NUM! | TOP2A |
| MIF Regulation of Innate Immunity | 0.76 | 0.0217 | #NUM! | FOS |
| Role of OCT4 in Mammalian Embryonic Stem Cell Pluripotency | 0.76 | 0.0217 | #NUM! | SPP1 |
| Aryl Hydrocarbon Receptor Signaling | 0.756 | 0.0109 | #NUM! | FOS,Gstp1 (includes others) |
| Pyruvate metabolism | 0.735 | 0.0204 | #NUM! | UBA52 |
| IL-17 Signaling | 0.735 | 0.0106 | #NUM! | CEBPB,FOS |
| Signaling by ERBB2 | 0.728 | 0.02 | #NUM! | UBA52 |
| Signaling by NOTCH3 | 0.728 | 0.02 | #NUM! | UBA52 |
| Meiotic recombination | 0.728 | 0.02 | #NUM! | RPA3 |
| Macrophage Alternative Activation Signaling Pathway | 0.724 | 0.0104 | #NUM! | CEBPB,FOS |
| Transcriptional activity of SMAD2/SMAD3:SMAD4 heterotrimer | 0.72 | 0.0196 | #NUM! | UBA52 |
| Transcriptional regulation of granulopoiesis | 0.72 | 0.0196 | #NUM! | CEBPB |
| Glutathione-mediated Detoxification | 0.72 | 0.0196 | #NUM! | Gstp1 (includes others) |
| UVC-Induced MAPK Signaling | 0.72 | 0.0196 | #NUM! | FOS |
| Cell Cycle: G2/M DNA Damage Checkpoint Regulation | 0.72 | 0.0196 | #NUM! | TOP2A |
| Regulation of the Epithelial-Mesenchymal Transition Pathway | 0.718 | 0.0103 | #NUM! | FZD7,HMGA2 |

| | | | | |
|---|---|---|---|---|
| Regulation of the Epithelial Mesenchymal Transition by Growth Factors Pathway | 0.718 | 0.0103 | #NUM! | FOS,HMGA2 |
| TNFR1 Signaling | 0.712 | 0.0192 | #NUM! | FOS |
| Cachexia Signaling Pathway | 0.709 | 0.00815 | #NUM! | CEBPB,EIF2S3,S1PR1 |
| ILK Signaling | 0.708 | 0.0102 | #NUM! | FOS,MYL9 |
| Regulation of Apoptosis | 0.705 | 0.0189 | #NUM! | UBA52 |
| Signaling by EGFR | 0.705 | 0.0189 | #NUM! | UBA52 |
| Human Embryonic Stem Cell Pluripotency | 0.698 | 0.01 | #NUM! | BMP4,FZD7 |
| Signaling by TGF-beta Receptor Complex | 0.698 | 0.0185 | #NUM! | UBA52 |
| UVB-Induced MAPK Signaling | 0.698 | 0.0185 | #NUM! | FOS |
| Pulmonary Healing Signaling Pathway | 0.694 | 0.00995 | #NUM! | BMP4,FZD7 |
| Signaling by PTK6 | 0.69 | 0.0182 | #NUM! | UBA52 |
| CSDE1 Signaling Pathway | 0.683 | 0.0179 | #NUM! | FOS |
| NLR signaling pathways | 0.683 | 0.0179 | #NUM! | UBA52 |
| Metabolism of non-coding RNA | 0.683 | 0.0179 | #NUM! | SNRPF |
| Deadenylation-dependent mRNA decay | 0.683 | 0.0179 | #NUM! | LSM5 |
| Interleukin-3, Interleukin-5 and GM-CSF signaling | 0.683 | 0.0179 | #NUM! | UBA52 |
| E3 ubiquitin ligases ubiquitinate target proteins | 0.683 | 0.0179 | #NUM! | UBA52 |
| CD27 Signaling in Lymphocytes | 0.683 | 0.0179 | #NUM! | FOS |
| EGF Signaling | 0.683 | 0.0179 | #NUM! | FOS |
| Beta-catenin independent WNT signaling | 0.682 | 0.00976 | #NUM! | FZD7,UBA52 |
| Fc epsilon receptor (FCERI) signaling | 0.679 | 0.00971 | #NUM! | FOS,UBA52 |
| Neurexins and neuroligins | 0.677 | 0.0175 | #NUM! | APBA3 |
| TNF signaling | 0.677 | 0.0175 | #NUM! | UBA52 |
| IL-8 Signaling | 0.672 | 0.00962 | #NUM! | FOS,MYL9 |
| Role of Tissue Factor in Cancer | 0.672 | 0.00962 | #NUM! | F2,FOS |
| Circadian Clock | 0.67 | 0.0172 | #NUM! | UBA52 |
| Signaling by PDGF | 0.67 | 0.0172 | #NUM! | SPP1 |
| RUNX1 regulates megakaryocyte differentiation and platelet function | 0.67 | 0.0172 | #NUM! | MYL9 |
| Triacylglycerol Biosynthesis | 0.67 | 0.0172 | #NUM! | LPCAT4 |
| Clathrin-mediated Endocytosis Signaling | 0.669 | 0.00957 | #NUM! | F2,UBA52 |
| Integrin Signaling | 0.666 | 0.00952 | #NUM! | MYL9,TTN |

| | | | | |
|---|---|---|---|---|
| Signaling by ERBB4 | 0.663 | 0.0169 | #NUM! | UBA52 |
| DNA Double Strand Break Response | 0.663 | 0.0169 | #NUM! | UBA52 |
| Gustation Pathway | 0.663 | 0.00948 | #NUM! | GABRA6, GABRD |
| NIK-->noncanonical NF-kB signaling | 0.657 | 0.0167 | #NUM! | UBA52 |
| Iron uptake and transport | 0.657 | 0.0167 | #NUM! | UBA52 |
| GABA receptor activation | 0.657 | 0.0167 | #NUM! | GABRA6 |
| PCP (Planar Cell Polarity) Pathway | 0.657 | 0.0167 | #NUM! | FZD7 |
| S100 Family Signaling Pathway | 0.652 | 0.00639 | -1.342 | AHNAK,FOS,FZD7,GPRC5C,S1PR1 |
| Gamma carboxylation, hypusinylation, hydroxylation, and arylsulfatase activation | 0.651 | 0.0164 | #NUM! | F2 |
| Retinoic acid Mediated Apoptosis Signaling | 0.651 | 0.0164 | #NUM! | CRABP1 |
| Protein Kinase A Signaling | 0.647 | 0.00758 | #NUM! | ACP1,MYL9,TTN |
| MSP-RON Signaling Pathway | 0.644 | 0.0161 | #NUM! | KLK12 |
| NCAM signaling for neurite out-growth | 0.638 | 0.0159 | #NUM! | GFRA4 |
| Transcriptional Regulation by MECP2 | 0.638 | 0.0159 | #NUM! | SST |
| Triacylglycerol Degradation | 0.638 | 0.0159 | #NUM! | AARSD1 |
| Stearate Biosynthesis I (Animals) | 0.638 | 0.0159 | #NUM! | LPCAT4 |
| IL-2 Signaling | 0.638 | 0.0159 | #NUM! | FOS |
| RHOGDI Signaling | 0.637 | 0.00909 | #NUM! | ARHGEF5,MYL9 |
| Semaphorin interactions | 0.632 | 0.0156 | #NUM! | MYL9 |
| Peroxisomal protein import | 0.632 | 0.0156 | #NUM! | UBA52 |
| Thrombopoietin Signaling | 0.632 | 0.0156 | #NUM! | FOS |
| Mitochondrial protein import | 0.626 | 0.0154 | #NUM! | TOMM7 |
| Hedgehog ligand biogenesis | 0.626 | 0.0154 | #NUM! | UBA52 |
| Role of JAK2 in Hormone-like Cytokine Signaling | 0.626 | 0.0154 | #NUM! | FOS |
| Glucocorticoid Receptor Signaling | 0.626 | 0.00667 | #NUM! | FOS,POLR2K,POLR2L,TAF7L |
| Glycation Signaling Pathway | 0.623 | 0.00889 | #NUM! | FOS,HBA2 |
| WNT/Ca+ pathway | 0.621 | 0.0152 | #NUM! | FZD7 |
| CD40 Signaling | 0.615 | 0.0149 | #NUM! | FOS |
| TNFR2 non-canonical NF-kB pathway | 0.609 | 0.0147 | #NUM! | UBA52 |

| | | | | |
|---|---|---|---|---|
| Sensory processing of sound by inner hair cells of the cochlea | 0.604 | 0.0145 | #NUM! | OTOF |
| Neurovascular Coupling Signaling Pathway | 0.603 | 0.00862 | #NUM! | GABRA6, GABRD |
| Remodeling of Epithelial Adherens Junctions | 0.593 | 0.0141 | #NUM! | CTNND1 |
| Sertoli Cell-Germ Cell Junction Signaling Pathway (Enhanced) | 0.593 | 0.00847 | #NUM! | CTNND1, FOS |
| Regulation of RUNX2 expression and activity | 0.588 | 0.0139 | #NUM! | UBA52 |
| EIF2 Signaling | 0.588 | 0.0084 | #NUM! | EIF2S3,UBA52 |
| Growth Hormone Signaling | 0.583 | 0.0137 | #NUM! | FOS |
| Ephrin B Signaling | 0.583 | 0.0137 | #NUM! | ACP1 |
| ISG15 antiviral mechanism | 0.578 | 0.0135 | #NUM! | UBA52 |
| ERK5 Signaling | 0.578 | 0.0135 | #NUM! | FOS |
| IL-12 Signaling and Production in Macrophages | 0.578 | 0.00826 | #NUM! | CEBPB,FOS |
| Pancreatic Secretion Signaling Pathway | 0.575 | 0.00823 | #NUM! | ARHGEF5,VIP |
| Extra-nuclear estrogen signaling | 0.573 | 0.0133 | #NUM! | FOS |
| Cellular response to hypoxia | 0.568 | 0.0132 | #NUM! | UBA52 |
| Plasma lipoprotein assembly, remodeling, and clearance | 0.568 | 0.0132 | #NUM! | UBA52 |
| Interferon alpha/beta signaling | 0.568 | 0.0132 | #NUM! | UBA52 |
| Signaling by NOTCH1 | 0.563 | 0.013 | #NUM! | UBA52 |
| Antiproliferative Role of Somatostatin Receptor 2 | 0.563 | 0.013 | #NUM! | SST |
| PKR-mediated signaling | 0.558 | 0.0128 | #NUM! | EIF2S3 |
| Neurotrophin/TRK Signaling | 0.558 | 0.0128 | #NUM! | FOS |
| Hypoxia Signaling in the Cardiovascular System | 0.558 | 0.0128 | #NUM! | UBE2T |
| Signaling by MET | 0.554 | 0.0127 | #NUM! | UBA52 |
| G alpha (12/13) signalling events | 0.549 | 0.0125 | #NUM! | ARHGEF5 |
| Degradation of the extracellular matrix | 0.544 | 0.0123 | #NUM! | SPP1 |
| DDX58/IFIH1-mediated induction of interferon-alpha/beta | 0.544 | 0.0123 | #NUM! | UBA52 |
| IL-3 Signaling | 0.544 | 0.0123 | #NUM! | FOS |
| Chemokine Signaling | 0.544 | 0.0123 | #NUM! | FOS |
| Transcriptional regulation of white adipocyte differentiation | 0.531 | 0.0119 | #NUM! | CEBPB |
| Integrin cell surface interactions | 0.527 | 0.0118 | #NUM! | SPP1 |
| LPS-stimulated MAPK Signaling | 0.527 | 0.0118 | #NUM! | FOS |

| | | | | |
|---|---|---|---|---|
| VEGF Family Ligand-Receptor Interactions | 0.523 | 0.0116 | #NUM! | FOS |
| Regulation of the Epithelial Mesenchymal Transition in Development Pathway | 0.523 | 0.0116 | #NUM! | FZD7 |
| Regulation of mitotic cell cycle | 0.514 | 0.0114 | #NUM! | UBA52 |
| Amyloid fiber formation | 0.514 | 0.0114 | #NUM! | UBA52 |
| NRF2-mediated Oxidative Stress Response | 0.512 | 0.00741 | #NUM! | FOS,Gstp1 (includes others) |
| Regulation of mRNA stability by proteins that bind AU-rich elements | 0.51 | 0.0112 | #NUM! | UBA52 |
| Cell Cycle Checkpoints | 0.508 | 0.00735 | #NUM! | RPA3,UBA52 |
| Degradation of beta-catenin by the destruction complex | 0.502 | 0.011 | #NUM! | UBA52 |
| Cell junction organization | 0.498 | 0.0109 | #NUM! | CTNND1 |
| Unfolded protein response | 0.498 | 0.0109 | #NUM! | CEBPB |
| EPH-Ephrin signaling | 0.495 | 0.0108 | #NUM! | MYL9 |
| ERBB Signaling | 0.495 | 0.0108 | #NUM! | FOS |
| Protein Ubiquitination Pathway | 0.494 | 0.00717 | #NUM! | UBA52,UBE2T |
| Heparan Sulfate Biosynthesis (Late Stages) | 0.491 | 0.0106 | #NUM! | AARSD1 |
| Apelin Adipocyte Signaling Pathway | 0.491 | 0.0106 | #NUM! | Gstp1 (includes others) |
| Oxytocin Signaling Pathway | 0.481 | 0.00702 | #NUM! | FOS,MYL9 |
| VEGF Signaling | 0.48 | 0.0103 | #NUM! | EIF2S3 |
| BBSome Signaling Pathway | 0.476 | 0.00609 | #NUM! | FZD7,GPRC5C,S1PR1 |
| DNA Methylation and Transcriptional Repression Signaling | 0.476 | 0.0102 | #NUM! | CEBPB |
| IL-1 Signaling | 0.476 | 0.0102 | #NUM! | FOS |
| UVA-Induced MAPK Signaling | 0.476 | 0.0102 | #NUM! | FOS |
| Protein folding | 0.472 | 0.0101 | #NUM! | PFDN4 |
| S Phase | 0.469 | 0.01 | #NUM! | UBA52 |
| Apelin Cardiomyocyte Signaling Pathway | 0.469 | 0.01 | #NUM! | MYL9 |
| Cellular response to heat stress | 0.465 | 0.0099 | #NUM! | RPA3 |
| Heparan Sulfate Biosynthesis | 0.465 | 0.0099 | #NUM! | AARSD1 |
| Sumoylation Pathway | 0.465 | 0.0099 | #NUM! | FOS |
| DNA Replication Pre-Initiation | 0.455 | 0.00962 | #NUM! | UBA52 |
| Cargo recognition for clathrin-mediated endocytosis | 0.452 | 0.00952 | #NUM! | UBA52 |
| IGF-1 Signaling | 0.452 | 0.00952 | #NUM! | FOS |

| | | | | |
|---|---|---|---|---|
| Signaling by VEGF | 0.445 | 0.00935 | #NUM! | CTNND1 |
| Axonal Guidance Signaling | 0.443 | 0.0058 | #NUM! | BMP4,FZD7,MYL9 |
| PPAR Signaling | 0.442 | 0.00926 | #NUM! | FOS |
| Xenobiotic Metabolism AHR Signaling Pathway | 0.439 | 0.00917 | #NUM! | Gstp1 (includes others) |
| Lung Ionic Balance Signaling Pathway | 0.436 | 0.00575 | #NUM! | FZD7,GPRC5C,S1PR1 |
| Regulation of Actin-based Motility by Rho | 0.433 | 0.00901 | #NUM! | MYL9 |
| Phase I - Functionalization of compounds | 0.427 | 0.00885 | #NUM! | CYP26B1 |
| Hedgehog 'off' state | 0.424 | 0.00877 | #NUM! | UBA52 |
| Nonsense-Mediated Decay (NMD) | 0.415 | 0.00855 | #NUM! | UBA52 |
| PAK Signaling | 0.412 | 0.00847 | #NUM! | MYL9 |
| Regulation of lipid metabolism by PPARalpha | 0.409 | 0.0084 | #NUM! | GLIPR1 |
| Sphingosine-1-phosphate Signaling | 0.409 | 0.0084 | #NUM! | S1PR1 |
| Glutaminergic Receptor Signaling Pathway (Enhanced) | 0.408 | 0.00613 | #NUM! | GABRA6, GABRD |
| Eukaryotic Translation Elongation | 0.401 | 0.0082 | #NUM! | UBA52 |
| Eukaryotic Translation Termination | 0.401 | 0.0082 | #NUM! | UBA52 |
| Renin-Angiotensin Signaling | 0.401 | 0.0082 | #NUM! | FOS |
| IL-13 Signaling Pathway | 0.398 | 0.00813 | #NUM! | SERPINE3 |
| Asparagine N-linked glycosylation | 0.393 | 0.008 | #NUM! | UBA52 |
| TCR signaling | 0.39 | 0.00794 | #NUM! | UBA52 |
| Glycerophospholipid biosynthesis | 0.385 | 0.00781 | #NUM! | LPCAT4 |
| Endocannabinoid Developing Neuron Pathway | 0.385 | 0.00781 | #NUM! | PAX6 |
| Interleukin-1 family signaling | 0.382 | 0.00775 | #NUM! | UBA52 |
| Clathrin-mediated endocytosis | 0.382 | 0.00775 | #NUM! | UBA52 |
| Mitochondrial Dysfunction | 0.378 | 0.0058 | #NUM! | Gstp1 (includes others),TOMM7 |
| TR/RXR Activation | 0.377 | 0.00763 | #NUM! | TRH |
| HGF Signaling | 0.377 | 0.00763 | #NUM! | FOS |
| Response to elevated platelet cytosolic Ca2+ | 0.375 | 0.00758 | #NUM! | TTN |
| 14-3-3-mediated Signaling | 0.375 | 0.00758 | #NUM! | FOS |
| P2Y Purinergic Receptor Signaling Pathway | 0.37 | 0.00746 | #NUM! | FOS |

| | | | | |
|---|---|---|---|---|
| CDC42 Signaling | 0.368 | 0.00519 | #NUM! | CDC42EP5,FOS,MYL9 |
| Complement cascade | 0.365 | 0.00735 | #NUM! | F2 |
| SNARE Signaling Pathway | 0.36 | 0.00725 | #NUM! | MYL9 |
| CGAS-STING Signaling Pathway | 0.36 | 0.00725 | #NUM! | Atp6ap1l |
| Role of PKR in Interferon Induction and Antiviral Response | 0.36 | 0.00725 | #NUM! | FOS |
| Reelin Signaling in Neurons | 0.36 | 0.00725 | #NUM! | ARHGEF5 |
| White Adipose Tissue Browning Pathway | 0.358 | 0.00719 | #NUM! | CEBPB |
| Signaling by NOTCH4 | 0.353 | 0.00709 | #NUM! | UBA52 |
| Th2 Pathway | 0.353 | 0.00709 | #NUM! | S1PR1 |
| Hedgehog 'on' state | 0.347 | 0.00694 | #NUM! | UBA52 |
| G alpha (s) signalling events | 0.344 | 0.0069 | #NUM! | VIP |
| C-type lectin receptors (CLRs) | 0.344 | 0.0069 | #NUM! | UBA52 |
| Apelin Endothelial Signaling Pathway | 0.344 | 0.0069 | #NUM! | FOS |
| Selenoamino acid metabolism | 0.342 | 0.00685 | #NUM! | UBA52 |
| Gαi Signaling | 0.34 | 0.0068 | #NUM! | S1PR1 |
| Semaphorin Neuronal Repulsive Signaling Pathway | 0.34 | 0.0068 | #NUM! | MYL9 |
| PTEN Regulation | 0.334 | 0.00667 | #NUM! | UBA52 |
| WNT/SHH Axonal Guidance Signaling Pathway | 0.332 | 0.00662 | #NUM! | FZD7 |
| Transcriptional regulation by RUNX1 | 0.33 | 0.00658 | #NUM! | UBA52 |
| KEAP1-NFE2L2 pathway | 0.33 | 0.00658 | #NUM! | UBA52 |
| IL-10 Signaling | 0.323 | 0.00645 | #NUM! | FOS |
| Corticotropin Releasing Hormone Signaling | 0.323 | 0.00645 | #NUM! | FOS |
| Relaxin Signaling | 0.321 | 0.00641 | #NUM! | FOS |
| Epithelial Adherens Junction Signaling | 0.317 | 0.00633 | #NUM! | CTNND1 |
| HMGB1 Signaling | 0.314 | 0.00625 | #NUM! | FOS |
| HEY1 Signaling Pathway | 0.312 | 0.00621 | #NUM! | BMP4 |
| Necroptosis Signaling Pathway | 0.308 | 0.00613 | #NUM! | TOMM7 |
| Xenobiotic Metabolism General Signaling Pathway | 0.308 | 0.00613 | #NUM! | Gstp1 (includes others) |
| HOTAIR Regulatory Pathway | 0.304 | 0.00606 | #NUM! | SPP1 |
| Mitochondrial Division Signaling Pathway | 0.302 | 0.00602 | #NUM! | S1PR1 |
| Signaling by the B Cell Receptor (BCR) | 0.295 | 0.00588 | #NUM! | UBA52 |
| Estrogen Receptor Signaling | 0.294 | 0.00487 | #NUM! | FOS,MYL9 |

| | | | | |
|---|---|---|---|---|
| Germ Cell-Sertoli Cell Junction Signaling | 0.293 | 0.00585 | #NUM! | CTNND1 |
| Ribonucleotide Reductase Signaling Pathway | 0.288 | 0.00575 | #NUM! | FOS |
| Th1 and Th2 Activation Pathway | 0.285 | 0.00568 | #NUM! | S1PR1 |
| Ion channel transport | 0.273 | 0.00546 | #NUM! | UBA52 |
| Major pathway of rRNA processing in the nucleolus and cytosol | 0.269 | 0.00538 | #NUM! | UBA52 |
| Regulation of eIF4 and p70S6K Signaling | 0.266 | 0.00532 | #NUM! | EIF2S3 |
| D-myo-inositol (1,4,5,6)-Tetrakisphosphate Biosynthesis | 0.264 | 0.00529 | #NUM! | ACP1 |
| D-myo-inositol (3,4,5,6)-tetrakisphosphate Biosynthesis | 0.264 | 0.00529 | #NUM! | ACP1 |
| Production of Nitric Oxide and Reactive Oxygen Species in Macrophages | 0.26 | 0.00521 | #NUM! | FOS |
| NOD1/2 Signaling Pathway | 0.258 | 0.00518 | #NUM! | FOS |
| GNRH Signaling | 0.258 | 0.00518 | #NUM! | FOS |
| Leukocyte Extravasation Signaling | 0.257 | 0.00515 | #NUM! | CTNND1 |
| Endothelin-1 Signaling | 0.257 | 0.00515 | #NUM! | FOS |
| IL-33 Signaling Pathway | 0.255 | 0.00513 | #NUM! | FOS |
| FXR/RXR Activation | 0.255 | 0.00513 | #NUM! | Gstp1 (includes others) |
| TCF dependent signaling in response to WNT | 0.251 | 0.00505 | #NUM! | UBA52 |
| Mitotic G2-G2/M phases | 0.248 | 0.005 | #NUM! | UBA52 |
| 3-phosphoinositide Degradation | 0.248 | 0.005 | #NUM! | ACP1 |
| ID1 Signaling Pathway | 0.245 | 0.00495 | #NUM! | BMP4 |
| Ephrin Receptor Signaling | 0.245 | 0.00495 | #NUM! | ACP1 |
| D-myo-inositol-5-phosphate Metabolism | 0.243 | 0.0049 | #NUM! | ACP1 |
| Cell surface interactions at the vascular wall | 0.23 | 0.00467 | #NUM! | F2 |
| Immunoregulatory interactions between a Lymphoid and a non-Lymphoid cell | 0.228 | 0.00465 | #NUM! | CRTAM |
| 3-phosphoinositide Biosynthesis | 0.228 | 0.00465 | #NUM! | ACP1 |
| Keratinization | 0.227 | 0.00463 | #NUM! | KLK12 |
| Calcium Signaling | 0.225 | 0.00459 | #NUM! | MYL9 |
| Autophagy | 0.223 | 0.00457 | #NUM! | FOS |
| ERK/MAPK Signaling | 0.222 | 0.00455 | #NUM! | FOS |
| Xenobiotic Metabolism PXR Signaling Pathway | 0.221 | 0.00452 | #NUM! | Gstp1 (includes others) |

| | | | | |
|---|---|---|---|---|
| Agranulocyte Adhesion and Diapedesis | 0.217 | 0.00446 | #NUM! | MYL9 |
| Xenobiotic Metabolism CAR Signaling Pathway | 0.207 | 0.00429 | #NUM! | Gstp1 (includes others) |
| Mitotic Metaphase and Anaphase | 0.204 | 0.00424 | #NUM! | UBA52 |
| Superpathway of Inositol Phosphate Compounds | 0.198 | 0.00415 | #NUM! | ACP1 |
| cAMP-mediated signaling | 0.198 | 0.00415 | #NUM! | S1PR1 |
| CLEAR Signaling Pathway | 0 | 0.00351 | #NUM! | Atp6ap1l |
| Neutrophil Extracellular Trap Signaling Pathway | 0 | 0.00244 | #NUM! | TOMM7 |
| Circadian Rhythm Signaling | 0 | 0.0037 | #NUM! | VIP |
| Chaperone Mediated Autophagy Signaling Pathway | 0 | 0.00157 | #NUM! | Atp6ap1l |
| Orexin Signaling Pathway | 0 | 0.00413 | #NUM! | Atp6ap1l |
| LPS/IL-1 Mediated Inhibition of RXR Function | 0 | 0.00336 | #NUM! | Gstp1 (includes others) |
| Generic Transcription Pathway | 0 | 0.00233 | #NUM! | NR4A2 |
| Signaling by ROBO receptors | 0 | 0.00407 | #NUM! | UBA52 |
| Deubiquitination | 0 | 0.00377 | #NUM! | UBA52 |
| Neutrophil degranulation | 0 | 0.0021 | #NUM! | GLIPR1 |
| Neddylation | 0 | 0.00407 | #NUM! | UBA52 |
| Class I MHC mediated antigen processing and presentation | 0 | 0.00262 | #NUM! | UBA52 |
| Histone Modification Signaling Pathway | 0 | 0.00333 | #NUM! | KDM6A |
| Ribosomal Quality Control Signaling Pathway | 0 | 0.00379 | #NUM! | UBA52 |
| Chromatin organization | 0 | 0.00392 | #NUM! | KDM6A |
| TRIM21 Intracellular Antibody Signaling Pathway | 0 | 0.00353 | #NUM! | FOS,UBA52 |
| ID3 Signaling Pathway | 0 | 0.00103 | #NUM! | BMP4 |
| Role of NFAT in Regulation of the Immune Response | 0 | 0.000955 | #NUM! | FOS |
| CCR5 Signaling in Macrophages | 0 | 0.002 | #NUM! | FOS |
| CTLA4 Signaling in Cytotoxic T Lymphocytes | 0 | 0.00164 | #NUM! | FOS |
| CD28 Signaling in T Helper Cells | 0 | 0.00192 | #NUM! | FOS |
| p70S6K Signaling | 0 | 0.00172 | #NUM! | F2 |
| FAK Signaling | 0 | 0.00386 | -2 | FOS,FZD7,GPRC5C,S1PR1 |
| Phospholipase C Signaling | 0 | 0.00267 | #NUM! | AHNAK,ARHGEF5,MYL9 |

| | | | | |
|---|---|---|---|---|
| Regulation of IL-2 Expression in Activated and Anergic T Lymphocytes | 0 | 0.00214 | #NUM! | FOS |
| PKCθ Signaling in T Lymphocytes | 0 | 0.00179 | #NUM! | FOS |
| PI3K Signaling in B Lymphocytes | 0 | 0.00169 | #NUM! | FOS |
| TEC Kinase Signaling | 0 | 0.00173 | #NUM! | FOS |
| Xenobiotic Metabolism Signaling | 0 | 0.00302 | #NUM! | Gstp1 (includes others) |
| Serotonin Receptor Signaling | 0 | 0.00213 | #NUM! | TTN |
| NF-κB Signaling | 0 | 0.00175 | #NUM! | BMP4 |
| T Cell Receptor Signaling | 0 | 0.00161 | #NUM! | FOS |
| Sirtuin Signaling Pathway | 0 | 0.00337 | #NUM! | TOMM7 |
| Opioid Signaling Pathway | 0 | 0.00356 | #NUM! | FOS |
| Cardiac Hypertrophy Signaling (Enhanced) | 0 | 0.00186 | #NUM! | FZD7 |
| T Cell Exhaustion Signaling Pathway | 0 | 0.00176 | #NUM! | FOS |
| Synaptogenesis Signaling Pathway | 0 | 0.00316 | #NUM! | CTNND1 |
| Senescence Pathway | 0 | 0.00332 | #NUM! | CEBPB |
