## Supplemental Table 11 for "Early Life Adversity Produces Enduring Molecular and Functional Disruption of Developing Vagal Circuits"

**P9 EXCITATORY NEURONS FEMALE**

| Category | Term | PValue | Genes | Fold Enrichment | FDR |
|---|---|---|---|---|---|
| KEGG_PATHWAY | mmu04080:Neuroactive ligand-receptor interaction | 1.14783E-07 | 21334, 15564, 12918, 14599, | 10.4583333 | 8.37914E-06 |
| UP_KW_PTM | KW-0027~Amidation | 1.5372E-06 | 21334, 12918, | 55.4124579 | 1.38348E-05 |
| GOTERM_CC_DIRECT | GO:0005576~extracellular region | 2.71716E-06 | 13590, 21334, 14560, | 5.34980748 | 0.00029617 |
| UP_KW_MOLECULAR_FUNCTION | KW-0372~Hormone | 1.05431E-05 | 12918, 14599, | 19.5326954 | 0.000305751 |
| UP_KW_PTM | KW-0165~Cleavage on pair of basic residues | 0.000142805 | 21334, 12918, 14560, | 11.1246303 | 0.000642622 |
| GOTERM_MF_DIRECT | GO:0005179~hormone activity | 5.61821E-05 | 12918, 14599, | 23.574668 | 0.006067665 |
| UP_KW_CELLULAR_COMPONENT | KW-0964~Secreted | 0.000528508 | 13590, 21334, 12918, | 3.26927716 | 0.010041649 |
| GOTERM_MF_DIRECT | GO:1990837~sequence-specific double-stranded DNA binding | 0.00027435 | 18424, 65100, 18012, 110648, | 7.57481135 | 0.014814913 |
| UP_KW_PTM | KW-1015~Disulfide bond | 0.007290033 | 13590, 245827, 15564, | 1.9973127 | 0.021870098 |
| UP_KW_DOMAIN | KW-0371~Homeobox | 0.002117445 | 18424, 110648, | 8.85938802 | 0.025409334 |
| SMART | SM00389:HOX | 0.001671168 | 18424, | 9.08065051 | 0.035094521 |
| SMART | SM00039:CRF | 0.00401661 | 12918, | 475.826087 | 0.042174406 |
| UP_KW_MOLECULAR_FUNCTION | KW-9996~Developmental protein | 0.004456487 | 13590, 18424, 65100, | 3.20238822 | 0.043079374 |
| UP_KW_MOLECULAR_FUNCTION | KW-0217~Developmental protein | 0.004456487 | 13590, 18424, 65100, | 3.20238822 | 0.043079374 |
| UP_KW_MOLECULAR_FUNCTION | KW-0869~Chloride channel | 0.008084655 | 243634, 14399, | 21.4108392 | 0.058613752 |
| INTERPRO | IPR017970:Homeobox_CS | 0.000388053 | 18424, 110648, | 14.139662 | 0.058984053 |
| GOTERM_BP_DIRECT | GO:0090280~positive regulation of calcium ion import | 0.000524051 | 12918, 14526, 22226 | 86.8075802 | 0.091940088 |

| | | | | | |
|---|---|---|---|---|---|
| GOTERM_BP_DIRECT | GO:0032355~response to estradiol | 0.000656712 | 16409, 14599, | 23.148688 | 0.091940088 |
| GOTERM_BP_DIRECT | GO:0007565~female pregnancy | 0.000656712 | 12918, 14599, | 23.148688 | 0.091940088 |
| GOTERM_BP_DIRECT | GO:0007417~central nervous system development | 0.000855257 | 18424, 65100, 18012, | 21.1357587 | 0.091940088 |
| GOTERM_BP_DIRECT | GO:0009952~anterior/posterior pattern specification | 0.001139261 | 18424, 18012, 15416, 15415 | 19.1386791 | 0.097976488 |
| UP_KW_PTM | KW-0325~Glycoprotein | 0.04371183 | 13590, 243634, 245827, | 1.61466765 | 0.098351617 |
| GOTERM_BP_DIRECT | GO:0071542~dopaminergic neuron differentiation | 0.001548103 | 18424, 110648, 13799 | 50.6377551 | 0.105899298 |
| GOTERM_BP_DIRECT | GO:0030901~midbrain development | 0.001723942 | 18424, 110648, | 47.9726101 | 0.105899298 |
| UP_KW_DOMAIN | KW-0732~Signal | 0.017901712 | 13590, 245827, 21334, 14560, | 1.60988163 | 0.107410269 |
| UP_KW_LIGAND | KW-0868~Chloride | 0.01114592 | 243634, | 17.0369748 | 0.111459197 |
| INTERPRO | IPR001356:HD | 0.001584992 | 18424, | 9.69122899 | 0.114647349 |
| INTERPRO | IPR009057:Homeodomain-like_sf | 0.003675092 | 18424, 110648, | 7.67821406 | 0.114647349 |
| INTERPRO | IPR003620:Urocortin_CRF | 0.003771294 | 12918, 22226 | 517.511628 | 0.114647349 |
| INTERPRO | IPR018446:Corticotropin-releasing_fac_CS | 0.003771294 | 12918, 22226 | 517.511628 | 0.114647349 |
| GOTERM_MF_DIRECT | GO:0051430~corticotropin-releasing hormone receptor 1 binding | 0.006579714 | 12918, 22226 | 297.040816 | 0.147807506 |
| GOTERM_MF_DIRECT | GO:0051431~corticotropin-releasing hormone receptor 2 binding | 0.006579714 | 12918, 22226 | 297.040816 | 0.147807506 |
| GOTERM_MF_DIRECT | GO:0005254~chloride channel activity | 0.00684294 | 243634, 14399, | 23.7632653 | 0.147807506 |
| INTERPRO | IPR050948:Antp_homeobox_TF | 0.007528703 | 15416, 15415 | 258.755814 | 0.163480418 |
| INTERPRO | IPR000187:CRF | 0.007528703 | 12918, | 258.755814 | 0.163480418 |
| GOTERM_CC_DIRECT | GO:0034707~chloride channel complex | 0.003161272 | 243634, 14399, | 35.2900943 | 0.1722893 |

| | | | | | |
|---|---|---|---|---|---|
| GOTERM_MF_DIRECT | GO:0000981~DNA-binding transcription factor activity, RNA polymerase II-specific | 0.009608408 | 18424, 65100, 18012, 110648, 15416, 15415, | 3.73636247 | 0.172951351 |
| UP_SEQ_FEATURE | DNA_BIND:Homeobox | 0.001234272 | 18424, 110648, | 10.3613445 | 0.188843614 |
| INTERPRO | IPR020479:HD_metazoa | 0.012806184 | 15416, 15415, | 17.0608229 | 0.24331749 |
| GOTERM_CC_DIRECT | GO:0045202~synapse | 0.007184439 | 15564, 12918, | 4.82677419 | 0.261034615 |
| GOTERM_BP_DIRECT | GO:0021549~cerebellum development | 0.004961449 | 14560, 18012, | 28.0455259 | 0.266677904 |
| GOTERM_CC_DIRECT | GO:0005615~extracellular space | 0.011876159 | 13590, 21334, | 2.80696598 | 0.323625345 |
| GOTERM_MF_DIRECT | GO:0051378~serotonin binding | 0.024456124 | 15564, 15567 | 79.2108844 | 0.348318816 |
| GOTERM_MF_DIRECT | GO:0008083~growth factor activity | 0.025801394 | 13590, 14560, | 11.8029463 | 0.348318816 |
| SMART | SM00110:C1Q | 0.058657209 | 23829, | 31.7217391 | 0.357568919 |
| SMART | SM00204:TGFB | 0.068108366 | 13590, | 27.1900621 | 0.357568919 |
| GOTERM_MF_DIRECT | GO:0004890~GABA-A receptor activity | 0.030878225 | 14399, 14403 | 62.5349087 | 0.370538705 |
| INTERPRO | IPR050822:Cerebellin_Synaptic_Org | 0.022420482 | 23829, 56410 | 86.251938 | 0.378657032 |
| UP_SEQ_FEATURE | DOMAIN:Homeobox | 0.005183443 | 18424, 110648, | 11.124812 | 0.396533376 |
| INTERPRO | IPR017995:Homeobox_antennapedia | 0.02610921 | 15416, 15415 | 73.9302326 | 0.396859986 |
| GOTERM_BP_DIRECT | GO:0007611~learning or memory | 0.008533084 | 12918, 13411, | 21.1971998 | 0.407691805 |
| GOTERM_CC_DIRECT | GO:0043005~neuron projection | 0.022416663 | 243634, 15567, | 6.49435764 | 0.423397126 |
| GOTERM_CC_DIRECT | GO:0043196~varicosity | 0.023306264 | 12918, 22226 | 83.1277778 | 0.423397126 |
| GOTERM_MF_DIRECT | GO:0005148~prolactin receptor binding | 0.045176974 | 14599, 19109 | 42.4344023 | 0.4539972 |
| GOTERM_MF_DIRECT | GO:0005125~cytokine activity | 0.046240456 | 13590, 14560, | 8.56848509 | 0.4539972 |
| GOTERM_CC_DIRECT | GO:1902711~GABA-A receptor complex | 0.029430942 | 14399, 14403 | 65.627193 | 0.458281814 |
| GOTERM_BP_DIRECT | GO:0030900~forebrain development | 0.010748669 | 18424, 16409, | 18.7933936 | 0.462192784 |
| INTERPRO | IPR018116:Somatotropin_CS | 0.035271685 | 14599, 19109 | 54.4749082 | 0.46418722 |

| | | | | | |
|---|---|---|---|---|---|
| INTERPRO | IPR001827:Homeobox_Antennapedia_CS | 0.038913066 | 15416, 15415 | 49.2868217 | 0.46418722 |
| INTERPRO | IPR006028:GABAA/Glycine_rcpt | 0.042541028 | 14399, 14403 | 45.0010111 | 0.46418722 |
| INTERPRO | IPR001400:Somatotropin/Prolactin | 0.051552544 | 14599, 19109 | 36.9651163 | 0.46418722 |
| INTERPRO | IPR017948:TGFb_CS | 0.058702095 | 13590, 14560 | 32.3444767 | 0.46418722 |
| INTERPRO | IPR001073:C1q_dom | 0.058702095 | 23829, 56410 | 32.3444767 | 0.46418722 |
| INTERPRO | IPR015615:TGF-beta-rel | 0.058702095 | 13590, 14560 | 32.3444767 | 0.46418722 |
| INTERPRO | IPR001839:TGF-b_C | 0.06756508 | 13590, | 27.9736015 | 0.46418722 |
| INTERPRO | IPR018000:Neurotransmitter_ion_chnl_CS | 0.074596765 | 14399, 14403 | 25.2444697 | 0.46418722 |
| INTERPRO | IPR006029:Neurotrans-gated_channel_TM | 0.076346582 | 14399, 14403 | 24.6434109 | 0.46418722 |
| INTERPRO | IPR038050:Neuro_actylchol_rec | 0.076346582 | 14399, 14403 | 24.6434109 | 0.46418722 |
| INTERPRO | IPR006201:Neur_channel | 0.076346582 | 14399, 14403 | 24.6434109 | 0.46418722 |
| INTERPRO | IPR036719:Neuro-gated_channel_TM_sf | 0.076346582 | 14399, 14403 | 24.6434109 | 0.46418722 |
| INTERPRO | IPR006202:Neur_chan_lig-bd | 0.076346582 | 14399, 14403 | 24.6434109 | 0.46418722 |
| INTERPRO | IPR036734:Neur_chan_lig-bd_sf | 0.076346582 | 14399, 14403 | 24.6434109 | 0.46418722 |
| GOTERM_BP_DIRECT | GO:0051461~positive regulation of corticotropin secretion | 0.012825691 | 12918, 22226 | 151.913265 | 0.476887649 |
| GOTERM_BP_DIRECT | GO:1902476~chloride transmembrane transport | 0.014132265 | 243634, 14399, 14403 | 16.2764213 | 0.476887649 |
| GOTERM_BP_DIRECT | GO:0061743~motor learning | 0.014417534 | 23829, 64378 | 135.034014 | 0.476887649 |
| GOTERM_CC_DIRECT | GO:0030141~secretory granule | 0.038050764 | 14599, 22044, | 9.54272959 | 0.482335409 |
| GOTERM_CC_DIRECT | GO:0042734~presynaptic membrane | 0.039825859 | 243634, 15567, | 9.30534826 | 0.482335409 |
| GOTERM_MF_DIRECT | GO:0030594~neurotransmitter receptor activity | 0.054594493 | 15564, 14403 | 34.9459784 | 0.491350437 |

| | | | | | |
|---|---|---|---|---|---|
| GOTERM_BP_DIRECT | GO:2000987~positive regulation of behavioral fear response | 0.016006862 | 12918, 22226 | 121.530612 | 0.491639342 |
| GOTERM_CC_DIRECT | GO:0030286~dynein complex | 0.047580311 | 13411, 21648 | 40.2231183 | 0.501129847 |
| GOTERM_CC_DIRECT | GO:0043083~synaptic cleft | 0.050572737 | 23829, 56410 | 37.7853535 | 0.501129847 |
| GOTERM_MF_DIRECT | GO:0045505~dynein intermediate chain binding | 0.063921043 | 13411, 21648 | 29.7040816 | 0.519605718 |
| GOTERM_MF_DIRECT | GO:0001228~DNA-binding transcription activator activity, RNA polymerase II-specific | 0.069870752 | 18424, 18012, 110648, 15415 | 4.11129158 | 0.519605718 |
| GOTERM_MF_DIRECT | GO:0003700~DNA-binding transcription factor activity | 0.072167461 | 18424, 65100, 18012, 15416 | 4.05516473 | 0.519605718 |
| UP_KW_BIOLOGICAL_PROCESS | KW-0130~Cell adhesion | 0.0517366 | 245827, 16409, | 4.44920635 | 0.526284704 |
| UP_KW_BIOLOGICAL_PROCESS | KW-0813~Transport | 0.056409267 | 243634, 17718, | 2.09271832 | 0.526284704 |
| UP_KW_BIOLOGICAL_PROCESS | KW-0524~Neurogenesis | 0.087939218 | 65100, 18012, | 5.72040816 | 0.526284704 |
| UP_KW_BIOLOGICAL_PROCESS | KW-0406~Ion transport | 0.095688128 | 243634, 52898, | 3.43900008 | 0.526284704 |
| GOTERM_MF_DIRECT | GO:1904315~transmitter-gated monoatomic ion channel activity involved in regulation of postsynaptic membrane potential | 0.080786298 | 14399, 14403 | 23.2973189 | 0.545307514 |
| INTERPRO | IPR008983:Tumour_necrosis_fac-like_dom | 0.093668057 | 23829, 56410 | 19.9042934 | 0.547597869 |
| GOTERM_BP_DIRECT | GO:0032099~negative regulation of appetite | 0.020759806 | 14526, 22226 | 93.4850863 | 0.557919794 |
| GOTERM_BP_DIRECT | GO:2000252~negative regulation of feeding behavior | 0.020759806 | 22044, 22226 | 93.4850863 | 0.557919794 |

| | | | | | |
|---|---|---|---|---|---|
| GOTERM_CC_DIRECT | GO:1902495~transmembrane transporter complex | 0.063925171 | 14399, 14403 | 29.6884921 | 0.580653634 |
| GOTERM_CC_DIRECT | GO:0045211~postsynaptic membrane | 0.070754841 | 15567, 14399, | 6.72796763 | 0.593252131 |
| GOTERM_CC_DIRECT | GO:0030425~dendrite | 0.07996172 | 15564, 14399, | 3.87843442 | 0.622559106 |
| GOTERM_CC_DIRECT | GO:0043025~neuronal cell body | 0.087319317 | 12918, 21648, | 3.73328343 | 0.634520373 |
| UP_KW_CELLULAR_C | KW-0243~Dynein | 0.071385976 | 13411, | 26.3456615 | 0.678166774 |
| GOTERM_BP_DIRECT | GO:0099558~maintenance of synapse structure | 0.027062111 | 23829, 56410 | 71.4885954 | 0.684512221 |
| UP_SEQ_FEATURE | CARBOHYD:N-linked (GlcNAc...) asparagine | 0.014128256 | 13590, 243634, 245827, | 1.98125335 | 0.720541068 |
| GOTERM_BP_DIRECT | GO:0030278~regulation of ossification | 0.034884163 | 14560, 19109 | 55.2411874 | 0.789483679 |
| GOTERM_BP_DIRECT | GO:0043950~positive regulation of cAMP-mediated signaling | 0.034884163 | 12918, 22226 | 55.2411874 | 0.789483679 |
| GOTERM_BP_DIRECT | GO:0030902~hindbrain development | 0.037995712 | 18012, 13799 | 50.6377551 | 0.816907814 |
| GOTERM_BP_DIRECT | GO:0048265~response to pain | 0.041097439 | 12918, 22226 | 46.7425432 | 0.826149129 |
| GOTERM_BP_DIRECT | GO:0051412~response to corticosterone | 0.042644628 | 12918, 22044 | 45.0113379 | 0.826149129 |
| GOTERM_BP_DIRECT | GO:0051932~synaptic transmission, GABAergic | 0.044189372 | 14399, 14403 | 43.4037901 | 0.826149129 |
| GOTERM_BP_DIRECT | GO:0021542~dentate gyrus development | 0.048808976 | 18012, 110648 | 39.2034233 | 0.869941051 |
| GOTERM_BP_DIRECT | GO:0060828~regulation of canonical Wnt signaling pathway | 0.051876559 | 14275, 14369 | 36.8274583 | 0.869941051 |
| GOTERM_BP_DIRECT | GO:0007214~gamma-aminobutyric acid signaling pathway | 0.053406716 | 14399, 14403 | 35.7442977 | 0.869941051 |
| GOTERM_BP_DIRECT | GO:0045944~positive regulation of transcription by RNA polymerase II | 0.054624206 | 18424, 18012, 110648, 22226, | 2.84393008 | 0.869941051 |

| | | | | | |
|---|---|---|---|---|---|
| GOTERM_BP_DIRECT | GO:0060078~regulation of postsynaptic membrane potential | 0.065561232 | 14399, 14403 | 28.9358601 | 0.982921281 |
| GOTERM_BP_DIRECT | GO:0007165~signal transduction | 0.066605765 | 13590, 14560, | 3.19481105 | 0.982921281 |
| GOTERM_BP_DIRECT | GO:0061564~axon development | 0.068575903 | 18012, 21648 | 27.6205937 | 0.982921281 |
| GOTERM_BP_DIRECT | GO:0008306~associative learning | 0.076070979 | 12918, 22226 | 24.8021658 | 0.993071594 |
| GOTERM_BP_DIRECT | GO:0035774~positive regulation of insulin secretion involved in cellular response to glucose stimulus | 0.08202449 | 12918, 14526 | 22.9303042 | 0.993071594 |
| GOTERM_BP_DIRECT | GO:0007188~adenylate cyclase-modulating G protein-coupled receptor signaling pathway | 0.083506988 | 64378, 14526 | 22.5056689 | 0.993071594 |
| GOTERM_BP_DIRECT | GO:0048704~embryonic skeletal system morphogenesis | 0.089413575 | 15416, 15415 | 20.9535538 | 0.993071594 |
| GOTERM_BP_DIRECT | GO:0046427~positive regulation of receptor signaling pathway via JAK-STAT | 0.092352875 | 14599, 19109 | 20.255102 | 0.993071594 |
| GOTERM_BP_DIRECT | GO:0009410~response to xenobiotic stimulus | 0.095110728 | 12918, 18012, 15567 | 5.66136392 | 0.993071594 |
| UP_SEQ_FEATURE | TOPO_DOM:Extracellular | 0.034039823 | 243634, 245827, 16409, | 2.13420263 | 1 |
| UP_SEQ_FEATURE | MOTIF:Antp-type hexapeptide | 0.043429075 | 15416, 15415 | 44.0357143 | 1 |
| UP_SEQ_FEATURE | DOMAIN:TGF-beta family profile | 0.046963944 | 13590, 14560 | 40.6483516 | 1 |
| KEGG_PATHWAY | mmu04723:Retrograde endocannabinoid signaling | 0.049557225 | 17718, 14399, 14403 | 7.99754902 | 1 |
| UP_SEQ_FEATURE | DOMAIN:C1q | 0.05749226 | 23829, | 33.0267857 | 1 |
| KEGG_PATHWAY | mmu04814:Motor proteins | 0.078031125 | 13411, 21648, | 6.17992424 | 1 |

| | | | | | |
|---|---|---|---|---|---|
| UP_SEQ_FEATURE | REGION:Disordered | 0.082940222 | 68499, 243634, 65100, 15564, 23829, 18012, 110648, 54698, | 1.21134386 | 1 |
| KEGG_PATHWAY | mmu05033:Nicotine addiction | 0.089973711 | 14399, 14403 | 20.39375 | 1 |
