## Supplemental Table 12 for "Early Life Adversity Produces Enduring Molecular and Functional Disruption of Developing Vagal Circuits"

**P120 EXCITATORY NEURONS MALES**

| Category | Term | PValue | Genes | Fold Enrichment | FDR |
|---|---|---|---|---|---|
| GOTERM_CC_DIRECT | GO:0005833~hemoglobin complex | 4.5344E-07 | 101488143, 110257, 15122, 100503605 | 248.8648649 | 2.73898E-05 |
| GOTERM_CC_DIRECT | GO:0031838~haptoglobin-hemoglobin complex | 5.76628E-07 | 101488143, 110257, 15122, 100503605 | 231.0888031 | 2.73898E-05 |
| INTERPRO | IPR050056:Hemoglobin_oxygen_transport | 6.93489E-07 | 101488143, 110257, 15122, 100503605 | 213.9711538 | 4.23138E-05 |
| INTERPRO | IPR000971:Globin | 1.35404E-06 | 101488143, 110257, 15122, 100503605 | 173.8515625 | 4.23138E-05 |
| INTERPRO | IPR009050:Globin-like_sf | 1.35404E-06 | 101488143, 110257, 15122, 100503605 | 173.8515625 | 4.23138E-05 |
| INTERPRO | IPR012292:Globin/Proto | 1.35404E-06 | 101488143, 110257, 15122, 100503605 | 173.8515625 | 4.23138E-05 |
| GOTERM_MF_DIRECT | GO:0031720~haptoglobin binding | 5.8164E-07 | 101488143, 110257, 15122, 100503605 | 229.6646943 | 6.92837E-05 |

| | | | | | |
|---|---|---|---|---|---|
| GOTERM_MF_DIRECT | GO:0005344~oxygen carrier activity | 1.1358E-06 | 101488143, 110257, 15122, 100503605 | 186.6025641 | 6.92837E-05 |
| GOTERM_MF_DIRECT | GO:0019825~oxygen binding | 5.22601E-06 | 101488143, 110257, 15122, 100503605 | 114.8323471 | 0.000212524 |
| UP_SEQ_FEATURE | DOMAIN:Globin | 1.65516E-06 | 101488143, 110257, 15122, 100503605 | 163.1911765 | 0.000249929 |
| GOTERM_CC_DIRECT | GO:0005615~extracellular space | 1.45013E-05 | 101488143, 20202, 110257, 21334, 20201, 14061, 14560, 15122, 56410, 54615, 76293, 100503605 | 4.855292511 | 0.000459207 |
| GOTERM_BP_DIRECT | GO:0098869~cellular oxidant detoxification | 1.63131E-06 | 101488143 , 20202, 20201, 15122, 100503605 | 56.97474168 | 0.000510599 |
| GOTERM_MF_DIRECT | GO:0004601~peroxidase activity | 1.81554E-05 | 101488143, 110257, 15122, 100503605 | 76.55489809 | 0.00055374 |

| | | | | | |
|---|---|---|---|---|---|
| GOTERM_MF_DIRECT | GO:0020037~heme binding | 0.000115891 | 101488143, 110257, 13078, 15122, 100503605 | 19.33705327 | 0.002607834 |
| GOTERM_MF_DIRECT | GO:0043177~organic acid binding | 0.000128254 | 101488143, 15122, 100503605 | 172.2485207 | 0.002607834 |
| UP_KW_LIGAND | KW-0349~Heme | 0.000516809 | 101488143, 110257, 13078, 15122, 100503605 | 11.79818436 | 0.006201703 |
| UP_KW_LIGAND | KW-0408~Iron | 0.0093705 | 101488143, 110257, 13078, 15122, 100503605 | 5.346518987 | 0.056222999 |
| GOTERM_CC_DIRECT | calprotectin complex | 0.002404528 | 20202, 20201 | 808.8108108 | 0.057107532 |
| KEGG_PATHWAY | mmu04080:Neuroactive ligand-receptor interaction | 0.004867546 | 21334, 14061, 54615, 14399, 14403 | 6.605263158 | 0.066523129 |
| INTERPRO | IPR002339:Hemoglobin_pi | 0.002784263 | 110257, 15122 | 695.40625 | 0.069606577 |
| GOTERM_MF_DIRECT | G protein-coupled receptor | 0.004798502 | 110257, 15122, 54615 | 28.34469328 | 0.079477296 |
| GOTERM_MF_DIRECT | GO:0031722~hemoglobin beta binding | 0.005211626 | 101488143, 100503605 | 373.2051282 | 0.079477296 |
| GOTERM_BP_DIRECT | GO:0042744~hydrogen peroxide catabolic process | 0.000630051 | 101488143, 15122, 100503605 | 78.97877984 | 0.098602966 |

| | | | | | |
|---|---|---|---|---|---|
| UP_KW_PTM | KW-0165~Cleavage on pair of basic residues | 0.009163782 | 21334, 14061, 14560, 54615 | 8.706232435 | 0.109965388 |
| KEGG_PATHWAY | mmu04657:IL-17 signaling pathway | 0.012375232 | 20202, 20201, 12608 | 16.6196944 | 0.126846133 |
| GOTERM_BP_DIRECT | GO:0043542~endothelial cell migration | 0.001322243 | 20202, 20201, 13078 | 54.53296703 | 0.137954014 |
| GOTERM_MF_DIRECT | GO:0031721~hemoglobin alpha binding | 0.010396802 | 101488143, 100503605 | 186.6025641 | 0.140934427 |
| GOTERM_MF_DIRECT | GO:0044877~protein-containing complex binding | 0.011964034 | 101488143, 22097, 15122, 14399, 100503605 | 5.440307991 | 0.143954589 |
| GOTERM_MF_DIRECT | GO:0030492~hemoglobin binding | 0.012979512 | 101488143, 100503605 | 149.2820513 | 0.143954589 |
| INTERPRO | emoglobin_a-typ | 0.006946596 | 110257, 15122 | 278.1625 | 0.144720753 |
| UP_KW_BIOLOGICAL_PROCESS | KW-0813~Transport | 0.004435476 | 101488143, 110257, 17718, 15122, 52898, 14399, 20538, 21648, 67473, 14403, 100503605 | 2.517801731 | 0.150806196 |
| GOTERM_CC_DIRECT | GO:0005576~extracellular region | 0.008796747 | 20202, 21334, 14061, 14560, 56410, 54615, 76293 | 3.737079654 | 0.1671382 |

| | | | | | |
|---|---|---|---|---|---|
| INTERPRO | IPR002337:Hemoglobin_b | 0.011092114 | 101488143, 100503605 | 173.8515625 | 0.198073465 |
| GOTERM_MF_DIRECT | GABA-A receptor | 0.024520718 | 14399, 14403 | 78.56950067 | 0.233529298 |
| GOTERM_MF_DIRECT | antioxidant activity | 0.025794934 | 20202, 20201 | 74.64102564 | 0.233529298 |
| GOTERM_MF_DIRECT | GO:0005506~iron ion binding | 0.026798444 | 110257, 13078, 15122 | 11.48323471 | 0.233529298 |
| GOTERM_BP_DIRECT | neutrophil aggregation | 0.005095446 | 20202, 20201 | 381.7307692 | 0.249044385 |
| GOTERM_BP_DIRECT | positive regulation of inflammatory | 0.006046236 | 20202, 20201, 12608 | 25.16906171 | 0.249044385 |
| GOTERM_BP_DIRECT | peptidyl-cysteine S- | 0.006365352 | 20202, 20201 | 305.3846154 | 0.249044385 |
| GOTERM_BP_DIRECT | positive regulation of peptide | 0.006365352 | 20202, 20201 | 305.3846154 | 0.249044385 |
| GOTERM_BP_DIRECT | peptide secretion | 0.006365352 | 20202, 20201 | 305.3846154 | 0.249044385 |
| KEGG_PATHWAY | etrograde endocannabinoid signaling | 0.031522555 | 17718, 14399, 14403 | 10.10216718 | 0.258484952 |
| GOTERM_BP_DIRECT | autocrine signaling | 0.010165608 | 20202, 20201 | 190.8653846 | 0.331524112 |
| GOTERM_BP_DIRECT | regulation of toll-like receptor | 0.011429212 | 20202, 20201 | 169.6581197 | 0.331524112 |
| GOTERM_BP_DIRECT | proton transmembrane transport | 0.011651007 | 17718, 52898, 67473 | 17.89362981 | 0.331524112 |
| GOTERM_CC_DIRECT | GABA-A receptor | 0.022617328 | 14399, 14403 | 85.13798009 | 0.358107691 |
| INTERPRO | 00/CaBP7/8-like_CS | 0.026149541 | 20202, 20201 | 73.20065789 | 0.408586584 |
| INTERPRO | ABAA/Glycine_rcpt | 0.031569765 | 14399, 14403 | 60.4701087 | 0.418526877 |
| INTERPRO | 00_Ca-bd_sub | 0.039645399 | 20202, 20201 | 47.95905172 | 0.418526877 |
| INTERPRO | eurotransmitter_ion_chnl_ | 0.055601613 | 14399, 14403 | 33.9222561 | 0.418526877 |
| INTERPRO | euro-gated_chann | 0.056919655 | 14399, 14403 | 33.11458333 | 0.418526877 |
| INTERPRO | IPR006201:Neur_channel | 0.056919655 | 14399, 14403 | 33.11458333 | 0.418526877 |

| | | | | | |
|---|---|---|---|---|---|
| INTERPRO | eurotrans-gated_chann | 0.056919655 | 14399, 14403 | 33.11458333 | 0.418526877 |
| INTERPRO | eur_chan_lig-bd | 0.056919655 | 14399, 14403 | 33.11458333 | 0.418526877 |
| INTERPRO | eur_chan_lig-bd_sf | 0.056919655 | 14399, 14403 | 33.11458333 | 0.418526877 |
| INTERPRO | euro_actylchol_rec | 0.056919655 | 14399, 14403 | 33.11458333 | 0.418526877 |
| UP_KW_PTM | 0027~Amidation | 0.071054061 | 21334, 54615 | 26.01976285 | 0.426324366 |
| UP_KW_CELLULAR_COMPONENT | KW-0964~Secreted | 0.02365476 | 20202, 21334, 20201, 14560, 56410, 54615, 76293 | 2.898759083 | 0.473095192 |
| KEGG_PATHWAY | icotine addiction | 0.071111749 | 14399, 14403 | 25.76052632 | 0.485930284 |
| GOTERM_CC_DIRECT | GO:0072562~blood microparticle | 0.036644872 | 101488143, 100503605 | 52.18134263 | 0.497323265 |
| GOTERM_BP_DIRECT | positive regulation of blood | 0.020230568 | 20202, 14061 | 95.43269231 | 0.51721227 |
| GOTERM_BP_DIRECT | astrocyte development | 0.02148166 | 20202, 20201 | 89.81900452 | 0.51721227 |
| GOTERM_CC_DIRECT | neuronal cell body | 0.047036883 | 14399, 20538 | 40.44054054 | 0.520720066 |
| GOTERM_CC_DIRECT | transmembrane | 0.049331375 | 14399, 14403 | 38.51480051 | 0.520720066 |
| GOTERM_MF_DIRECT | transmitter-gated monoatomic ion channel activity | 0.064501623 | 14399, 14403 | 29.27099045 | 0.524613197 |
| GOTERM_MF_DIRECT | GO:0005509~calcium ion binding | 0.070055849 | 20202, 245827, 20201, 14061 | 4.073180117 | 0.534175852 |
| ULAR_FUNCTION | 0049~Antioxidant | 0.024550528 | 20202, 20201 | 76.54375 | 0.567464218 |
| ULAR_FUNCTION | 0527~Neuropeptide | 0.036610595 | 21334, 54615 | 51.02916667 | 0.567464218 |
| GOTERM_BP_DIRECT | response to lipopolysaccharide | 0.025704638 | 20202, 20201, 12608 | 11.74556213 | 0.574682259 |

| | | | | | |
|---|---|---|---|---|---|
| GOTERM_BP_DIRECT | leukocyte migration involved in | 0.027713823 | 20202, 20201 | 69.40559441 | 0.578295098 |
| GOTERM_CC_DIRECT | chloride channel | 0.061855389 | 14399, 14403 | 30.52116267 | 0.587626195 |
| SMART | SM01394:S_100 | 0.032772115 | 20202, 20201 | 55.83673469 | 0.65544231 |
| GOTERM_BP_DIRECT | synaptic transmission, | 0.035141415 | 14399, 14403 | 54.53296703 | 0.665340141 |
| GOTERM_BP_DIRECT | positive regulation of | 0.041288845 | 21334, 54615 | 46.27039627 | 0.665340141 |
| GOTERM_BP_DIRECT | erythrocyte development | 0.041288845 | 110257, 15122 | 46.27039627 | 0.665340141 |
| GOTERM_BP_DIRECT | neuron differentiation | 0.041374919 | 15285, 104382, 12608 | 9.052903618 | 0.665340141 |
| GOTERM_BP_DIRECT | gamma-aminobutyric acid signaling | 0.042513747 | 14399, 14403 | 44.90950226 | 0.665340141 |
| GOTERM_MF_DIRECT | chloride channel | 0.093435659 | 14399, 14403 | 19.9042735 | 0.67053826 |
| GOTERM_BP_DIRECT | positive regulation of reactive | 0.046179316 | 14061, 13078 | 41.26819127 | 0.68829171 |
| GOTERM_BP_DIRECT | positive regulation of intrinsic | 0.048615434 | 20202, 20201 | 39.15187377 | 0.691665037 |
| GOTERM_CC_DIRECT | chromosome, centromeric | 0.080895772 | 21973, 20957 | 23.10888031 | 0.698645305 |
| GOTERM_BP_DIRECT | regulation of postsynaptic membrane | 0.052258255 | 14399, 14403 | 36.35531136 | 0.711166692 |
| ULAR_FUNCTION | 0869~Chloride channel | 0.077717928 | 14399, 14403 | 23.55192308 | 0.803085256 |
| GOTERM_BP_DIRECT | neutrophil chemotaxis | 0.076198991 | 20202, 20201 | 24.62779156 | 0.993761839 |
| GOTERM_BP_DIRECT | positive regulation of osteoblast | 0.096084767 | 14560, 12608 | 19.32814021 | 1 |
